# Animal-encoded nonribosomal pathway to bursatellin analogs

**DOI:** 10.1101/2024.11.12.622736

**Authors:** Aarthi Venugopalan, Eric W. Schmidt

## Abstract

The bursatellin-oxazinin family is a series of tyrosine-derived, nitrile-containing marine natural products from gastropod and bivalve molluscs. Although the first analogs were identified and associated with toxicity forty years ago, their biosynthetic origins were unknown. During an investigation of published mollusc genomes and transcriptomes, we serendipitously identified a putative bursatellin biosynthetic gene cluster (referred hereafter as the *bur-ox* pathway). Through biochemical characterization of some *bur-ox* genes, we provide evidence suggesting that bursatellin-type metabolites are produced by molluscs themselves rather than by their microbial symbionts. We show that the reductive domain from a monomodular nonribosomal peptide synthetase (NRPS) protein FmtATR performs a four-electron reduction to produce tyrosinols from tyrosine derivatives. Moreover, an aminocarboxypropyltransferase enzyme, ACT, uses *S*-adenosylmethionine (SAM) to transform tyrosinols into their phenolic homoserine ethers, which in bursatellin is further modified to the nitrile. Widespread occurrence of *bur-ox* in molluscs suggests a common biosynthetic origin for bursatellins and oxazinins as well as an important but currently unidentified physiological role for this metabolite family in molluscs inhabiting diverse ecological niches. Further, the presence of *bur-ox* pathway homologs in many culinary bivalves such as mussels and geoducks suggests that possible impacts on human consumers should be investigated. As one of the few NRPS pathways of animal origin to be characterized, *bur-ox* sheds light on underappreciated chemical and biochemical diversity in animals.

## Introduction

Bursatellin (**1a**) is a tyrosine-derived, formylated diol nitrile reported from gastropod molluscs, including the sea hare *Bursatella leachii*^1,2^ and later from nudibranchs of genera *Spurilla*^3^ and *Jorunna*.^4^ Oxazinins (**1c-1i**), originally reported from the bivalve mollusc *Mytilus galloprovincialis*, are composed of an indole moiety linked via a morpholinone ring to tyrosinol or deformylated bursatellin (Figures 1A and S1).^5–7^ Oxazinin-1 (**1c**) has recently also been found in marine snails (gastropods) *Planaxus sulcatus*^8^ and *Mauritia arabica*.^9^ These compounds have been synthesized in several studies.^10–14^ However, despite their presence across a wide taxonomic group of molluscs, a systematic investigation of the true distribution of this compound family, their biological role in molluscs, and any ecological impact has not been reported. Molluscs such as *Mytilus* spp. mussels act as environmental engineers and indicators of water quality and are also commonly consumed by people, while other molluscs produce therapeutics or lead compounds.^15,16^ Therefore, understanding the provenance of chemical components such as bursatellin and relatives is both scientifically and economically important.

**Figure 1.**
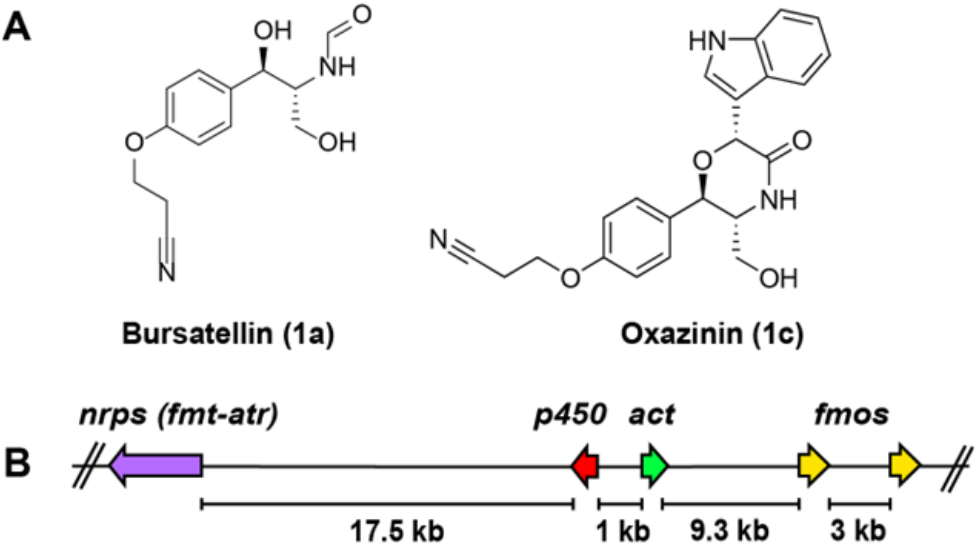
Representative structures and genes for bursatellin and oxazinin metabolites. A) Structures of the major members of each class (related compounds shown in Figure S1). B) Putative *bur-ox* BGC in the mussel *Mytilus galloprovincialis* genome. Distance between genes (in kbp) is indicated. The gene organization is conserved across sequenced *Mytilus* mussel species. *nrps*: non-ribosomal peptide synthetase; *act*: aminocarboxypropyltransferase; *fmos*: flavin monooxygenases; *p450*: cytochrome p450.

While investigating the animal kingdom for uncharacterized nonribosomal peptide synthetases (NRPSs) using NCBI datasets, we serendipitously encountered a putative NRPS-containing biosynthetic gene cluster (BGC), *bur-ox*, in the genome of the blue mussel, *Mytilus edulis*, a common food species (Figure 1B). Only a few NRPS-type genes have so far been characterized in animals,^17–22^ while for the most part NRPS genes have been studied in microbes. The scarcity of experimental data for animal NRPSs despite their widespread occurrence across the animal kingdom,^23^ drove us to characterize the orphan pathway in molluscs.

Here, we describe the discovery and characterization of *bur-ox* and its widespread occurrence across phylum Mollusca. We expressed and purified biosynthetic enzymes, employing synthetic substrates to show that the mussel NRPS catalyzes reduction of *N*-formyl-L-tyrosine (**2a**) to *N-*formyl-L-tyrosinol (**4a**), while an aminocarboxypropyltransferase (ACT) transfers homoserine from *S*-adenosylmethionine (SAM) to the tyrosinol substrate **4a** to produce a phenolic ether (**5a**). These transformations are consistent with early steps in the formation of bursatellin-type metabolites. The *bur-ox* pathway is present not just in mussels inhabiting diverse habitats but also in other bivalves such as oysters, clams, abalones, and scallops, as well as in several gastropods. While NRPS BGCs are a common feature in microbes, this is a rare instance of an NRPS being observed to cluster with tailoring enzymes in an animal genome.

## Results

### Bioinformatic identification of putative bursatellin cluster

In a campaign to discover animal NRPS genes, we assembled publicly available transcriptome read data from NCBI for mussels of genus *Mytilus* (Table S1) and encountered an NRPS gene (*fmt-atr*) in the transcriptomes. This gene was distinct from the few characterized animal NRPSs and the NRPS-like molluscan glycine-betaine reductase^24^ homolog in its domain organization. To understand the genomic context, we extracted the relevant genomic contigs/scaffolds and annotated them using fungal antiSMASH.^25^ Surprisingly, *fmtatr* was annotated as being part of a putative BGC with a few other genes, including those encoding a methyltransferase domain-containing enzyme (*act*), a cytochrome p450 (*p450*), and two flavin monoxygenases (*fmos*) (Figures 1B and S2). Unlike in bacteria, fungi, and some plants, BGCs have only very rarely been described in animals, making it difficult to identify all the genes involved in a biosynthetic pathway.

Analysis of *fmt-atr* and its protein sequence using antiSMASH and InterProScan^26^ revealed that the gene had an N*-*terminal formylation domain (Fmt), a tyrosine-activating adenylation domain (A), a thiolation (peptidyl carrier protein) domain (T or PCP), and a NAD-binding reductase domain (R) (Figure S2). Based upon this architecture, we predicted that the product of FmtATR enzyme should be *N*-formyl-L-tyrosinol (**4a**). A SciFinder (https://scifinder.cas.org) search for formyl-tyrosinol-type metabolites turned up reports on bursatellins and oxazinins from several molluscs, leading to the hypothesis that FmtATR might underlie their biosyntheses.

From the structure of bursatellin, we expected that the biosynthetic pathway would involve an atypical methyltransferase-like enzyme that employs SAM to catalyze phenol aminocarboxypropylation. A BLAST analysis of the clustered methyltransferase domain-containing protein revealed that the methyltransferase was an isonocardicin synthase homolog (Figure S3A). Isonocardicin synthase is an ACT that catalyzes the formation of nocardicin A from nocardicin E in bacteria.^27^ ACTs are rare in nature with few characterized examples besides isonocardicin synthase such as MccD in the microcin gene cluster, nicotianamine synthase (in plants) and its bacterial homolog CntL in the staphylopine gene cluster, BtaA (SAM-diacylglycerol 3-amino-3-carboxypropyl transferase) in microalgal betaine lipid biosynthesis, and TYW2, a tRNA-modifying enzyme found widely in the tree of life.^28,29^ Their activity cannot be easily predicted because of lack of pairwise sequence identity between the few biochemically characterized enzymes and lack of structural information.^30^ The bivalve gene, however, is annotated as ‘isonocardicin synthase-like’ by the NCBI eukaryotic genome annotation pipeline. Superimposition of *ab initio* models of isonocardicin synthase and the bivalve transferase suggested that the two enzymes share a conserved fold (Figure S3B).

A cytochrome p450 is present in proximity to the ACT and is predicted to be globular unlike typical animal cytochrome p450s that have an N*-*terminal transmembrane domain. A BLAST analysis of the bivalve p450 against the NCBI nr database produces hits from *Streptomyces* when molluscan hits are excluded from the search (Figure S4). This suggests that, similar to the NRPS and ACT, the cytochrome p450 is also bacteria-like. The two Fmos, on the other hand, are homologous to other animal Fmos. We predict that the cytochrome p450 and Fmos are involved in installing the hydroxyl and nitrile groups, respectively, in bursatellin, although their exact roles can only be determined through rigorous biochemical characterization. Based upon this analysis, we predicted a potential biosynthetic pathway to bursatellin (Figure 2).

**Figure 2.**
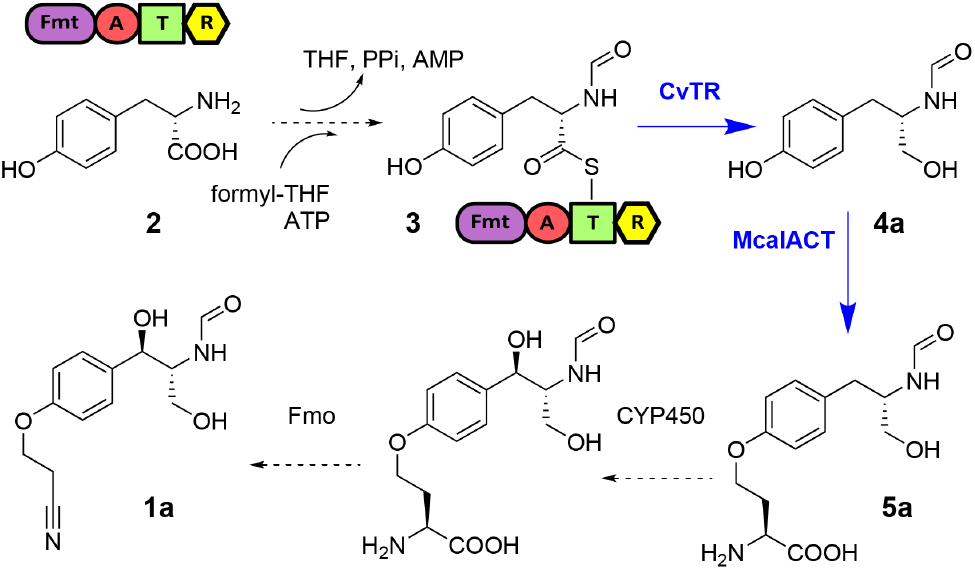
Proposed biosynthesis of bursatellin based upon *bur-ox*. Dashed arrows indicate putative enzymatic reactions; Solid/blue arrows indicate experimentally validated biosynthetic steps in this study. THF: tetrahydrofolate; PPi: pyrophosphate; Fmt: formyltransferase. NRPS domains: A: adenylation; T: thiolation; R: reductase.

A search for the *bur-ox* pathway genes in other bivalve genomes and transcriptomes derived from NCBI datasets revealed that a partial/full machinery is present in most of the published mussel, oyster, clam, scallop, cockle and pen shell genomes (Figure 3 and Table S2). Among gastropods, evidence for the genes is seen in vetigastropods and in some caenogastropod and heterobranch genomes (Figure 3 and Table S3), but the distribution is not as clear as in bivalve genomes. Some patellogastropod genomes, for example, harbor the ACT homolog but not the NRPS gene. This might be due to a stark contrast in the number and quality of published genomes for bivalves versus gastropods, since larger genes are often challenging to assemble and sometimes cannot be identified in lower quality genomes and transcriptomes despite their presence in an organism.

**Figure 3.**
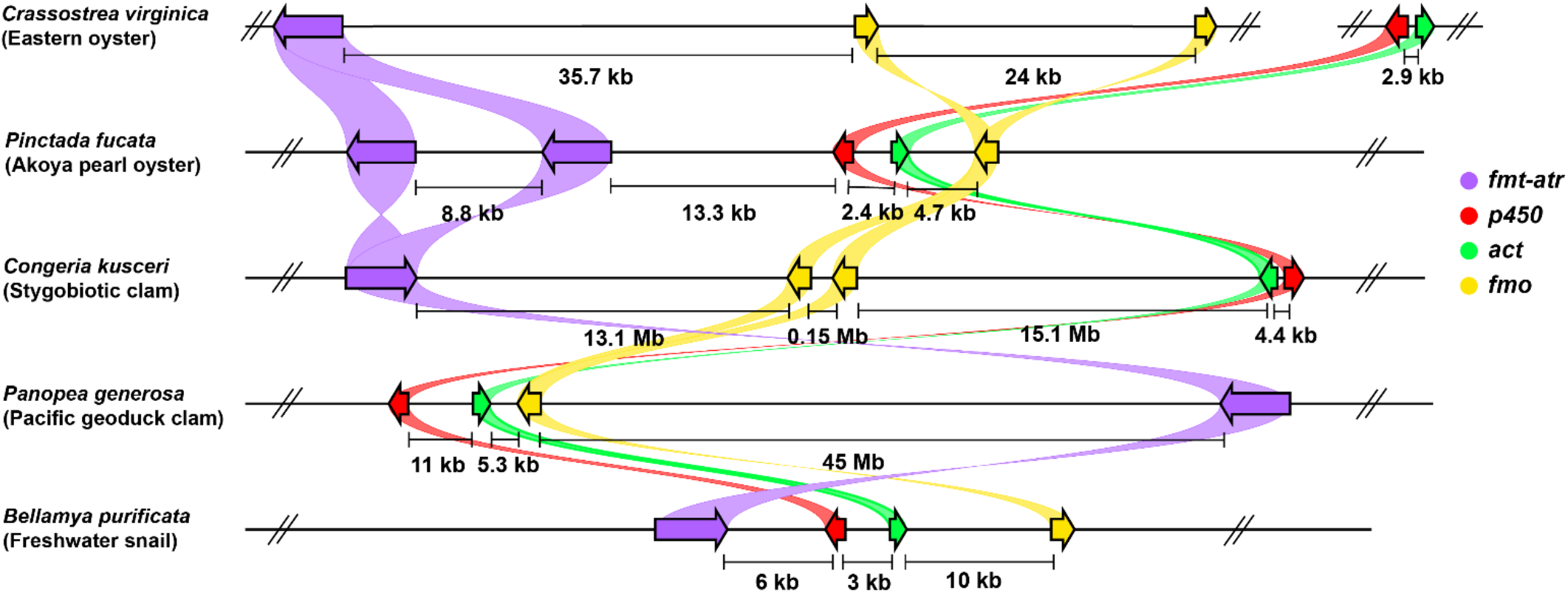
Distribution of the putative *bur-ox* pathway genes in a few representative bivalve and gastropod genomes. The figure is not drawn to scale, but the distance between genes (in kbp) is indicated.

### Chemical and biochemical strategy for enzyme characterization

To determine whether *bur-ox* was plausibly involved in bursatellin and oxazinin biosynthesis, key enzymes were expressed and used in reactions with synthetic, fully characterized substrates. We synthesized 7 substrate mimics of covalently tethered thioester **3**, consisting of coenzyme A (CoA) and *N*-acetylcysteamine (SNAC) thioesters, as well as 5 substrate analogs of tyrosinol compound **4a** (see Scheme S1). Four commercial compounds were also used in enzyme assays. In all enzymatic reduction reactions described below, unless otherwise stated these well-characterized and purified compounds were used as substrates and as standards to validate product formation. We also enzymatically synthesized 5 homoserine ethers, analogs of compound **5**. One of these (**5b\***) was fully characterized by NMR spectroscopy, while the remainder were analyzed based upon MS and diagnostic MS^2^ fragments shared with **5b** (see Scheme S2). Using these characterized compounds, enzymatic reactions could be performed using robust controls.

### Tyrosinols synthesized by the bivalve NRPS reductase construct (CvTR)

In our attempts at biochemical characterization of key *bur-ox* genes, we faced failure of cloning and soluble expression, respectively, with the NRPS genes from *Mytilus edulis* (*Mefmt-atr*) and *Mytilus californianus* (*Mcalfmt-atr*). Similarly, expression of codon optimized, excised T-R and R domains with or without solubility tags in *E. coli* produced very little soluble protein with abundant chaperone copurification suggesting misfolding of target proteins. Consequently, assays with purified proteins did not show any sign of enzyme activity.

Some FmtATR homologs in bivalves such as oysters showed a greater predicted hydrophilicity, which predicts greater solubility. We assembled the full NRPS gene from *Crassostrea virginica* (*Cvfmt-atr*) and expressed it in *Saccharomyces cerevisiae*. An expected ∼180 kDa protein was observed by SDS-PAGE, but it was largely in the aggregated form (Figure S5A). Assaying the purified protein with L-tyrosine, ATP, NADPH, and folinic acid did not provide reproducible results, potentially reflecting poor protein stability. Screening CvATR, a construct lacking the N*-*terminal formylation domain, also produced low levels of soluble but misfolded protein (Figure S5B). Finally, a T-R construct was generated from the same gene (CvTR), which showed a high level of expression, solubility, and folding (Figure S6).

Pure CvTR was used as a surrogate for the full NRPS with suitable substrates in assays. Assay results with various substrates are summarized as a qualitative heatmap in Scheme S3. We chemically synthesized a series of tyrosyl CoA (**3Ca**-**3Cc**) and SNAC (**3Sa**-**3Sd**) substrates, as well as predicted reduction products, and characterized them spectroscopically. The goal was to investigate whether CvTR catalyzed the reduction of the native substrate in thioester form as hypothesized and whether it demonstrated any tolerance towards substrate analogs. Enzyme reactions were monitored by high resolution mass spectrometry. All enzymatic reaction products were verified by comparison and coelution with well characterized synthetic standards (Figures S48-S52). CoA esters were tethered to the T-domain using the phosphopantetheinyl transferase, Sfp (R4-4 mutant).^31^ Incubation of CvTR with *N*-formyl-L-tyrosine CoA (**3Ca**) in the presence of Sfp and NADPH produced a peak with the expected mass of *N*-formyl-L-tyrosinol **4a** (*m/z* 218.0788 [M+Na^+^]) (Figures 4D and S8C). No product peak was seen in controls lacking either CvTR or NADPH (Figures 4A-B and S8A-B). CvTR also accepted the much bulkier CoA substrates, *N*-Boc-L-tyrosine CoA (**3Cb**) and *N*-acetyl-*O-*tert-butyl tyrosine CoA (**3Cc**). Peaks matching *N*-Boc-L-tyrosinol (*m/z* 290.1363 [M+Na^+^]) (**4b**) and *N*-acetyl-*O-*tert-butyl-L-tyrosinol (*m/z* 288.1571 [M+Na^+^]) (**4c\***) were present in the respective chromatograms (Figures 5A-B and S9-S14). Controls lacking Sfp still showed product formation (Figures S11C, S12C, and S23B). This prompted us to investigate if thioester mimics such as aminoacyl SNAC substrates are also reduced by CvTR.

**Figure 4.**
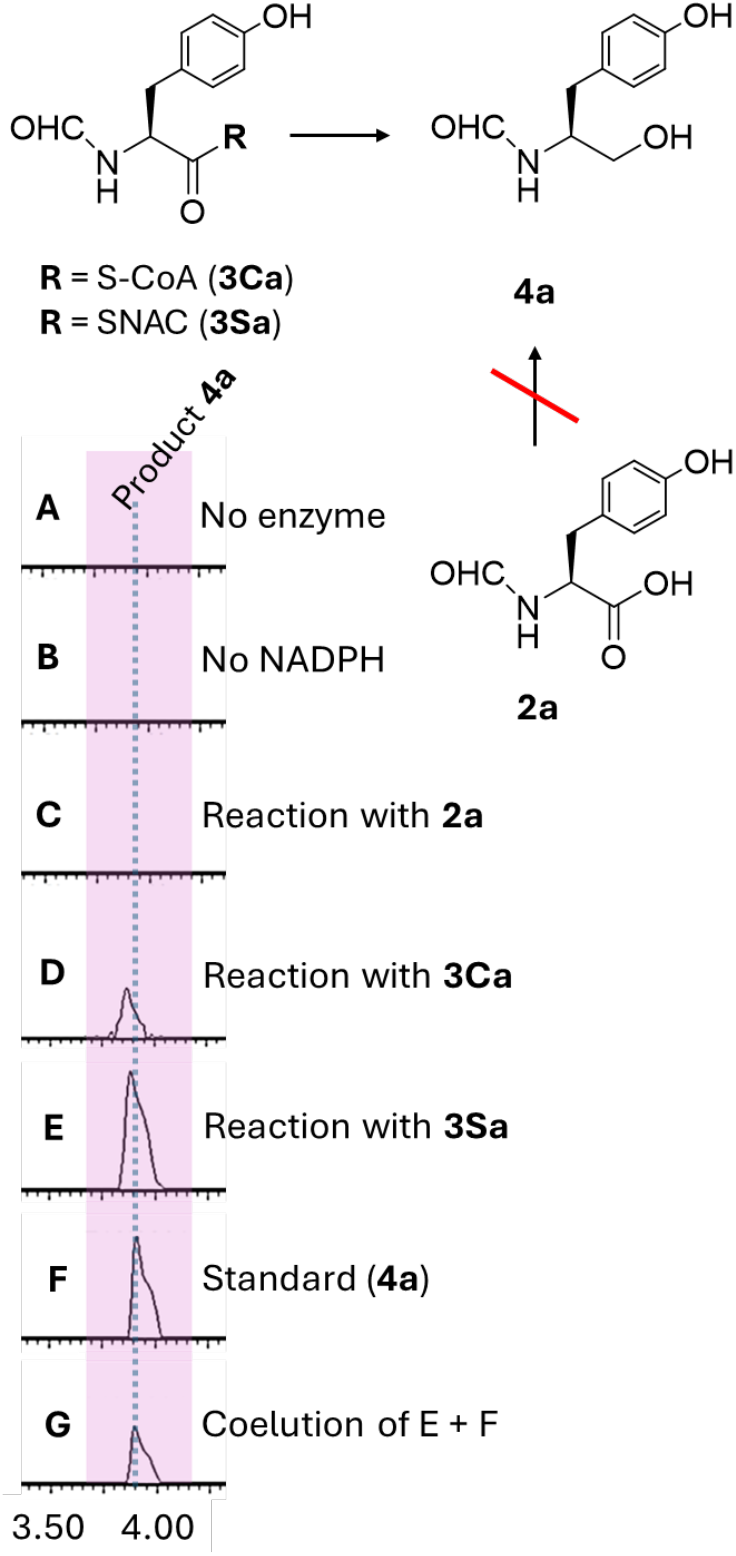
CvTR-catalyzed reduction of *N*-formyl-L-tyrosyl-CoA (**3Ca**) and -SNAC (**3Sa**) produces *N*-formyl-L-tyrosinol (**4a**). Reactions were run as described in methods section 1.6. Shown are UPLC-MS ion chromatograms extracted with the mass of product **4a** (*m/z* 218.078). A) No enzyme control containing substrate **3Ca** and all reagents except for enzyme CvTR. B) No NADPH control containing substrate **3Ca** and all reaction components except for reductant. C) No thioester control using substrate **2a** (*N*formyl-L-tyrosine) and all reaction components. D) Reaction mixture containing substrate **3Ca**, all reagents, and Sfp. E) Reaction mixture containing substrate **3Sa** and all reagents. F) Synthetic standard of **4a**. G) Coelution of samples shown in panels E + F.

**Figure 5.**
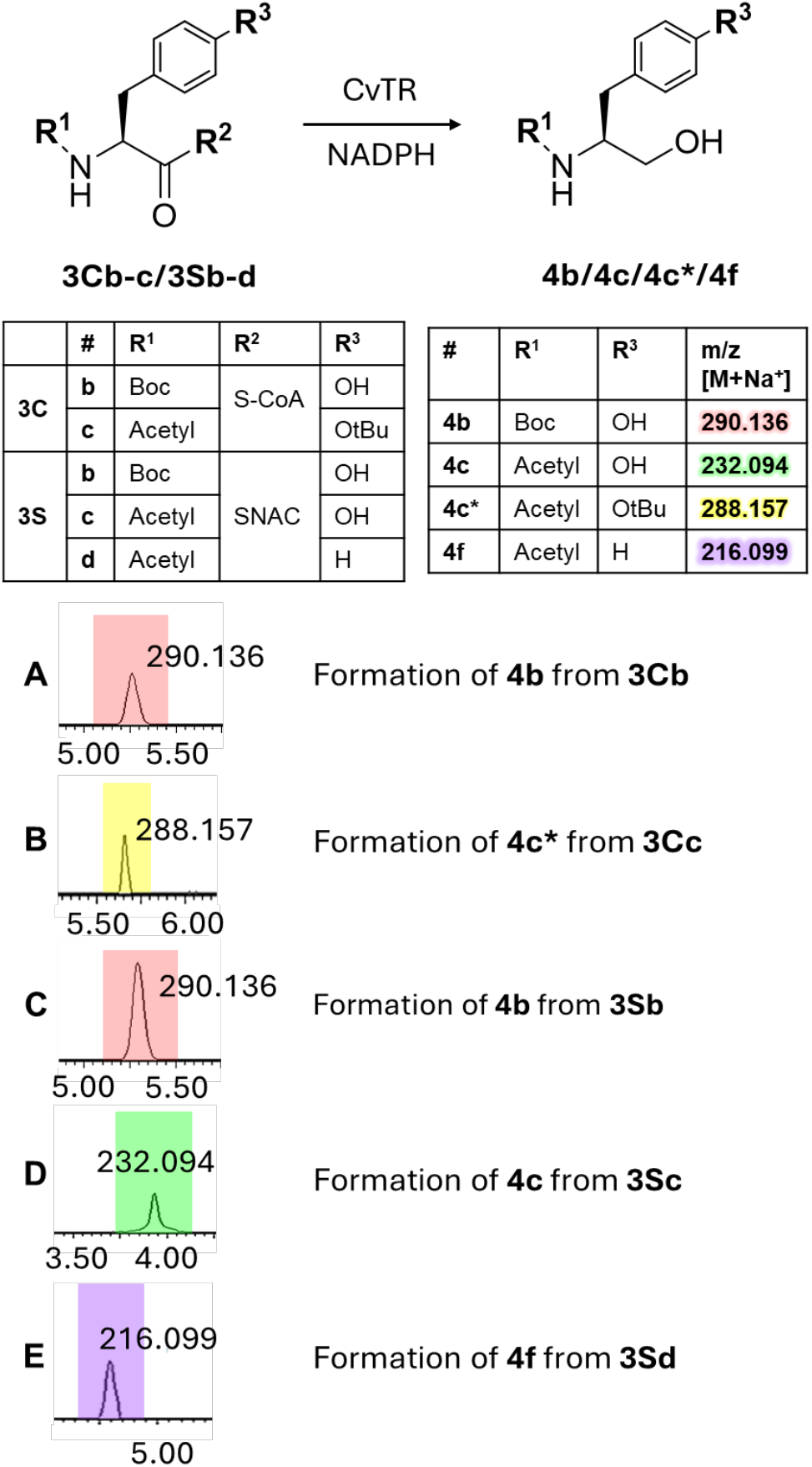
CvTR accepts thioester substrates with diverse *N*- and *O*-modifying groups. Reactions were run as described in methods section 1.6. Shown are UPLC-MS ion chromatograms extracted with the mass of product for reaction mixture containing all reagents, Sfp, and CoA substrate **3Cb** (A) or **3Cc** (B) and reaction mixture containing all reagents and SNAC substrate **3Sb** (C), **3Sc** (D), or **3Sd** (E).

Unlike CoA substrates that are covalently tethered to the T-domain by a phosphopantetheinyl arm, the freely diffusible SNAC substrates often allow for multiple turnovers while efficiently mimicking aminoacyl-S-PCP.^32^ Incubation of CvTR with *N*-formyl-L-tyrosine SNAC (**3Sa**) and NADPH produced a peak with the expected mass of **4a** (*m/z* 218.0788 [M+Na^+^]) (Figures 4E, S15D, and S16D), while no product peak was seen in controls lacking either enzyme or NADPH (Figures S15B-C and S16B-C) or in reactions of CvTR with *N-*formyl-L-tyrosine (**2a**) and NADPH (Figures 4C, S15A, and S16A). In reactions with other SNAC substrates, i.e. *N-*Boc-L-tyrosine SNAC (**3Sb**), *N-*acetyl-L-tyrosine SNAC (**3Sc**), and *N-*acetyl-L-phenylalanine SNAC (**3Sd**), product peaks matching standards of **4b**, *N*-acetyl-L-tyrosinol (*m/z* 232.0945 [M+Na^+^]) (**4c**), and *N*-acetyl-L-phenylalaninol (*m/z* 216.0995 [M+Na^+^]) (**4f**) were present in the respective chromatograms (Figures 5C-E, S17-S20, S11E, and S12E).

We used *N-*Boc-L-tyrosine thioesters (**3Cb** and **3Sb**) as substrates for further experiments because the reduction product **4b** was easier to monitor chromatographically than the more polar acyl tyrosinols. We probed whether CvTR could utilize NADH as an alternative reductant in place of NADPH by incubating **3Cb** and **3Sb** in mixtures containing CvTR and NADH. The formation of **4b** was readily observed in these conditions, indicating that CvTR tolerates NADH as an alternative reductant (Figures S21-S22). We compared the yield of **4b** from 100 µM **3Cb** and **3Sb** after 20 h in the presence of either NADPH or NADH by monitoring UV absorption at 280 nm. While CvTR showed a slight preference for NADPH over NADH with the CoA substrate **3Cb**, almost no conversion of the SNAC substrate **3Sb** was observed with NADH as the reductant (Figures 6 and S24).

**Figure 6.**
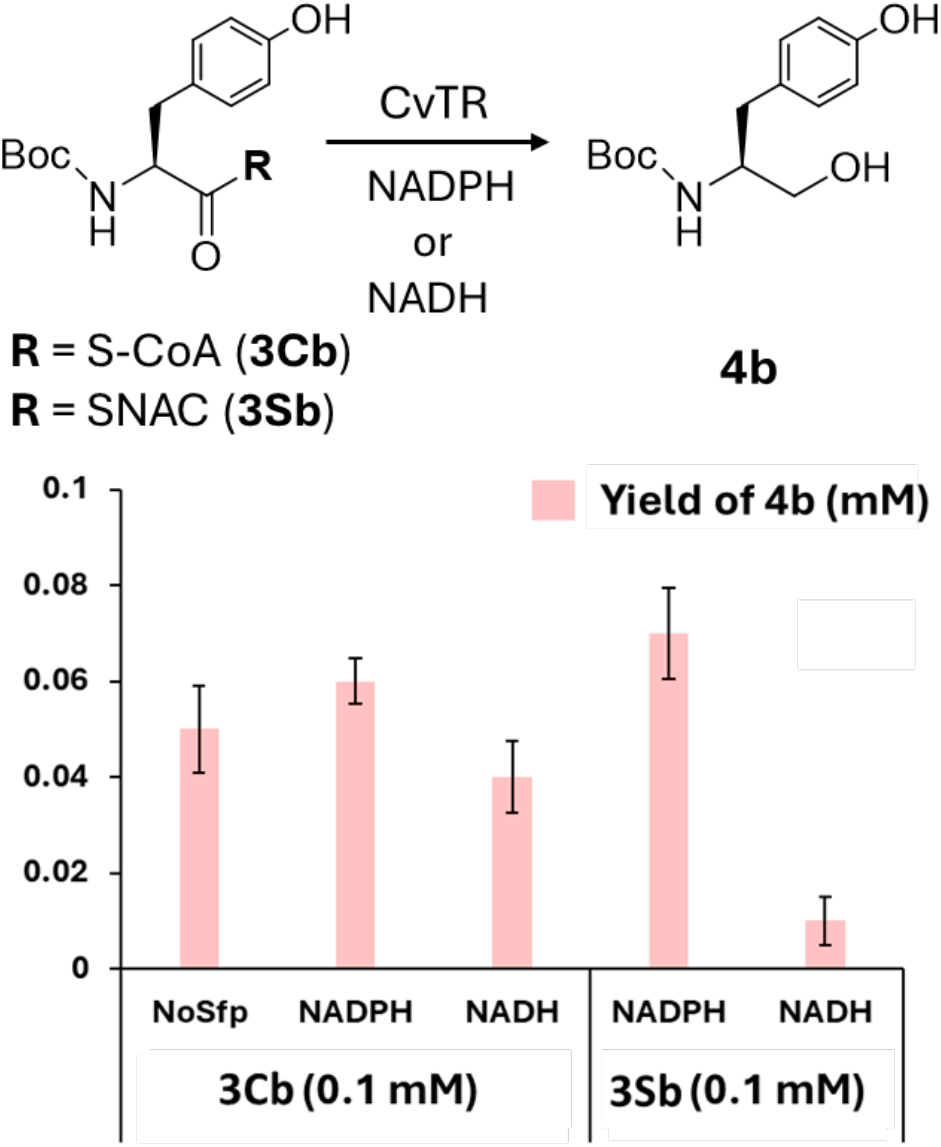
Yield estimate for CvTR catalyzed conversion of 0.1 mM **3Cb** and **3Sb** with NADPH or NADH as reductant. The y-axis shows yield of compound **4b** (in mM).

In summary, CvTR catalyzed the four-electron reduction of several aromatic thioester substrates but not free amino acids, consistent with its hypothesized role in bursatellin biosynthesis but suggesting that amino acid substrate selectivity might be imposed by the adenylation domain. Use of NADH as the sole reductant has been reported with NRPS/PKS-associated thioester reductases such as CpkC-TR in coelimycin P1 biosynthesis^33^ and the *Tg*TR domain from *Toxoplasma gondii Tg*PKS2^34^ while it is a less preferred alternative to NADPH in some bacterial carboxylic acid reductase (CAR) catalysis.^35^ Our demonstration of CvTR catalysis using NADH as an alternative reductant is the first example for such activity from an animal NRPS reductase domain.

### McalACT catalyzes tyrosyl aminocarboxypropylation

Recombinant McalACT from *M. californianus* showed soluble expression in *E. coli* and eluted at the expected monomeric elution volume (Figure S7). Results of assaying McalACT with various substrates are summarized as a qualitative heatmap in Scheme S4. Whether aminocarboxypropylation occurs on T-domain-tethered substrates or after product offloading by the reductase domain cannot be predicted bioinformatically. In the event, incubation of McalACT with **4a**, SAM, and MgCl_2_ and analysis by UPLC-MS led to a new peak with *m/z* 297.1445 in the reaction but not in controls (Figures 7A-C and S25-S26). The new peak was shifted in mass by 101 Da from that of **4a** (*m/z* 196.0969 [M+H^+^]), consistent with addition of an aminocarboxypropyl group. HR-MS/MS fragmentation data for the product, *N*-formyl-L-tyrosinol *O*-homoserine ether (**5a**), revealed key fragments corresponding to the transferred aminocarboxypropyl group (*m/z* 102.0546) and tyrosinol backbone (*m/z* 107.0485) as well as other conserved fragmentation products (Figure 7D-E and SI section 1.8).

**Figure 7.**
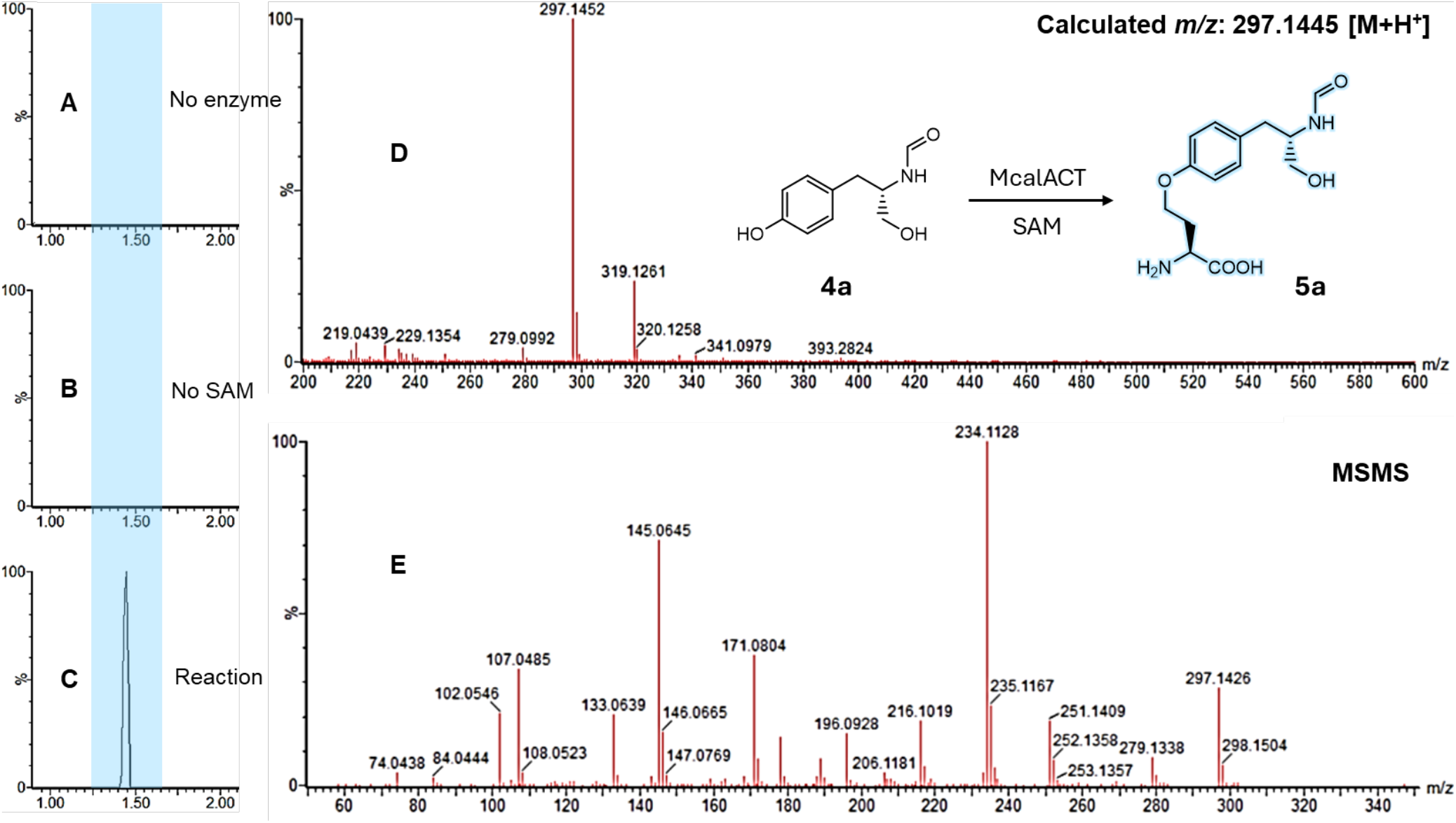
McalACT catalyzes aminocarboxypropylation of *N*-formyl-L-tyrosinol (**4a**) in vitro. Reactions were run as described in methods section 1.8. Shown are UPLC-MS ion chromatograms extracted with the mass of product **5a** (*m/z* 297.1445). A) No enzyme control containing substrate **4a** and all reagents except for enzyme McalACT. B) No SAM control containing substrate **4a** and all reaction components except for SAM. C) Reaction mixture containing substrate **4a** and all reagents. D) MS spectrum of reaction product (**5a**). E) MS/MS spectrum of the precursor ion (*m/z* 297.1452) from D shows key diagnostic fragments (*m/z* 102.0546, 107.0485, 133.0693, and 145.0645).

Product **5a** was challenging to separate from various SAM byproducts. The assay was therefore run in the presence of less polar *N*-protected tyrosinols (**4b-4e**) to investigate substrate tolerance. Boc-protected substrate **4b** was aminocarboxypropylated (Figures 8B and S33-S34), providing a similar MS^2^ fragmentation as found for **5a** (Figure S37 panels D1 and D2). As it was the easiest product to purify, scale up and purification yielded *N*-Boc-L-tyrosinol *O*-homoserine ether (**5b**), a compound that readily lost Boc upon drying generating L-tyrosinol *O*-homoserine ether (**5b\***). This latter compound was characterized by NMR, definitively identifying the product as the *O*-aminocarboxypropylated phenol derivative (Figures S46-47). Other *N-*acyl tyrosinols (**4c-e**) were also accepted as substrates by McalACT and transformed into the corresponding *O-*homoserine ethers **5c**-**5e** (Figures 8A-C and S27-S32). The identities of the various homoserine ethers were confirmed by MS^2^ analysis which supported the assignment of modifications on the phenol group (Figure S37). No measurable product was observed when **4f** (Figures 8A and S35-S36) was used as the substrate indicating that the phenolic hydroxyl group is required. Thus, McalACT modifies the phenolic hydroxyl group at a biosynthetic step after tyrosine activation, consistent with its role en route to bursatellin analogs.

**Figure 8.**
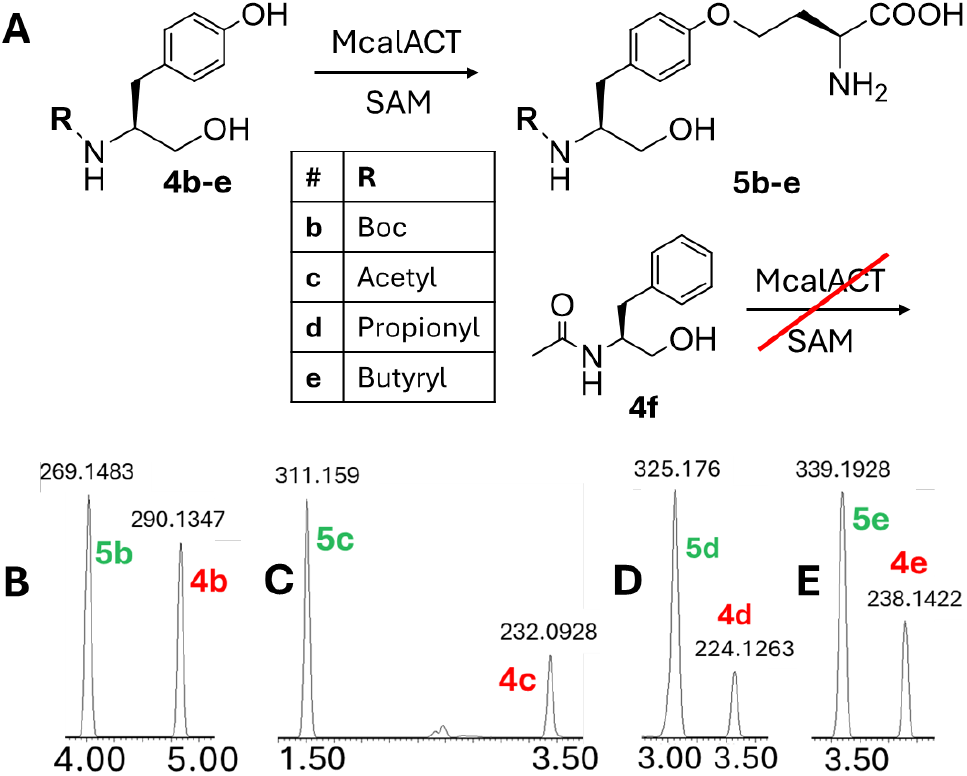
McalACT accepts various *N*-acyl tyrosinols (**4b-e**) as substrates. Reactions were run as described in methods section 1.8. A) Schematic summary of reaction results. UPLC-MS ion chromatograms extracted with respective product mass for reaction mixture containing all reagents and B) substrate **4b**, C) substrate **4c**, D) substrate **4d**, or E) substrate **4e**.

Prior to estimating kinetic parameters for McalACT-catalyzed formation of **5b**, conditions were screened at various incubation temperatures and concentrations of enzyme and SAM (Figure S38). Near complete conversion of **4b** (0.5 mM) was observed in 50 mM NaH_2_PO_4_, 150 mM NaCl, pH 8.0 buffer at 25 °C after 20+ h with 20 µM McalACT, 10 mM MgCl_2,_ and 10-fold excess SAM (5 mM) (Figure S38E and S39A). Thus, reactions containing **4b** (0.2-1 mM), 10 µM McalACT, 10 mM MgCl_2_, and 10-fold excess SAM (5 mM) at 25 °C were used for initial rate experiments. K_m_ and V_max_ values for McalACT with **4b** as substrate were estimated to be 0.16 mM and 0.03 mM h^-1^, respectively (Figure 9). The calculated k_cat_ was 3 h^-1^.

**Figure 9.**
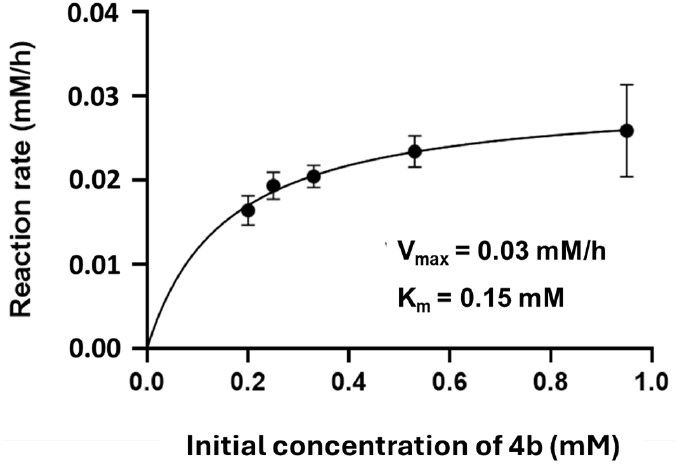
Michaelis-Menten plot for McalACT-catalyzed aminocarboxylpropylation of **4b**.

### One-pot generation of *N-*protected tyrosinol *O*-homoserine ethers from *N-*protected tyrosyl thioesters

We tested thioesters **3Sa, 3Sb**, and **3Cb** as substrates in one-pot reactions with CvTR and McalACT in the presence of SAM and NADPH. Parallel reactions containing only McalACT, SAM, and respective substrate were used for comparison. Results of these assays are summarized in Figure 10A and Scheme S5. When **3Sa** (or **3Sb**) was incubated with only McalACT, a new peak with *m/z* 412.1553 (or 484.2141) ([M+H^+^]) shifted by 101 Da mass from **3Sa** (*m/z* 311.1061 [M+H^+^]) (or **3Sb** (383.1636 [M+H^+^])) appeared (Figures 10B1 and 10C1; S40B, S41B, S42B, and S43B). In one-pot reactions containing **3Sa** (or **3Sb**), McalACT, and CvTR, an additional peak matching **5a** (or **5b**) was observed after 20 h (Figures 10B2 and 10C2; S40D, S41D, S42C, and S43C). Thus, McalACT aminocarboxypropylated both **3Sa** (or **3Sb**) producing **8Sa** (or **8Sb**) and the CvTR-generated **4a** (or **4b**). In contrast, in reactions of **3Cb** with McalACT, only the CoA hydrolysis byproduct, *N-*Boc-L-tyrosine (**2b**) (*m/z* 304.1156 [M+Na^+^]) was observed, suggesting no measurable enzyme activity after 20 h (Figure 10D1, S44A, and S45A). In one-pot reactions including CvTR, an additional peak matching **5b** (Figures 10D2, S44B, and S45B) was seen, recapitulating the expected sequence of CvTR-catalyzed reduction of CoA thioester followed by McalACT-catalyzed aminocarboxypropylation as is predicted for bursatellin biosynthesis. These experiments revealed that in addition to modifying tyrosinols, McalACT shows activity towards SNAC thioesters and to a much lesser extent towards tyrosines (Figure S42A and S43A). McalACT may therefore find use as a broad catalyst for this unusual reaction.

**Figure 10.**
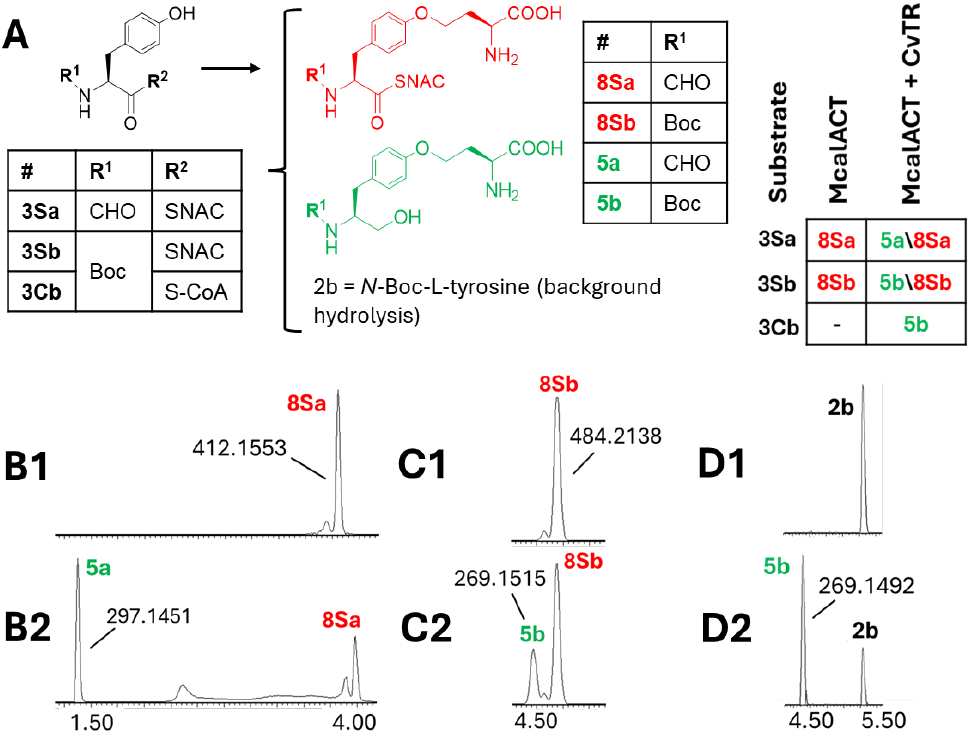
One-pot reactions with McalACT and CvTR. Reactions were run as described in methods section 1.10. A) Schematic summary of reaction results. UPLC-MS ion chromatograms extracted with respective product masses for reaction mixture containing B1) **3Sa**, McalACT, SAM, and MgCl_2_. B2) **3Sa**, CvTR, McalACT, NADPH, SAM, and MgCl_2_. C1) **3Sb**, McalACT, SAM, and MgCl_2_. C2) **3Sb**, CvTR, McalACT, NADPH, SAM, and MgCl_2_. D1) **3Cb**, McalACT, SAM, and MgCl_2_. D2) **3Cb**, CvTR, McalACT, NADPH, SAM, and MgCl_2_.

## Discussion

In the present study, we identified a pattern repeated in numerous bivalve and gastropod mollusc genomes of a set of genes clustered together. Homologs were identified in diverse molluscs, such as hydrothermal vent organisms (*Alviniconcha* and *Bathymodiolous*), stygiobiotic clams (*Congeria*), the blacktip shipworm (*Lyrodus pedicellatus*), the California sea hare (*Aplysia californica*), the giant clam (*Tridacna gigas*), and many food animals such as *Mytilus* mussels, geoducks (*Panopea*), and abalones (*Haliotis*). A retrobiosynthetic analysis led to the prediction that the genes represented a part of the bursatellin/oxazinin biosynthetic pathway. Expression of a T-R didomain construct from the NRPS (CvTR) demonstrated four-electron reduction of *N-*formyl-L-tyrosine thioester and analogs to the corresponding alcohols, consistent with bursatellin production, while biochemical characterization confirmed the activity of the McalACT enzyme on the phenolic hydroxyl group of tyrosinols. In addition, the oxidase genes in the cluster are plausibly involved in tyrosinol beta-oxidation and/or oxidation of the aminocarboxypropyl group to the nitrile. These data lead us to propose that gastropods and bivalves synthesize bursatellin/oxazinin using enzymes encoded in the *bur-ox* locus.

Outside of invertebrates such as molluscs, the closest homologs of individual *bur*-*ox* genes are found in bacteria (with the exception of the animal-derived Fmos), suggesting possible ancient horizontal gene transfer (HGT). The *bur*-*ox* proteins have several microbe-like features. A McalACT-like protein was characterized in the biosynthesis of the bacterial antibiotics, isonocardicins.^27^ Rare bacterial NRPS proteins harbor a formylation domain,^36^ while in bacteria and fungi, NRPS-like CARs catalyze the reduction of carboxylic acids to aldehydes.^37,38^ Despite some bacterial similarities, *bur*-*ox* clearly represents a true animal biosynthetic pathway. The pathway contains animal introns and animal-derived *fmo* genes and is widely distributed across phylum Mollusca. If *bur*-*ox* originated through HGT, it likely predated the divergence of molluscan classes. Indeed, eukaryotic monomodular NRPS-like proteins such as glycine-betaine reductases ^24^ and ACSF4-U26 proteins^17^ are encoded in mollusc genomes, while the FmtATR protein reported here represents a third class of NRPS proteins in molluscs, and the first to be characterized.

Further, the widespread distribution of *bur*-*ox* in molluscs hints at a fundamental role for the metabolites in invertebrate biology. In previous work, oxazinins were modestly toxic to a human cell line and were originally suspected to be toxins because of isolation from mussels that also contained more conventional shellfish toxins.^39^ Unlike true shellfish toxins that originate from microbes, bursatellin-derived metabolites are widespread molecules made by the animals themselves. A recent study reported that erythro-bursatellin B exhibited antibacterial activity in vitro and hypothesized that bursatellins may function as sunscreens in *B. leachii*.^2^ However, many molluscs harboring the *bur-ox* pathway live in dark environments, negating the sunscreen hypothesis. Based upon their widespread occurrence and structural similarity to compounds that act on neurons,^40^ G protein-coupled receptors, and ion channels, we propose that bursatellins and oxazinins might act as hormones, neurotransmitters, or pheromones. Our description of biosynthetic genes will enable rigorous approaches to understand the physiological and ecological functions of these widespread metabolites that are commonly found in human food sources.

The *bur-ox* pathway also revealed several features with potential biotechnological uses. For example, the clean production of alcohols from thioesters using relatively diverse substrates might have application in synthetic biology. More compelling is the possibility of applying this system to promiscuously install reactive nitriles on tyrosine sidechains, which would be useful in chemical biology applications. Further work is required to realize these technological goals.

This study has several limitations. As described above, we do not yet know the biological functions of the *bur-ox* enzyme products, a factor that we hope is aided by the gene descriptions and biochemical tools reported here. In addition, here we are dealing with a non-model system, for which little experimental work has been done and genetic tools are sparse. In our hands, it is still difficult to functionally express some classes of proteins from these sources, and our lab has an overall success rate of ∼10%. New technologies are urgently needed to fully capture and evaluate the biosynthetic diversity of the animal kingdom.

The burgeoning field of animal-derived biosynthetic pathways is leading to important advances in understanding basic animal biology and ecology and in providing new compounds and enzymes for technological applications. Here, we describe one of the few animal NRPS-like pathways to be characterized, revealing potentially useful enzymes and providing tools to study widespread molluscan natural products that are found in human food sources but are poorly understood.

## ACKNOWLEDGMENT

We are grateful to Jack Skalicky and the HSC NMR Core Facility for aid with running NMR spectra, and to Hsiaonung Chen (University of Utah Department of Chemistry Mass Spectrometry Core Facility) for MS^2^ data associated with *N-*acyl tyrosinols and homoserine ethers. We thank Nancy DaSilva and Yi Tang for permission to use strain BJ5464 and plasmid pxw55, respectively. We also thank Heidi Schubert and Chris Hill for providing access to their microfluidizer. This project was funded by NIH R35GM148283.

## ABBREVIATIONS

NRPS: nonribosomal peptide synthetase
ACT: aminocarboxypropyltransferase
SAM: S-adenosyl methionine
BGC: biosynthetic gene cluster
NCBI: National Centre for Biotechnology Information
antiSMASH: antibiotics and secondary metabolite analysis shell
p450/*p450*: cytochrome p450 monoxygenase
fmo/Fmo: flavin monooxygenase
InterPro: Integrative Protein Signature Database
Fmt/*fmt*-domain: formylation domain
A-domain: adenylation domain
T/PCP-domain: thiolation/peptidyl carrier protein domain
R-domain: reductase domain
BLAST: Basic Local Alignment Search Tool
tRNA: transfer ribonucleic acid
CoA: coenzyme A
SNAC: *N*-acetyl cysteamine
NMR: nuclear magnetic resonance
SDS-PAGE: sodium dodecyl sulfate-polyacrylamide gel electrophoresis
UPLC: ultra-performance liquid chromatography
MS: mass spectrometry
HR-MS/MS: high-resolution tandem mass spectrometry
ATP: adenosine triphosphate
NADPH: nicotinamide adenine dinucleotide phosphate hydrogen
NADH: nicotinamide adenine dinucleotide hydrogen
CAR: carboxylic acid reductase
UV: ultraviolet
K_m_: Michaelis constant
V_max_: maximum rate of reaction
k_cat_: catalytic rate constant
HGT: horizontal gene transfer
ACSF4-U26: Acyl CoA synthetase family member 4

