## Supplemental data for "Animal-encoded nonribosomal pathway to bursatellin analogs"

#### Supplementary information

#### 1. Materials and Methods

##### 1.1. Bivalve and gastropod transcriptome assembly and mining

BLAST searches using *Drosophila melanogaster* Ebony (CAA11962.1) and the fungal glycine betaine reductase (A0A1U8QWA2.1) limited to phylum Mollusca identified an NRPS with a formylation domain (CAG2199366.1). Genomic context and intron-exon organization of the NRPS hit was inspected manually and using fungal antiSMASH (as a surrogate for animal gene cluster prediction[1]), identifiying *fmt-atr* (NRPS), a methyltransferase domain-containing gene (*act*), a cytochrome p450 (*p450*), and one or two Fmos (*fmos*) in a contiguous genomic region (referred hereafter as the *bur-ox* pathway). Based on the prediction of *N*-formyl-L-tyrosinol as the product of FmtATR, a Scifinder substructure search (https://scifinder.cas.org) identified bursatellin as a potential product of *bur-ox*. To validate these BLAST hits and rule out the source as microbial contamination, several bivalve and gastropod transcriptomes were assembled from NCBI's SRA (Sequence Read Archive) datasets using rnaSPAdes v3.15.4[2] after read trimming with Trimmomatic v0.39[3] (Table S1). When multiple datasets were available for an organism, the largest SRA datasets (in terms of base size) were selected. Numerous whole organism, embryo, larvae, gill, mantle, digestive gland, and CNS transcriptomes were thus screened. If a transcriptome contained homologs of *act* and *fmt-atr*, we considered that animal as harboring the *bur-ox* pathway.

##### 1.2 Protein structure modeling

AlphaFold[4]-generated models for isonocardicin synthase (Q9RAH6) and McalACT (A0A6J8C2P8) were superimposed using the matchmaker feature in ChimeraX[5] to infer fold conservation. To identify a cut site for soluble expression of the T-R didomain construct CvTR from the full NRPS CvFmtATR, homology models generated via SWISS-MODEL[6] using multiple templates were used. The cut site was 53 residues away from the active site serine (part of the GGNS motif) within the T-domain.

##### 1.3 Strains used in this study

*E. coli* strains DH10β and StBl3 were used for cloning and sequencing of all plasmids constructed in this study. For recombinant protein expression in *E. coli*, strain C43(DE3) was used. *Saccharomyces cerevisiae* strain BJ5464-npgA (MATα ura3-52 trp1 leu2-Δ1 his3Δ200 pep::HIS3 prb1d1.6R can1 GAL) was used for recombinant protein expression in yeast when the expressed proteins required phosphopantetheinylation as a posttranslational modification. *Saccharomyces cerevisiae* strain BJ5464 without the chromosomally integrated phosphopantetheinyl transferase *npgA* was used for expression of non-phosphopantetheinylated proteins in yeast.

##### 1.4 Construction of plasmids

*Mcal-act* (isonocardicin synthase homolog from *Mytilus californianus*) was codon optimized and synthesized with a C-terminal 6xHis tag by Genewiz and cloned in-house into pET28b vector using NcoI and XhoI restriction sites to generate the plasmid, pET28b-McalACT. Yeast codon optimized gene fragments of *Mcalfmt-atr* and *Mefmt-atr* (the full NRPS homologs from *M. californianus* and *M. edulis*, respectively, were procured from Genewiz and assembled in-house using Gibson assembly or yeast homologous recombination into pXW55 expression vector.[7] *Cvfmt-atr* (NRPS homolog from *Crassostrea virginica*) was yeast codon optimized and synthesized with a N-terminal FLAG and C-terminal His tag by Genewiz and cloned in-house via SpeI and PmlI restriction sites into pXW55 to generate pXW55-CvFmtATR. pXW55-CvATR was generated from pXW55-CvFmtATR by site-directed mutagenesis. *Cv-tr* was PCR amplified from plasmid pXW55-CvFmtATR and cloned back into the pXW55 backbone via SpeI and PmlI restriction sites to generate pXW55-CvTR. All plasmids were sequence verified by nanopore sequencing at Plasmidsaurus (https://www.plasmidsaurus.com). The maps of all plasmids constructed for this study are presented in Figure S63. Primer sequences are presented in Table S4.

##### 1.5 Expression and purification of proteins

All bacterial expression plasmids were transformed into chemically competent *E. coli* C43(DE3) cells. A single transformant colony was inoculated into 10 mL LB medium containing 50 μg/mL kanamycin and grown overnight at 30 °C with shaking at 180 rpm. The overnight culture was used to inoculate 1 L of terrific broth containing 50 μg/mL kanamycin, and the culture was incubated at 37 °C and 220 rpm until the OD600 was between 0.4-0.6. The growth temperature was then maintained at 4 °C for 30 mins. Finally, 0.3 mM IPTG was added to induce protein expression, and the culture was grown at 16 °C and 220 rpm for 16-20 h. The bacterial pellet was collected by centrifugation at 3,739 × *g* for 10 mins at 4 °C and either used for protein purification immediately or stored at -80 °C. The cell pellet was resuspended in lysis buffer (50 mM Tris, 10 mM imidazole, 500 mM NaCl, pH 8.0) at 5 mL/g of pellet with addition of lysozyme (600 µg/mL), DNAse (20 µg/mL), and 10 mM $MgCl_2$, lysed by sonication on ice (5 cycles with pulse on for 1 s and off for 3 s at an amplitude of 50%), and centrifuged (32,913 × *g* for 35 mins at 4 °C). The supernatant was further clarified using a 0.45 µm syringe filter (Millex-HV, Sigma), incubated with 5 mL of nickel resin (pre-equilibriated with lysis buffer) for 20 mins in the cold-room, and washed twice with 50 mL of wash buffer (50 mM Tris, 30 mM imidazole, 500 mM NaCl, pH 8.0). The protein was eluted with 15 mL of elution buffer (50 mM Tris, 250 mM imidazole, 500 mM NaCl, pH 8.0), concentrated to 1 mL using an Amicon Ultra-15 10 kDa MWCO centrifugal filter (EMD Millipore), applied onto a Superose™ 6 Increase 10/300 sizing column (GE Healthcare), and eluted in 50 mM $NaH_2PO_4$, 150 mM NaCl, pH 8.0 buffer. The fraction containing the desired protein was concentrated again to 0.2 mL using centrifugal filtration and used for assays. Protein yields were determined by NanoDrop using the theoretical molar extinction coefficient and molecular weight values generated via the ProtParam tool from ExPasy (Expert Protein analysis system).[8] Yield of McalACT was 5.1 mg/mL (x 0.2 mL) from 25 g of bacterial cell pellet.

Competent cell preparation and plasmid transformation for *S. cerevisiae* strains BJ5464 and BJ5464-npgA were performed using the Frozen-EZ Yeast Transformation II kit (Zymo Research). Transformant colonies were selected on uracil-deficient agar (1.39 g/L yeast synthetic drop-out media supplements without uracil (Sigma-Aldrich), 6.7 g/L yeast nitrogen base (Sigma-Aldrich), 20 g/L D-glucose, and 20 g/L agar) after incubation of plates at 30 °C for 48 h. A single colony was inoculated into 10 mL uracil-deficient broth (1.39 g/L yeast synthetic drop-out media supplements without uracil (Sigma-Aldrich), 6.7 g/L yeast nitrogen base (Sigma-Aldrich), and 20 g/L D-glucose) and grown for 20 h at 30 °C and 180 rpm. The 10 mL starter culture was used to inoculate 1L yeast peptone dextrose broth (10 g/L yeast extract, 20 g/L peptone, and 20 g/L glucose), which was in turn grown at 28 °C for 96 h before the yeast cells were pelleted via centrifugation for 10 minutes at 4 °C and 3,739 × *g*. For protein purification, yeast pellets were resuspended in lysis buffer (50 mM Tris, 500 mM NaCl, 10 mM imidazole, pH 8.0) and extruded through a microfluidizer (LM20 Microfluidizer™ Processor) at 20,000 psi five times to lyse the cells. The subsequent steps of lysate processing were same as described above for protein purification from *E. coli*. The CvTR fractions from the S6 column contained an abundant lower molecular weight contaminant. Therefore, these fractions were cleaned using FLAG tag-based affinity purification (GenScript, L00432) via competitive elution with FLAG peptide (GLPBio, GP10149) according to manufacturer's instructions to get pure CvTR protein. Yield of CvTR was >5 mg/mL (x 1 mL) from 40 g of yeast cell pellet.

1.6 Reductase (CvTR) assay

For aminoacyl CoA substrates, reaction conditions for initial screening were as follows: CvTR (10 µM) was incubated with Sfp (1 µM), aminoacyl CoA (200 µM), NADPH (500 µM), and $MgCl_2$ (10 mM) in 50 mM HEPES, 150 mM NaCl, pH 8.0 buffer at 25 °C for 20 h in 50 µL scale. Three aminoacyl CoA substrates were tested: *N*-formyl-L-tyrosine CoA (**3Ca**), *N*-Boc-L-tyrosine CoA (**3Cb**), and *N*-acetyl-*O*-*tert*-butyl-L-tyrosine CoA (**3Cc**). Stock solutions (either 1 or 10 mM) of all CoA substrates were prepared in 1% DMSO. Controls used included a reaction with no enzyme, a reaction with no NADPH, and a reaction with no Sfp. All reactions were performed in triplicate. The reactions were quenched with MeCN (150 µL), centrifuged at 16,200 g for 10 min to precipitate the protein, and analyzed by UPLC-ESIMS (Waters™ Xevo™ G2-XS Q-Tof) in positive mode. Reactions were injected on to an Acquity UPLC BEH C18 1.7 µm column (2.1 x 150 mm) at 0.3 mL/min flowrate over a 10 min 5-100% mobile phase B gradient with water + 0.1% formic acid and MeCN as mobile phase A and B, respectively. The injection volume was varied from 1-4 µL to get the best peak shape. The identity of the products from reactions containing **3Ca** and **3Cb** were verified by comparison with synthetic standards of *N*-formyl-L-tyrosinol (**4a**) and *N*-Boc-L-tyrosinol (**4b**), respectively. The product from **3Cc** reaction was verified using HR-MS/MS fragmentation pattern (Figure S52B) by analyzing assay samples using a Waters™ Xevo™ G2-S Q-Tof mass spectrometer equipped with a diode array detector with data analysis using MassLynx 4.1 software. Conditions used for HR-MS/MS analysis of synthetic standards of **4a** and **4b** are mentioned in section 1.11.1.

CvTR (10 µM) was incubated with aminoacyl SNAC (500 µM), NADPH (500 µM), and $MgCl_2$ (10 mM) under the same initial conditions used for the CoA substrates. *N*-formyl-L-tyrosine SNAC (**3Sa**), *N*-acetyl-L-tyrosine SNAC (**3Sc**), and *N*-acetyl-L-phenylalanine SNAC (**3Sd**) were the substrates tested initially with stock solutions prepared as described above for the CoA substrates. All reactions were performed in triplicate. SNAC substrate reactions were processed similar to the CoA substrate reactions, and identity of the product peak was validated by coelution with the respective synthetic standard. An additional control in the form of *N*-formyl-L-tyrosine (**2a**) as a substrate was used to confirm whether the observed reductase activity was thioester-dependent.

1.7 Yield estimation for CvTR-catalyzed formation of **4b**

For additional experiments, based on ease of substrate synthesis and stability upon storage, **3Cb** and its cognate SNAC thioester equivalent, *N*-Boc-L-tyrosine SNAC (**3Sb**), were used as substrates. The product from these reactions, **4b**, was also easier to quantify chromatographically with no peak broadening or splitting as was often observed with the more polar acyl tyrosinols. Standards of **4b** diluted from a 50 mM (50% methanolic) stock solution in the concentration range of 0.05-4mM were injected (10 µL) in triplicate on an analytical C18 column (Phenomenex Luna) over a 5-50% solvent B gradient with water + 0.01% TFA and MeCN as solvents A and B, respectively. The AUC at 280 nm was used to generate a **4b** standard curve for yield estimation experiments (Figure S64).

To determine whether NADH could be utilized as an alternative reductant, CvTR (10 µM) was incubated with **3Cb** or **3Sb** (100 µM), Sfp (10 µM), NADH (1 mM), and $MgCl_2$ (10 mM) in 50 mM $NaH_2PO_4$, 150 mM NaCl, pH 8.0 buffer at 25 °C for 20 h in 50 µL scale. Parallel reactions with NADPH as reductant and a no-Sfp control containing **3Cb** and NADPH were also included in these experiments aimed at comparing the 20 h yield of **4b**. All reactions were performed in triplicate on two different days. Each reaction was quenched with 150 µL methanol after 20 h, dried by SpeedVac, resuspended in 50 µL of 4% methanol, and centrifuged at 16,200 x *g* for 10 min to precipitate the protein. The supernatant was analyzed using the same method described for the **4b** standards above. The average yield from six replicates was plotted as a bar graph using MS Excel.

1.8 Aminocarboxypropylation (McalACT) assay

McalACT (10 µM) was incubated with the appropriate substrate (1 mM), SAM (1 mM), and $MgCl_2$ (10 mM) in 50 mM $NaH_2PO_4$, 150 mM NaCl, pH 8.0 buffer at 25 °C for 20 h in 50 µL scale. The substrates tested included *N*-formyl-L-tyrosinol (**4a**), *N*-acetyl-L-tyrosinol (**4c**), *N*-propionyl-L-tyrosinol (**4d**), *N*-butyryl-L-tyrosinol (**4e**), *N*-Boc-L-tyrosinol (**4b**), and *N*-acetyl-L-phenylalaninol (**4f**). A stock solution of **4b** (100 mM) dissolved in DMSO or methanol was used to achieve the working concentration in assays such that no more than 1% DMSO or methanol was present in the final reaction mixture. All other substrates were dissolved in the reaction buffer itself to make 100 mM stock solutions. Controls for the reaction with **4a** as substrate included a reaction with no enzyme and a reaction with no SAM. For all other substrates, only the no enzyme control was used. All reactions were performed in triplicate. The reactions were quenched with MeCN (150 µL), centrifuged at 16,200 x *g* for 10 min to precipitate the protein, and diluted further with 50 µL water prior to analysis by UPLC-ESIMS in positive mode. For detection of the product, reactions were injected on to an Acquity UPLC BEH C18 1.7 µm column (2.1 x 150 mm) at 0.4 mL/min flowrate over a 10 min 5-100% mobile phase B gradient with water + 0.1% formic acid and MeCN as mobile phase A and B, respectively. The flowrate was reduced to 0.3 mL/min for reactions containing **2a** to allow for sufficient retention of the highly polar reactant and product. The injection volume was varied from 1-4 µL to get the best peak shape. Identity of the reaction product *N*-Boc-L-tyrosinol *O*-homoserine ether (**5b**) derived from substrate **4b** was verified after reaction scale-up by NMR-based characterization of the purified product. All NMR analyses described in this study were run on Varian Innova NMR spectrometers. For all other substrates, product identity was verified using HR-MS/MS fragmentation pattern based on observation of expected product *m/z* in reactions. HR-MS/MS analyses were performed on a Waters™ Xevo™ G2-S Q-Tof mass spectrometer equipped with a diode array detector with data analysis using MassLynx 4.1 software.

For large-scale purification of **5b**, 96 x 100 µL scale reactions with enzyme and substrate concentrations maintained as described above were incubated in a thermocycler. This was repeated thrice, and the reactions were combined and quenched with 2x volume of MeCN and centrifuged at 3,739 x g for 10 min to remove the precipitated protein. The organic layer was dried using a SpeedVac, and the residue was resuspended in 50% MeCN (200 µL). During purification by reverse phase HPLC using an analytical C18 column (Phenomenex Luna) over a 5-50% solvent B gradient with water + 0.01% TFA and MeCN as solvents A and B, respectively, **5b** underwent Boc deprotection. Therefore, the partially deprotected material was subject to another round of HPLC purification under the same conditions as described for **5b** with collection of L-tyrosinol *O*-homoserine ether (**5b***) for NMR analysis. The yield was ~ 1 mg.

*N*-formyl-L-tyrosinol *O*-homoserine ether (**5a**): HRMS (*m/z*): $[M + H]^+$ calcd for $C_{14}H_{21}N_2O_5^+$ 297.1445, found 297.1452. $MS^2$ diagnostic fragments: 102.0546, 107.0485, 133.0693, 145.0645.

*N*-acetyl-L-tyrosinol *O*-homoserine ether (**5c**): HRMS (*m/z*): $[M + H]^+$ calcd for $C_{15}H_{23}N_2O_5^+$ 311.1602, found 311.1608. $MS^2$ diagnostic fragments: 102.0561, 107.0504, 133.0654, 145.0655, 293.1501.

*N*-propionyl-L-tyrosinol *O*-homoserine ether (**5d**): HRMS (*m/z*): $[M + H]^+$ calcd for $C_{16}H_{25}N_2O_5^+$ 325.1758, found 325.1804. $MS^2$ diagnostic fragments: 102.0563, 107.0502, 133.0653, 145.0652, 307.1651.

*N*-butyryl-L-tyrosinol *O*-homoserine ether (**5e**): HRMS (*m/z*): $[M + H]^+$ calcd for $C_{17}H_{27}N_2O_5^+$ 339.1915, found 339.1987. $MS^2$ diagnostic fragments: 102.0560, 107.501, 133.0652, 145.0652, 321.1806.

*N*-Boc-L-tyrosinol *O*-homoserine ether (**5b**): HRMS (*m/z*): $[M + H]^+$ calcd for $C_{18}H_{29}N_2O_6^+$ 369.2021, found 369.2033. $MS^2$ diagnostic fragments: 102.0560, 107.0499, 133.0653, 145.0651.

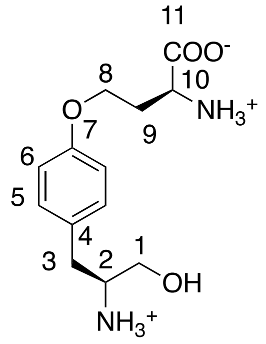


L-tyrosinol *O*-homoserine ether (**5b***): $^1$H NMR (500 MHz, $CD_3OD$): δ = 7.21 (d, *J* = 8.5 Hz, 2H), 6.96 (d, *J* = 8.5 Hz, 2H), 4.2 (m, 2H), 4.2 (m, 1H), 3.67 (dd, *J* = 11.5,3.5 Hz, 1H), 3.50 (dd, *J* = 11.5,6 Hz, 1H), 3.40 (m, 1H), 2.88 (d, *J* = 7.5 Hz, 2H), 2.57 (m, 1H), 2.35 (m, 1H). $^{13}$C NMR (125 MHz, $CD_3OD$) (assigned from HSQC of **5b** and **5b*** mixture): δ = 171.2 (C-10), 158.8 (C-7), 131.4 (C-5), 129.5 (C-4), 115.9 (C-6), 65.0 (C-1), 61.0 (C-8), 55.7 (C-2), 52.1 (C-10), 35.4 (C-3), 31.2 (C-9). HRMS (*m/z*): $[M + H]^+$ calcd for $C_{13}H_{21}N_2O_4^+$ 269.1496, found 269.1455.

1.9 Determination of kinetic parameters for aminocarboxypropylation of **4b**

As **4b** is a commercially available compound, the rate of consumption of **4b** rather than formation of **5b** was measured for kinetic parameter estimation. Prior to doing kinetic experiments for McalACT-catalyzed conversion of **4b** to **5b,** multiple reaction parameters were varied to determine the conditions for maximum conversion. In all cases, reactions containing **4b** (0.5 mM) and $MgCl_2$ (10 mM) in 50 mM $NaH_2PO_4$, 150 mM NaCl, pH 8.0 buffer were incubated for 20 h in 50 µL scale. The samples were processed as described in section 1.7 for analysis by RP-HPLC. Two reaction temperatures (30 and 37 °C) and McalACT concentrations (20 µM and 40 µM) were screened with 0.5 mM SAM (1 eq.). Four SAM concentrations (0.1–10 eq.) were screened with 20 µM McalACT at 25 °C. Based on the observation that product formation increased with SAM concentration and was inhibited at higher temperatures (Figure S38), an initial kinetic experiment was run by quenching reactions containing **4b** (0.5 mM), McalACT (20 µM), SAM (5 mM), and $MgCl_2$ (10 mM) and incubated at 25 °C every 25 mins until 200 min, at 6 h, and at 24 h (Figure S39A). As the conversion was ~50% at 200 mins with 20 µM McalACT and AUC estimation was challenging with shorter time intervals that would allow initial rate estimation, the enzyme concentration was halved for the final kinetic experiments to slow down the reaction. The final conditions thus chosen were **4b** (0.2-1 mM), 10 µM McalACT, 10 mM $MgCl_2$, and 10-fold excess SAM (5 mM) at 25 °C with reactions being quenched in triplicate every hour until 6 h such that a large excess of the substrate remained unconverted (Figure S39B). Using the standard curve generated for **4b** (section 1.7), the AUC for each replicate of a triplicate set was converted to residual **4b** concentration leading to initial rate estimation in triplicate for each initial concentration of **4b** tested. The initial concentration vs rate values were fitted using non-linear regression function (Prism 10.3.1, GraphPad Software, San Diego, CA, USA) to get kinetic constant estimates.

1.10 One-pot reactions with CvTR and McalACT

One-pot reactions containing McalACT (10 µM), CvTR (10 µM), **3Sa** (1 mM), SAM (5 mM), NADPH (5 mM) and $MgCl_2$ (10 mM) in 50 mM $NaH_2PO_4$, 150 mM NaCl, pH 8.0 buffer were incubated at 25 °C for 20 h in 50 µL scale. Parallel reactions containing McalACT (10 µM), **3Sa** (1 mM), SAM (5 mM), and $MgCl_2$ (10 mM) were also setup under the same conditions. For **3Sb** and **3Cb**, onepot reactions containing the respective substrate (100 µM), McalACT (10 µM), CvTR (10 µM), SAM (1 mM), NADPH (1 mM) and $MgCl_2$ (10 mM) in 50 mM $NaH_2PO_4$, 150 mM NaCl, pH 8.0 buffer were incubated at 25 °C for 20 h in 50 µL scale with parallel reactions containing only McalACT and substrate. Reactions containing *N*-Boc-L-tyrosine (**2b**) (100 µM), McalACT (10 µM), SAM (1 mM), and $MgCl_2$ (10 mM) were additionally setup to probe activity of McalACT against tyrosines. All reactions in triplicate were quenched with MeCN (150 µL), centrifuged at 16,200 x *g* for 10 min to precipitate the protein, and analyzed by UPLC-ESIMS in positive mode under the same conditions as described in section 1.8 for reactions containing **4a** as substrate.

1.11 Synthetic methods

1.11.1 Synthesis of acyl tyrosinols

L-tyrosinol hydrochloride (**4**) and compound **4b** were procured from Sigma-Aldrich (catalog no. 469998) and Chemimpex International (catalog no. 11897), respectively. **4c**, **4d**, and **4e** were synthesized by modifying a previously published protocol.[9] Briefly, 1 eq. of **4** dissolved in THF was mixed with 4 eq. of $K_2CO_3$ dissolved in water (THF:water = 2:1 v/v) followed by addition of the appropriate acyl chloride (1 eq.) after 45 mins. The reaction was thereafter allowed to proceed for < 1 h to minimize formation of undesired diacylated byproducts. The reaction mixtures were filtered and concentrated by rotary evaporation. The residue was resuspended in a minimal volume of 50% MeCN and purified by reverse phase HPLC using an analytical C18 column (Phenomenex Luna) over a 5-100% solvent B gradient with water + 0.01% TFA and MeCN as solvents A and B, respectively. For 50 mg scale synthesis, the yields of **4c**, **4d**, and **4e** were 11 mg, 27.7 mg, and 12.9 mg, respectively. HR-MS/MS analyses were performed on a Thermo Scientific™ Orbitrap Exploris™ 120 mass spectrometer with data analysis using Freestyle 1.7 software. NMR and HR-MS/MS spectra for acyl tyrosinols are reported in section 3.2 (Figures S48-S52).

*N*-acetyl-L-tyrosinol (**4c**): $^1$H NMR (500 MHz, Acetone-$d_6$): δ = 7.06 (d, *J* = 8.5 Hz, 2H), 6.74 (d, *J* = 8 Hz, 2H), 4.01 (m, 1H), 3.50 (d, *J* = 4.5 Hz, 2H), 2.79 (dd, *J* = 13.5,7 Hz, 1H), 2.68 (dd, *J* = 13.5,7.5 Hz, 1H), 1.85 (s, 3H) ppm. $^{13}$C NMR (125 MHz, Acetone-$d_6$): δ = 170.8, 157.2, 131.5, 131.0, 116.4, 64.3, 54.5, 37.4, 23.5 ppm. HRMS (*m/z*): [M + H]$^+$ calcd for $C_{11}H_{16}NO_3^+$ 210.1125, found 210.1121.

*N*-propionyl-L-tyrosinol (**4d**): $^1$H NMR (500 MHz, Acetone-$d_6$): δ = 7.06 (d, *J* = 9 Hz, 2H), 7.00 (d, *J* = 8 Hz, 1H), 6.74 (d, *J* = 9 Hz, 2H), 4.03 (m, 1H), 3.50 (d, *J* = 5.5 Hz, 2H), 2.80 (dd, *J* = 13.5,8 Hz, 1H), 2.68 (dd, *J* = 14.5,7.5 Hz, 1H), 2.14 (q, *J* = 7.5 Hz, 2H), 1.02 (t, *J* = 7.5 Hz, 3H) ppm. $^{13}$C NMR (125 MHz, Acetone-$d_6$): δ = 174.6, 157.2, 131.4, 130.8, 116.3, 64.3, 54.4, 37.3, 29, 10.6 ppm. HRMS (*m/z*): [M + H]$^+$ calcd for $C_{12}H_{18}NO_3^+$ 224.1282, found 224.1279.

*N*-butyryl-L-tyrosinol (**4e**): $^1$H NMR (500 MHz, Acetone-$d_6$): δ = 7.07 (d, *J* = 8.5 Hz, 2H), 6.91 (d, *J* = 8.5 Hz, 1H), 6.74 (d, *J* = 8.5 Hz, 2H) 4.06 (m, 1H), 3.51 (d, *J* = 5.5 Hz, 2H), 2.82 (dd, *J* = 13.5,6.5 Hz, 1H), 2.68 (dd, *J* = 14,8 Hz, 1H), 2.11 (t, *J* = 7.5 Hz, 2H), 1.55 (h, *J* = 7.5 Hz, 2H), 0.84 (t, *J* = 7.5 Hz, 3H) ppm. $^{13}$C NMR (125 MHz, Acetone-$d_6$): δ = 173.9, 157.1, 131.5, 131.0, 116.3, 64.5, 54.5, 39.2, 37.3, 20.3, 14.4 ppm. HRMS (*m/z*): [M + H]$^+$ calcd for $C_{13}H_{20}NO_3^+$ 238.1438, found 238.1438.

**4a** was synthesized as follows: To a mixture of **4** (1 eq.) and *N*-formylsaccharin (Ambeed, catalog no. A427512) (1.5 eq.) in THF, $K_2CO_3$ (2.5 eq.) dissolved in water (THF:water = 1:1 v/v) was added. After 1 h, the reaction mixture was filtered, concentrated to remove THF, and resuspended in a minimal volume of 50% MeCN for purification by reverse phase HPLC using an analytical C18 column (Phenomenex Luna) over a 5-100% mobile phase B gradient with water and MeCN as solvents A and B, respectively. For 150 mg scale synthesis, the purified yield of **4a** was 41 mg.

*N*-formyl-L-tyrosinol (**4a**): $^1$H NMR (500 MHz, Acetone-$d_6$): δ = 8.09 (s, 1H), 7.08 (d, *J* = 8 Hz, 2H), 6.75 (d, *J* = 7.5 Hz, 2H), 4.09 (m, 1H), 3.52 (d, *J* = 4.5 Hz, 2H), 2.84 (dd, J = 13.5,7 Hz, 1H), 2.71 (dd, J = 13.5,7.5 Hz, 1H) ppm. $^{13}$C NMR (125 MHz, Acetone-$d_6$): δ = 162.0, 157.3, 131.7, 130.8, 116.5, 64.1, 53.2, 37.4 ppm. HRMS (*m/z*): [M + H]$^+$ calcd for $C_{10}H_{14}NO_3^+$ 196.0969, found 196.0967.

1.11.2 Aminoacyl CoA synthesis via thiophenolation

*N*-formyl-L-tyrosine thiophenolate (**7a**), *N*-Boc-L-tyrosine thiophenolate (**7b**), and *N*-acetyl-*O*-*tert*-butyl-L-tyrosine thiophenolate (**7c**) were synthesized as follows: amino acid (1 eq.), DCC (1 eq.) and HOBt (1 eq.) were dissolved in up to 4 mL ethyl acetate (or THF in case of **2a**). Thiophenol (1.2 eq.) was added to the reaction mixture after formation of dicyclohexyl urea was visibly observed. The mixture was stirred for 22 h, filtered through a PTFE syringe filter (0.2 μm), and washed with 10% citric acid. In case of **7a**, THF was first removed by rotary evaporation, and the residue was resuspended in ethyl acetate prior to extraction with 10% citric acid. The organic layer was dried over $Na_2SO_4$ and concentrated by rotary evaporation. The residue was resuspended in a minimum volume of MeCN and injected on to a reverse phase C18 semipreparative column (Phenomenex Luna). Aminoacyl thiophenolates were thus purified over a 5-100% mobile phase B gradient with water + 0.01% TFA and MeCN as solvents A and B, respectively. For 50 mg scale synthesis, the yields of **7b** and **7c** were 44.68 mg and 49 mg, respectively. For **7a**, the yield from a 150 mg scale synthesis was 82 mg. NMR spectra for aminoacyl thiophenolates are reported in section 3.5 (Figures S60-S62).

*N*-Boc-L-tyrosine thiophenolate (**7b**): $^1$H NMR (500 MHz, $CDCl_3$): δ = 7.37 (br m, 3H), 7.33 (br m, 2H), 7.01 (d, *J* = 8 Hz, 2H), 6.74 (d, *J* = 8 Hz, 2H), 5.02 (d, *J* = 10 Hz, 1H), 4.69 (m, 1H), 3.05 (m, 2H), 1.43 (s, 9H) ppm. $^{13}$C NMR (125 MHz, $CDCl_3$): δ = 199.8, 155.4 (2C), 134.8, 130.8, 129.8, 129.5, 127.4, 115.9, 81.0, 61.4, 37.9, 28.6 ppm. HRMS (*m/z*): [M + H]$^+$ calcd for $C_{20}H_{24}NO_4S^+$ 374.1421, found 374.1429.

*N*-acetyl-*O*-*tert*-butyl tyrosine thiophenolate (**7c**): $^1$H NMR (500 MHz, $CDCl_3$): δ = 7.34 (br m, 3H), 7.30 (br m, 1H), 7.06 (d, *J* = 8 Hz, 2H), 6.90 (d, *J* = 8 Hz, 2H), 6.5 (br, 1H), 5.01 (m, 1H), 3.12 (dd, *J* = 14,6.5 Hz, 2H), 3.01 (dd, *J* = 15,8 Hz, 2H), 1.92 (s, 3H), 1.30 (s, 9H) ppm. $^{13}$C NMR (125 MHz, $CDCl_3$): δ = 198.6, 170.3, 154.6, 134.6, 130.6,129.9, 129.6, 129.3, 127.0, 125.4, 78.5, 60.0, 37.6, 28.9, 23.0 ppm. HRMS (*m/z*): [M + H]$^+$ calcd for $C_{21}H_{26}NO_3S^+$ 372.1628, found 372.1649.

*N*-formyl-L-tyrosine thiophenolate (**7a**): $^1$H NMR (500 MHz, Acetone-$d_6$): δ = 8.30 (br s, 1H), 8.23 (br s, 1H), 7.84 (d, *J* = 8 Hz, 1H), 7.45 (m, 3H), 7.40 (m, 2H), 7.12 (d, *J* = 10 Hz, 2H), 6.79 (d, *J* = 10 Hz, 2H), 4.95 (m, 1H), 3.16 (dd, *J* = 13.5,5 Hz, 1H), 2.99 (m, 1H) ppm. $^{13}$C NMR (125 MHz, acetone-$d_6$): δ = 197.0, 160.7, 156.0, 134.2, 130.0, 128.9, 128.8, 127.3, 126.5, 114.9, 58.8, 36.3 ppm. HRMS (*m/z*): [M + H]$^+$ calcd for $C_{16}H_{16}NO_3S^+$ 302.0846, found 302.0871.

*N*-formyl-L-tyrosine CoA (**3Ca**), *N*-Boc-L-tyrosine CoA (**3Cb**), and *N*-acetyl-*O*-*tert*-butyl-L-tyrosine CoA (**3Cc**) were made by exchanging the corresponding aminoacyl thiophenolates with coenzyme A as follows: aminoacyl thiophenolate (1 eq.) dissolved in 150 μL MeCN was mixed with 1 eq. of coenzyme A dissolved in 150 μL phosphate buffer (pH 8.0). The reaction was allowed to proceed for 2 h and directly injected on a reverse phase C18 semipreparative column (Phenomenex Luna) and purified under the same conditions used for the thiophenolates. From **7b** (7.4 mg) and coenzyme A (7.8 mg), the yield of

**3Cb** was 6.28 mg. From **7a** (4.5 mg) and coenzyme A (7.8 mg), the yield of **3Ca** was 5.5 mg. From **7c** (5 mg) and coenzyme A (7 mg), the yield of **3Cc** was 6 mg. NMR data for CoA esters were assigned using 2D data (section 3.3 – Figures S53-S55) and compared with those from literature (Table S5).[10]

*N*-formyl-L-tyrosine CoA (**3Ca**): HRMS (*m/z*): [M + H]+ calcd for $C_{31}H_{46}N_8O_{19}P_3S^+$ 959.1808, found 959.1804.

*N*-Boc-L-tyrosine CoA (**3Cb**): HRMS (*m/z*): [M + H]+ calcd for $C_{35}H_{54}N_8O_{20}P_3S^+$ 1031.2382, found 1031.2361.

*N*-acetyl-*O*-*tert*-butyl-L-tyrosine CoA (**3Cc**): HRMS (*m/z*): [M + H]+ calcd for $C_{36}H_{56}N_8O_{19}P_3S^+$ 1029.2590, found 1029.2573.

1.11.3 Aminoacyl *N*-acetyl cysteamine (aminoacyl SNAC) synthesis
**3Sb**, **3Sc**, and **3Sd** were synthesized based on a previously published protocol.[11] Briefly, the respective amino acid (1 eq.), DCC (1 eq.), HOBt (1 eq.), and $K_2CO_3$ (0.5 eq.) were dissolved in 4 mL THF. *N*-acetylcysteamine (2 eq. for *N*-acetyl-*O*-*tert*-butyl-L-tyrosine (**2c**) and *N*-acetyl-L-phenylalanine (**2d**) and 5 eq. for *N*-Boc-L-tyrosine (**2b**)) was added to the reaction mixture after formation of dicyclohexyl urea was visibly observed. The reaction was stirred for 3 h, filtered, and concentrated by rotary evaporation. The residue was resuspended in ethyl acetate, washed with 10% aqueous $NaHCO_3$, dried over $Na_2SO_4$, and purified on a reverse phase C18 semipreparative column (Phenomenex Luna) under the same conditions used for the thiophenolates. *N*-acetyl-*O*-tert-butyl-L-tyrosine SNAC was deprotected overnight in 1 mL of 50% TFA/$CH_2Cl_2$, concentrated by rotary evaporation with redissolution in $CH_2Cl_2$ at least twice for TFA removal, and subject to another round of RP-HPLC purification under the same conditions used for the thiophenolates to yield **3Sc**. For 50 mg scale synthesis, the yields of **3Sc, 3Sd,** and **3Sb** were 48 mg, 34 mg, and 49 mg respectively. **3Sa** was made by exchanging **8a** with *N*-acetylcysteamine using the same procedure as described for thiophenolate-coenzyme A exchange. The yield of **3Sa** from 50 mg of **8a** was 27 mg. NMR spectra for aminoacyl SNAC substrates are reported in section 3.4 (Figures S56-S59).

*N*-formyl-L-tyrosine SNAC (**3Sa**): $^1$H NMR (500 MHz, Acetone-*d*$_6$): δ = 8.16 (br s, 1H), 7.08 (d, *J* = 8 Hz, 2H), 6.76 (d, *J* = 8.5 Hz, 2H), 4.80 (dd, *J* = 9.5,5.5 Hz, 1H), 3.30 (m, 2H), 3.11 (dd, *J* = 14.5,5 Hz, 1H), 2.98 (t, *J* = 6.5 Hz, 2H), 2.89 (dd, *J* = 14.5,9 Hz, 1H), 1.86 (s, 3H) ppm. $^{13}$C NMR (125 MHz, Acetone-*d*$_6$): δ = 200.8, 170.7, 165.3, 162.3, 157.7, 131.9, 128.5, 116.5, 60.6, 39.9, 38.1, 29.5, 23.3 ppm. HRMS (*m/z*): [M + H]+ calcd for $C_{14}H_{19}N_2O_4S^+$ 311.1061, found 311.1064.

*N*-acetyl-L-tyrosine SNAC (**3Sc**): $^1$H NMR (500 MHz, Acetone-d$_6$): δ = 7.87 (d, *J* = 9 Hz, 1H), 7.53 (br t, *J* = 7 Hz, 1H), 7.08 (d, *J* = 8.5 Hz, 2H), 6.75 (d, *J* = 8.5 Hz, 2H), 4.71 (m, 1H), 3.33 (m, 2H), 3.08 (dd, *J* = 15,5 Hz, 1H), 2.98 (t, *J* = 6.8 Hz, 2H), 2.82 (dd *J* = 15,9.5 Hz, 1H), 1.92 (s, 1H), 1.91 (s, 1H) ppm. $^{13}$C NMR (125 MHz, Acetone-*d*$_6$): δ = 201.1, 173.4, 171.3, 157.2, 130.9, 128.3, 116.1, 62.1, 39.6, 37.5, 28.9, 22.9, 22.8 ppm. HRMS (*m/z*): [M + H]+ calcd for $C_{15}H_{21}N_2O_4S^+$ 325.1217, found 325.1217.

*N*-acetyl-L-phenylalanine SNAC (**3Sd**): $^1$H NMR (500 MHz, DMSO-*d*$_6$): δ = 8.59 (d, *J* = 8 Hz, 1H), 8.03 (br t, *J* = 5 Hz, 1H), 7.2 (m, 5H), 4.54 (m, 1H), 3.14 (m, 2H), 3.08 (dd, *J* = 14.5,5 Hz, 1H), 2.88 (t, *J* = 4 Hz, 2H), 2.80 (dd, *J* = 14,10.5 Hz, 1H), 1.81 (s, 3H), 1.79 (s, 3H) ppm. $^{13}$C NMR (125 MHz, DMSO-*d*$_6$): δ = 200.2, 169.2, 168.8, 136.8, 128.6, 127.8, 126.1, 60.2, 37.6, 36.3, 27.4, 22.1, 21.9 ppm. HRMS (*m/z*): [M + H]+ calcd for $C_{15}H_{21}N_2O_3S^+$ 309.1268, found 309.1261.

*N*-Boc-L-tyrosine SNAC (**3Sb**): $^1$H NMR (500 MHz, DMSO-*d*$_6$): δ = 9.19 (br s, 1H), 8.01 (t, *J* = 5.9 Hz, 1H), 7.58 (d, *J* = 8.5 Hz, 1H), 7.02 (d, *J* = 8.7, 2H) 6.65 (d, *J* = 9 Hz, 2H), 4.13 (m, 1H), 3.14 (m, 2H), 2.91 (m, 1H), 2.86 (t, *J* = 7.1 Hz, 2H), 2.67 (dd, *J* = 10.7, 3 Hz, 1H), 1.79 (s, 3H), 1.33 (s, 9H). $^{13}$C NMR (125 MHz, DMSO-*d*$_6$): δ = 204.9, 172.4, 159.1, 158.5, 133.2, 130.6, 118.1, 81.8, 65.9, 41.3, 38.9, 31.3, 30.9, 25.7 ppm. HRMS (*m/z*): [M + Na]+ calcd for $C_{18}H_{26}N_2NaO_5S^+$ 405.1455, found 405.1462.

#### 2. Supplementary figures

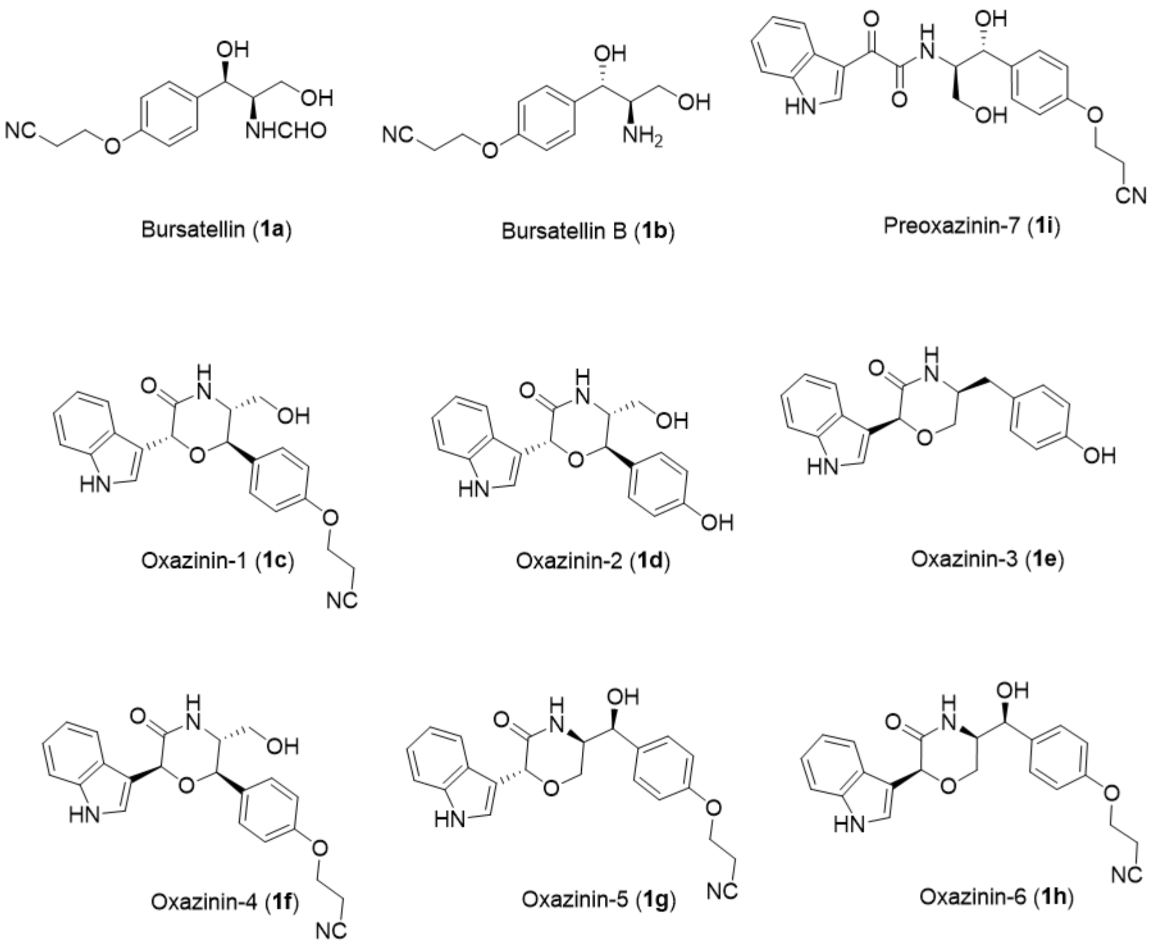


**Figure S1**. Structures of currently known bursatellins and oxazinins from bivalves and gastropods.

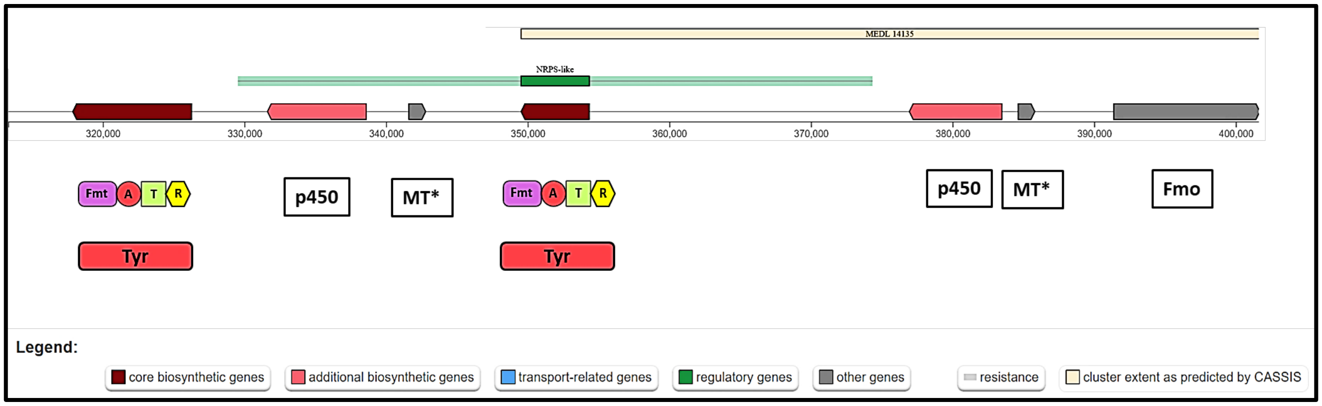


**Figure S2**. Putative bursatellin cluster in the genome of *M. edulis* as annotated by antiSMASH. In addition to duplicate copies of some genes being present on the same contig, the NCBI genome annotation pipeline annotates the two sets of *fmo* exons as an Fmo didomain protein in *M. edulis* and *M. galloprovincialis* genomes but as two separate Fmo proteins in *M. californianus* genome.

A

**Sequences producing significant alignments** Download Select columns Show 100

select all *100 sequences selected* GenPept Graphics Distance tree of results Multiple alignment MSA Viewer

| Description | Scientific Name | Max Score | Total Score | Query Cover | E value | Per. Ident | Acc. Len | Accession |
|---|---|---|---|---|---|---|---|---|
| unnamed protein product [Mytilus coruscus] | Mytilus coruscus | 770 | 770 | 100% | 0.0 | 93.20% | 397 | CAC5390678.1 |
| Hypothetical predicted protein [Mytilus galloprovincialis] | Mytilus galloprovincialis | 763 | 763 | 100% | 0.0 | 91.44% | 397 | VDI72202.1 |
| unnamed protein product [Mytilus edulis] | Mytilus edulis | 760 | 760 | 100% | 0.0 | 91.69% | 397 | CAG2199368.1 |
| unnamed protein product [Mytilus edulis] | Mytilus edulis | 629 | 629 | 97% | 0.0 | 74.55% | 397 | CAG2231838.1 |
| unnamed protein product [Mytilus edulis] | Mytilus edulis | 627 | 627 | 97% | 0.0 | 74.04% | 397 | CAG2199365.1 |
| Hypothetical predicted protein [Mytilus galloprovincialis] | Mytilus galloprovincialis | 583 | 583 | 91% | 0.0 | 74.38% | 380 | VDI72207.1 |
| hypothetical protein KUTeg_012996 [Tegillarca granosa] | Tegillarca granosa | 473 | 473 | 99% | 2e-162 | 57.54% | 403 | KAJ8308122.1 |
| uncharacterized protein LOC111138433 [Crassostrea virginica] | Crassostrea virginica | 458 | 458 | 99% | 1e-156 | 55.53% | 399 | XP_022346107.1 |
| hypothetical protein KP79_PYT09613 [Mizuhopecten yessoensis] | Mizuhopecten yessoensis | 454 | 454 | 99% | 3e-155 | 56.50% | 403 | OWF55694.1 |
| isonocardicin synthase-like [Ostrea edulis] | Ostrea edulis | 454 | 454 | 98% | 5e-155 | 54.16% | 399 | XP_056017047.1 |
| hypothetical protein CHS0354_017497 [Potamilus streckersoni] | Potamilus streckersoni | 465 | 465 | 99% | 5e-154 | 57.25% | 795 | KAK3610896.1 |
| hypothetical protein KP79_PYT01546 [Mizuhopecten yessoensis] | Mizuhopecten yessoensis | 448 | 448 | 99% | 9e-153 | 57.04% | 399 | OWF45616.1 |
| hypothetical protein DPMN_097580 [Dreissena polymorpha] | Dreissena polymorpha | 442 | 442 | 100% | 3e-150 | 55.33% | 404 | KAH3855021.1 |
| hypothetical protein CHS0354_024261 [Potamilus streckersoni] | Potamilus streckersoni | 433 | 433 | 95% | 3e-147 | 55.73% | 382 | KAK3594325.1 |
| hypothetical protein KP79_PYT18090 [Mizuhopecten yessoensis] | Mizuhopecten yessoensis | 434 | 434 | 99% | 3e-147 | 53.75% | 399 | OWF42937.1 |
| isonocardicin synthase-like [Ylistrum balloti] | Ylistrum balloti | 433 | 433 | 99% | 4e-147 | 54.75% | 399 | XP_060086363.1 |
| uncharacterized protein LOC127872473 [Dreissena polymorpha] | Dreissena polymorpha | 440 | 440 | 100% | 1e-143 | 55.33% | 871 | XP_052271766.1 |

B

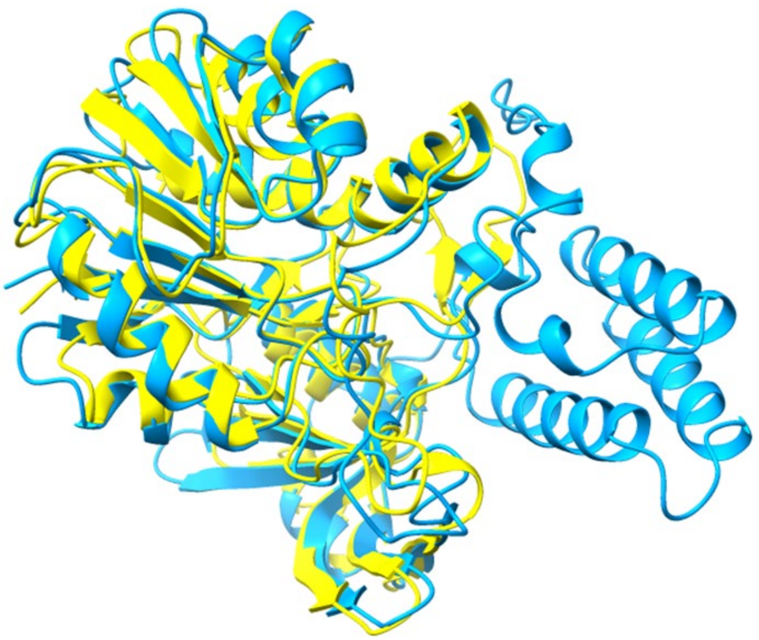

**Figure S3**. **A)** BLAST hits for the methyl transferase domain-containing protein in the putative bursatellin cluster include hits that are annotated as isonocardicin synthase-like (boxed in red). **B)** Superimposition of AlphaFold models of nocardicin 3-amino-3-carboxypropyl transferase (Q9RAH6; yellow ribbon) and McalACT (A0A6J8C2P8; blue ribbon) shows fold conservation from residues 94-397 in the latter protein. McalACT is longer than isonocardicin synthase by 96 residues.

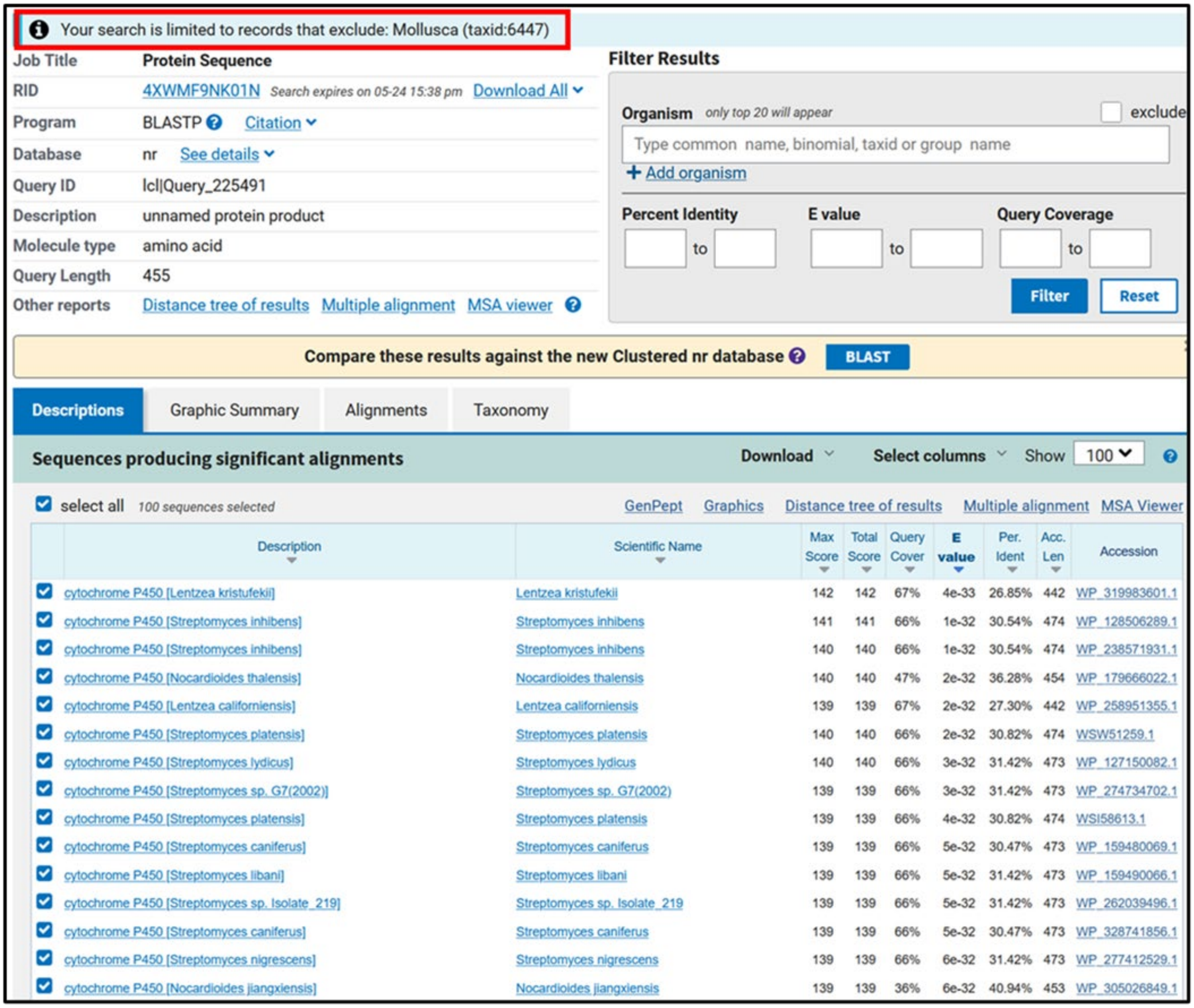


| Description | Scientific Name | Max Score | Total Score | Query Cover | E value | Per. Ident | Acc. Len | Accession |
|---|---|---|---|---|---|---|---|---|
| cytochrome P450 [Lentzea kristufekii] | Lentzea kristufekii | 142 | 142 | 67% | 4e-33 | 26.85% | 442 | WP_319983601.1 |
| cytochrome P450 [Streptomyces inhibens] | Streptomyces inhibens | 141 | 141 | 66% | 1e-32 | 30.54% | 474 | WP_128506289.1 |
| cytochrome P450 [Streptomyces inhibens] | Streptomyces inhibens | 140 | 140 | 66% | 1e-32 | 30.54% | 474 | WP_238571931.1 |
| cytochrome P450 [Nocardioides thalensis] | Nocardioides thalensis | 140 | 140 | 47% | 2e-32 | 36.28% | 454 | WP_179666022.1 |
| cytochrome P450 [Lentzea californiensis] | Lentzea californiensis | 139 | 139 | 67% | 2e-32 | 27.30% | 442 | WP_258951355.1 |
| cytochrome P450 [Streptomyces platensis] | Streptomyces platensis | 140 | 140 | 66% | 2e-32 | 30.82% | 474 | WSW51259.1 |
| cytochrome P450 [Streptomyces lydicus] | Streptomyces lydicus | 140 | 140 | 66% | 3e-32 | 31.42% | 473 | WP_127150082.1 |
| cytochrome P450 [Streptomyces sp. G7(2002)] | Streptomyces sp. G7(2002) | 139 | 139 | 66% | 3e-32 | 31.42% | 473 | WP_274734702.1 |
| cytochrome P450 [Streptomyces platensis] | Streptomyces platensis | 139 | 139 | 66% | 4e-32 | 30.82% | 474 | WSI58613.1 |
| cytochrome P450 [Streptomyces caniferus] | Streptomyces caniferus | 139 | 139 | 66% | 5e-32 | 30.47% | 473 | WP_159480069.1 |
| cytochrome P450 [Streptomyces libani] | Streptomyces libani | 139 | 139 | 66% | 5e-32 | 31.42% | 473 | WP_159490066.1 |
| cytochrome P450 [Streptomyces sp. Isolate_219] | Streptomyces sp. Isolate_219 | 139 | 139 | 66% | 5e-32 | 31.42% | 473 | WP_262039496.1 |
| cytochrome P450 [Streptomyces caniferus] | Streptomyces caniferus | 139 | 139 | 66% | 5e-32 | 30.47% | 473 | WP_328741856.1 |
| cytochrome P450 [Streptomyces nigrescens] | Streptomyces nigrescens | 139 | 139 | 66% | 6e-32 | 31.42% | 473 | WP_277412529.1 |
| cytochrome P450 [Nocardioides jiangxiensis] | Nocardioides jiangxiensis | 139 | 139 | 36% | 6e-32 | 40.94% | 453 | WP_305026849.1 |

**Figure S4**. BLAST hits for the cytochrome p450 in the putative bursatellin cluster are bacterial upon exclusion of molluscan hits from the nr database.

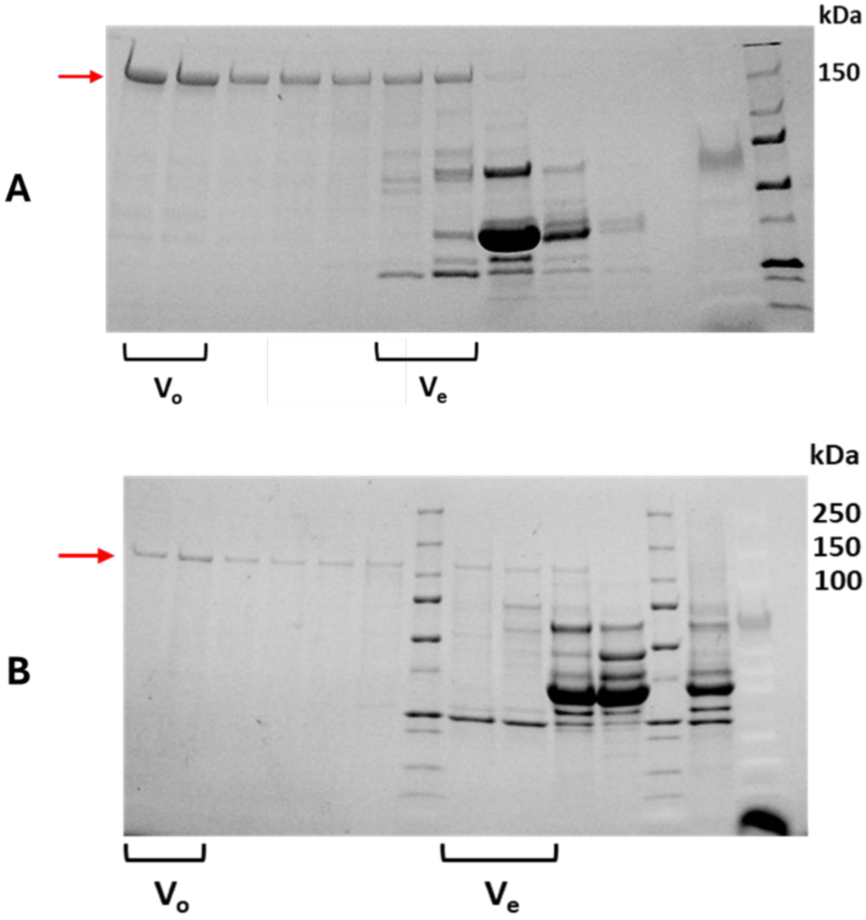


**Figure S5**. Protein fractions from size exclusion column (Superose 6 increase 10/300 GL) visualized on a TGX stain-free 4-20% gradient gel (Bio-Rad). **A)** Fractions of CvFmtATR (calculated molecular weight: 183.8 kDa). **B)** Fractions of CvATR (calculated molecular weight: 119.2 kDa). ($V_o$ = void volume; $V_e$ = expected elution volume).

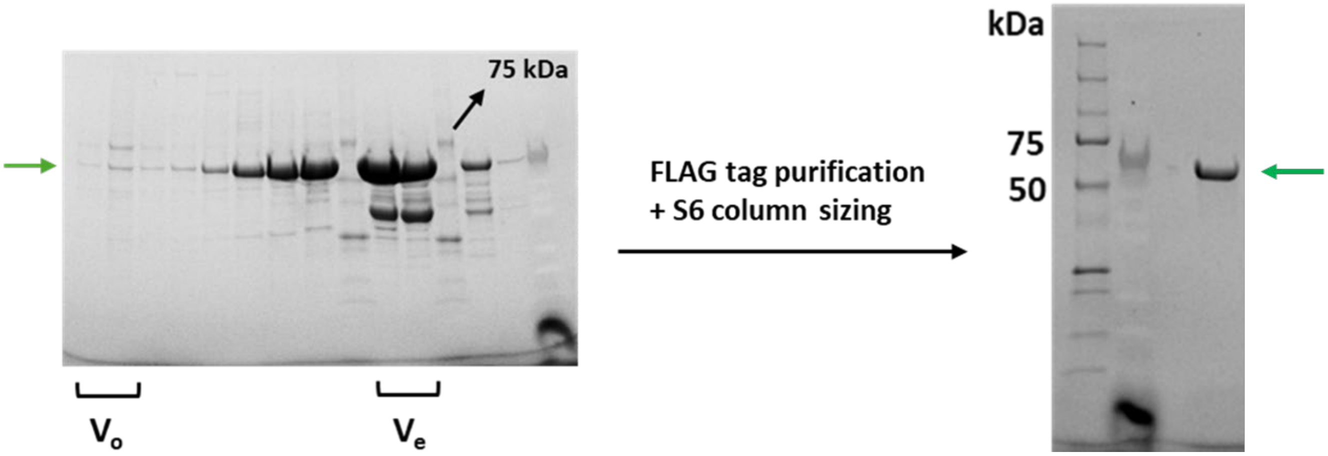


**Figure S6**. Left image - Protein fractions of CvTR (calculated molecular weight: 60.9 kDa) from size exclusion column (Superose 6 increase 10/300 GL) visualized on a TGX stain-free 4-20% gradient gel (Bio-Rad). Right image – CvTR after FLAG tag-based elution and subsequent sizing purification. ($V_o$ = void volume; $V_e$ = expected elution volume).

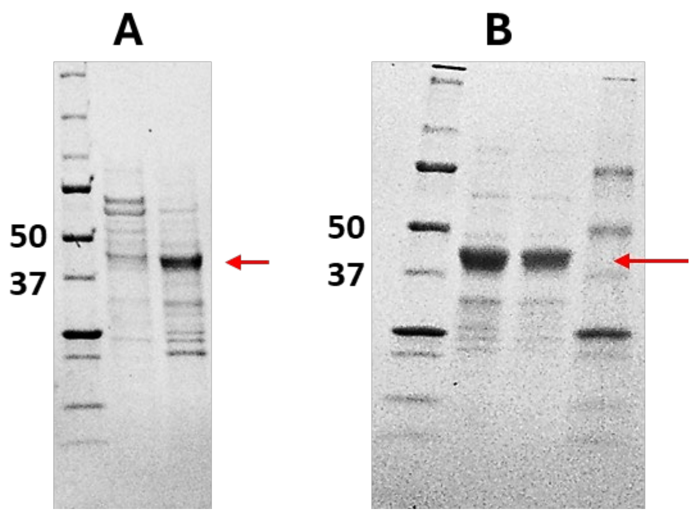


**Figure S7**. McalACT (46.1 kDa) visualized on a TGX stain-free 4-20% gradient gel (Bio-Rad). **A)** McalACT fraction separated on a Superose 6 increase 10/300 GL column (rightmost lane). **B)** Pooled and concentrated McalACT fractions after separation on a HiPrep 26/60 Sephacryl S-200 HR column (the two middle lanes represent the same sample at different loading volumes).

#### 3. Chemical and enzymatic synthesis schemes

**Scheme S1.** Synthetic substrates and standards prepared in this study. All boxed compounds were purified and characterized by NMR prior to use in biochemical experiments.

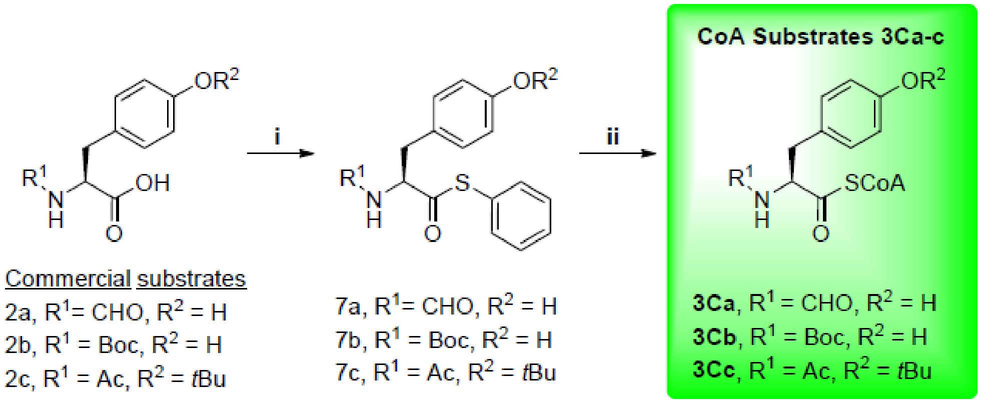


**i:** 2a/2b/2c (1 eq.), DCC (1 eq.), HOBt (1 eq.), thiophenol (1.2 eq.) in ethyl acetate for 22 h.
**ii:** 7a/7b/7c (1 eq.), coenzyme A (1 eq.) in 1:1 v/v MeCN:phosphate buffer (pH 8.0) for 2 h.

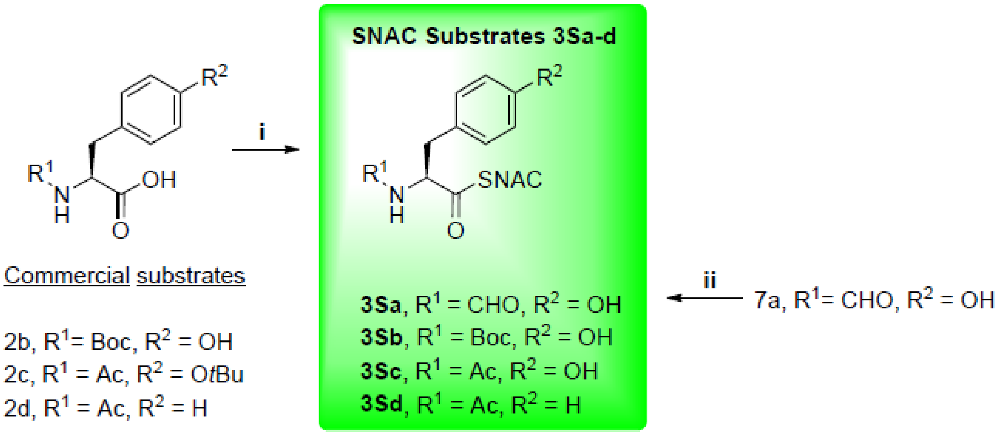


**i:** 2b/2c/2d (1 eq.), DCC (1 eq.), HOBt (1 eq.), $K_2CO_3$ (0.5 eq.), *N*-acetylcysteamine (2 eq. for 2c and 2d and 5 eq. for 2b) in THF for 3 h; *tert*-butyl deprotection in 1 mL of 50% TFA/$CH_2Cl_2$ overnight.
**ii:** 7a (1 eq.), *N*-acetylcysteamine (1 eq.) in 1:1 v/v MeCN:phosphate buffer (pH 8.0) for 2 h.

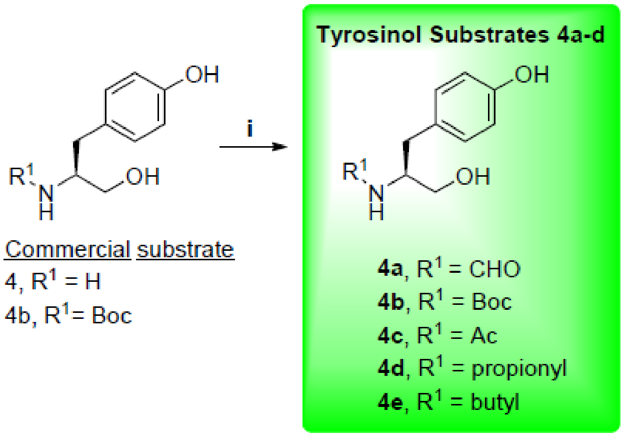


**i:** 4 (1 eq.), $K_2CO_3$ (4 eq.), acyl chloride (1 eq.) in THF:water (2:1 v/v) for 1 h (**4c**, **4d**, and **4e**).

**i:** 4 (1 eq.), *N*-formylsaccharin (1.5 eq.), $K_2CO_3$ (2.5 eq.) in THF:water (1:1 v/v) for 1 h (for **4a**).

**Scheme S2.** Characterization of enzymatically prepared tyrosinol derivatives. Each substrate was individually reacted with McalACT, and compound **5b** was purified and characterized by NMR. Characteristic $MS^2$ fragmentation patterns in comparison to those of **5b** confirmed the remaining products.

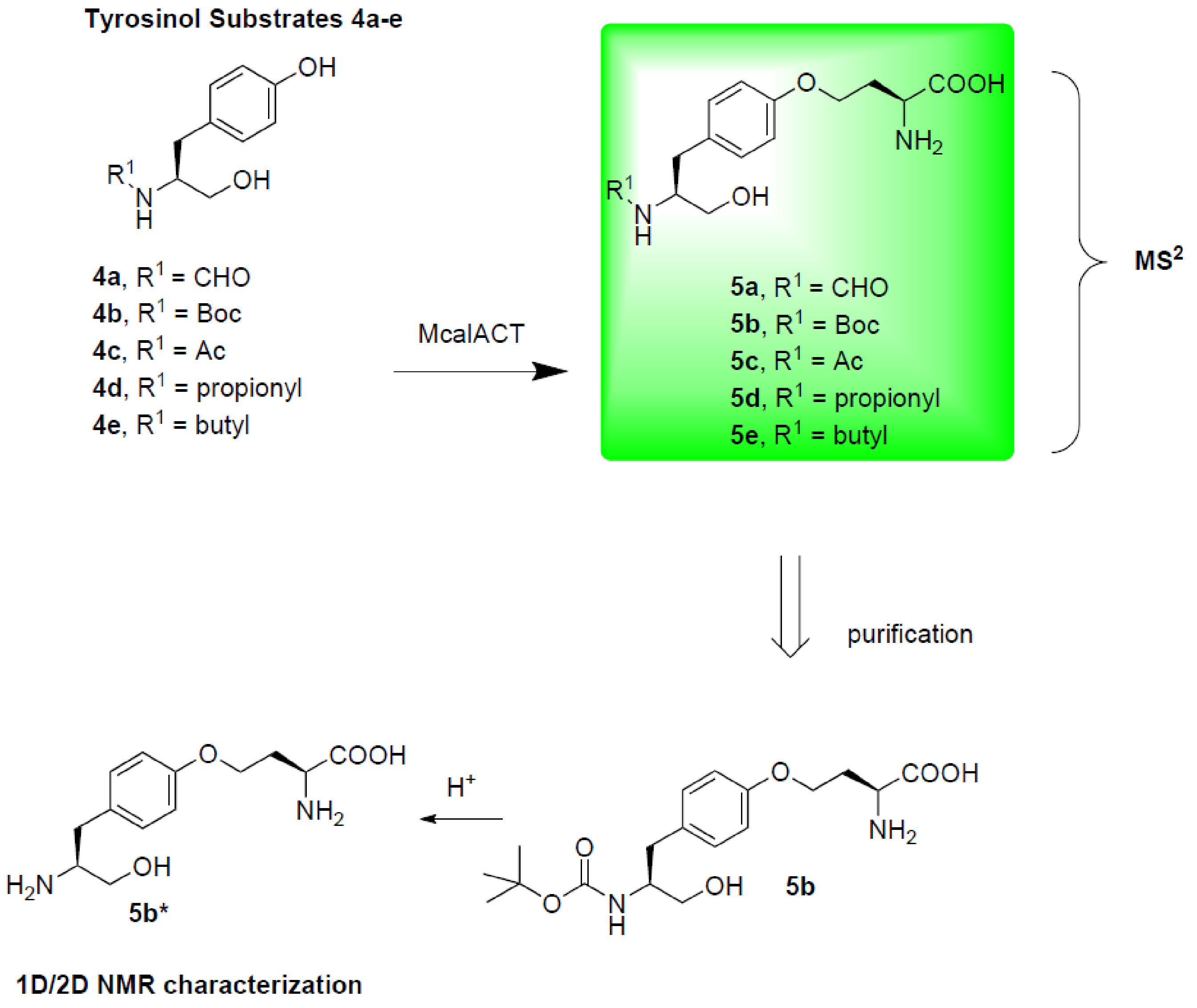

#### 4. Enzyme assay summary scheme and raw data

**Scheme S3.** Summary of CvTR assay results as a heatmap of respective product molecular ion peak intensity. As mass peak intensity is affected by the polarity and ionizability of a compound as well as the assay matrix, this heatmap is qualitative and cannot be used for comparing enzyme activity against different substrates.

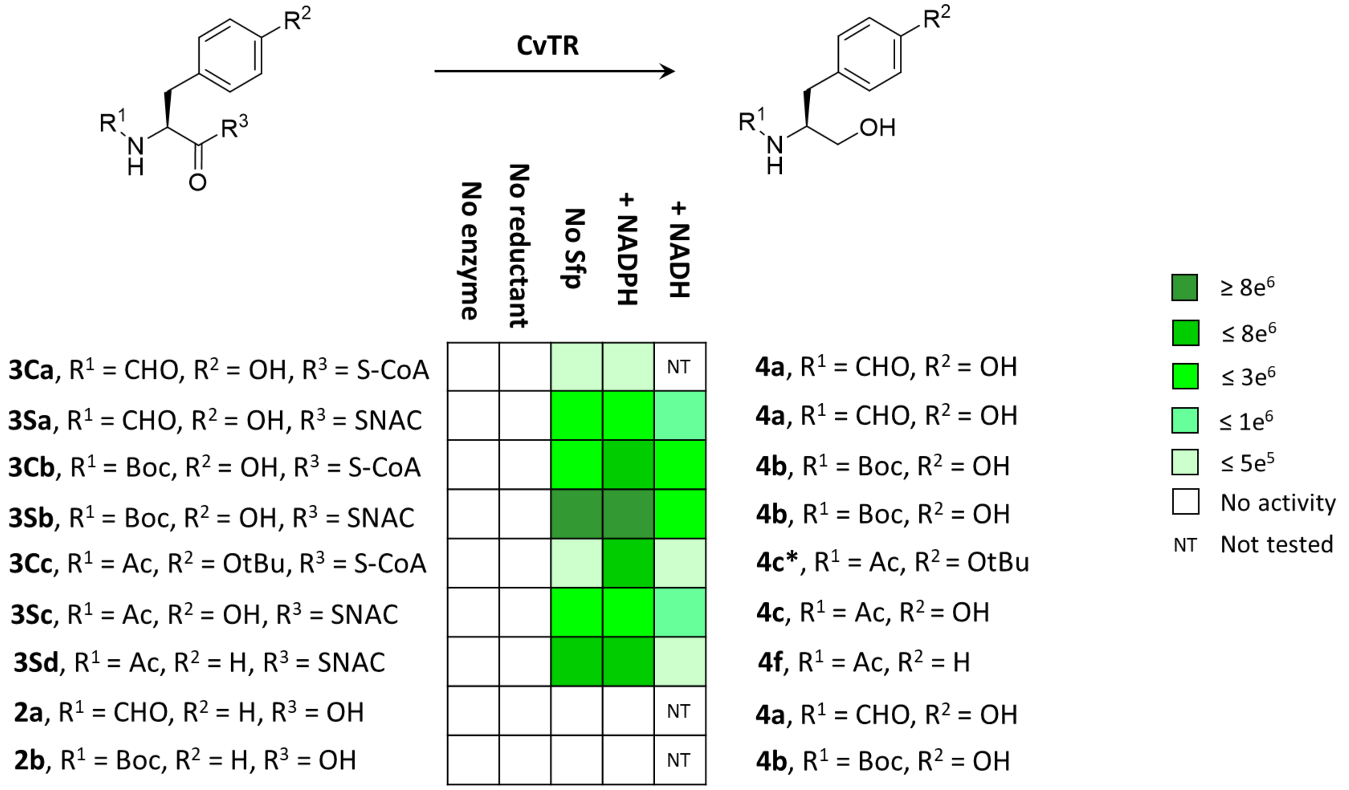

**Scheme S4.** Summary of McalACT assay results as a heatmap of respective product molecular ion peak intensity. As mass peak intensity is affected by the polarity and ionizability of a compound as well as the assay matrix, this heatmap is qualitative and cannot be used for comparing enzyme activity against different substrates.

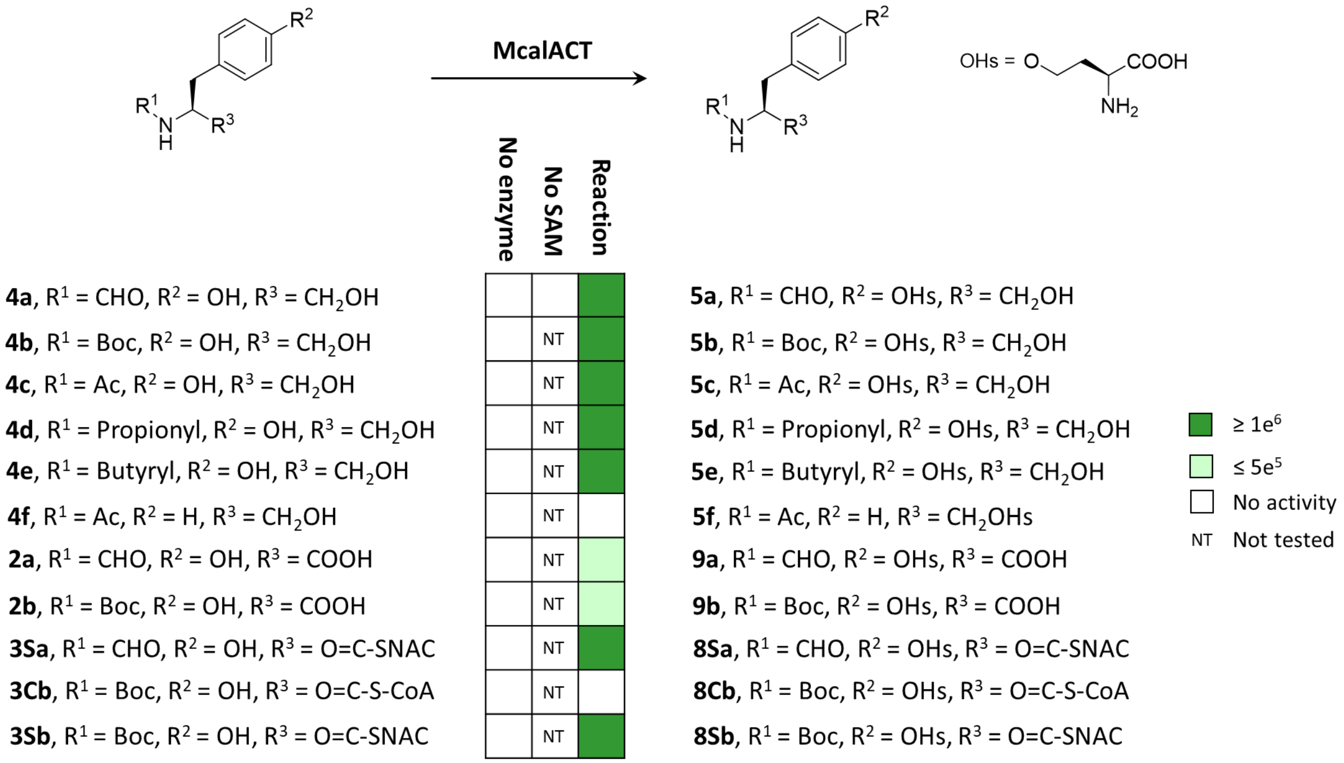


**Scheme S5.** Summary of results of one-pot assays containing both CvTR and McalACT.

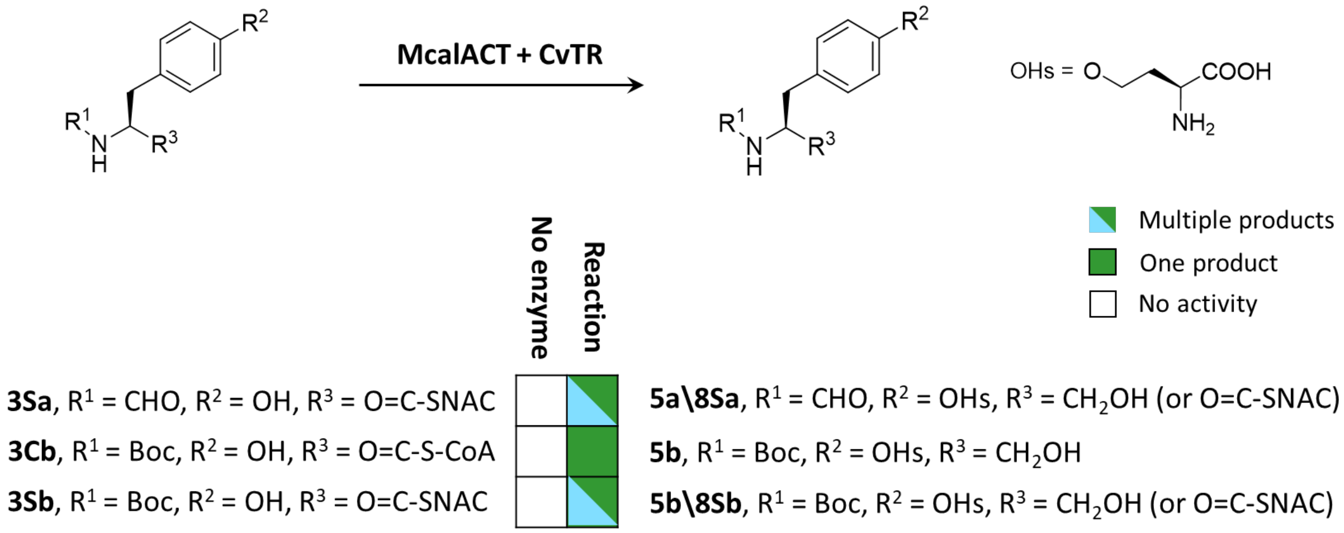

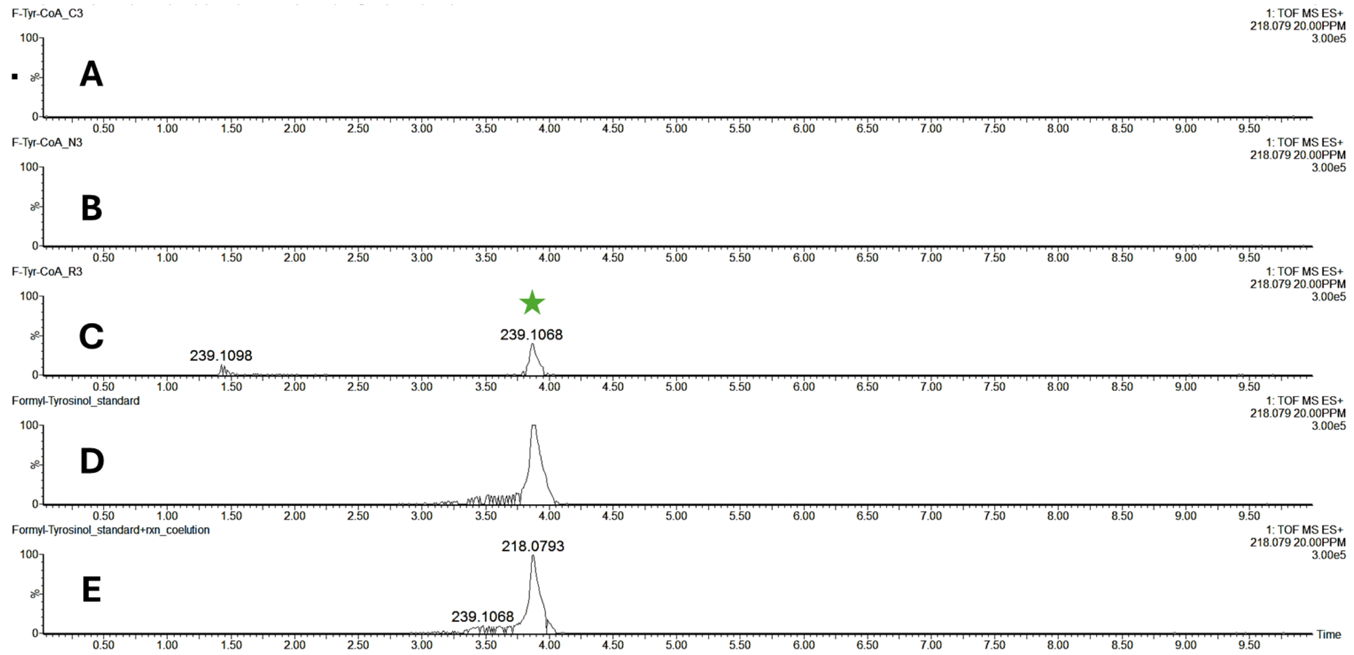


**Figure S8**. CvTR reduces *N*-formyl-L-tyrosine CoA (**3Ca**) in vitro. UPLC-MS extracted ion chromatograms of **A)** Substrate control containing Sfp (1 μM), **3Ca** (200 μM), NADPH (500 μM), and $MgCl_2$ (10 mM). **B)** No NADPH control containing CvTR (10 μM), Sfp (1 μM), **3Ca** (200 μM), and $MgCl_2$ (10 mM). **C)** Reaction mixture containing CvTR (10 μM), Sfp (1 μM), **3Ca** (200 μM), NADPH (500 μM), and $MgCl_2$ (10 mM). **D)** Synthetic standard of **4a**. **E)** Coelution of samples C + D.

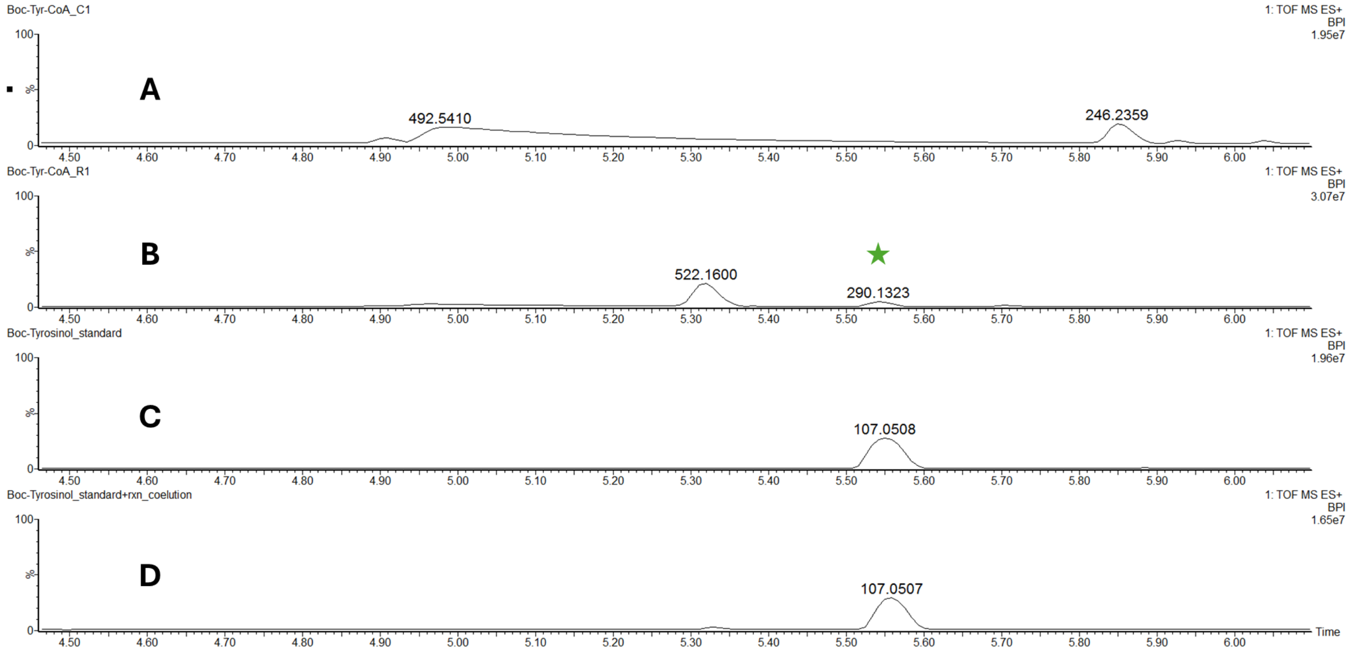


**Figure S9.** CvTR reduces *N*-Boc-L-tyrosine CoA (**3Cb**) in vitro. Zoomed-in UPLC-MS base peak intensity chromatograms of **A)** Substrate control containing Sfp (1 µM), **3Cb** (200 µM), NADPH (500 µM), and $MgCl_2$ (10 mM). **B)** Reaction mixture containing CvTR (10 µM), Sfp (1 µM), **3Cb** (200 µM), NADPH (500 µM), and $MgCl_2$ (10 mM). **C)** Synthetic standard of **4b**. **D)** Coelution of samples B + C.

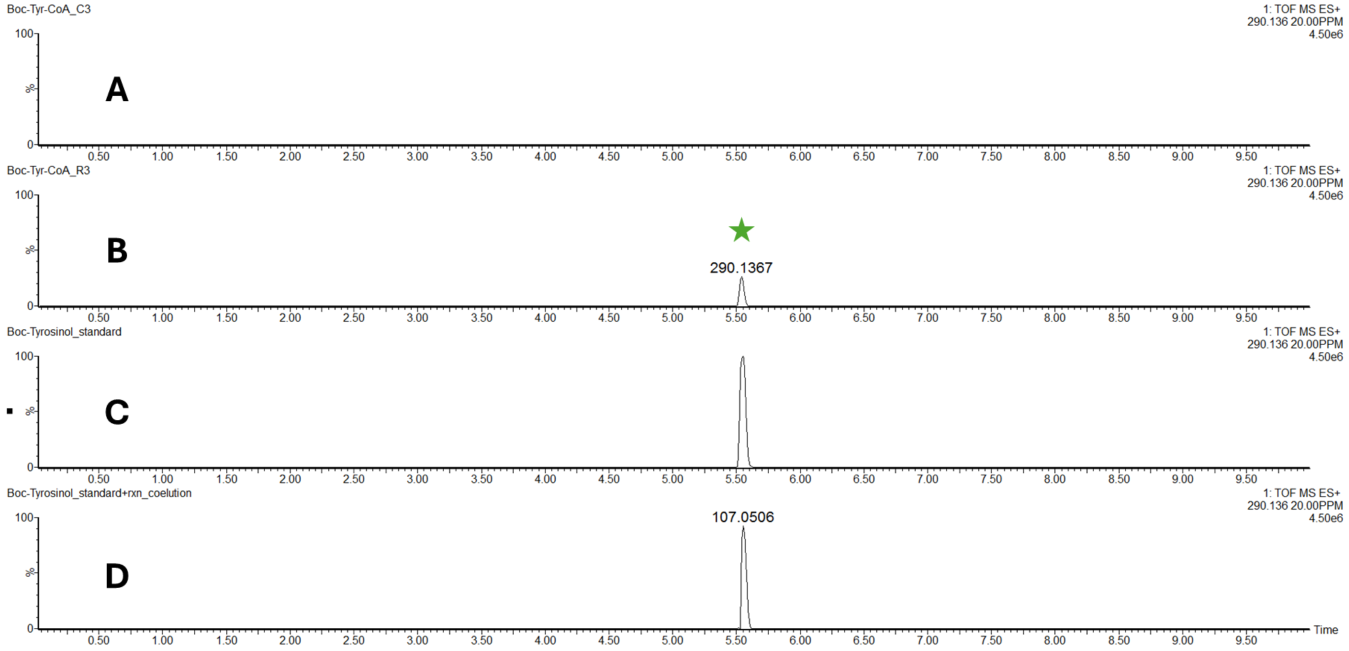


**Figure S10.** CvTR reduces *N*-Boc-L-tyrosine CoA (**3Cb**) in vitro. UPLC-MS extracted ion chromatograms of **A)** Substrate control containing Sfp (1 µM), **3Cb** (200 µM), NADPH (500 µM), and $MgCl_2$ (10 mM). **B)** Reaction mixture containing CvTR (10 µM), Sfp (1 µM), **3Cb** (200 µM), NADPH (500 µM), and $MgCl_2$ (10 mM). **C)** Synthetic standard of **4b**. **D)** Coelution of samples B + C.

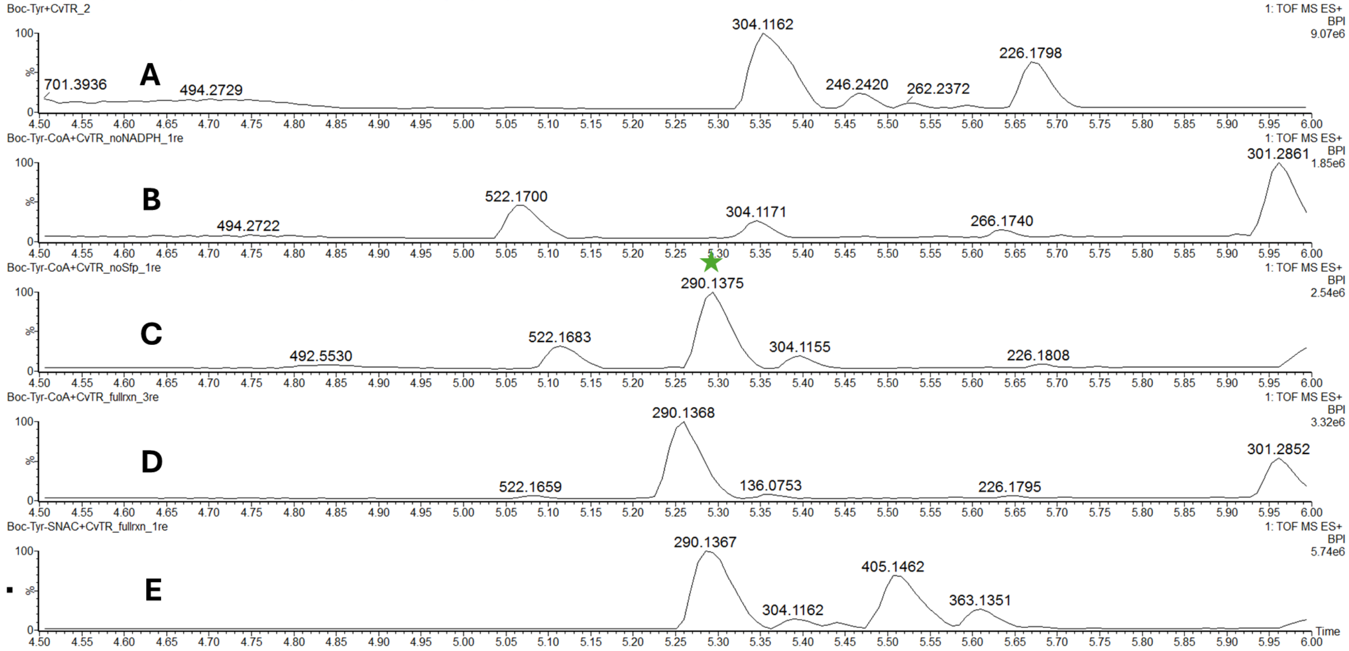


**Figure S11.** CvTR functions independent of Sfp activity to reduce thioester substrates but not free amino acids in vitro. UPLC-MS base peak intensity chromatograms of **A)** Reaction mixture containing CvTR (10 μM), Sfp (10 μM), *N*-Boc-L-tyrosine (**2b**) (100 μM), NADPH (1 mM), and $MgCl_2$ (10 mM). **B)** No NADPH control containing CvTR (10 μM), Sfp (10 μM), **3Cb** (100 μM), and $MgCl_2$ (10 mM). **C)** No Sfp control containing CvTR (10 μM), **3Cb** (100 μM), NADPH (1 mM), and $MgCl_2$ (10 mM). **D)** Reaction mixture containing CvTR (10 μM), Sfp (10 μM), **3Cb** (100 μM), NADPH (1 mM), and $MgCl_2$ (10 mM). **E)** Reaction mixture containing CvTR (10 μM), **3Sb** (100 μM), NADPH (1 mM), and $MgCl_2$ (10 mM).

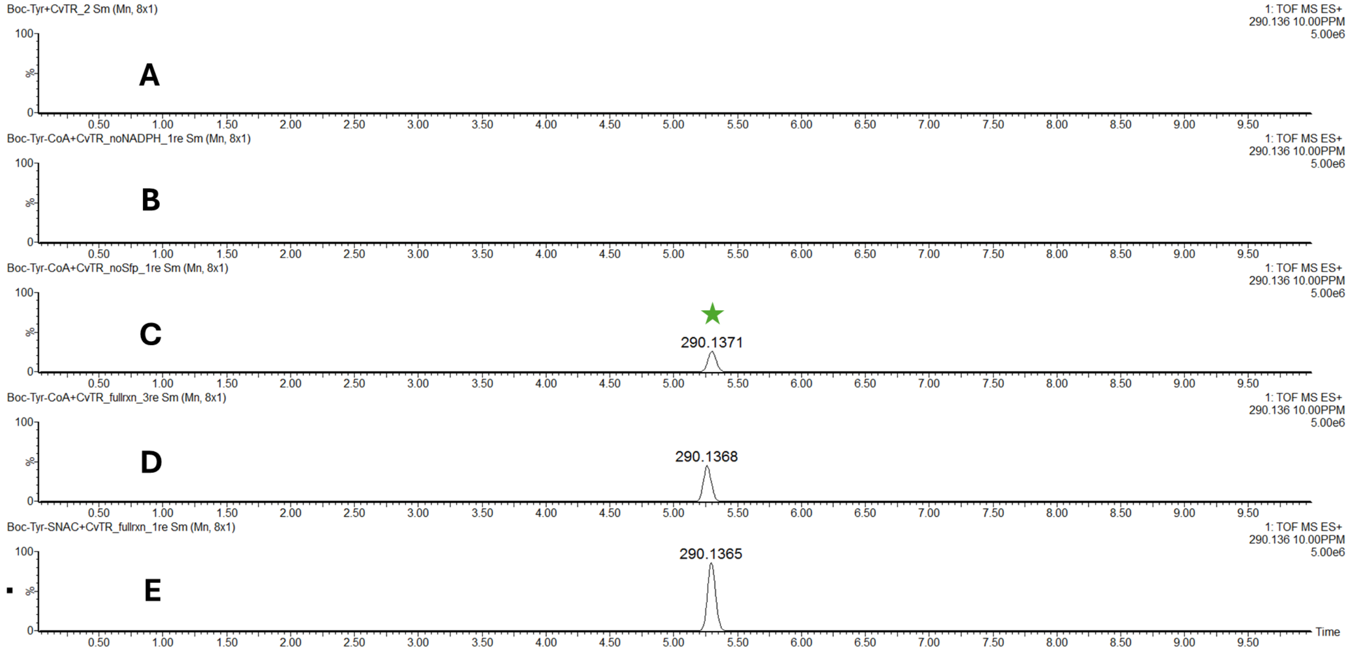


**Figure S12.** CvTR functions independent of Sfp activity to reduce thioester substrates but not free amino acids in vitro. UPLC-MS extracted ion chromatograms of **A)** Reaction mixture containing CvTR (10 μM), Sfp (10 μM), *N*-Boc-L-tyrosine (**2b**) (100 μM), NADPH (1 mM), and $MgCl_2$ (10 mM). **B)** No NADPH control containing CvTR (10 μM), Sfp (10 μM), **3Cb** (100 μM), and $MgCl_2$ (10 mM). **C)** No Sfp control containing CvTR (10 μM), **3Cb** (100 μM), NADPH (1 mM), and $MgCl_2$ (10 mM). **D)** Reaction mixture containing CvTR (10 μM), Sfp (10 μM), **3Cb** (100 μM), NADPH (1 mM), and $MgCl_2$ (10 mM). **E)** Reaction mixture containing CvTR (10 μM), **3Sb** (100 μM), NADPH (1 mM), and $MgCl_2$ (10 mM).

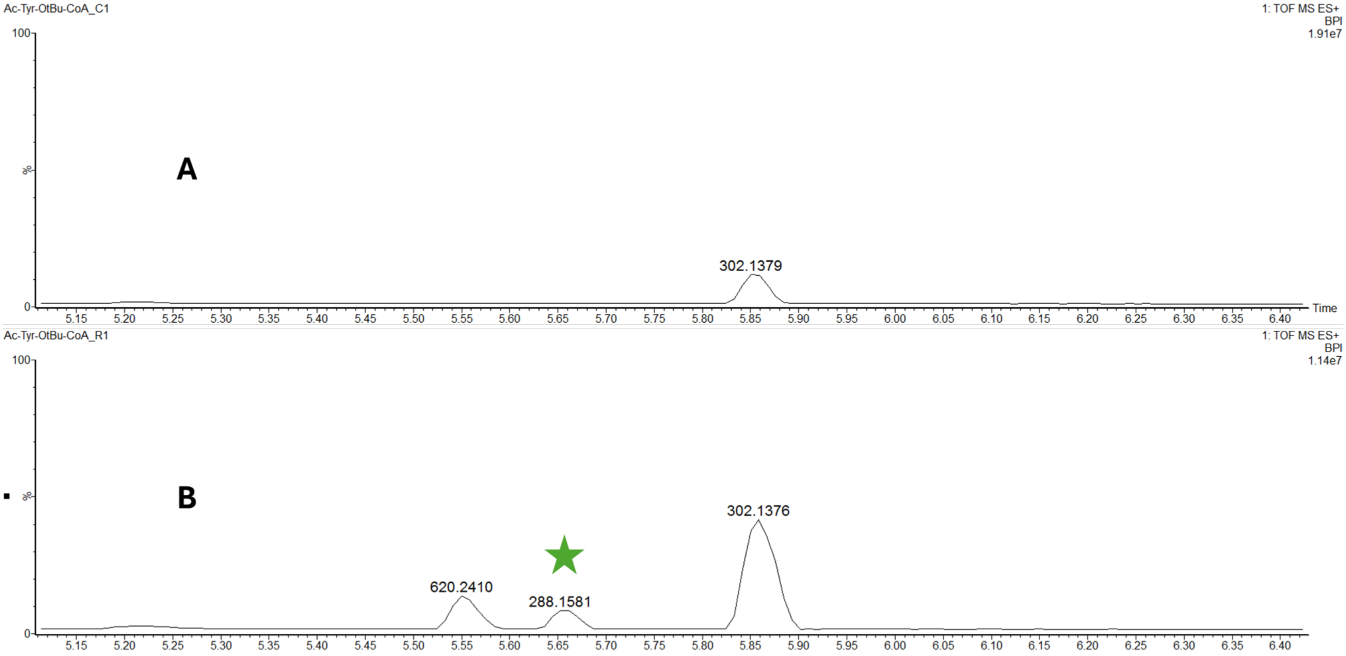


**Figure S13**. CvTR reduces *N*-acetyl-O-*tert*-butyl-L-tyrosine CoA (**3Cc**) in vitro. Zoomed-in UPLC-MS base peak intensity chromatograms of **A)** Substrate control containing Sfp (1 µM), **3Cc** (200 µM), NADPH (500 µM), and $MgCl_2$ (10 mM). **B)** Reaction mixture containing CvTR (10 µM), Sfp (1 µM), **3Cc** (200 µM), NADPH (500 µM), and $MgCl_2$ (10 mM).

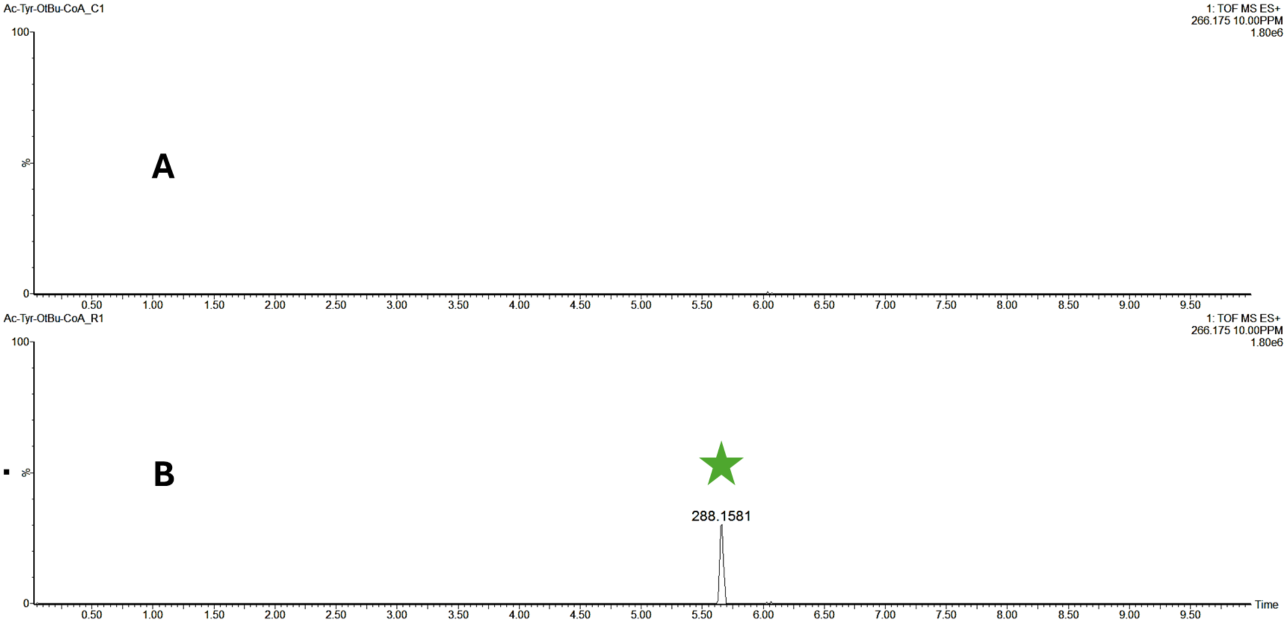


**Figure S14**. CvTR reduces *N*-acetyl-O-*tert*-butyl-L-tyrosine CoA (**3Cc**) in vitro. UPLC-MS extracted ion chromatograms of **A)** Substrate control containing Sfp (1 µM), **3Cc** (200 µM), NADPH (500 µM), and $MgCl_2$ (10 mM). **B)** Reaction mixture containing CvTR (10 µM), Sfp (1 µM), **3Cc** (200 µM), NADPH (500 µM), and $MgCl_2$ (10 mM).

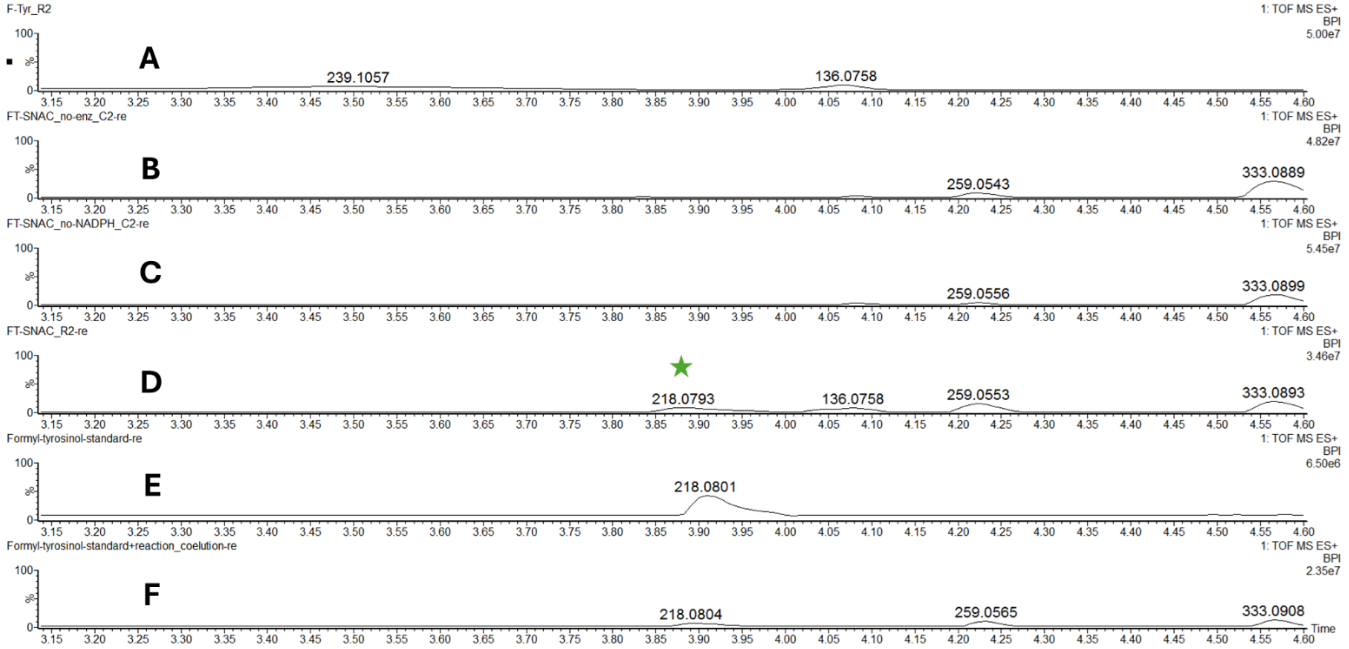


**Figure S15**. CvTR reduces *N*-formyl-L-tyrosine SNAC (**3Sa**) in vitro. Zoomed-in UPLC-MS base peak intensity chromatograms of **A)** Reaction mixture containing CvTR (10 μM), **2a** (500 μM), NADPH (500 μM), and $MgCl_2$ (10 mM). **B)** Substrate control containing **3Sa** (500 μM), NADPH (500 μM), and $MgCl_2$ (10 mM). **C)** No NADPH control containing CvTR (10 μM), **3Sa** (500 μM), and $MgCl_2$ (10 mM). **D)** Reaction mixture containing CvTR (10 μM), **3Sa** (500 μM), NADPH (500 μM), and $MgCl_2$ (10 mM). **E)** Synthetic standard of **4a**. **F)** Coelution of samples D + E.

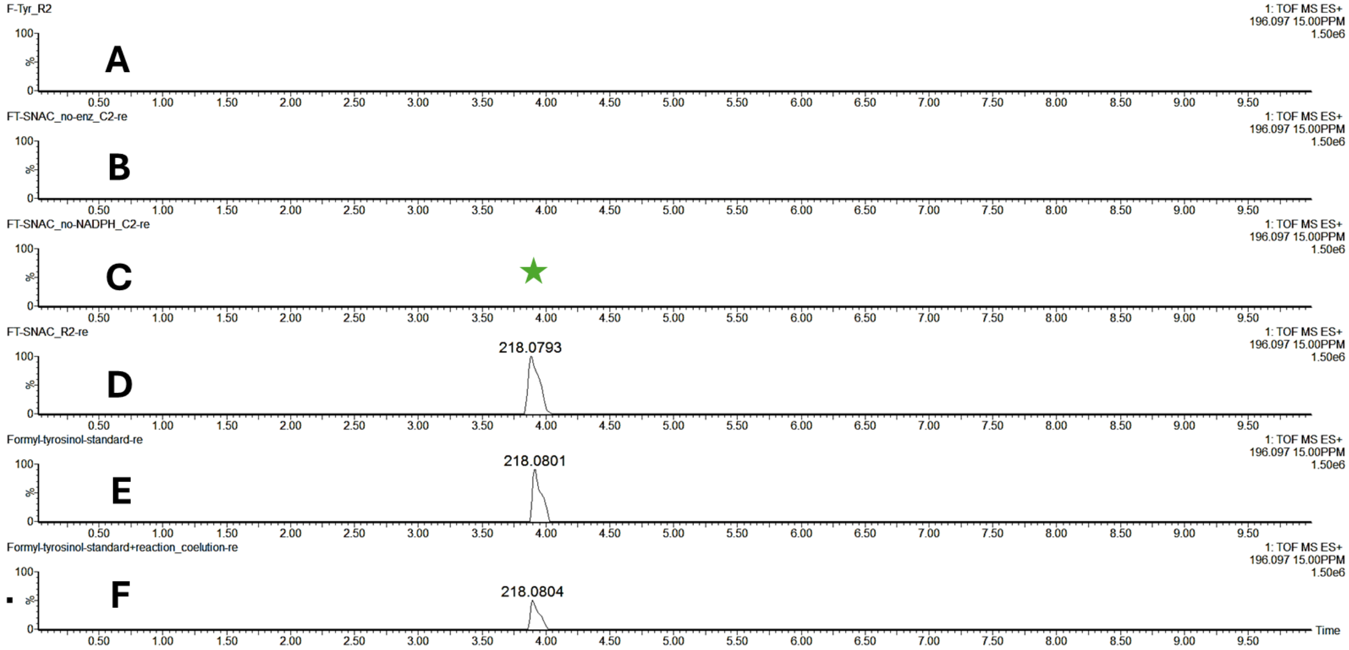


**Figure S16**. CvTR reduces *N*-formyl-L-tyrosine SNAC (**3Sa**) in vitro. UPLC-MS extracted ion chromatograms of **A)** Reaction mix-turecontaining CvTR (10 μM), **2a** (500 μM), NADPH (500 μM), and $MgCl_2$ (10 mM). **B)** Substrate control containing **3Sa** (500 μM), NADPH (500 μM), and $MgCl_2$ (10 mM). C**)** No NADPH control containing CvTR (10 μM), **3Sa** (500 μM), and $MgCl_2$ (10 mM). **D)** Reaction mixture containing CvTR (10 μM), **3Sa** (500 μM), NADPH (500 μM), and $MgCl_2$ (10 mM). **E)** Synthetic standard of **4a**. **F)** Coelution of samples D + E.

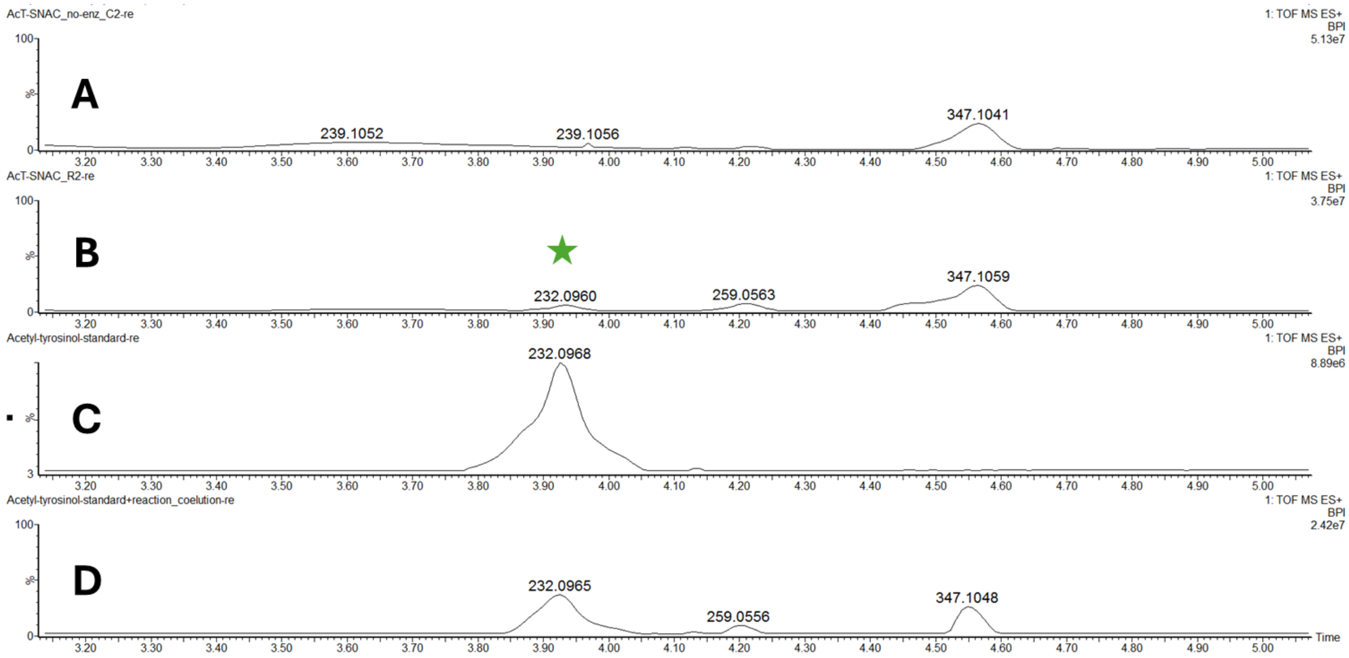


**Figure S17**. CvTR reduces *N*-acetyl-L-tyrosine SNAC (**3Sc**) in vitro. Zoomed-in UPLC-MS base peak intensity chromatograms of **A)** Substrate control containing **3Sc** (500 μM), NADPH (500 μM), and $MgCl_2$ (10 mM). **B)** Reaction mixture containing CvTR (10 μM), **3Sc** (500 μM), NADPH (500 μM), and $MgCl_2$ (10 mM). **C)** Synthetic standard of **4c**. **D)** Coelution of samples B + C.

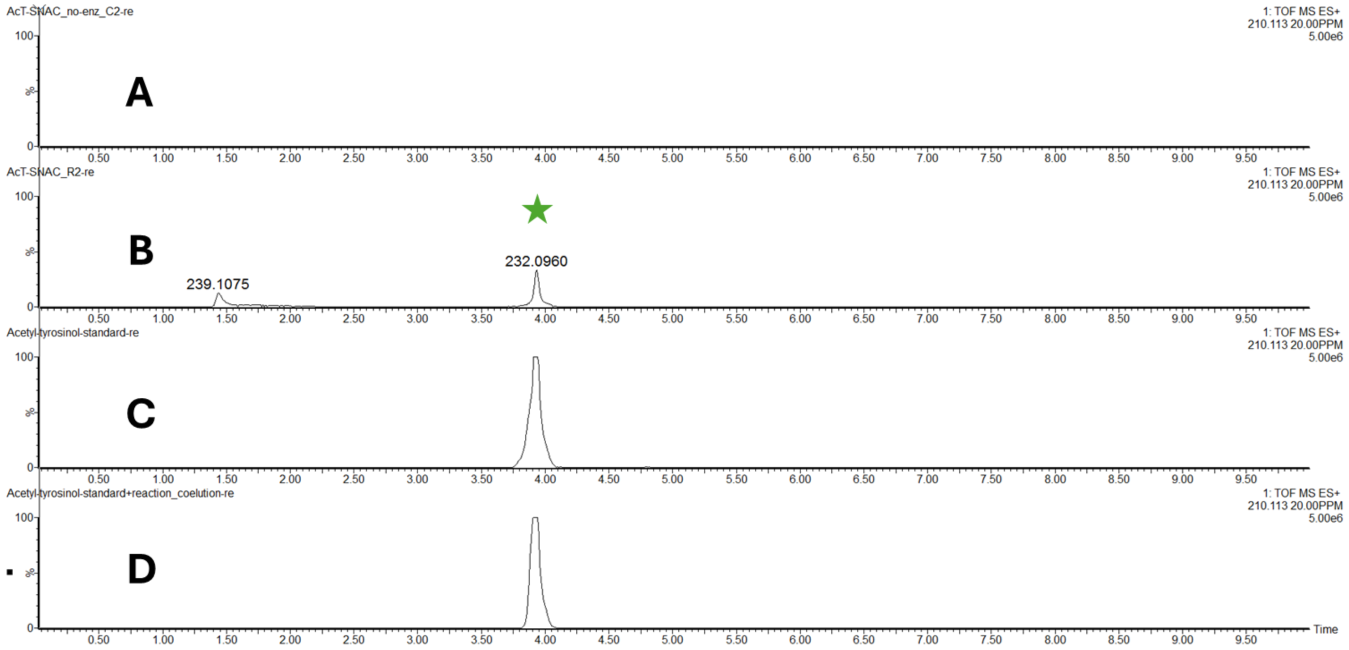


**Figure S18**. CvTR reduces *N*-acetyl-L-tyrosine SNAC (**3Sc**) in vitro. UPLC-MS extracted ion chromatograms of **A)** Substrate control containing **3Sc** (500 μM), NADPH (500 μM), and $MgCl_2$ (10 mM). **B)** Reaction mixture containing CvTR (10 μM), **3Sc** (500 μM), NADPH (500 μM), and $MgCl_2$ (10 mM). **C)** Synthetic standard of **4c**. **D)** Coelution of samples B + C.

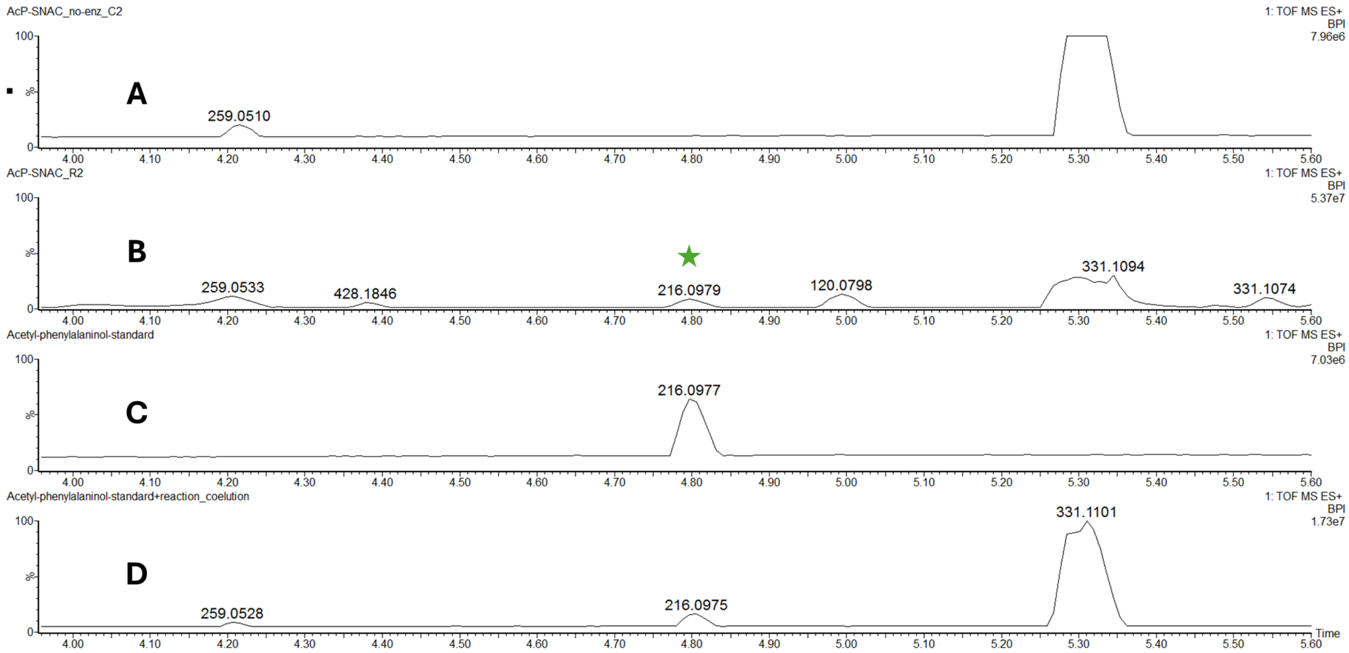


**Figure S19**. CvTR reduces *N*-acetyl-L-phenylalanine SNAC (**3Sd**) in vitro. Zoomed-in UPLC-MS base peak intensity chromatograms of **A)** Substrate control containing **3Sd** (500 μM), NADPH (500 μM), and $MgCl_2$ (10 mM). **B)** Reaction mixture containing CvTR (10 μM), **3Sd** (500 μM), NADPH (500 μM), and $MgCl_2$ (10 mM). **C)** Synthetic standard of **4f**. **D)** Coelution of samples B + C.

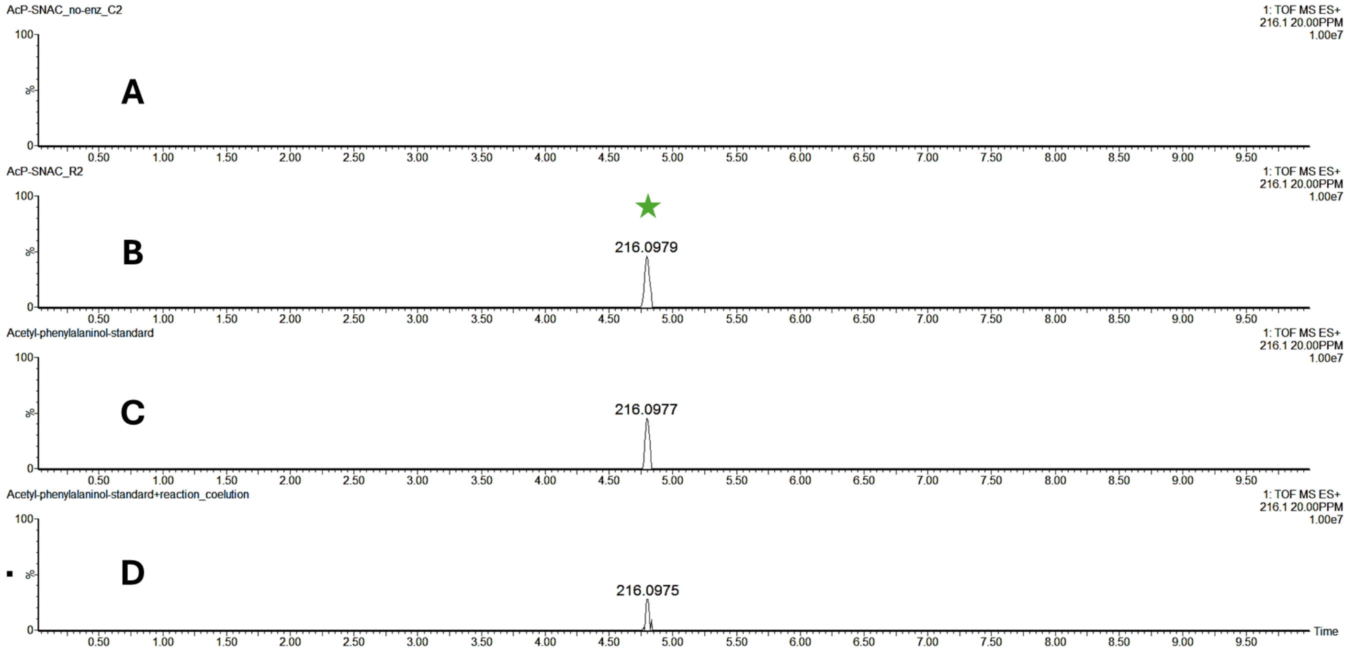


**Figure S20**. CvTR reduces *N*-acetyl-L-phenylalanine SNAC (**3Sd**) in vitro. UPLC-MS extracted ion chromatograms of **A)** Substrate control containing **3Sd** (500 μM), NADPH (500 μM), and $MgCl_2$ (10 mM). **B)** Reaction mixture containing CvTR (10 μM), **3Sd** (500 μM), NADPH (500 μM), and $MgCl_2$ (10 mM). **C)** Synthetic standard of **4f**. **D)** Coelution of samples B + C.

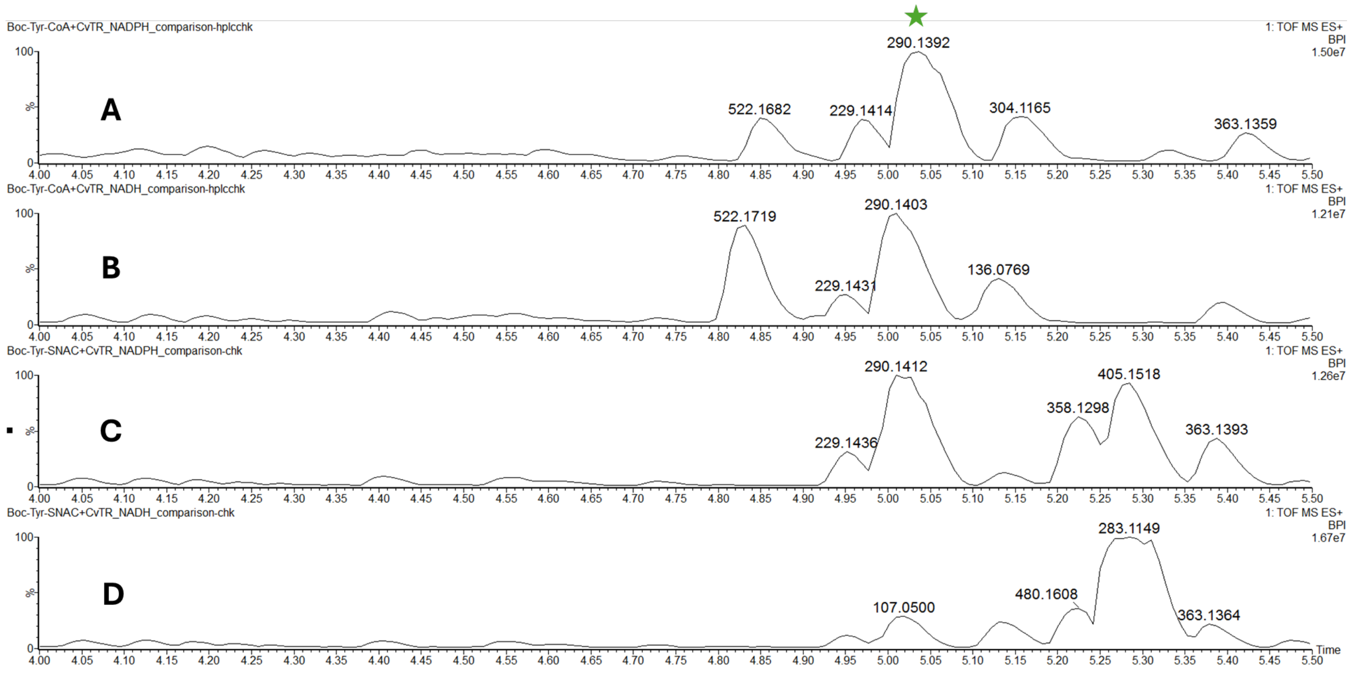


**Figure S21.** CvTR uses NADH as an alternative reductant. UPLC-MS base peak intensity chromatograms of **A)** Reaction mixture containing CvTR (10 µM), Sfp (10 µM), **3Cb** (100 µM), NADPH (1mM), and $MgCl_2$ (10 mM). **B)** Reaction mixture containing CvTR (10 µM), Sfp (10 µM), **3Cb** (100 µM), NADH (1mM), and $MgCl_2$ (10 mM). **C)** Reaction mixture containing CvTR (10 µM), **3Sb** (100 µM), NADPH (1mM), and $MgCl_2$ (10 mM). **D)** Reaction mixture containing CvTR (10 µM), **3Sb** (100 µM), NADH (1mM), and $MgCl_2$ (10 mM).

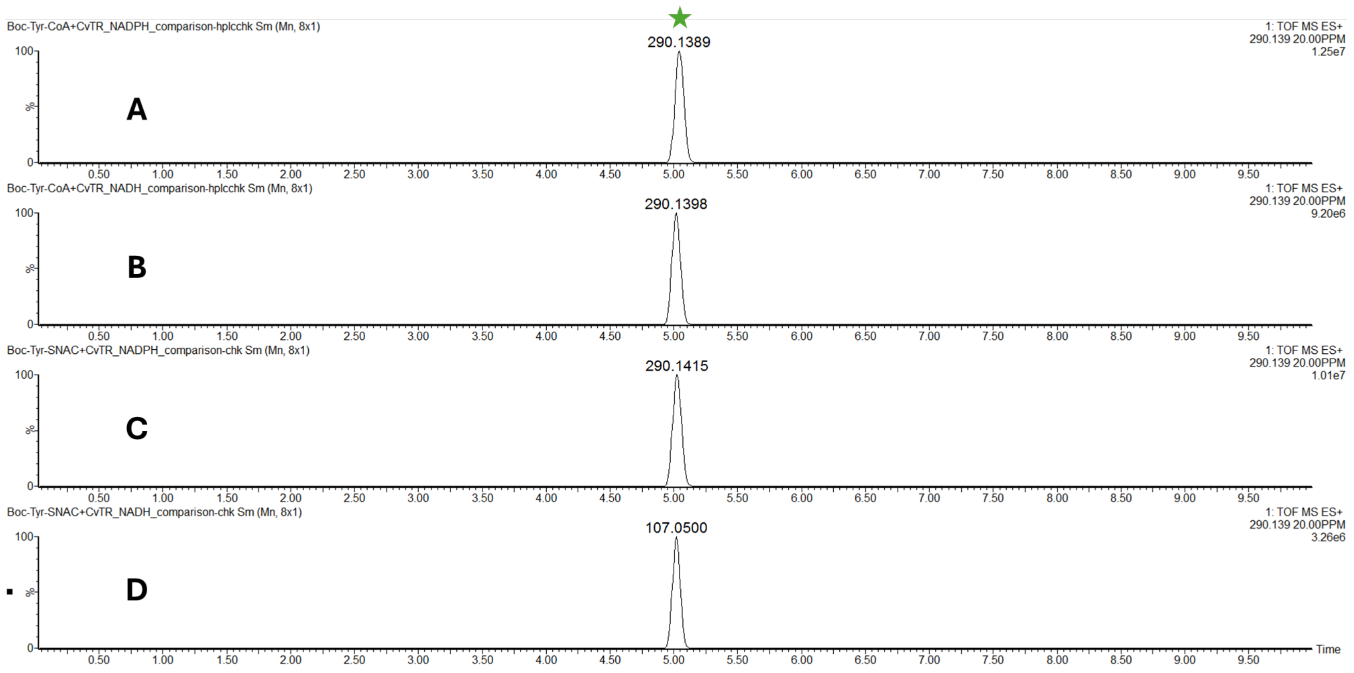


**Figure S22.** CvTR uses NADH as an alternative reductant. UPLC-MS extracted ion chromatograms of **A)** Reaction mixture containing CvTR (10 µM), Sfp (10 µM), **3Cb** (100 µM), NADPH (1mM), and $MgCl_2$ (10 mM). **B)** Reaction mixture containing CvTR (10 µM), Sfp (10 µM), **3Cb** (100 µM), NADH (1mM), and $MgCl_2$ (10 mM). **C)** Reaction mixture containing CvTR (10 µM), **3Sb** (100 µM), NADPH (1mM), and $MgCl_2$ (10 mM). **D)** Reaction mixture containing CvTR (10 µM), **3Sb** (100 µM), NADH (1mM), and $MgCl_2$ (10 mM).

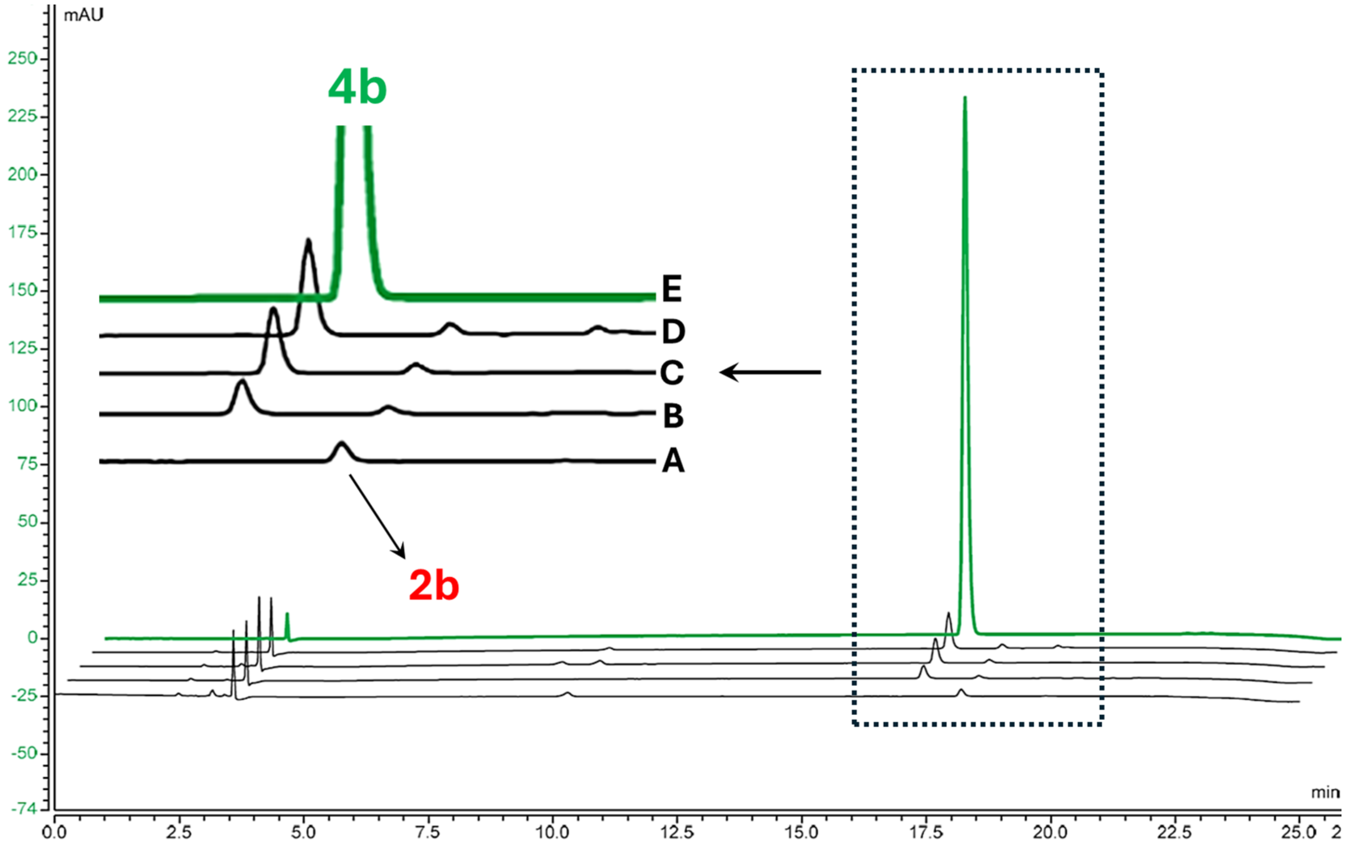


**Figure S23**. HPLC-UV chromatograms at 280 nm of **A)** No NADPH control containing CvTR (10 µM), Sfp (10 µM), **3Cb** (100 µM), and $MgCl_2$ (10 mM). **B)** No Sfp control containing CvTR (10 µM), **3Cb** (100 µM), NADPH (1 mM), and $MgCl_2$ (10 mM). **C)** Reaction mixture containing CvTR (10 µM), Sfp (10 µM), **3Cb** (100 µM), NADPH (1 mM), and $MgCl_2$ (10 mM). **D)** Reaction mixture containing CvTR (10 µM), **3Sb** (100 µM), NADPH (1 mM), and $MgCl_2$ (10 mM). **E)** Synthetic standard of **4b**.

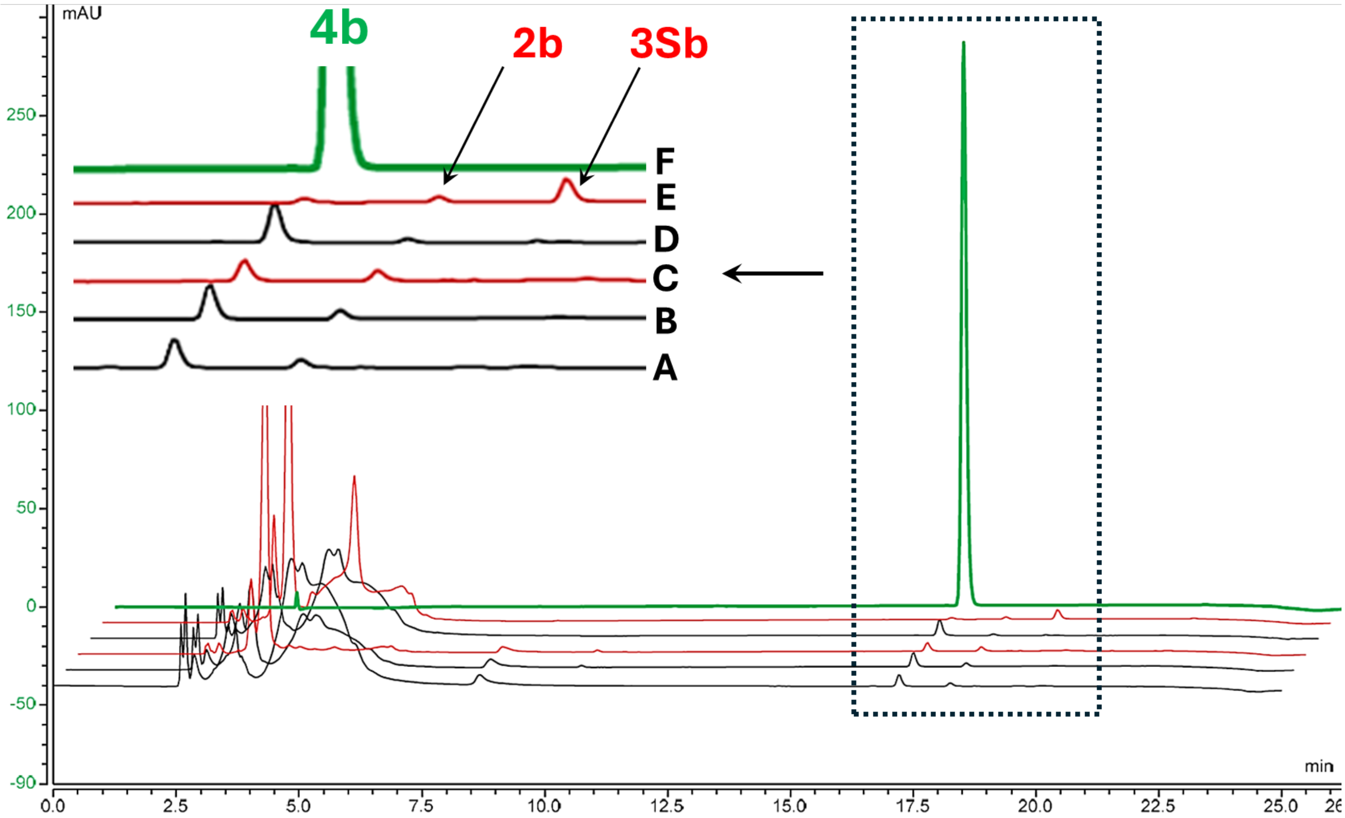


**Figure S24**. HPLC-UV chromatograms at 280 nm of **A)** No Sfp control containing CvTR (10 μM), **3Cb** (100 μM), NADPH (1 mM), and $MgCl_2$ (10 mM). **B)** Reaction mixture containing CvTR (10 μM), Sfp (10 μM), **3Cb** (100 μM), NADPH (1 mM), and $MgCl_2$ (10 mM). **C)** Reaction mixture containing CvTR (10 μM), Sfp (10 μM), **3Cb** (100 μM), NADH (1 mM), and $MgCl_2$ (10 mM). **D)** Reaction mixture containing CvTR (10 μM), **3Sb** (100 μM), NADPH (1 mM), and $MgCl_2$ (10 mM). **E)** Reaction mixture containing CvTR (10 μM), **3Sb** (100 μM), NADH (1 mM), and $MgCl_2$ (10 mM). **F)** Synthetic standard of **4b**.

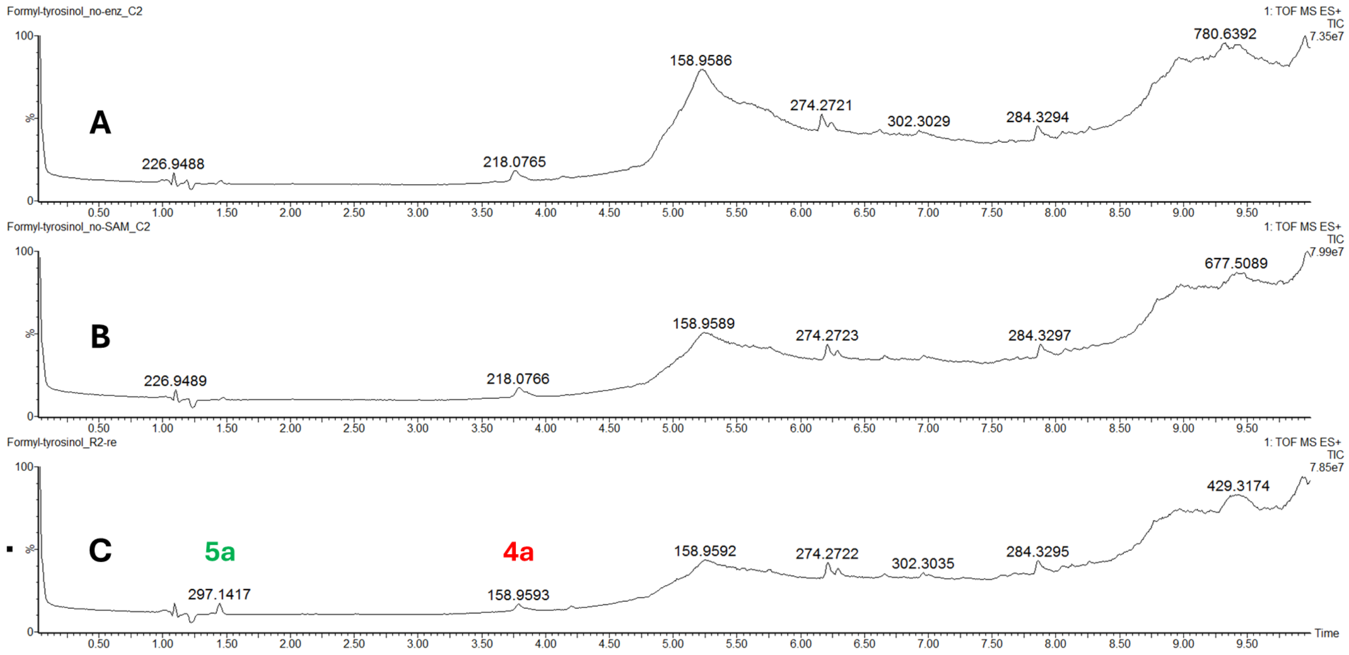


**Figure S25.** McalACT catalyzes aminocarboxypropylation of *N*-formyl-L-tyrosinol (**4a**) in vitro. UPLC-MS total ion chromatograms of **A)** Substrate control containing **4a** (1 mM), SAM (1 mM), and $MgCl_2$ (10 mM). **B)** No SAM control containing McalACT (10 µM), **4a** (1 mM), and $MgCl_2$ (10 mM). **C)** Reaction mixture containing McalACT (10 µM), **4a** (1 mM), SAM (1 mM), and $MgCl_2$ (10 mM).

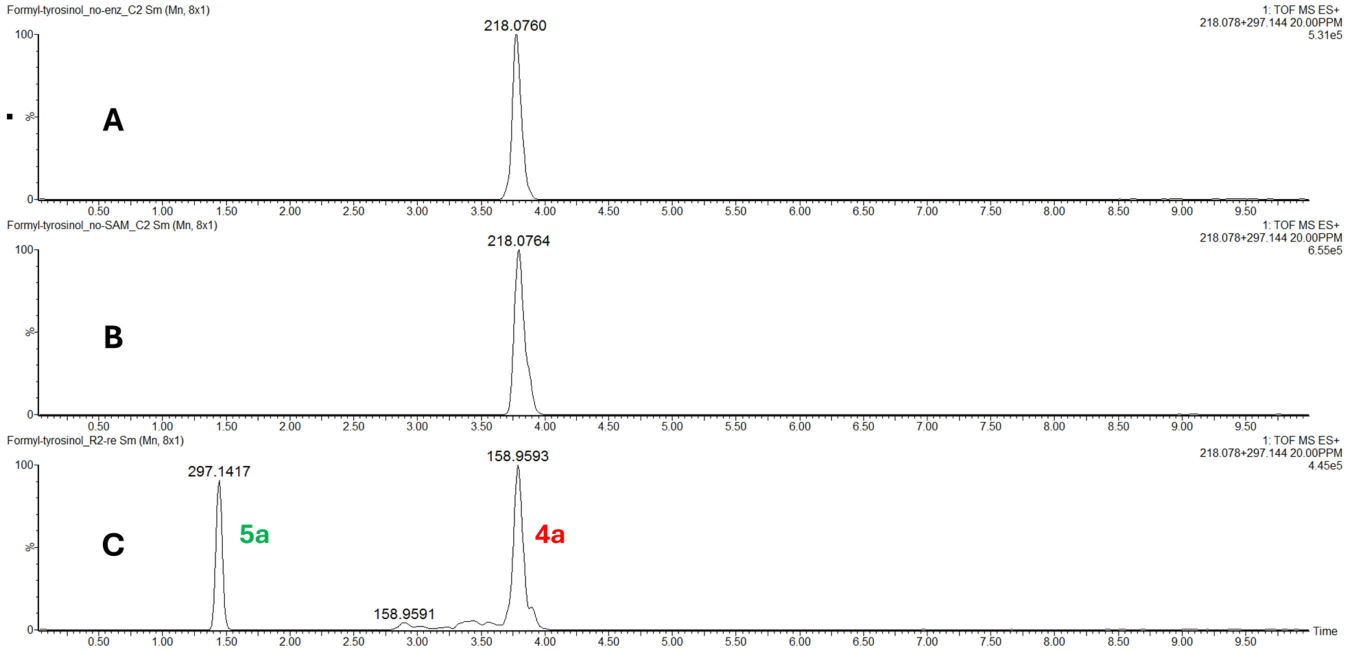


**Figure S26.** McalACT catalyzes aminocarboxypropylation of *N*-formyl-L-tyrosinol (**4a**) in vitro. UPLC-MS extracted ion chromatograms of **A)** Substrate control containing **4a** (1 mM), SAM (1 mM), and $MgCl_2$ (10 mM). **B)** No SAM control containing McalACT (10 µM), **4a** (1 mM), and $MgCl_2$ (10 mM). **C)** Reaction mixture containing McalACT (10 µM), **4a** (1 mM), SAM (1 mM), and $MgCl_2$ (10 mM).

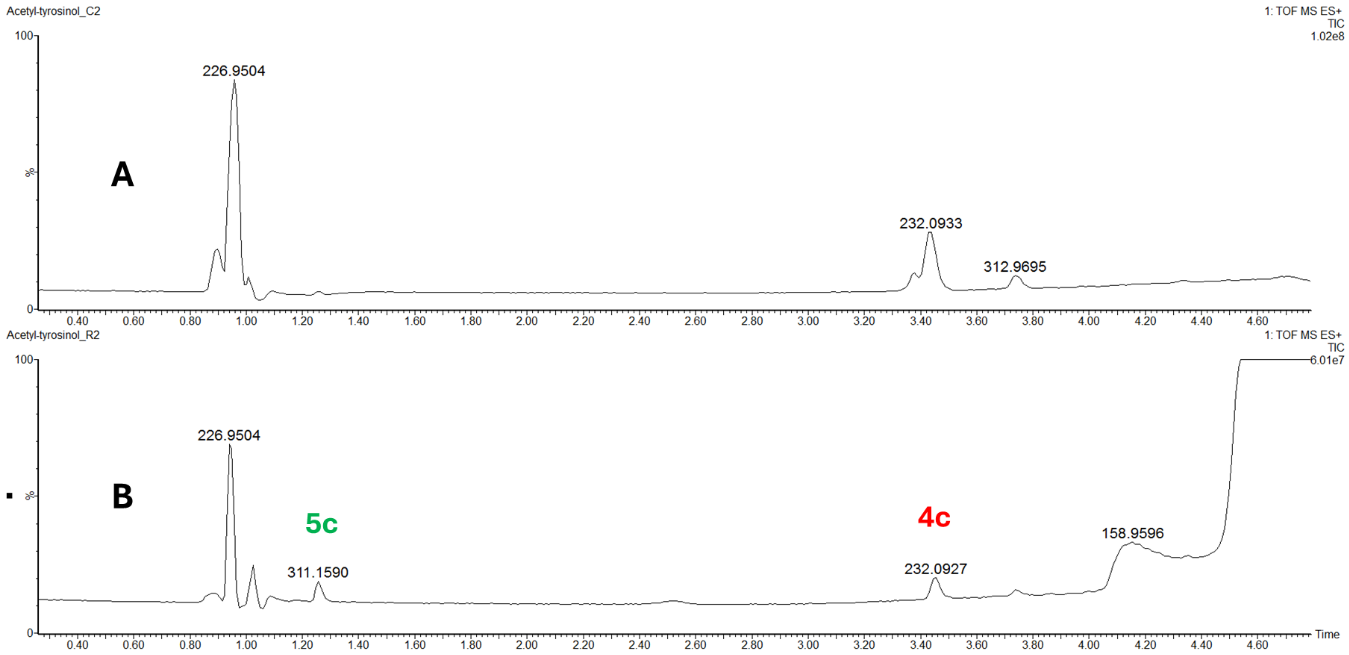


**Figure S27.** McalACT catalyzes aminocarboxypropylation of *N*-acetyl-L-tyrosinol (**4c**) in vitro. Zoomed-in UPLC-MS total ion chromatograms of **A)** Substrate control containing **4c** (1 mM), SAM (1 mM), and $MgCl_2$ (10 mM). **B)** Reaction mixture containing McalACT (10 µM), **4c** (1 mM), SAM (1 mM), and $MgCl_2$ (10 mM).

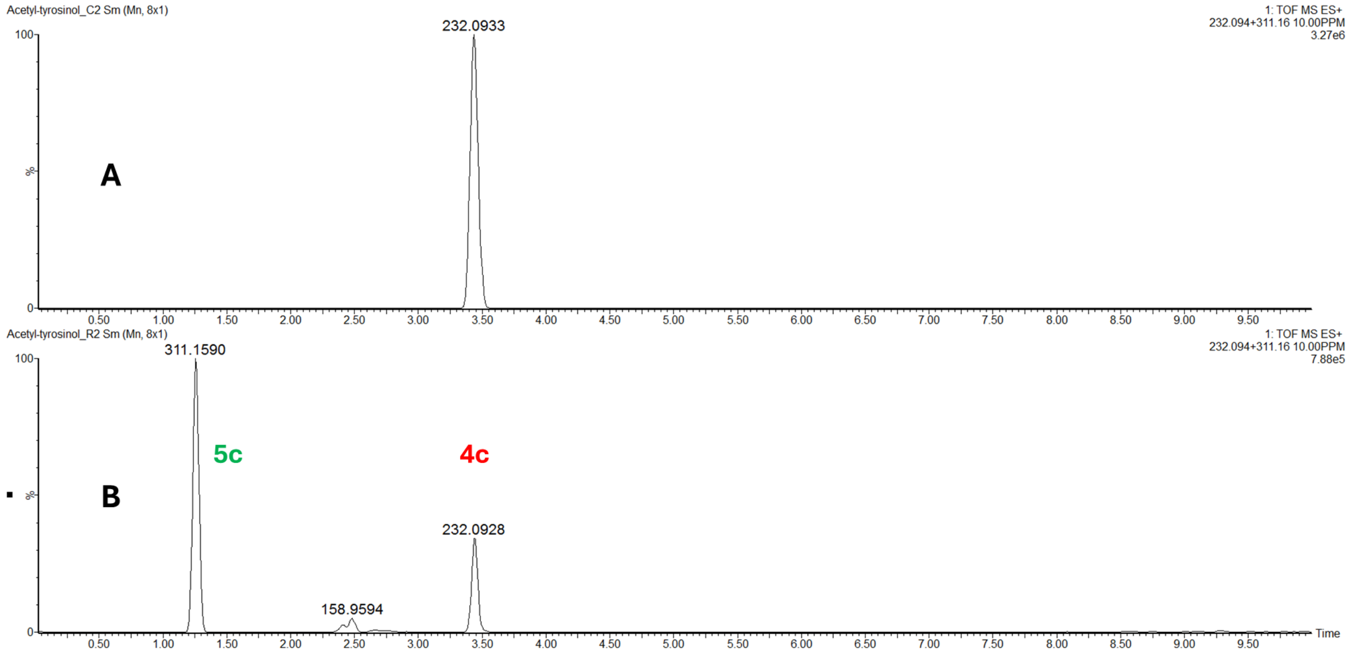


**Figure S28.** McalACT catalyzes aminocarboxypropylation of *N*-acetyl-L-tyrosinol (**4c**) in vitro. UPLC-MS extracted ion chromatograms of **A)** Substrate control containing **4c** (1 mM), SAM (1 mM), and $MgCl_2$ (10 mM). **B)** Reaction mixture containing McalACT (10 µM), **4c** (1 mM), SAM (1 mM), and $MgCl_2$ (10 mM).

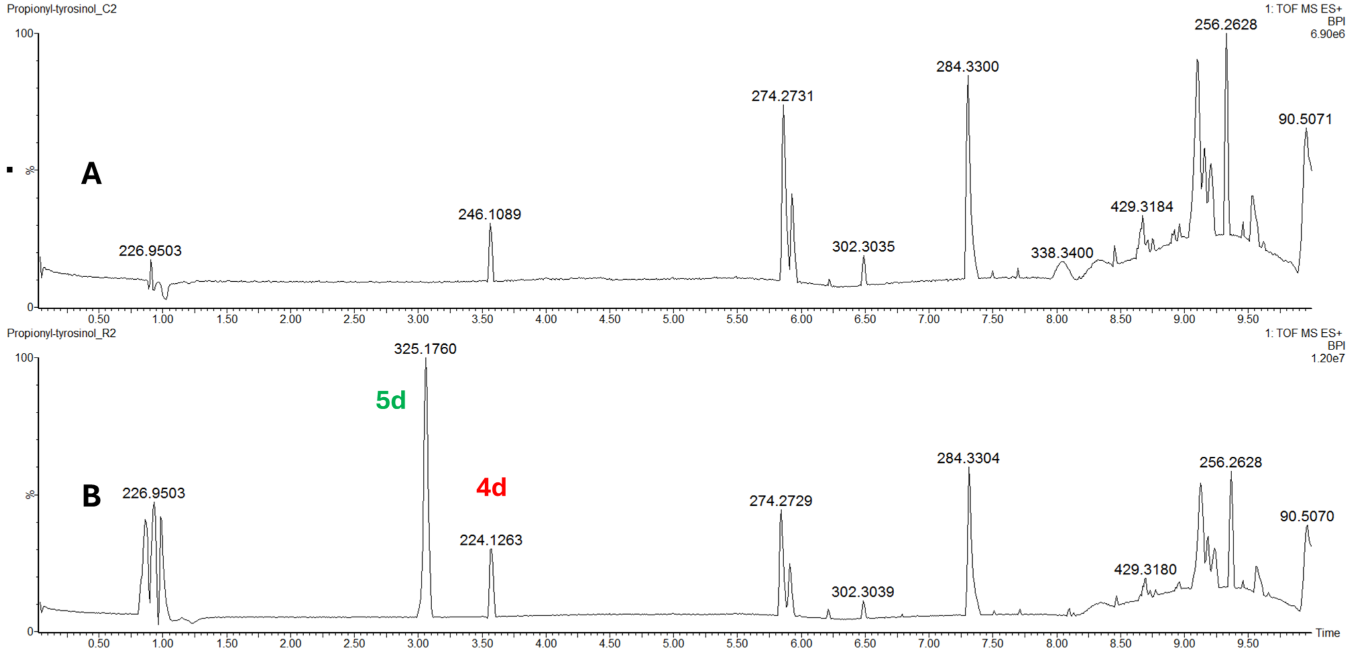


**Figure S29.** McalACT catalyzes aminocarboxypropylation of *N*-propionyl-L-tyrosinol (**4d**) in vitro. UPLC-MS base peak intensity chromatograms of **A)** Substrate control containing **4d** (1 mM), SAM (1 mM), and $MgCl_2$ (10 mM). **B)** Reaction mixture containing McalACT (10 μM), **4d** (1 mM), SAM (1 mM), and $MgCl_2$ (10 mM).

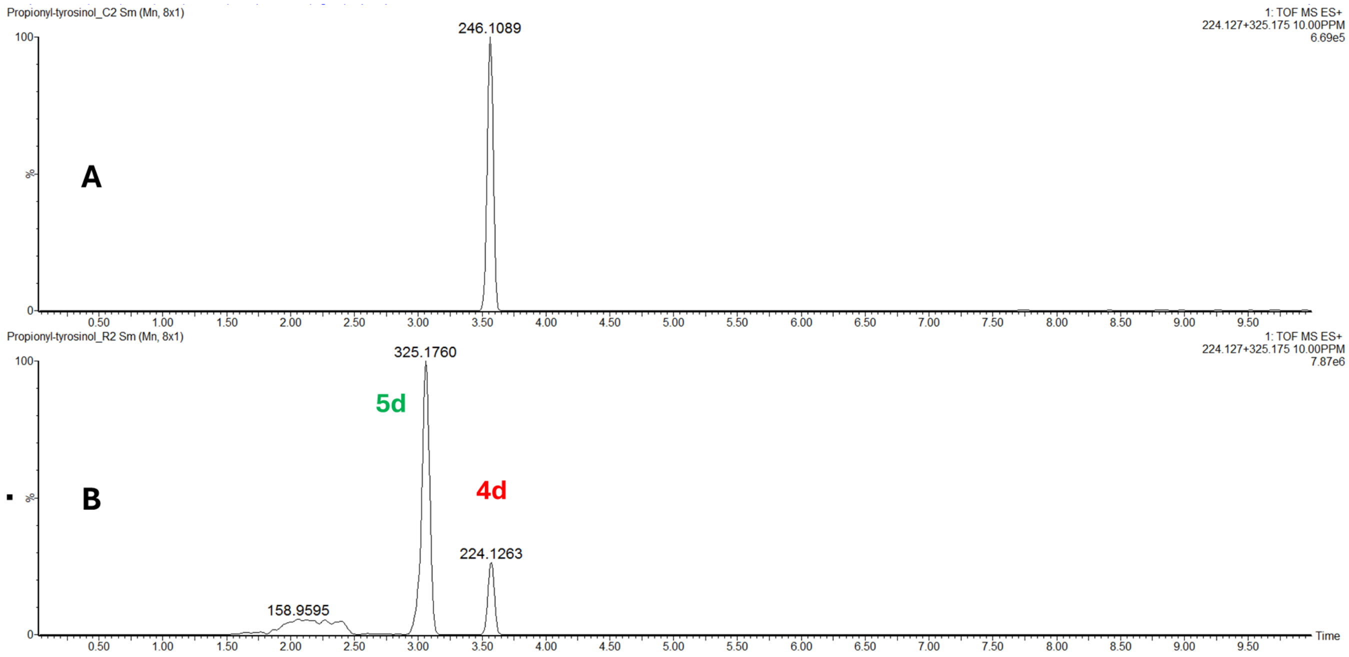


**Figure S30.** McalACT catalyzes aminocarboxypropylation of *N*-propionyl-L-tyrosinol (**4d**) in vitro. UPLC-MS extracted ion chromatograms of **A)** Substrate control containing **4d** (1 mM), SAM (1 mM), and $MgCl_2$ (10 mM). **B)** Reaction mixture containing McalACT (10 μM), **4d** (1 mM), SAM (1 mM), and $MgCl_2$ (10 mM).

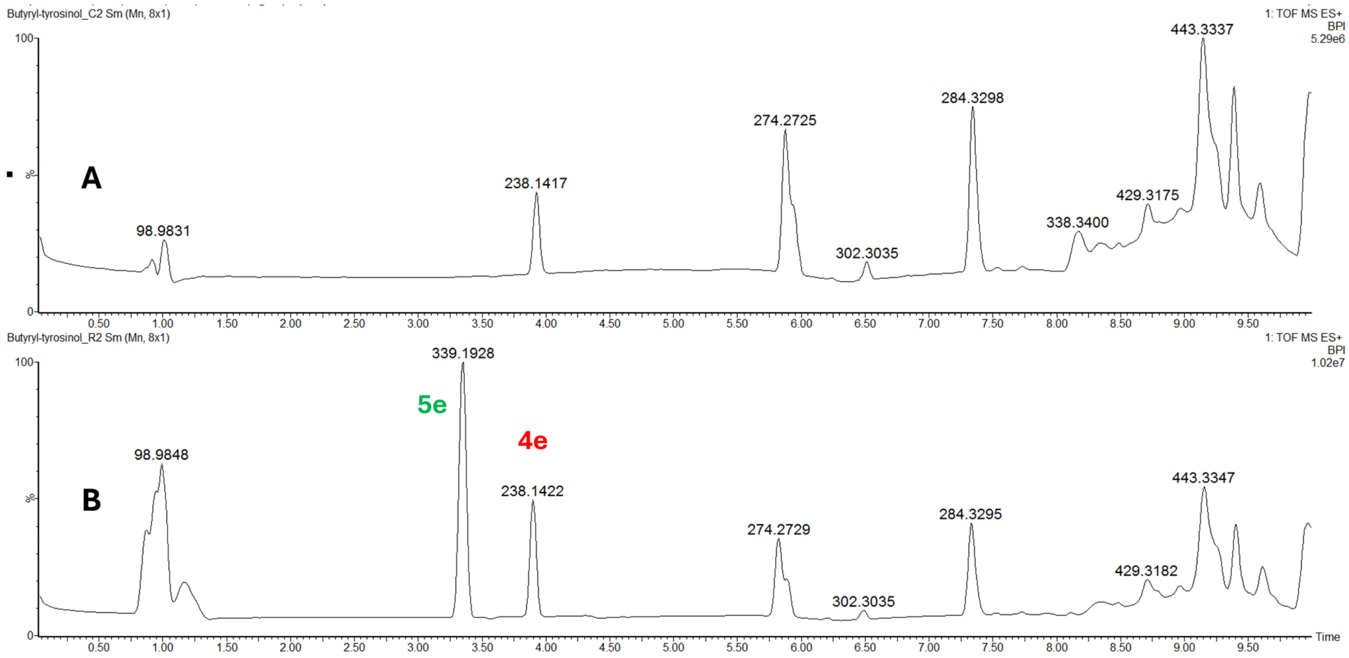


**Figure S31.** McalACT catalyzes aminocarboxypropylation of *N*-butyryl-L-tyrosinol (**4e**) in vitro. UPLC-MS base peak intensity chromatograms of **A)** Substrate control containing **4e** (1 mM), SAM (1 mM), and $MgCl_2$ (10 mM). **B)** Reaction mixture containing McalACT (10 µM), **4e** (1 mM), SAM (1 mM), and $MgCl_2$ (10 mM).

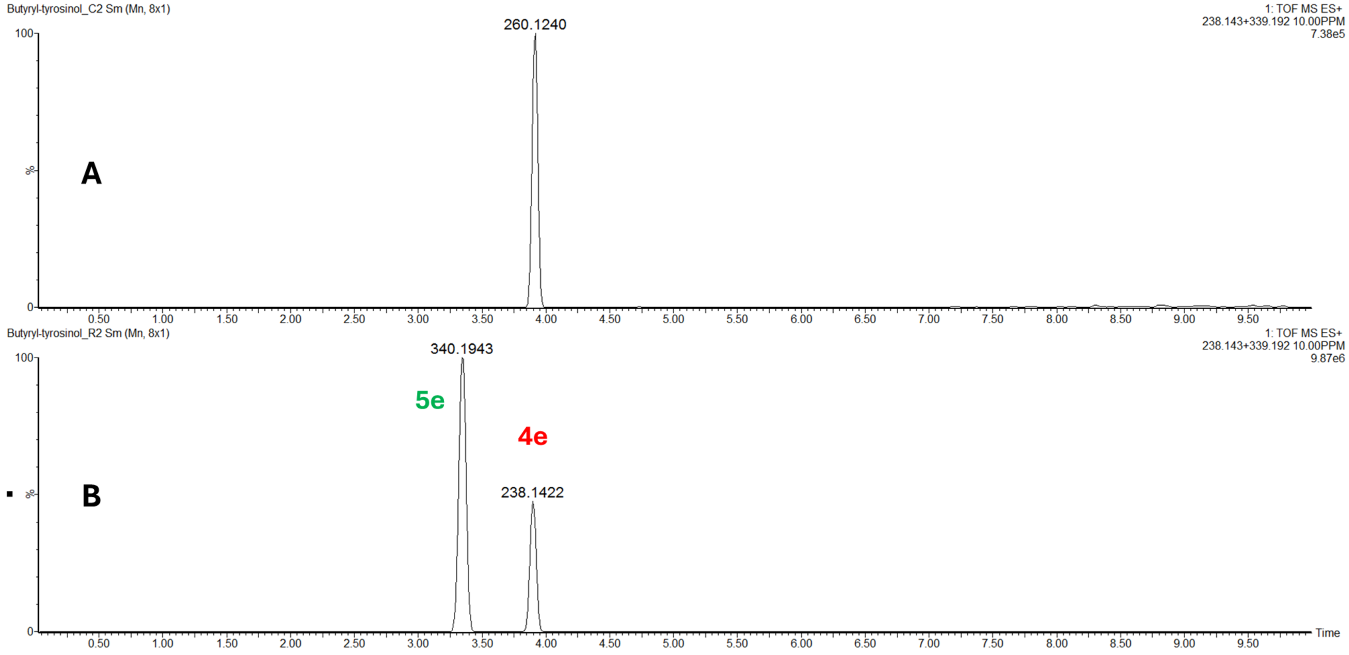


**Figure S32.** McalACT catalyzes aminocarboxypropylation of *N*-butyryl-L-tyrosinol (**4e**) in vitro. UPLC-MS extracted ion chromatograms of **A)** Substrate control containing **4e** (1 mM), SAM (1 mM), and $MgCl_2$ (10 mM). **B)** Reaction mixture containing McalACT (10 µM), **4e** (1 mM), SAM (1 mM), and $MgCl_2$ (10 mM).

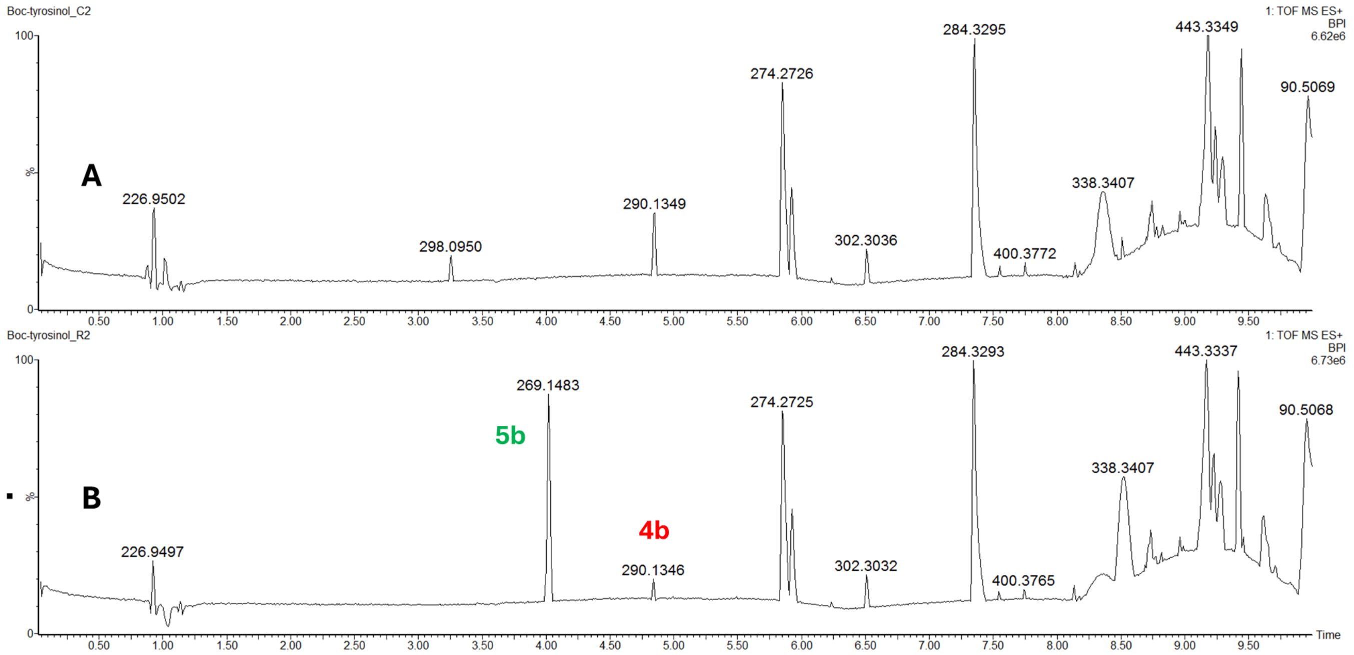


**Figure S33.** McalACT catalyzes aminocarboxypropylation of *N*-Boc-L-tyrosinol (**4b**) in vitro. UPLC-MS base peak intensity chromatograms of **A)** Substrate control containing **4b** (1 mM), SAM (1 mM), and $MgCl_2$ (10 mM). **B)** Reaction mixture containing McalACT (10 μM), **4b** (1 mM), SAM (1 mM), and $MgCl_2$ (10 mM).

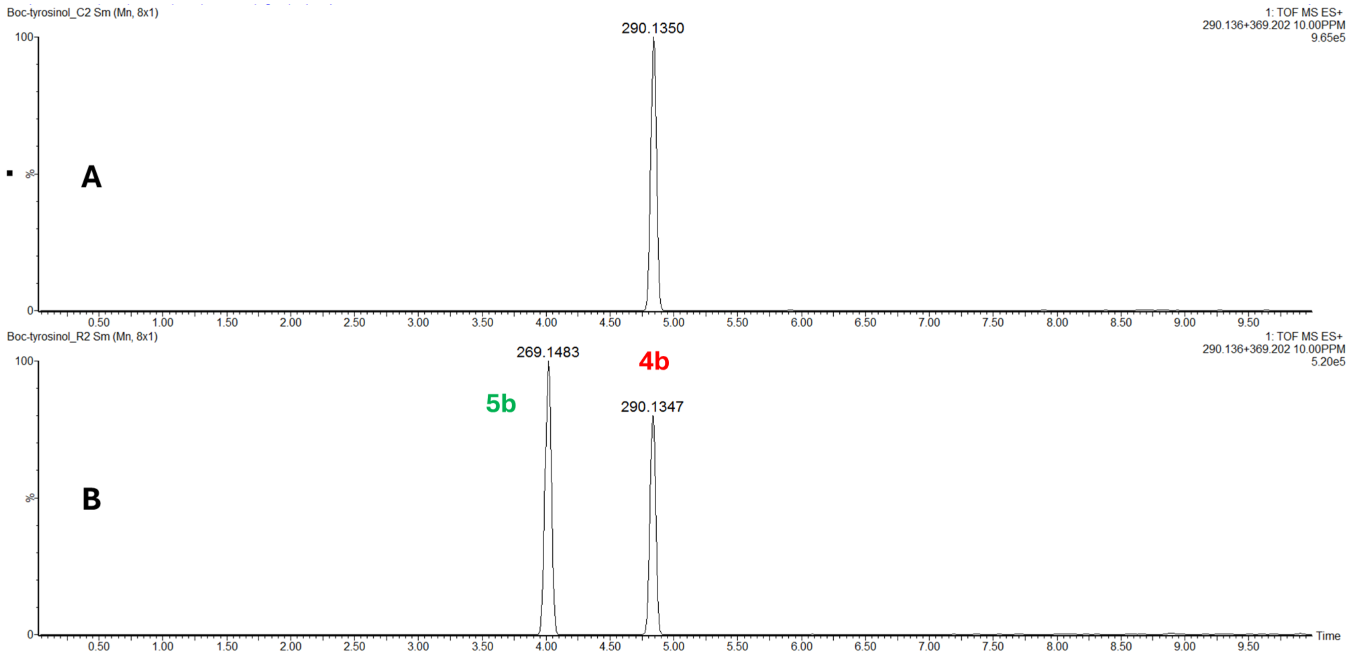


**Figure S34.** McalACT catalyzes aminocarboxypropylation of *N*-Boc-L-tyrosinol (**4b**) in vitro. UPLC-MS extracted ion chromatograms of **A)** Substrate control containing **4b** (1 mM), SAM (1 mM), and $MgCl_2$ (10 mM). **B)** Reaction mixture containing McalACT (10 μM), **4b** (1 mM), SAM (1 mM), and $MgCl_2$ (10 mM).

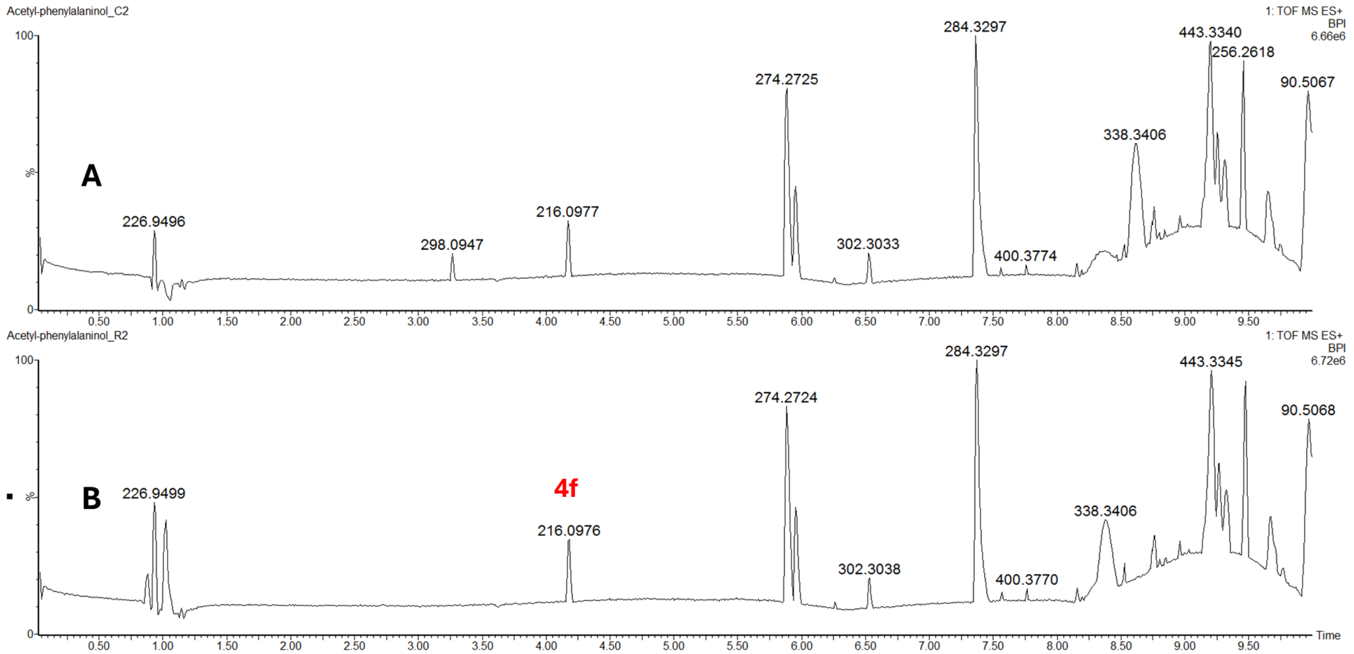


**Figure S35.** McalACT does not catalyze aminocarboxypropylation of *N*-acetyl-L-phenylalaninol (**4f**) in vitro. UPLC-MS base peak intensity chromatograms of **A)** Substrate control containing **4f** (1 mM), SAM (1 mM), and $MgCl_2$ (10 mM). **B)** Reaction mixture containing McalACT (10 µM), **4f** (1 mM), SAM (1 mM), and $MgCl_2$ (10 mM).

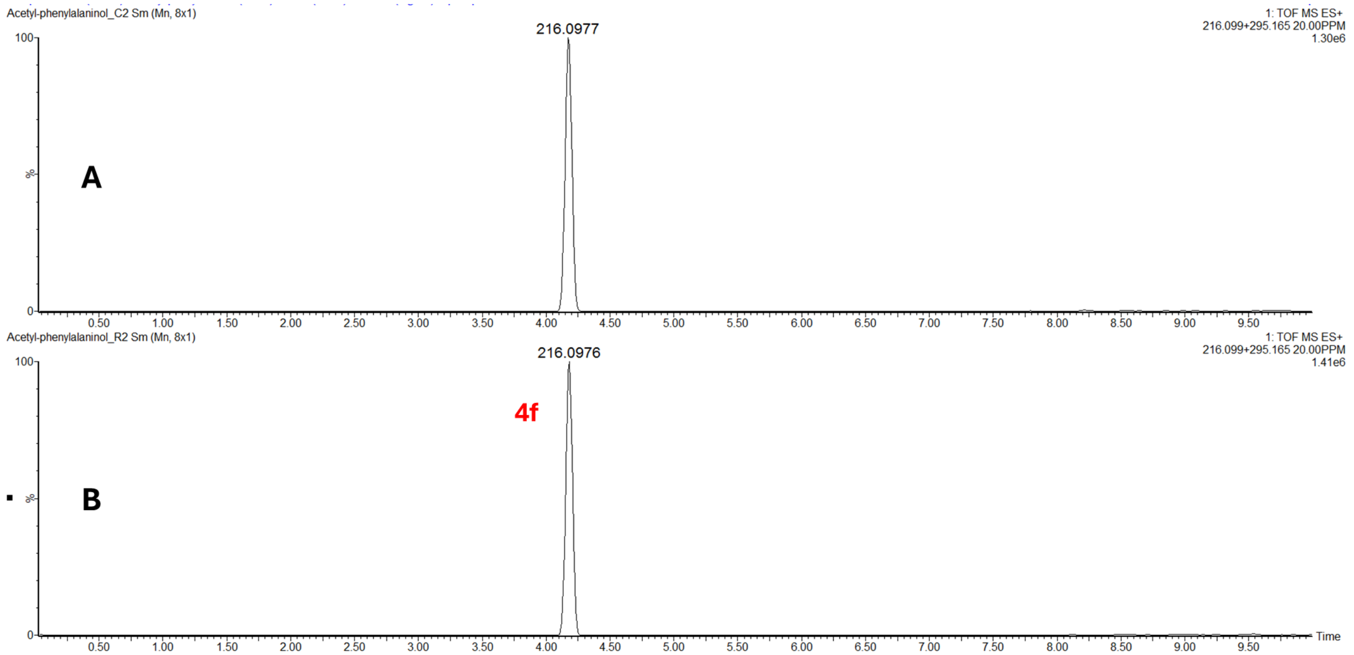


**Figure S36.** McalACT does not catalyze aminocarboxypropylation of *N*-acetyl-L-phenylalaninol (**4f**) in vitro. UPLC-MS extracted ion chromatograms of **A)** Substrate control containing **4f** (1 mM), SAM (1 mM), and $MgCl_2$ (10 mM). **B)** Reaction mixture containing McalACT (10 µM), **4f** (1 mM), SAM (1 mM), and $MgCl_2$ (10 mM).

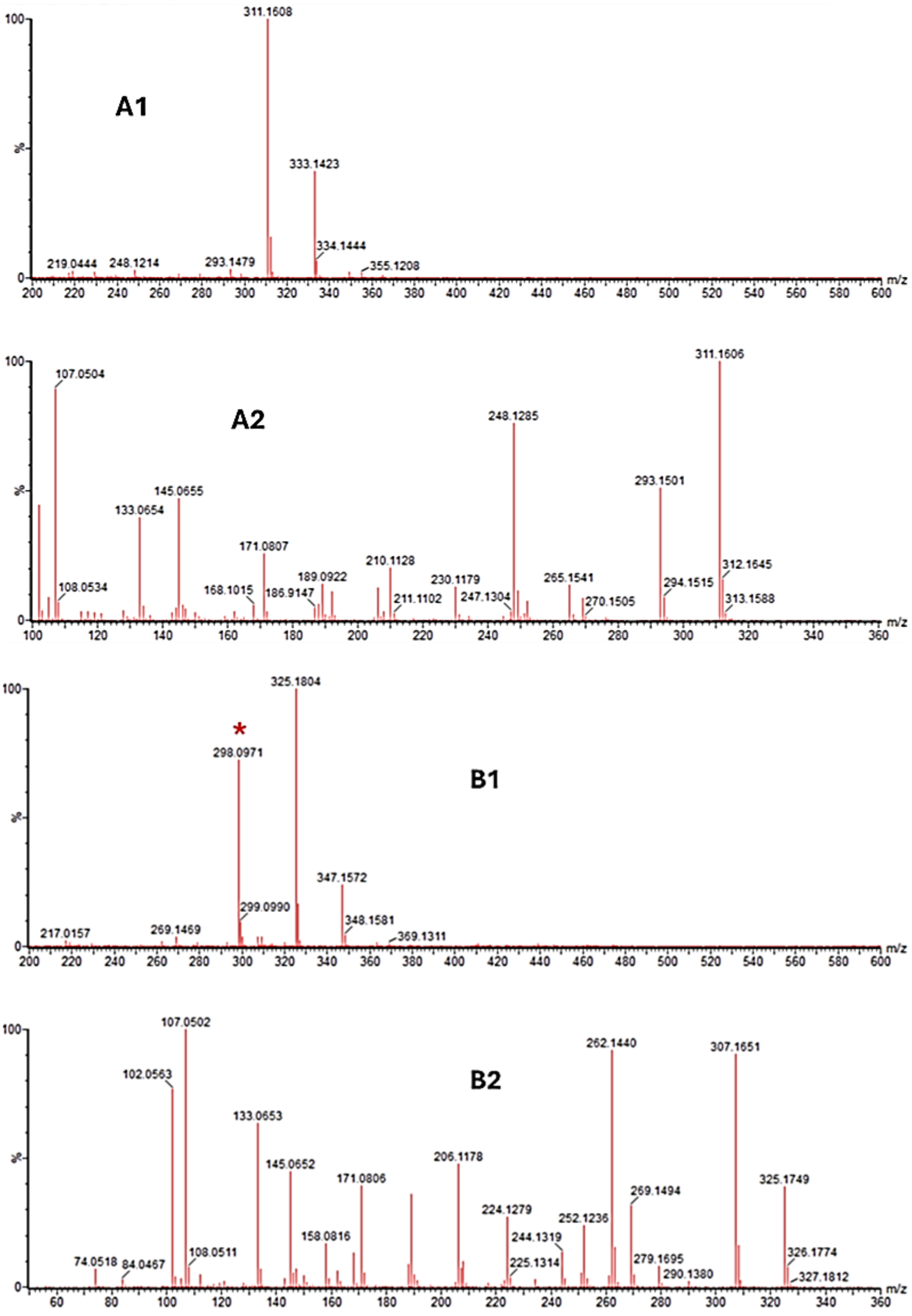
A1
311.1608
333.1423
334.1444
355.1208
219.0444
248.1214
293.1479
A2
107.0504
108.0534
133.0654
145.0655
168.1015
171.0807
186.9147
189.0922
210.1128
211.1102
230.1179
247.1304
248.1285
265.1541
270.1505
293.1501
294.1515
311.1606
312.1645
313.1588
B1
*
298.0971
299.0990
325.1804
347.1572
348.1581
369.1311
217.0157
269.1469
B2
74.0518
84.0467
102.0563
107.0502
108.0511
133.0653
145.0652
158.0816
171.0806
206.1178
224.1279
225.1314
244.1319
252.1236
262.1440
269.1494
279.1695
290.1380
307.1651
325.1749
326.1774
327.1812
%
m/z

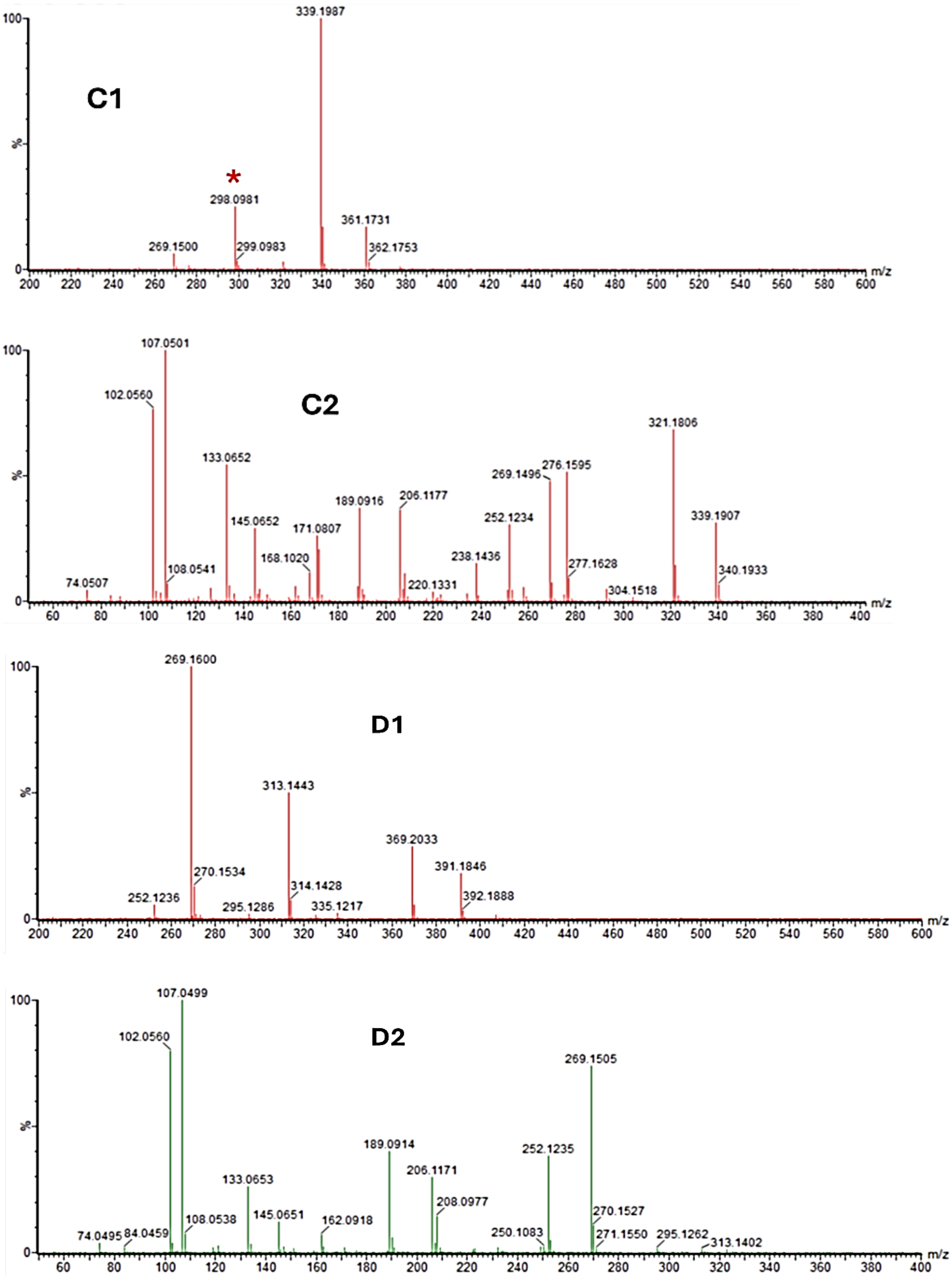


**Figure S37**. McalACT accepts various *N*-acyl tyrosinols as substrates. **A1)** MS spectrum of **5c** in reaction. **A2)** MS/MS spectrum of precursor ion (*m/z* 311.1608) from A1. **B1)** MS spectrum of **5d** in reaction. **B2)** MS/MS spectrum of precursor ion (*m/z* 325.1804) from B1. **C1)** MS spectrum of **5e** in reaction. **C2)** MS/MS spectrum of precursor ion (*m/z* 339.1987) from C1. **D1)** MS spectrum of **5b** in reaction. **D2)** MS/MS spectrum of precursor ion (*m/z* 369.2033) from D1. *Methylthioadenosine precursor ion (*m/z* 298.09) contamination.

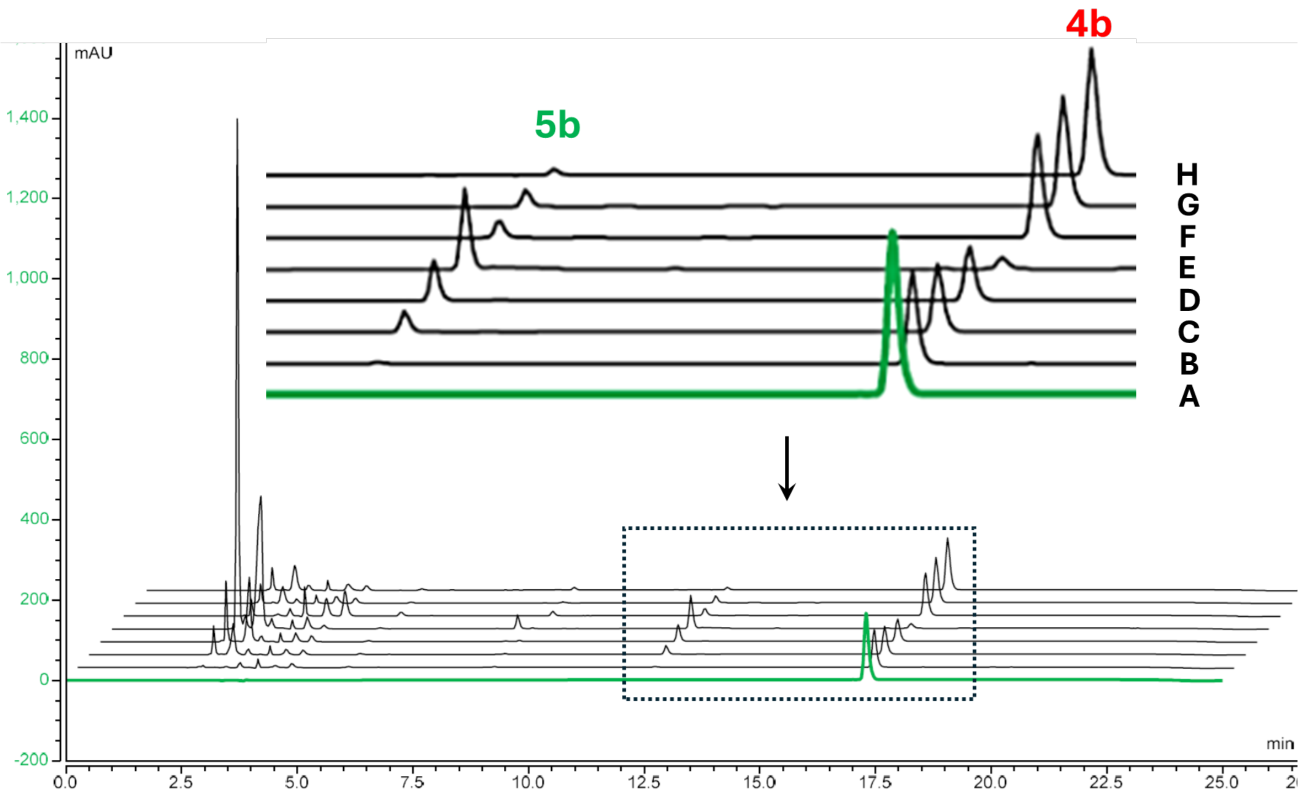


**Figure S38**. Screening of conditions for McalACT-catalyzed aminocarboxypropylation of **4b**. HPLC-UV chromatograms at 280 nm of **A)** Synthetic standard of **4b**. **B)** **0.1 eq. SAM:** Reaction mixture containing McalACT (20 µM), **4b** (0.5 mM), SAM (0.05 mM), and $MgCl_2$ (10 mM) incubated at 25 °C. **C)** **1 eq. SAM:** Reaction mixture containing McalACT (20 µM), **4b** (0.5 mM), SAM (0.5 mM), and $MgCl_2$ (10 mM) incubated at 25 °C. **D)** **2 eq. SAM:** Reaction mixture containing McalACT (20 µM), **4b** (0.5 mM), SAM (1 mM), and $MgCl_2$ (10 mM) incubated at 25 °C. **E)** **10 eq. SAM:** Reaction mixture containing McalACT (20 µM), **4b** (0.5 mM), SAM (5 mM), and $MgCl_2$ (10 mM) incubated at 25 °C. **F)** **2x enzyme + 1 eq. SAM:** Reaction mixture containing McalACT (40 µM), **4b** (0.5 mM), SAM (0.5 mM), and $MgCl_2$ (10 mM) incubated at 25 °C. **G)** **30 °C + 1 eq. SAM** Reaction mixture containing McalACT (20 µM), **4b** (0.5 mM), SAM (0.05 mM), and $MgCl_2$ (10 mM) incubated at 30 °C. **H)** **37 °C + 1 eq. SAM** Reaction mixture containing McalACT (20 µM), **4b** (0.5 mM), SAM (0.05 mM), and $MgCl_2$ (10 mM) incubated at 37 °C.

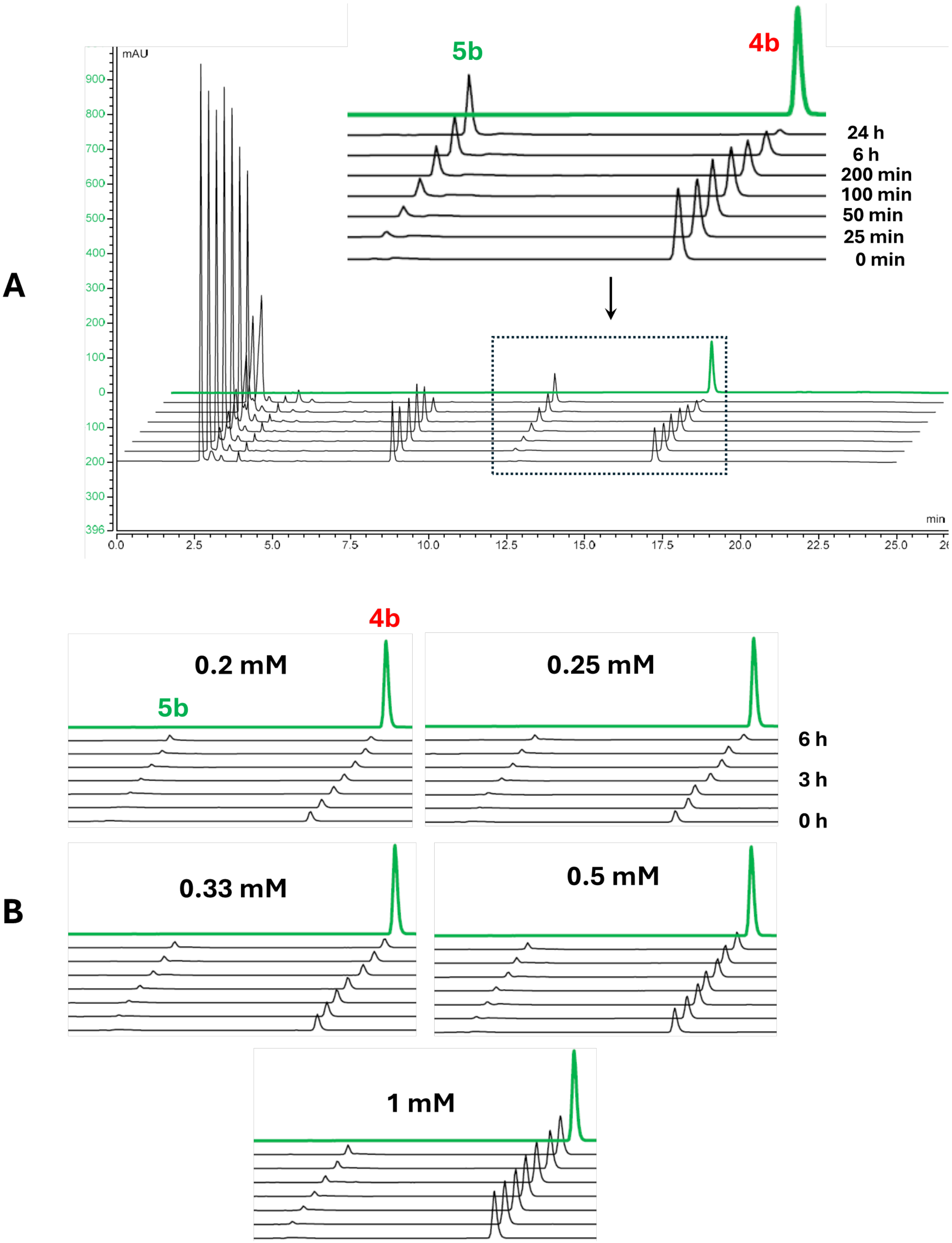


**Figure S39**. HPLC-UV chromatograms at 280 nm for initial rate experiments of McalACT-catalyzed **4b** aminocarboxypropylation. **A)** Reaction mixtures contain McalACT (20 µM), **4b** (0.5 mM), SAM (5 mM), and $MgCl_2$ (10 mM) incubated at 25 °C for 24 h. **B)** Reaction mixtures contain McalACT (10 µM), **4b** (0.2 -1 mM), SAM (5 mM), and $MgCl_2$ (10 mM) incubated at 25 °C for 6 h (for initial reaction rate estimation).

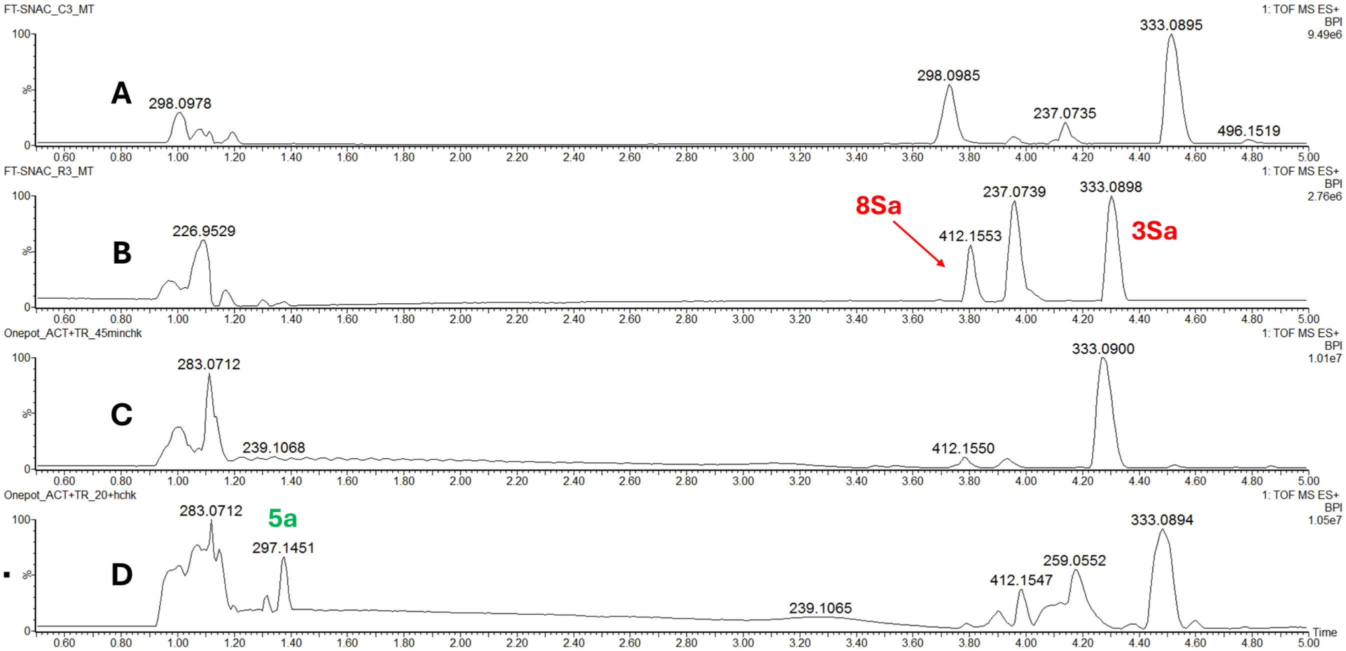


**Figure S40**. One-pot reactions for **3Sa** with McalACT and CvTR. UPLC-MS base peak intensity chromatograms of **A)** Substrate control containing **3Sa** (100 µM), SAM (1 mM), and $MgCl_2$ (10 mM). **B)** Reaction mixture containing McalACT (10 µM**), 3Sa** (100 µM), SAM (1 mM), and $MgCl_2$ (10 mM). **C)** Reaction mixture containing CvTR (10 µM), McalACT (10 µM), **3Sa** (100 µM), NADPH (1 mM), SAM (1 mM), and $MgCl_2$ (10 mM) quenched at 45 min. **D)** Reaction mixture containing CvTR (10 µM), McalACT (10 µM), **3Sa** (100 µM), NADPH (1 mM), SAM (1 mM), and $MgCl_2$ (10 mM) quenched at 20 h.

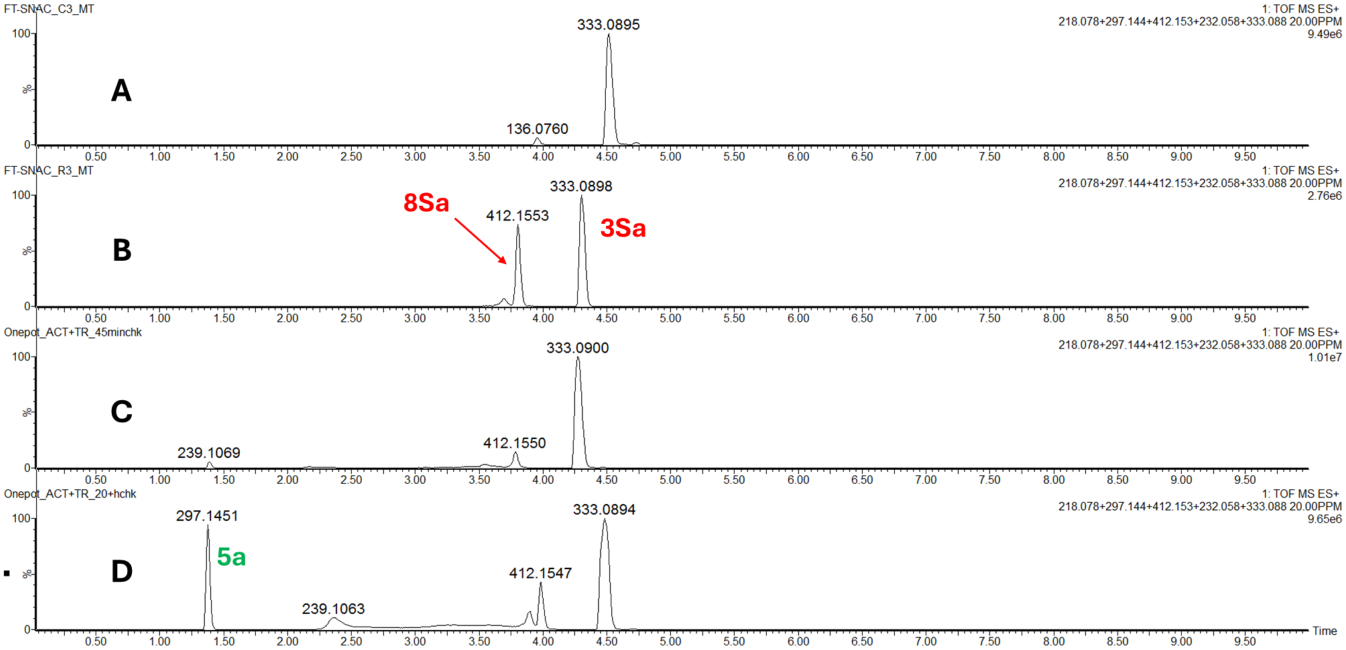


**Figure S41**. One-pot reactions for **3Sa** with McalACT and CvTR. UPLC-MS extracted ion chromatograms of **A)** Substrate control containing **3Sa** (100 µM), SAM (1 mM), and $MgCl_2$ (10 mM). **B)** Reaction mixture containing McalACT (10 µM**), 3Sa** (100 µM), SAM (1 mM), and $MgCl_2$ (10 mM). **C)** Reaction mixture containing CvTR (10 µM), McalACT (10 µM), **3Sa** (100 µM), NADPH (1 mM), SAM (1 mM), and $MgCl_2$ (10 mM) quenched at 45 min. **D)** Reaction mixture containing CvTR (10 µM), McalACT (10 µM), **3Sa** (100 µM), NADPH (1 mM), SAM (1 mM), and $MgCl_2$ (10 mM) quenched at 20 h.

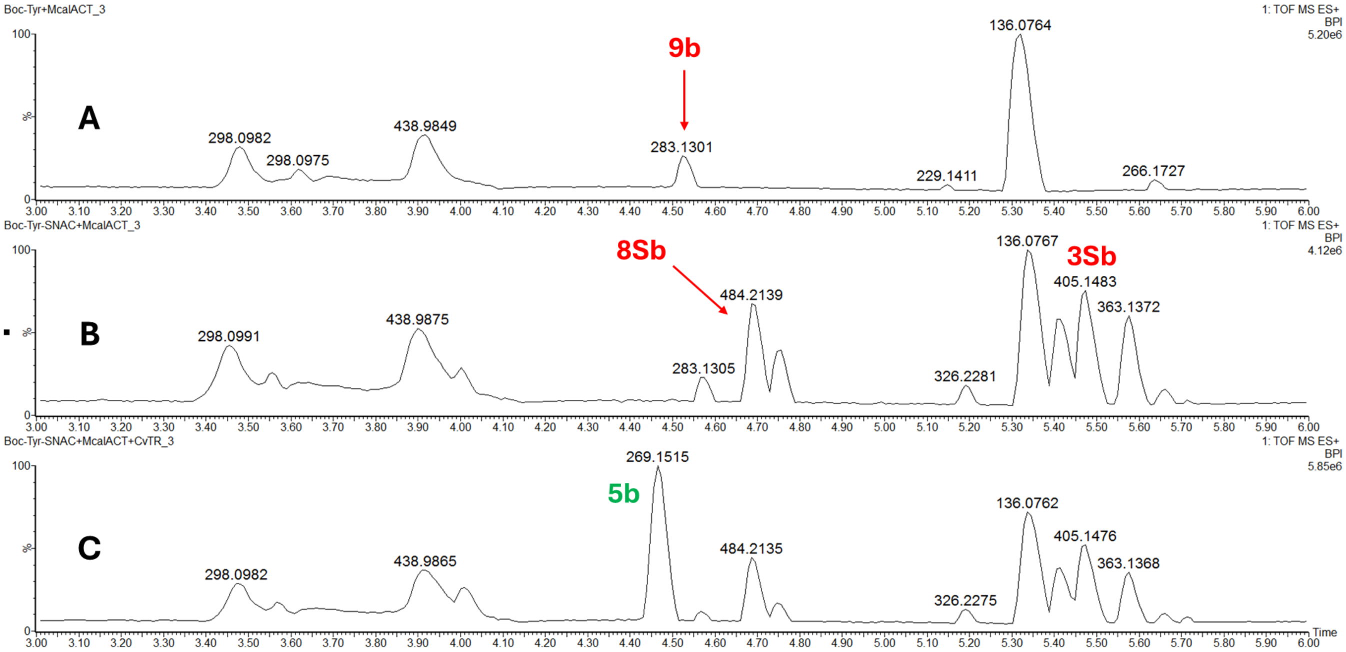


**Figure S42**. One-pot reactions for **3Sb** with McalACT and CvTR. UPLC-MS base peak intensity chromatograms of **A)** Reaction mixture containing McalACT (10 μM**), 2b** (100 μM), SAM (1 mM), and $MgCl_2$ (10 mM). **B)** Reaction mixture containing McalACT (10 μM**), 3Sb** (100 μM), SAM (1 mM), and $MgCl_2$ (10 mM). **C)** Reaction mixture containing CvTR (10 μM), McalACT (10 μM), **3Sb** (100 μM), NADPH (1 mM), SAM (1 mM), and $MgCl_2$ (10 mM) quenched at 20 h.

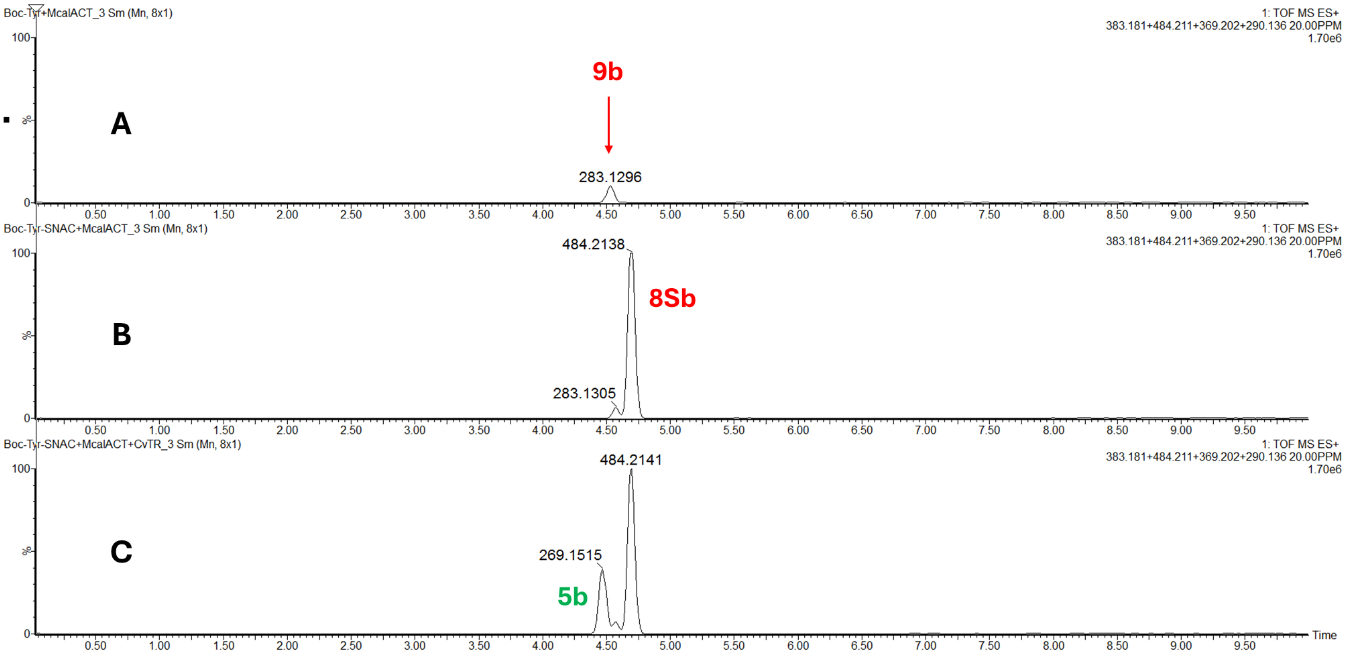


**Figure S43**. One-pot reactions for **3Sb** with McalACT and CvTR. UPLC-MS extracted ion chromatograms of **A)** Reaction mixture containing McalACT (10 μM**), 2b** (100 μM), SAM (1 mM), and $MgCl_2$ (10 mM). **B)** Reaction mixture containing McalACT (10 μM**), 3Sb** (100 μM), SAM (1 mM), and $MgCl_2$ (10 mM). **C)** Reaction mixture containing CvTR (10 μM), McalACT (10 μM), **3Sb** (100 μM), NADPH (1 mM), SAM (1 mM), and $MgCl_2$ (10 mM) quenched at 20 h.

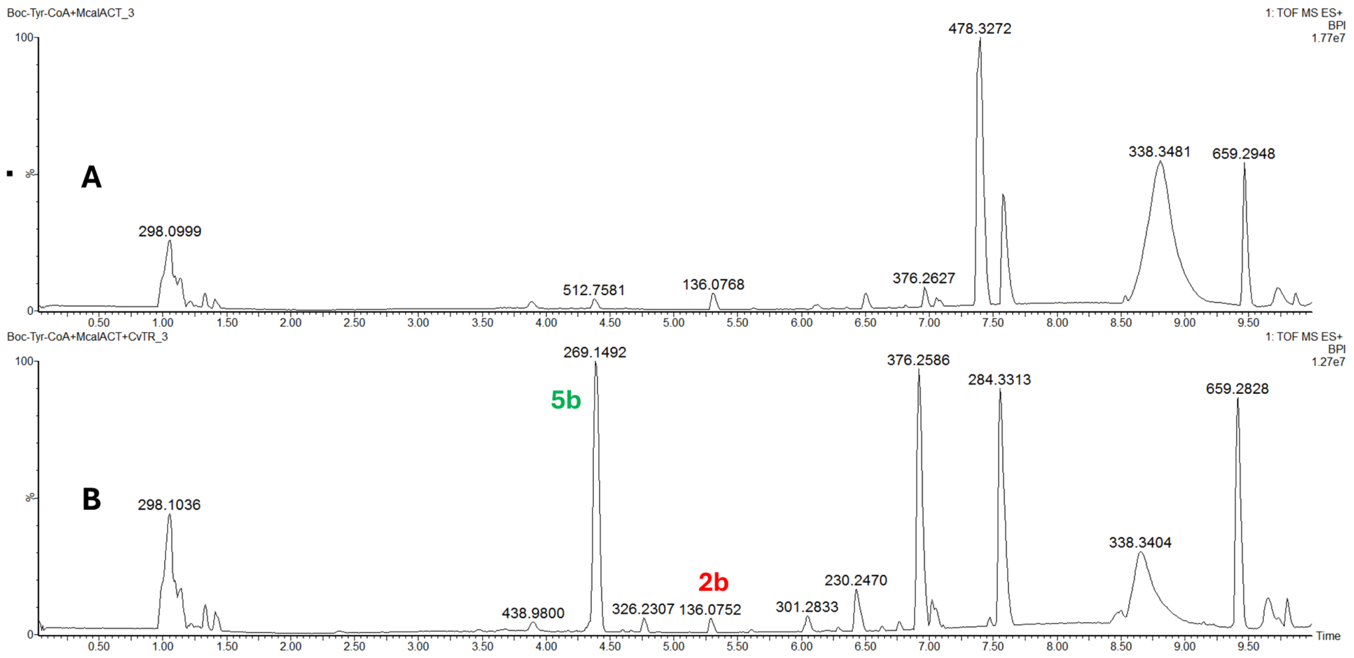


**Figure S44**. One-pot reactions for **3Cb** with McalACT and CvTR. UPLC-MS base peak intensity chromatograms of **A)** Reaction mixture containing McalACT (10 µM**), 3Cb** (100 µM), SAM (1 mM), and $MgCl_2$ (10 mM). **B)** Reaction mixture containing CvTR (10 µM), McalACT (10 µM), **3Cb** (100 µM), NADPH (1 mM), SAM (1 mM), and $MgCl_2$ (10 mM) quenched at 20 h.

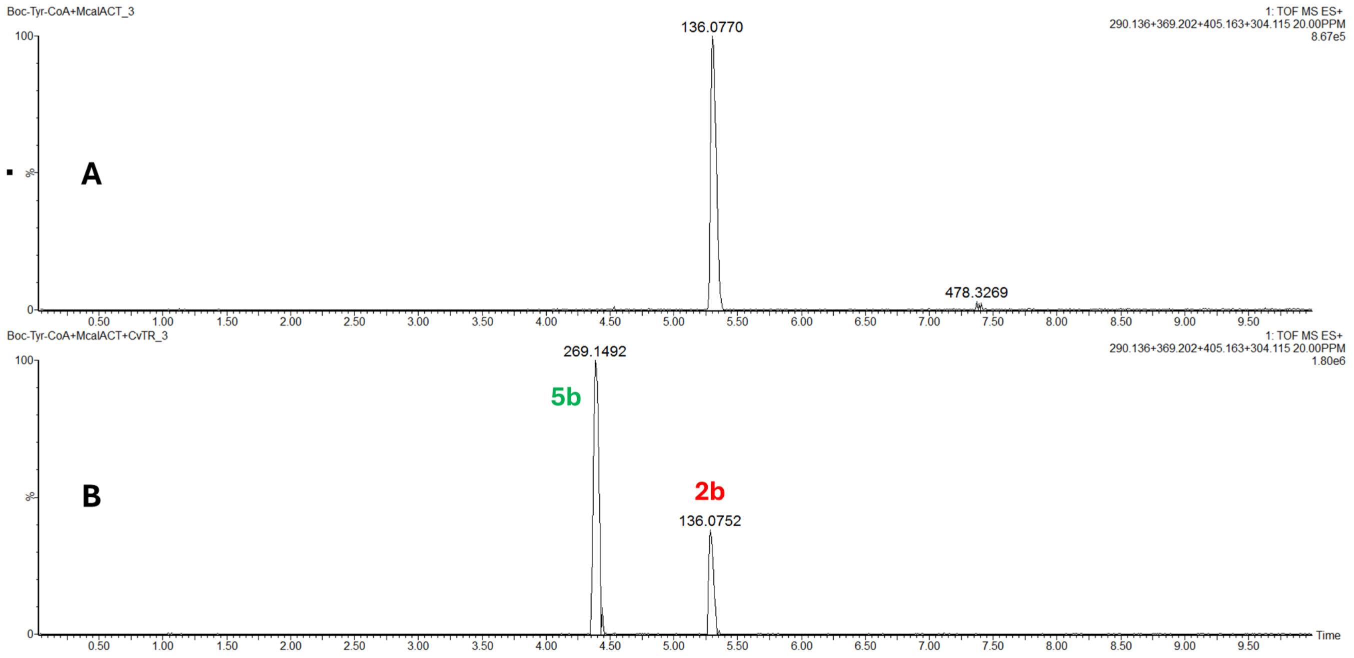


**Figure S45**. One-pot reactions for **3Cb** with McalACT and CvTR. UPLC-MS extracted ion chromatograms of **A)** Reaction mixture containing McalACT (10 µM**), 3Cb** (100 µM), SAM (1 mM), and $MgCl_2$ (10 mM). **B)** Reaction mixture containing CvTR (10 µM), McalACT (10 µM), **3Cb** (100 µM), NADPH (1 mM), SAM (1 mM), and $MgCl_2$ (10 mM) quenched at 20 h.

5. NMR and MS data for synthetic substrates, intermediates, and standards

5.1 NMR data for L-tyrosyl *O*-homoserine ethers

5.1.1 NMR data for **5b***

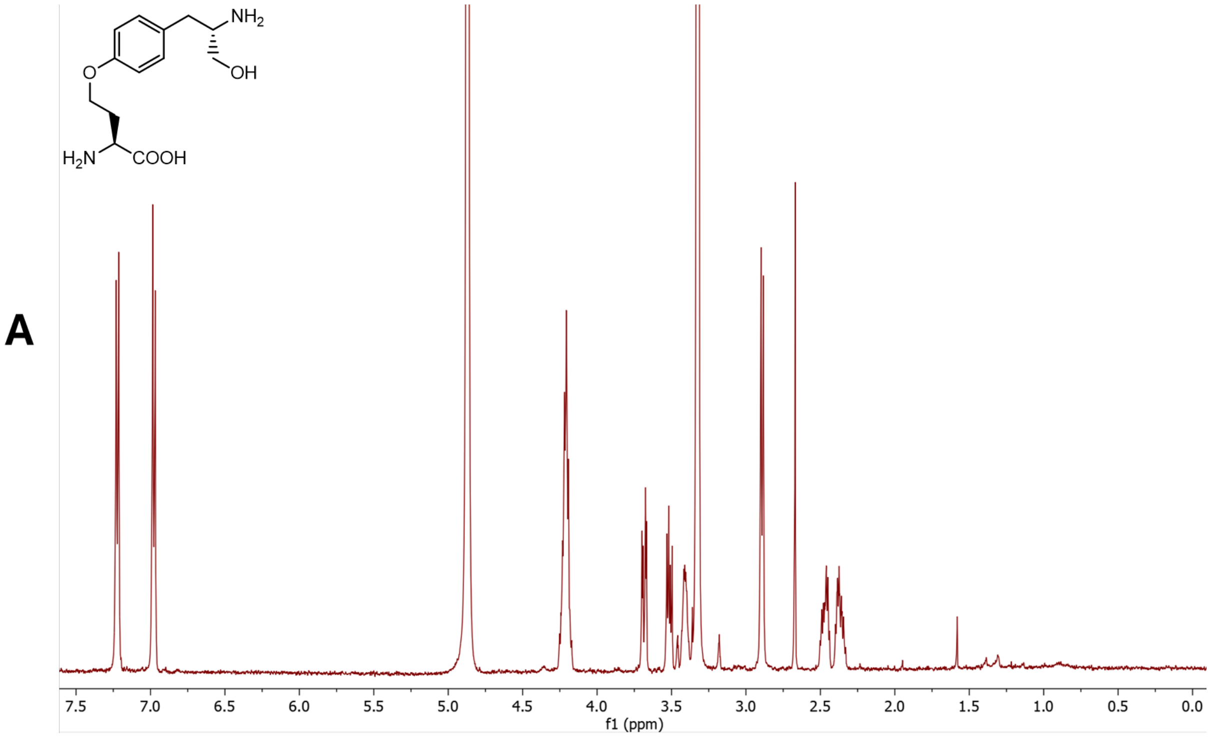

B

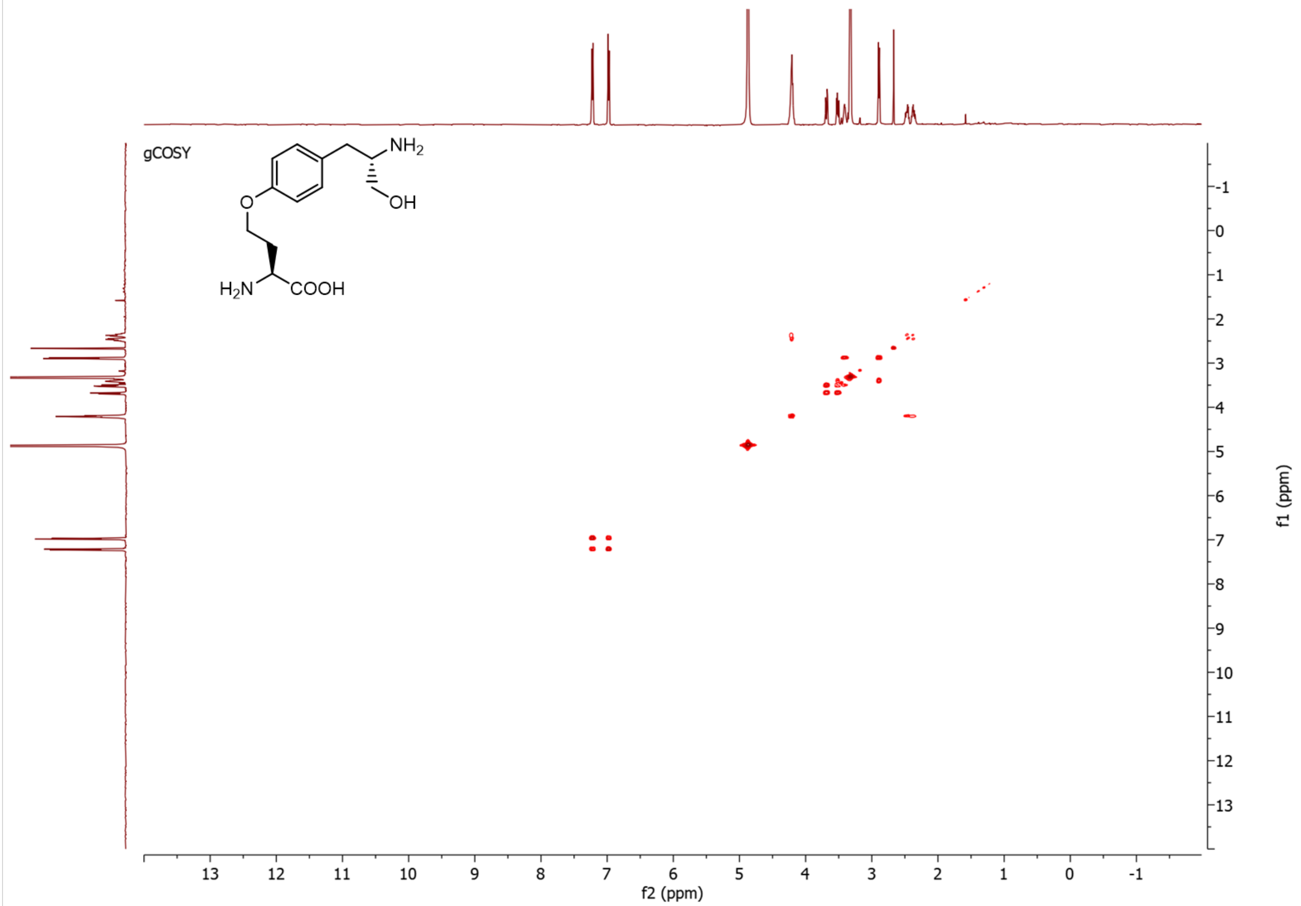
gCOSY
NH2
OH
O
H2N
COOH
f2 (ppm)
f1 (ppm)

C

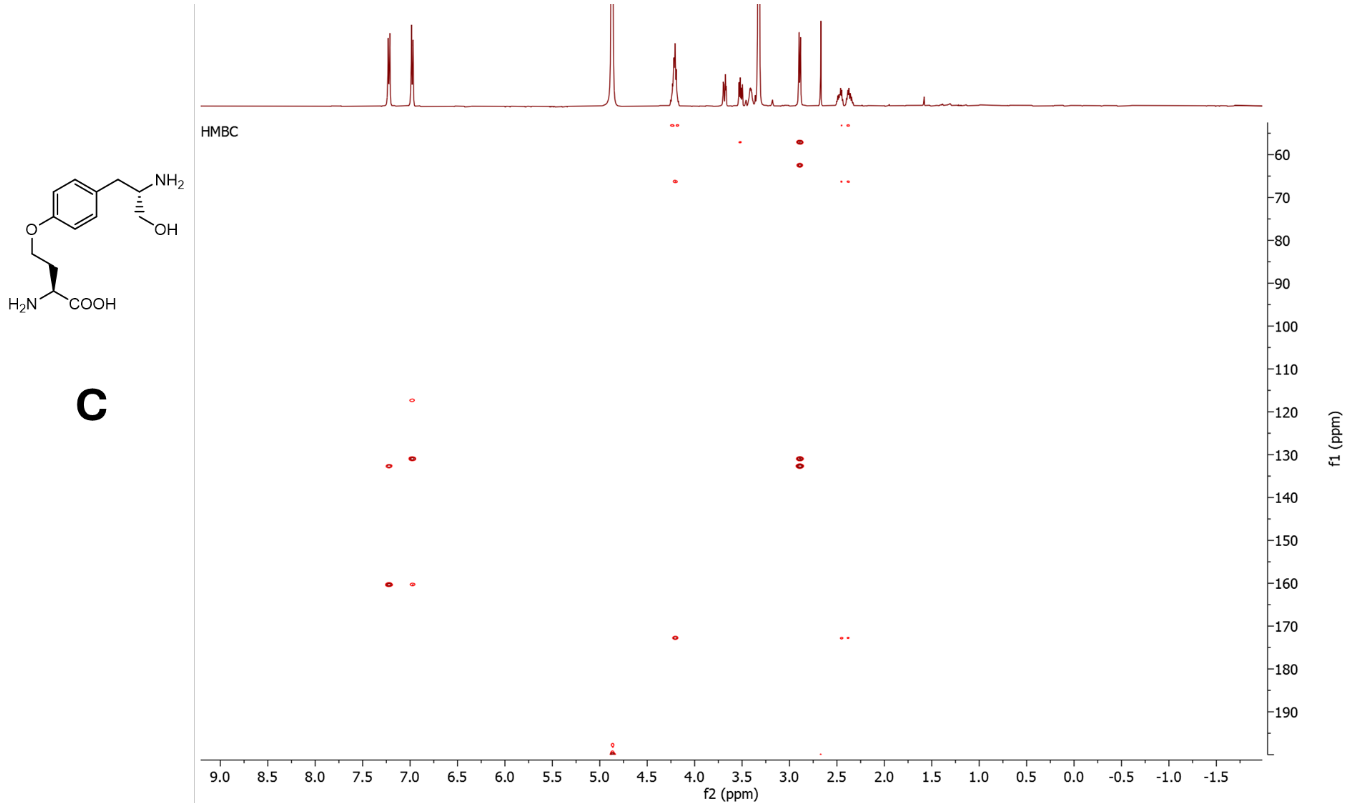
HMBC
NH2
OH
O
H2N
COOH
f2 (ppm)
f1 (ppm)

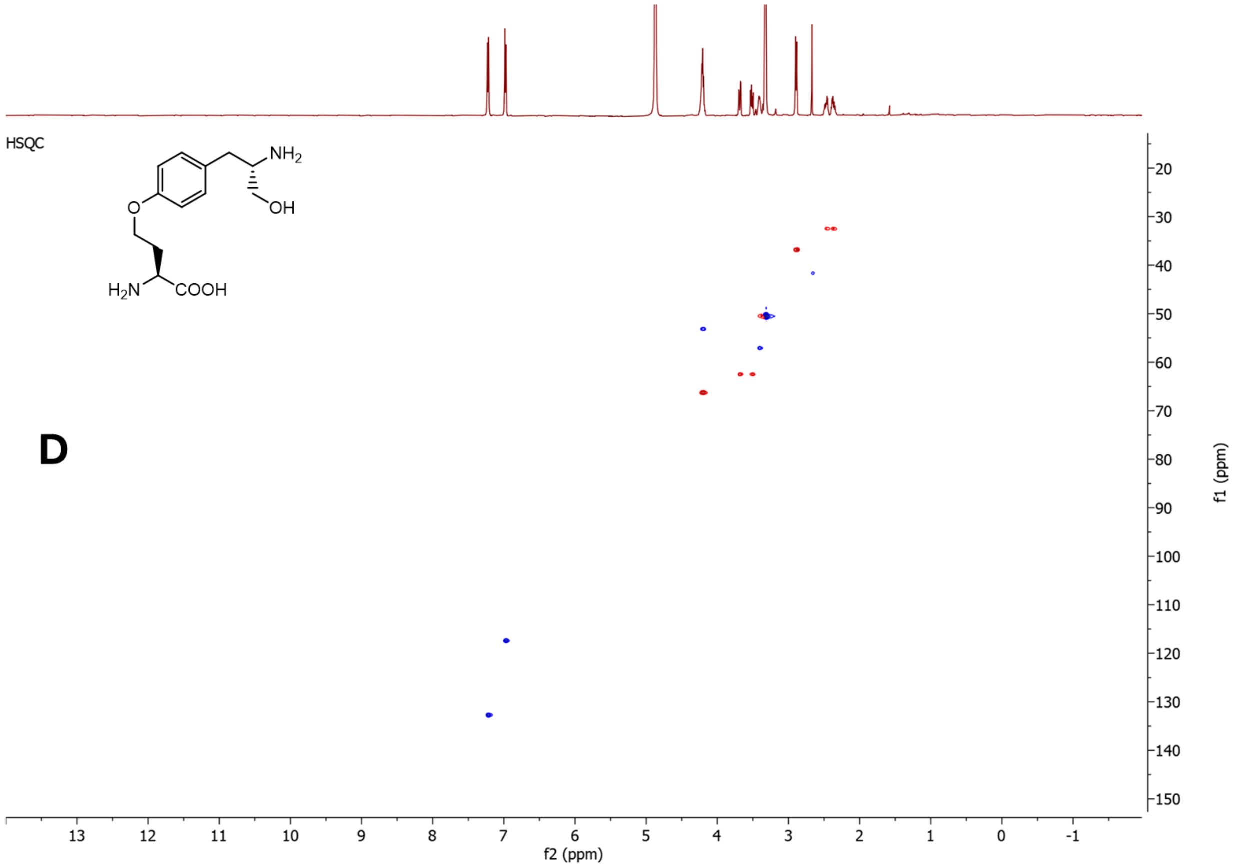


**Figure S46**. NMR data for **5b*** in $CD_3OD$, 500 MHz. **A)** $^1H$-NMR spectrum. **B)** COSY spectrum. **C)** HMBC spectrum. **D)** HSQC spectrum.

5.1.2 NMR data for **5b + 5b***

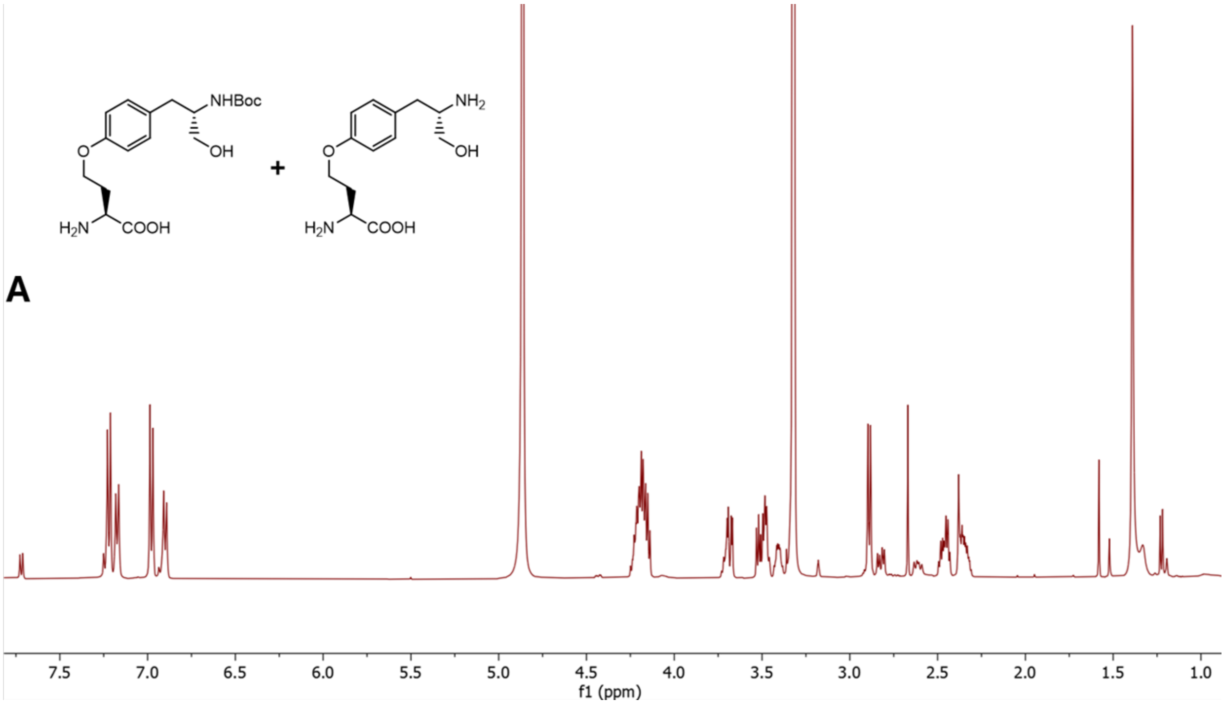


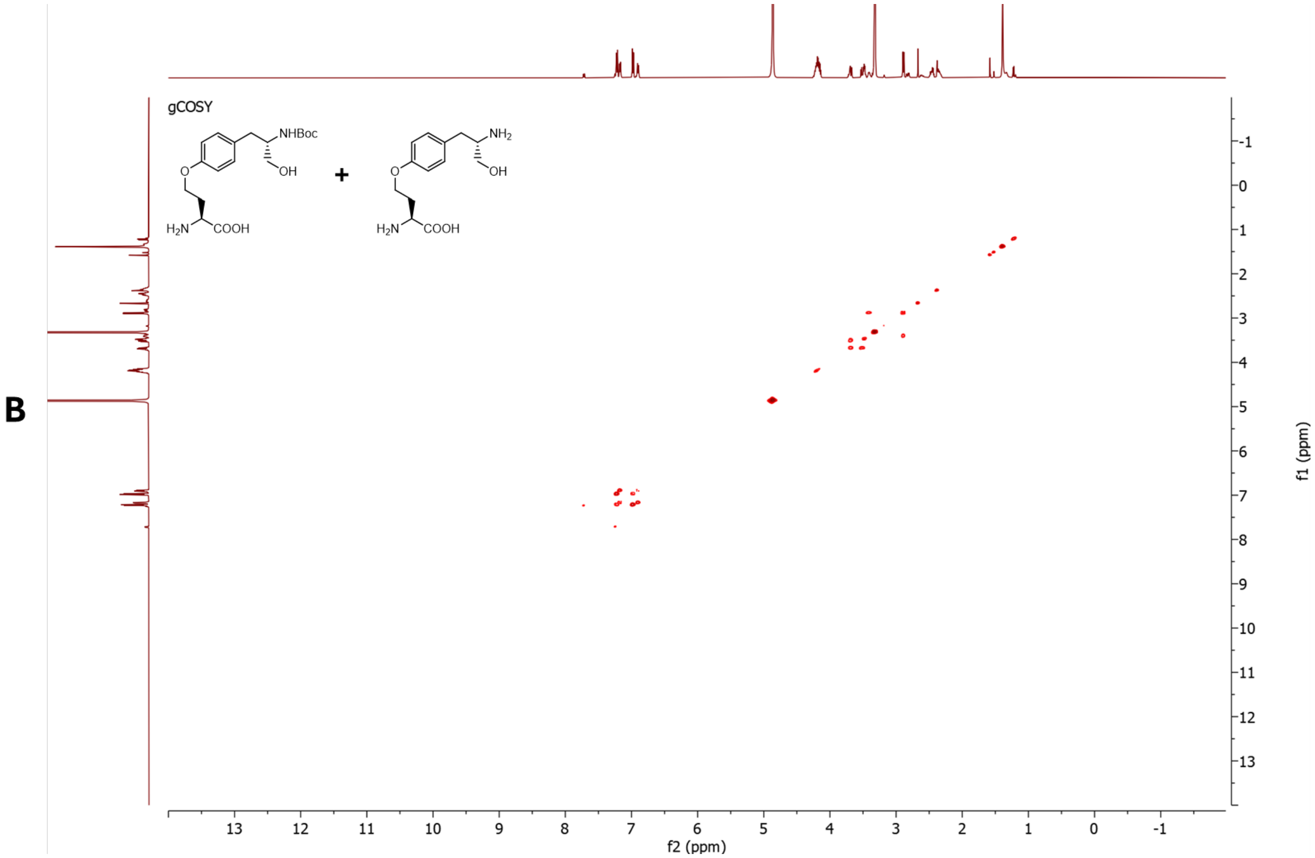

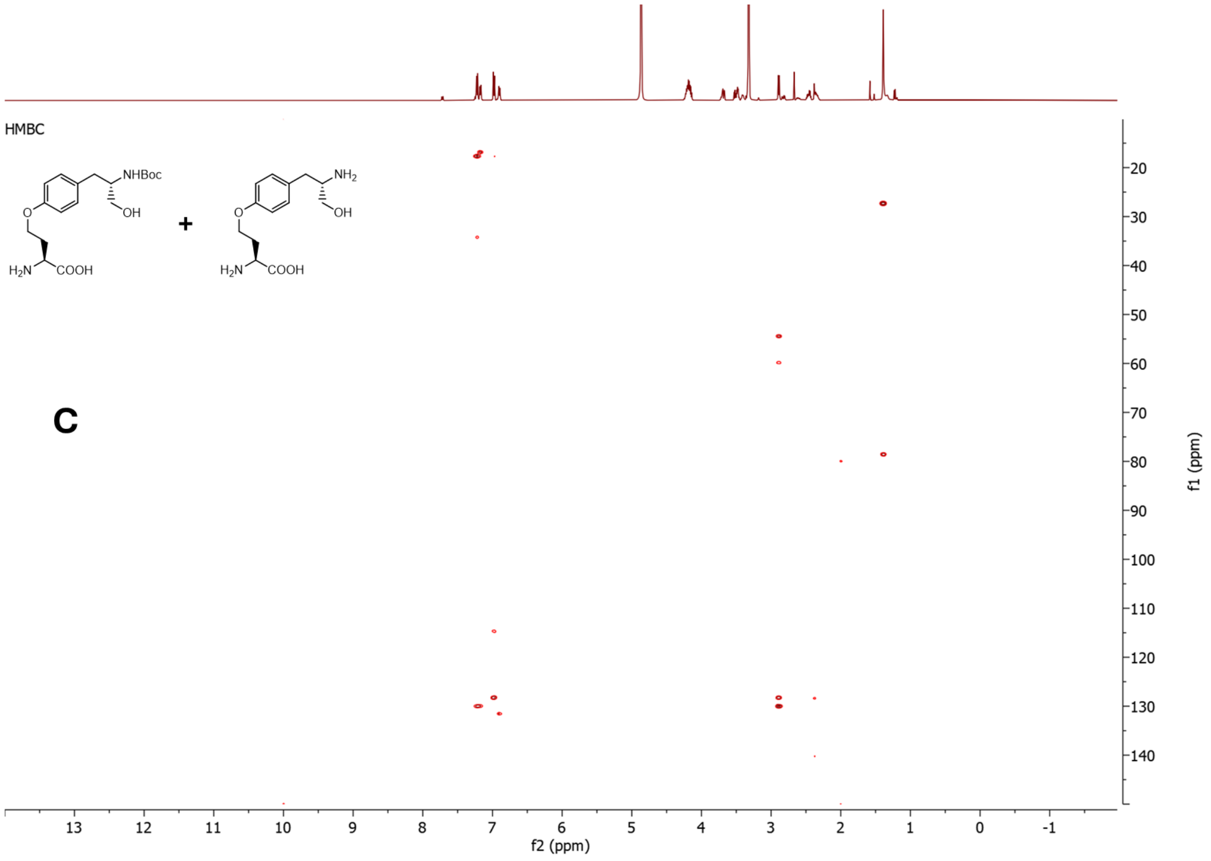
HMBC
NHBoc
OH
O
$H_2N$
COOH
+
$NH_2$
OH
O
$H_2N$
COOH
C
f1 (ppm)
f2 (ppm)

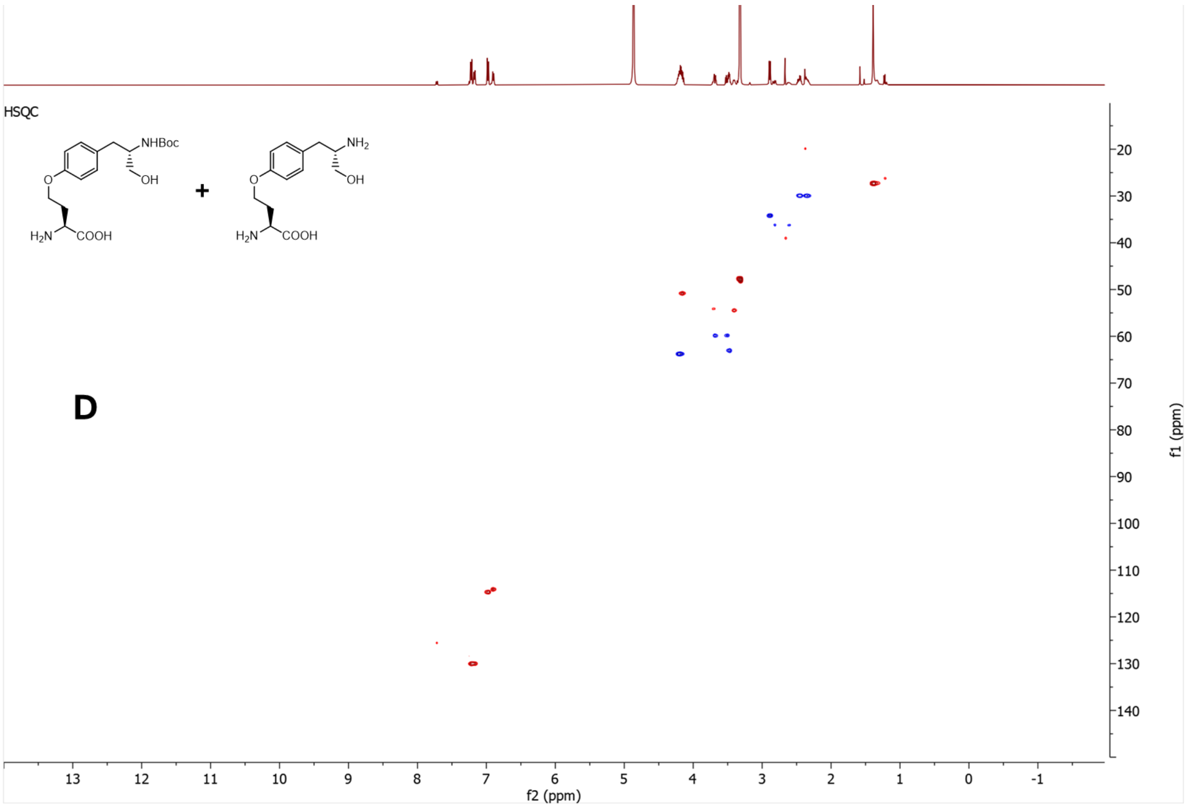
HSQC
NHBoc
OH
O
$H_2N$
COOH
+
$NH_2$
OH
O
$H_2N$
COOH
D
f1 (ppm)
f2 (ppm)

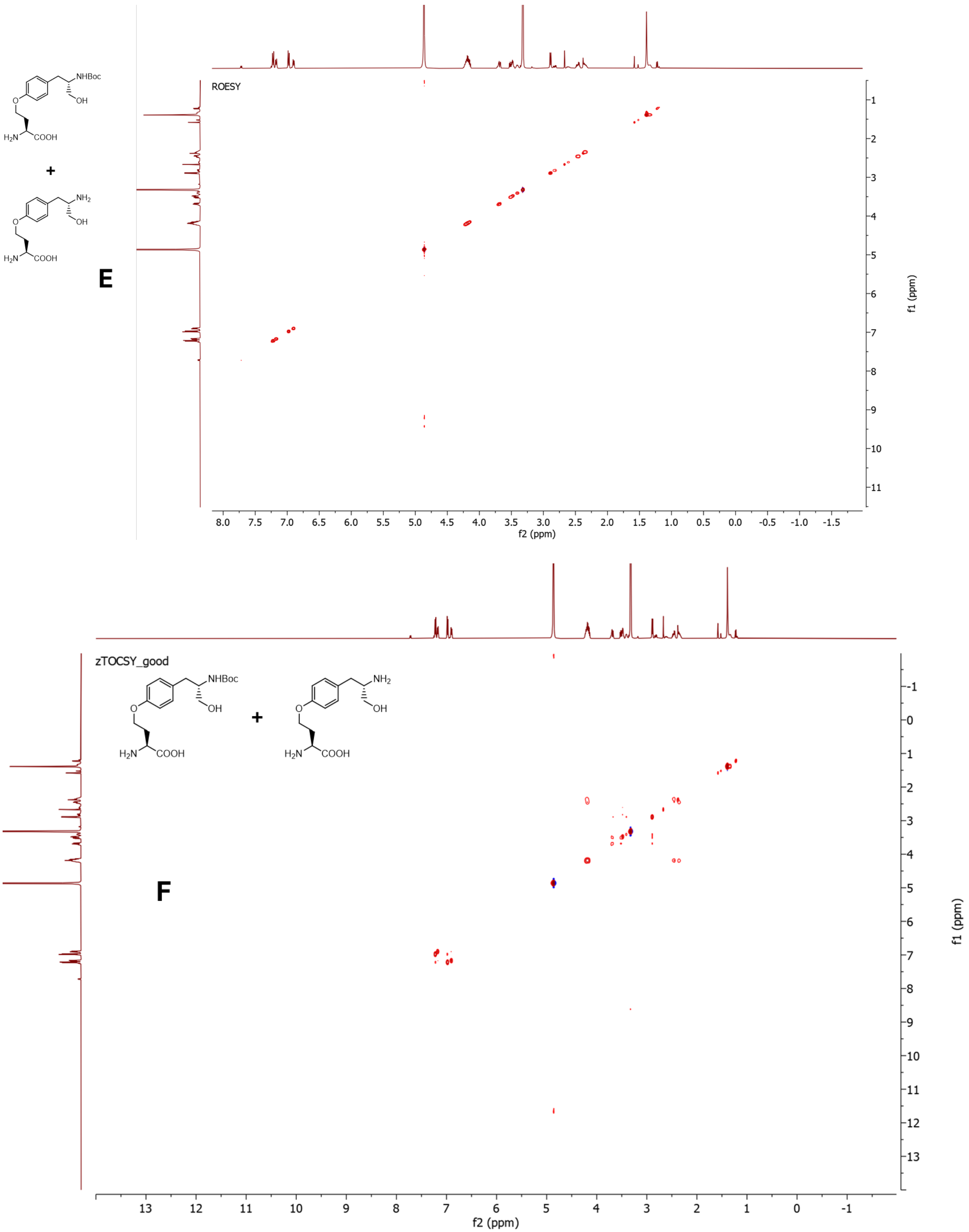


**Figure S47**. NMR data for mixture of **5b** and **5b*** in $CD_3OD$, 500 MHz. **A)** $^1$H-NMR spectrum. **B)** COSY spectrum. **C)** HMBC spectrum. **D)** HSQC spectrum. **E)** ROESY spectrum. **F)** TOCSY spectrum.

5.2 NMR and HR-MS/MS data for *N*-acyl-L-tyrosinols

5.2.1 NMR and HR-MS/MS data for **4a**

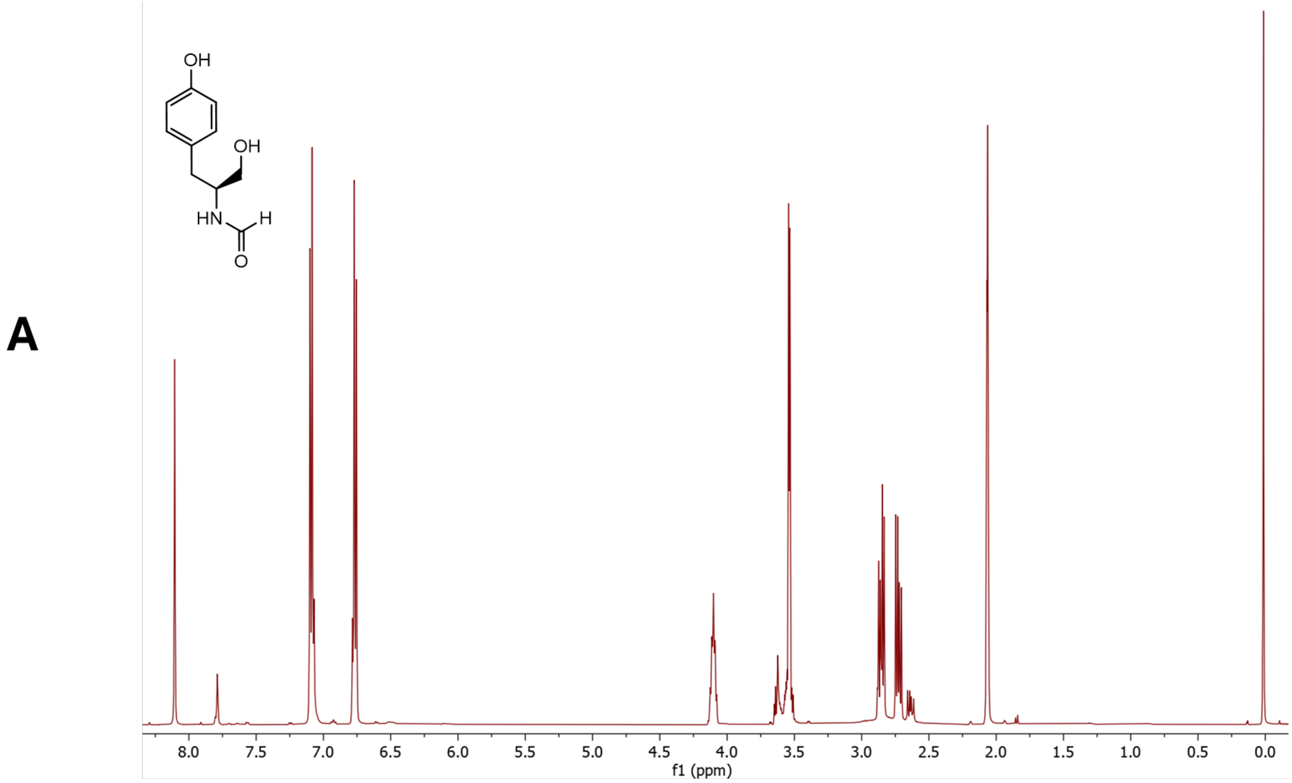


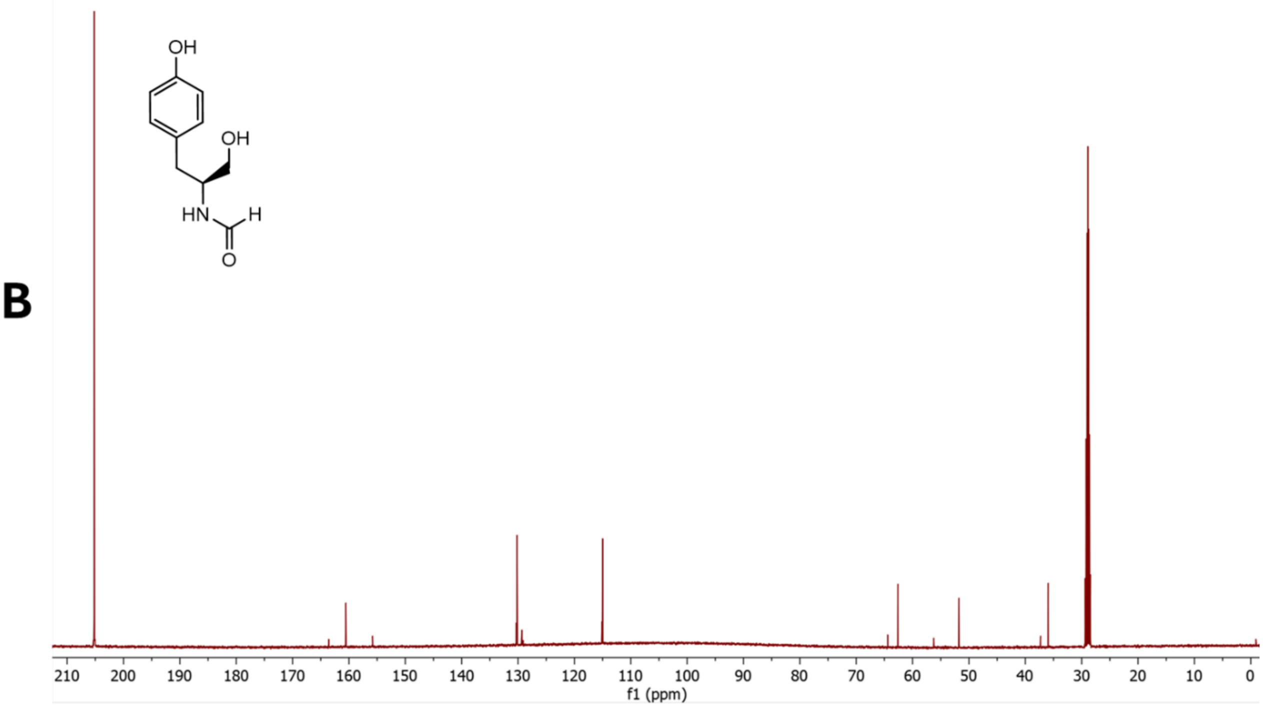

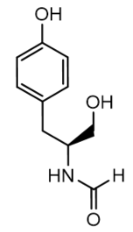

C

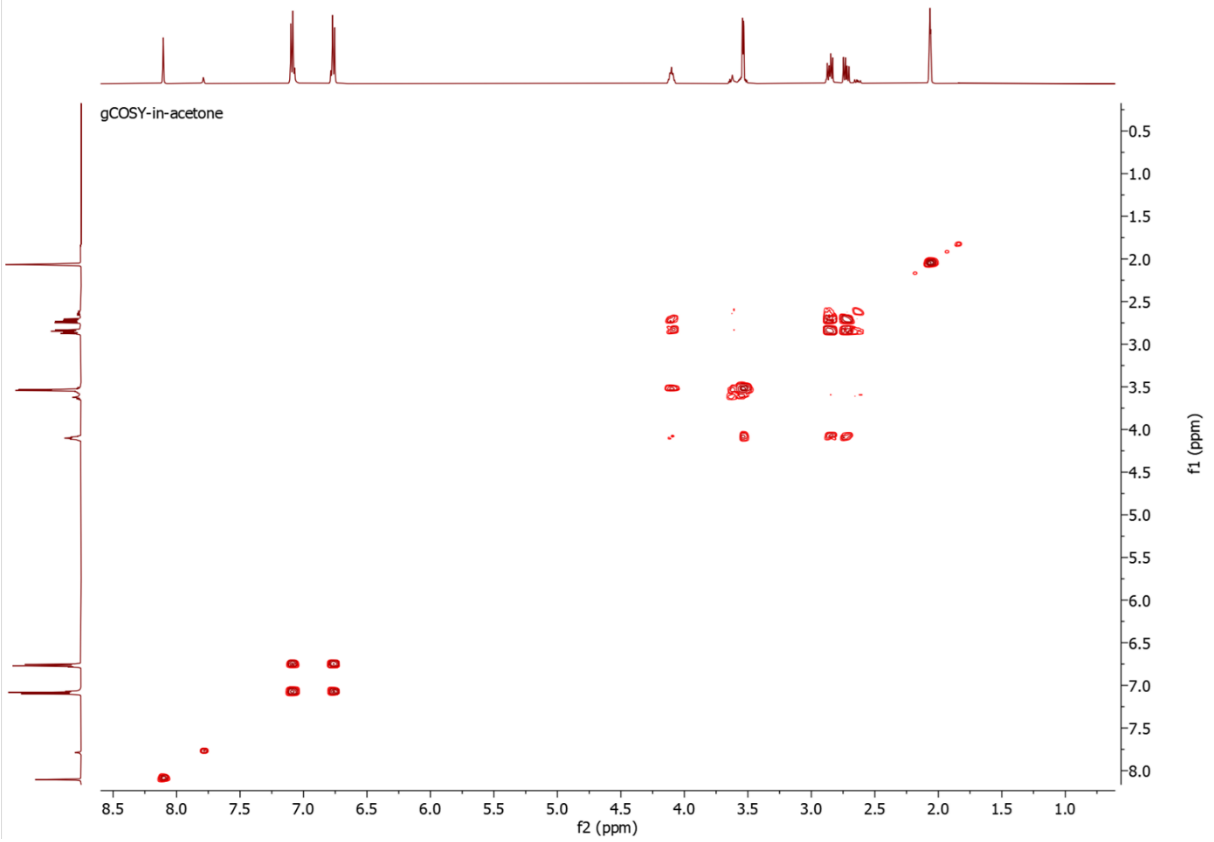


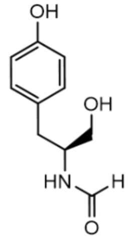

D

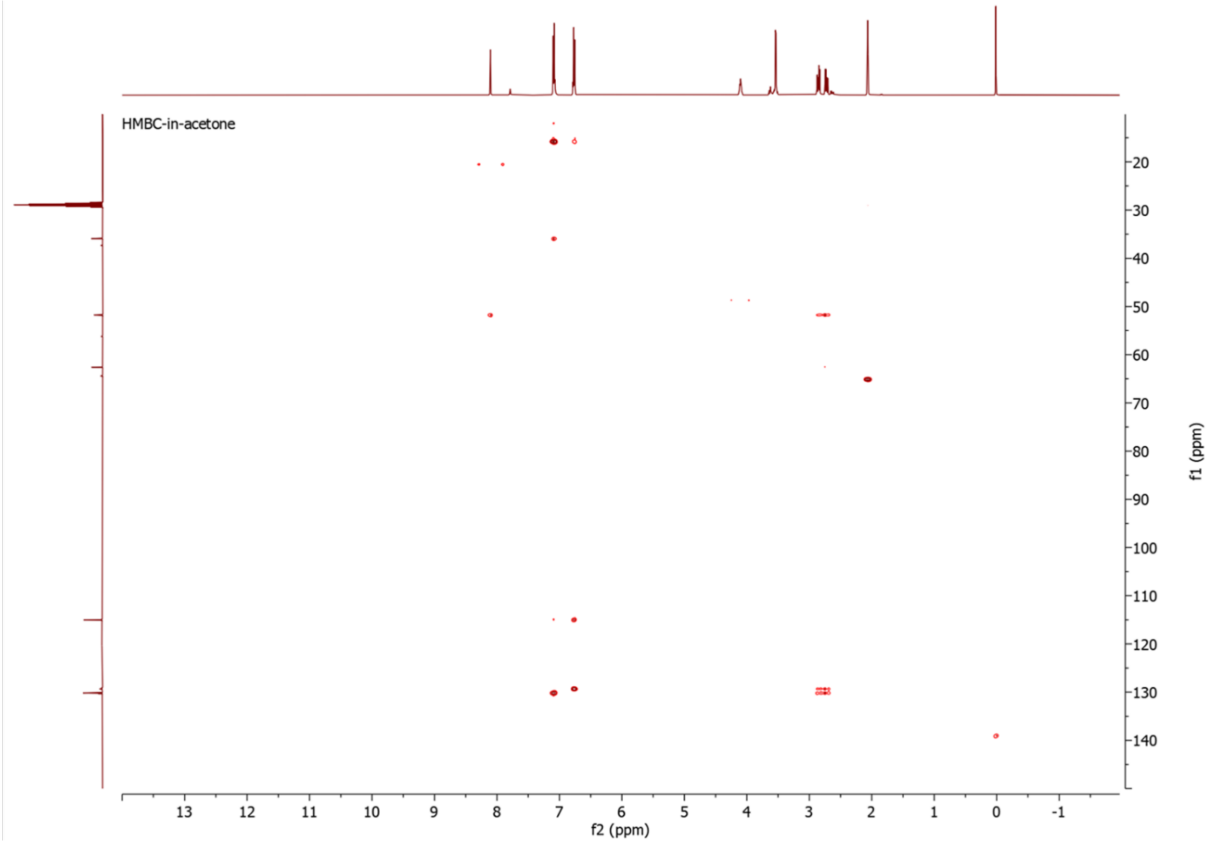

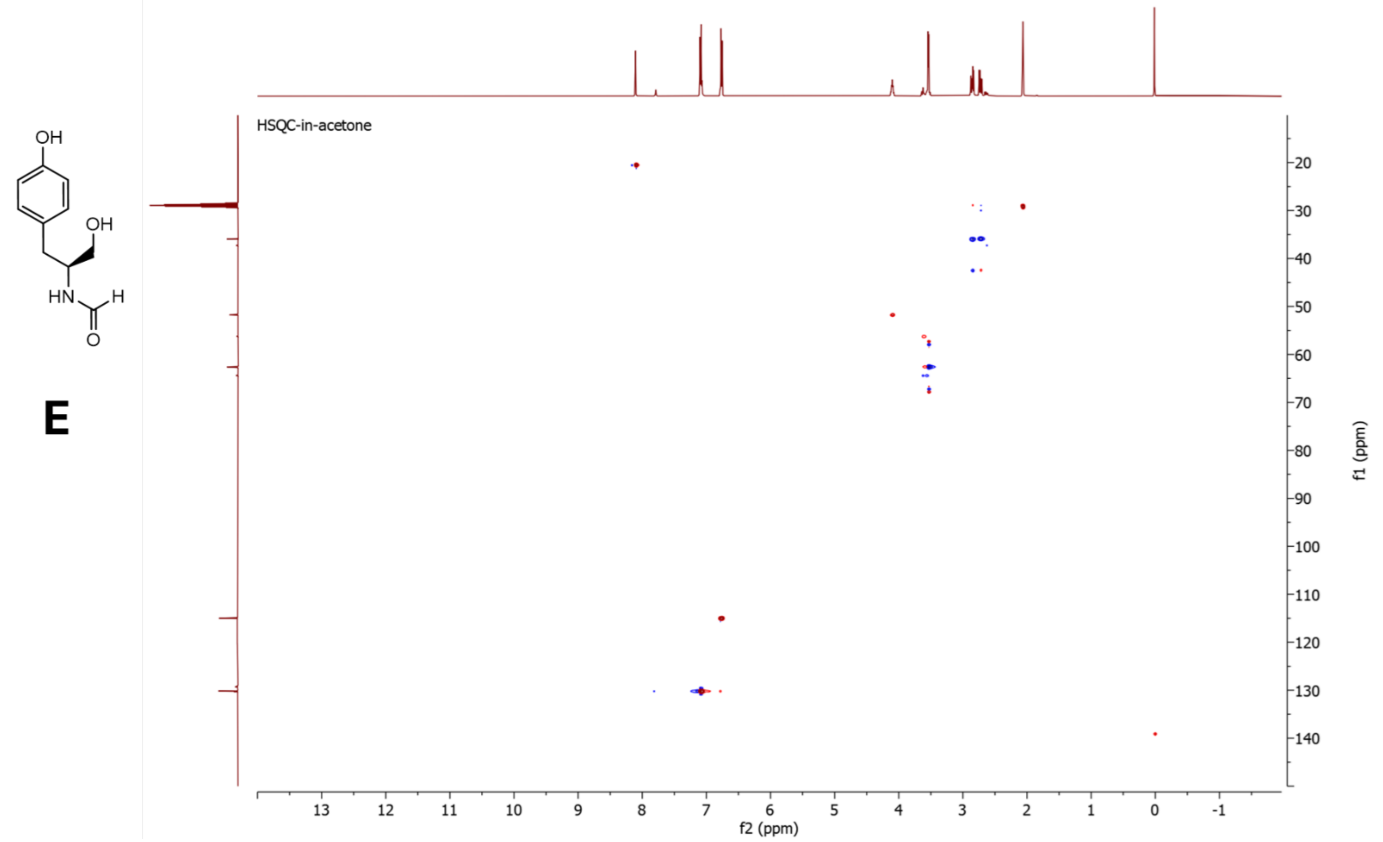


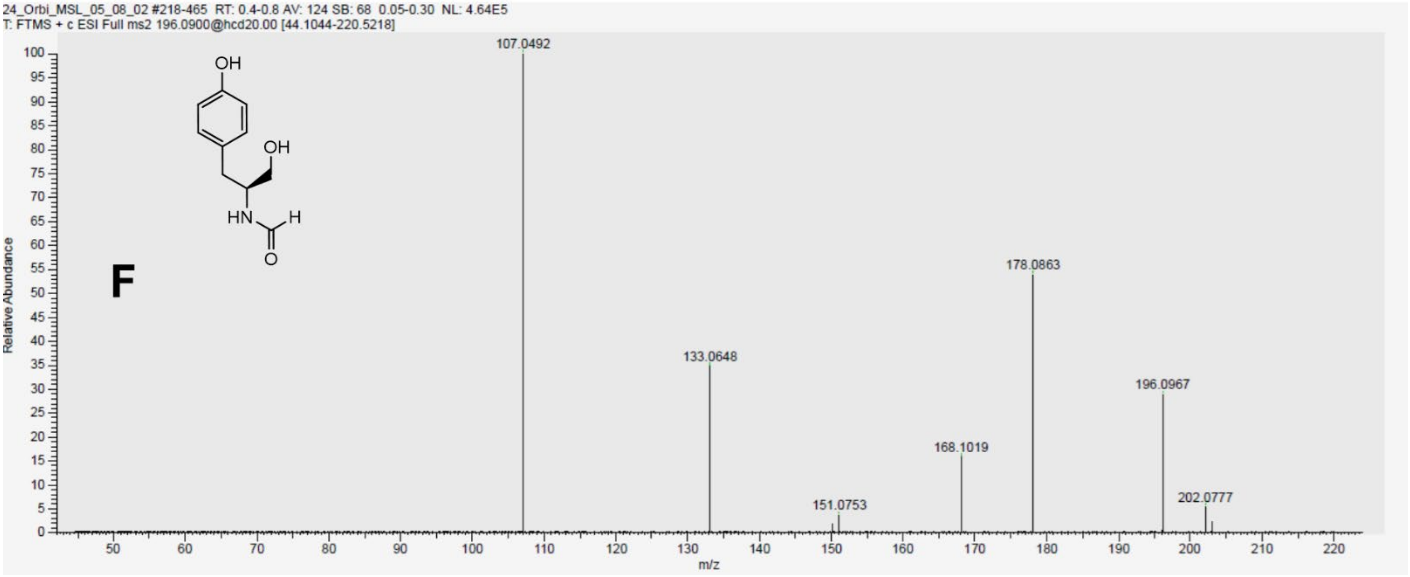


**Figure S48**. NMR data for **4a** in acetone-$d_6$, 500 MHz. **A)** $^{1}$H-NMR spectrum. **B)** $^{13}$C-NMR spectrum. **C)** COSY spectrum. **C)** HMBC spectrum. **E)** HSQC spectrum. **F)** HR-MS/MS spectrum. The CHO carbon in the HSQC is aliased

5.2.2 NMR and HR-MS/MS data for **4c**

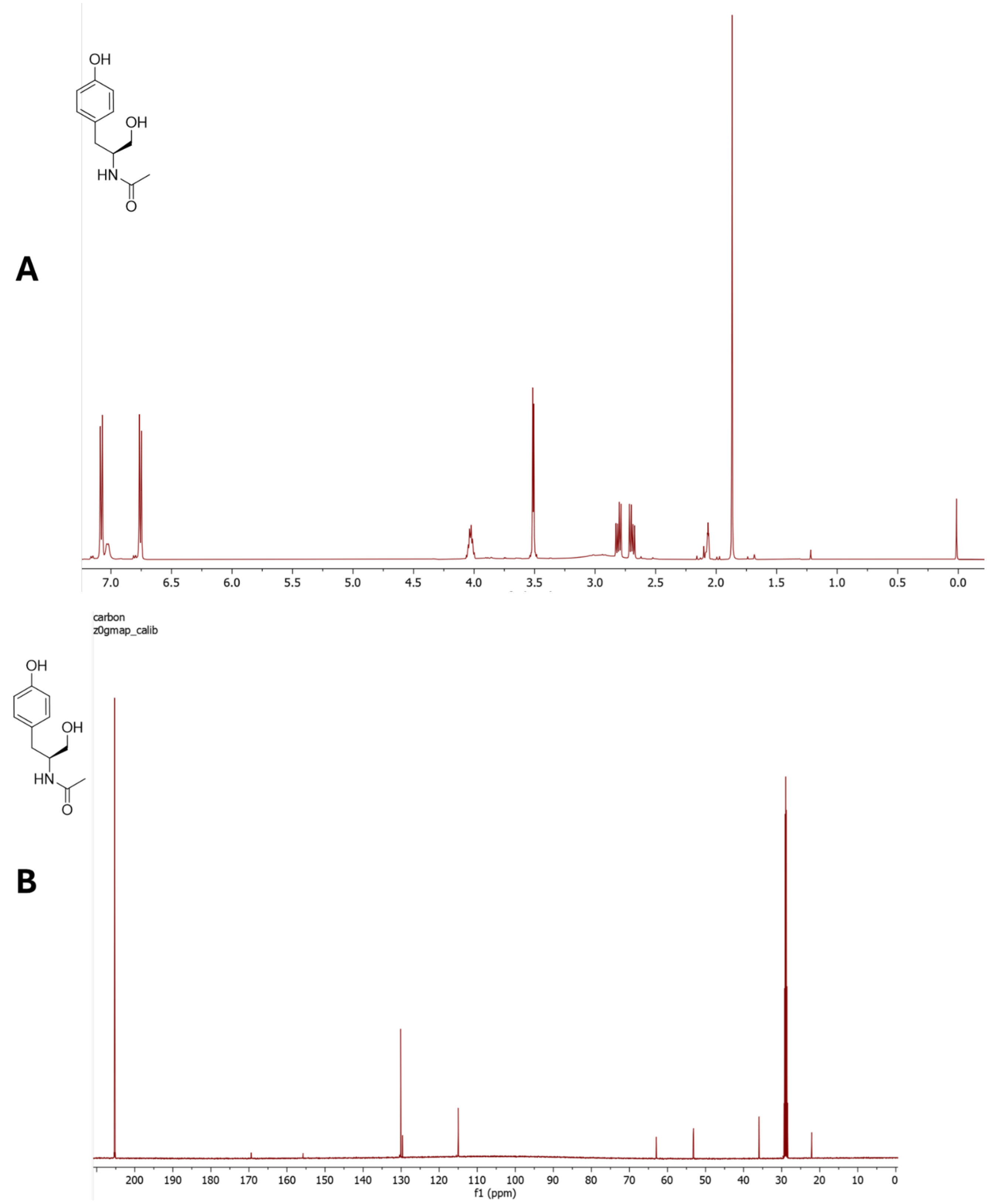

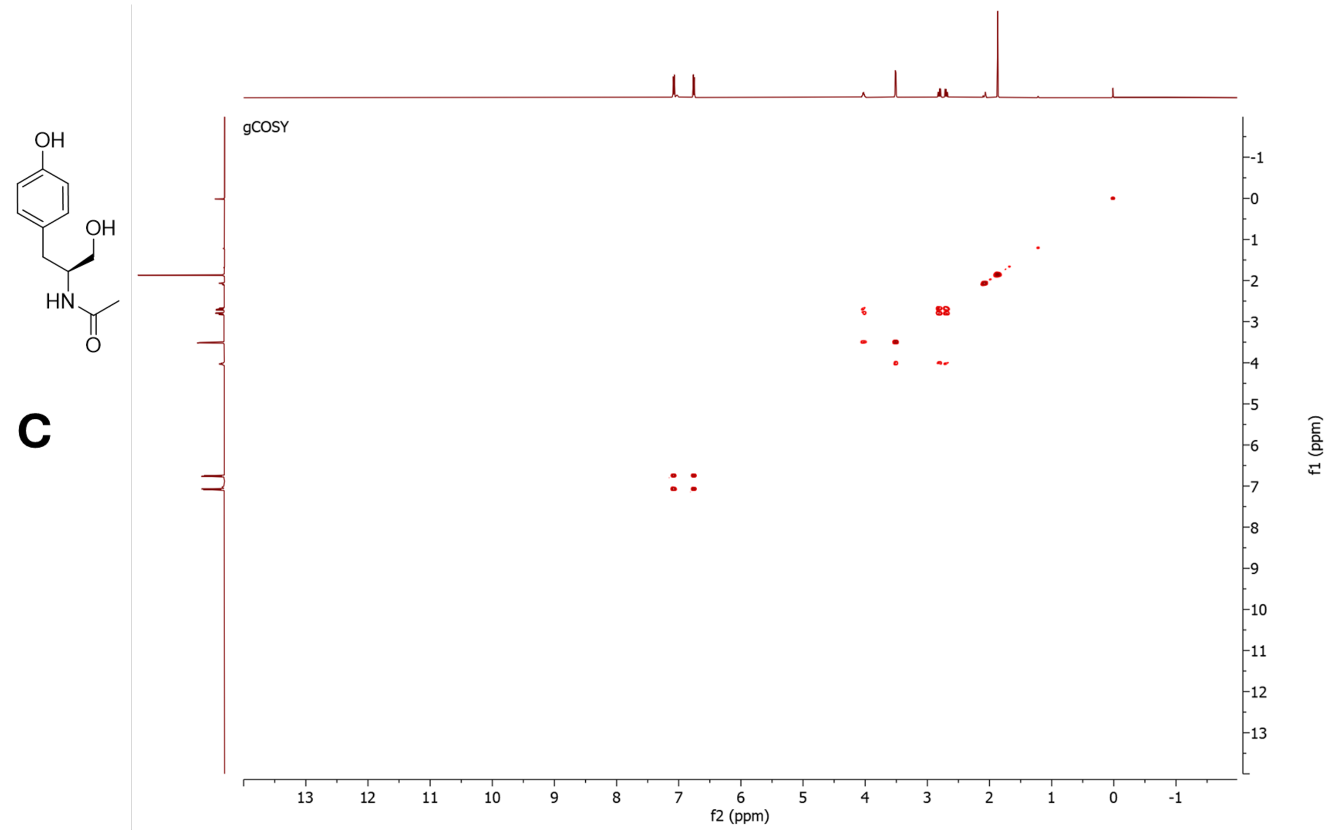


C

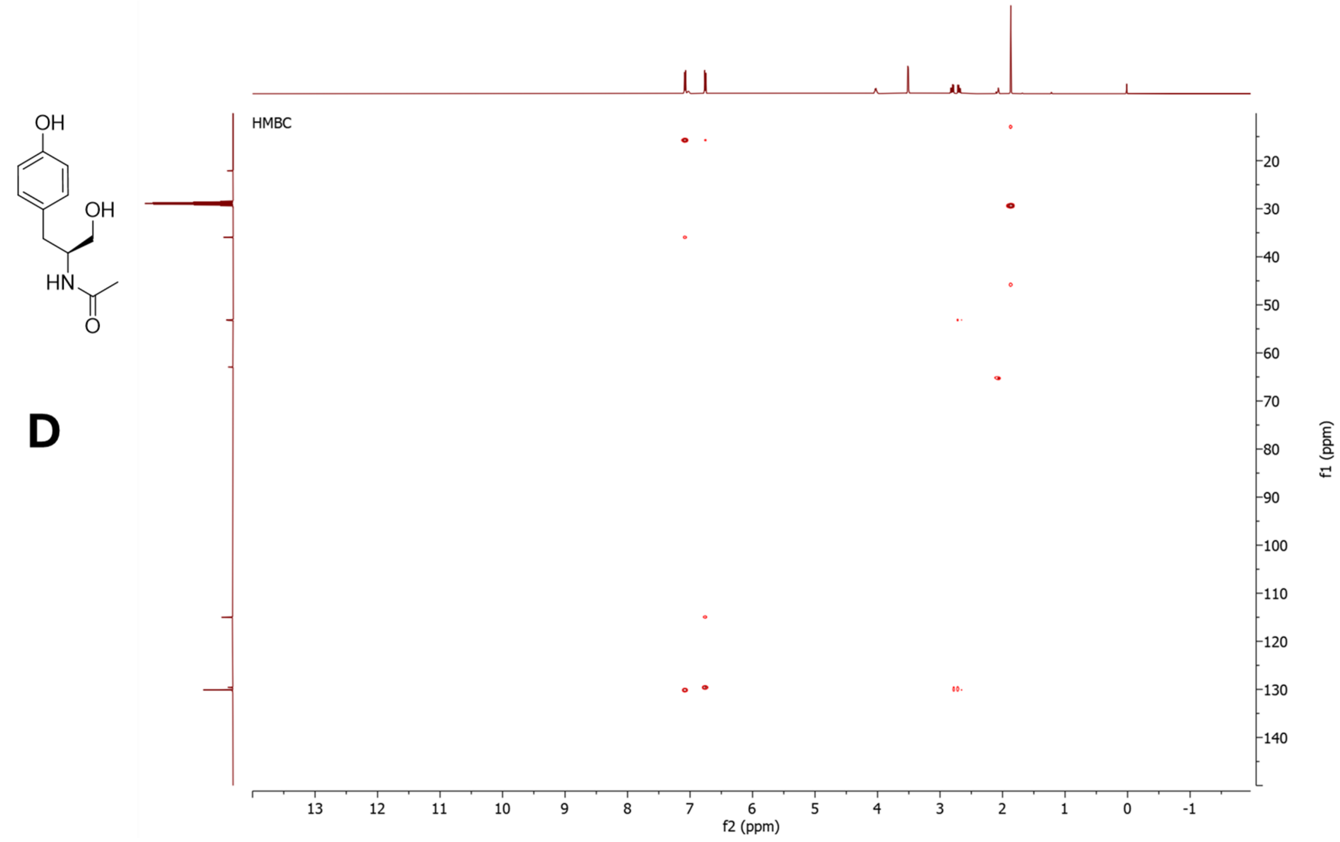


D

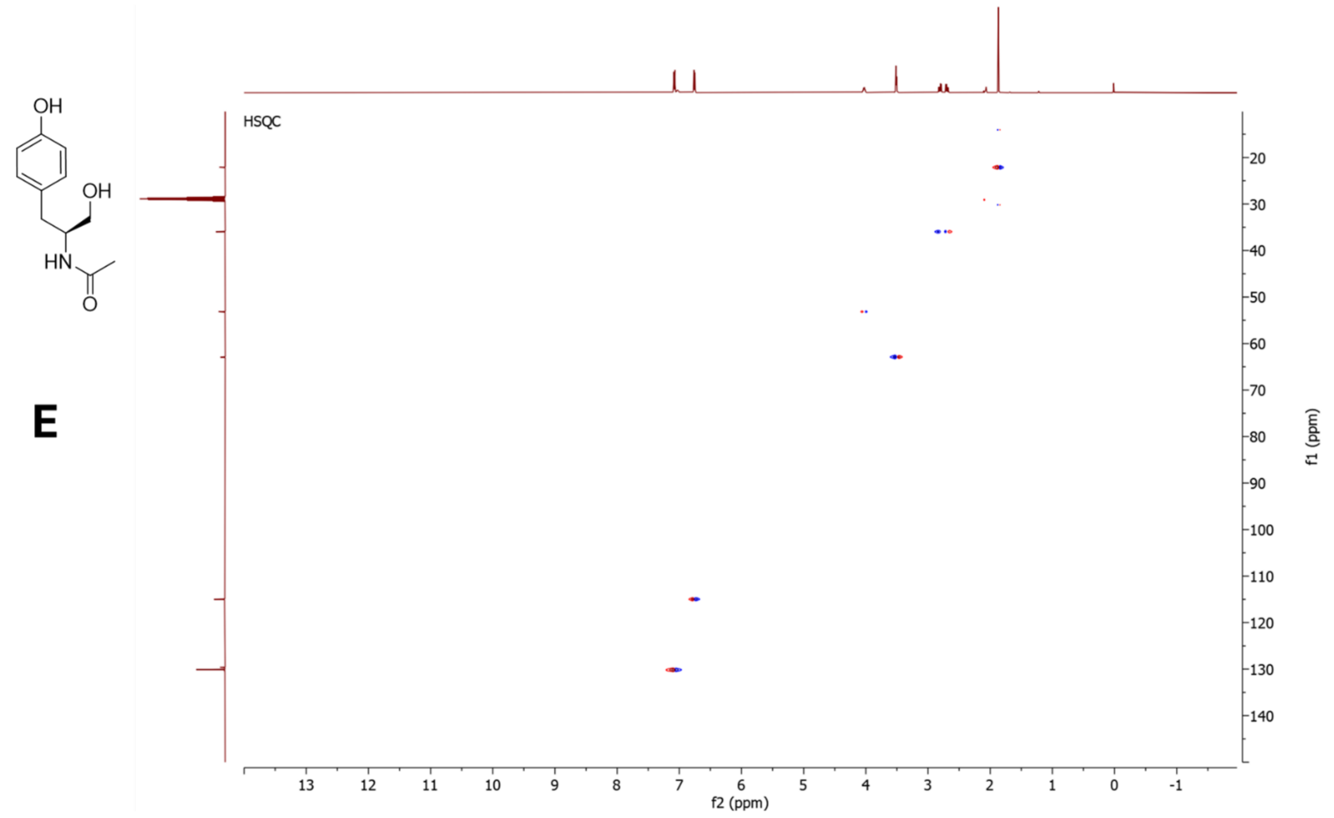


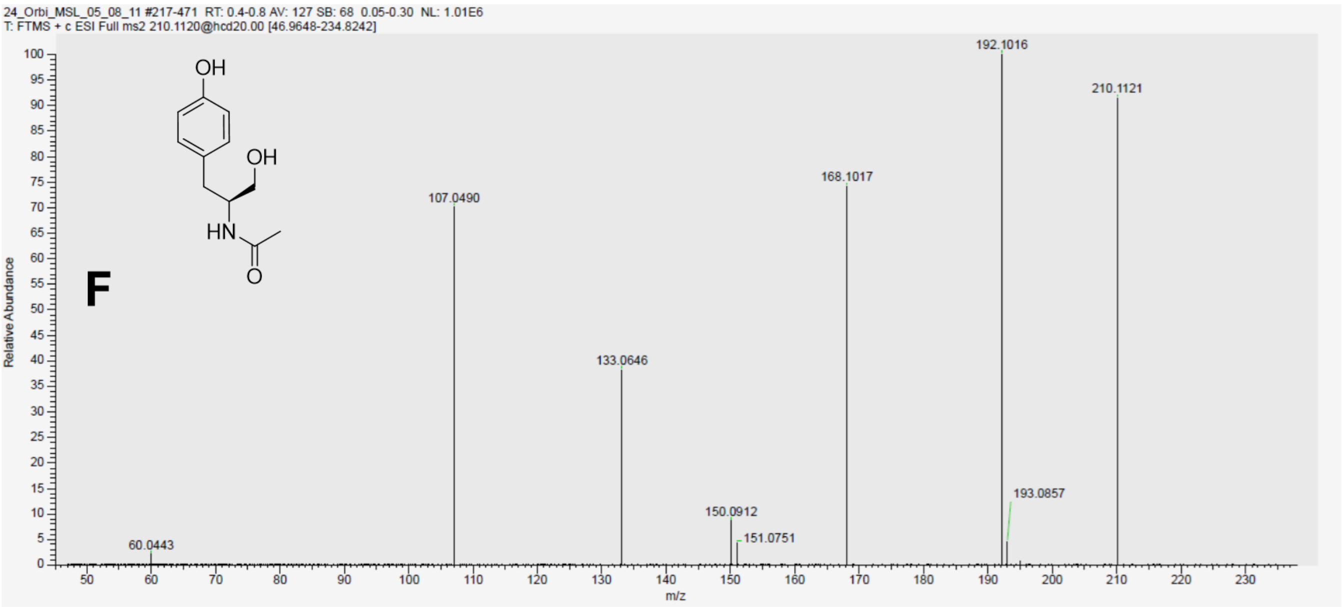


**Figure S49**. NMR data for **4c** in acetone-$d_6$, 500 MHz. **A)** $^1$H-NMR spectrum. **B)** $^{13}$C-NMR spectrum. **C)** COSY spectrum. **C)** HMBC spectrum. **E)** HSQC spectrum. **F)** HR-MS/MS spectrum.

5.2.3 NMR and HR-MS/MS data for **4d**

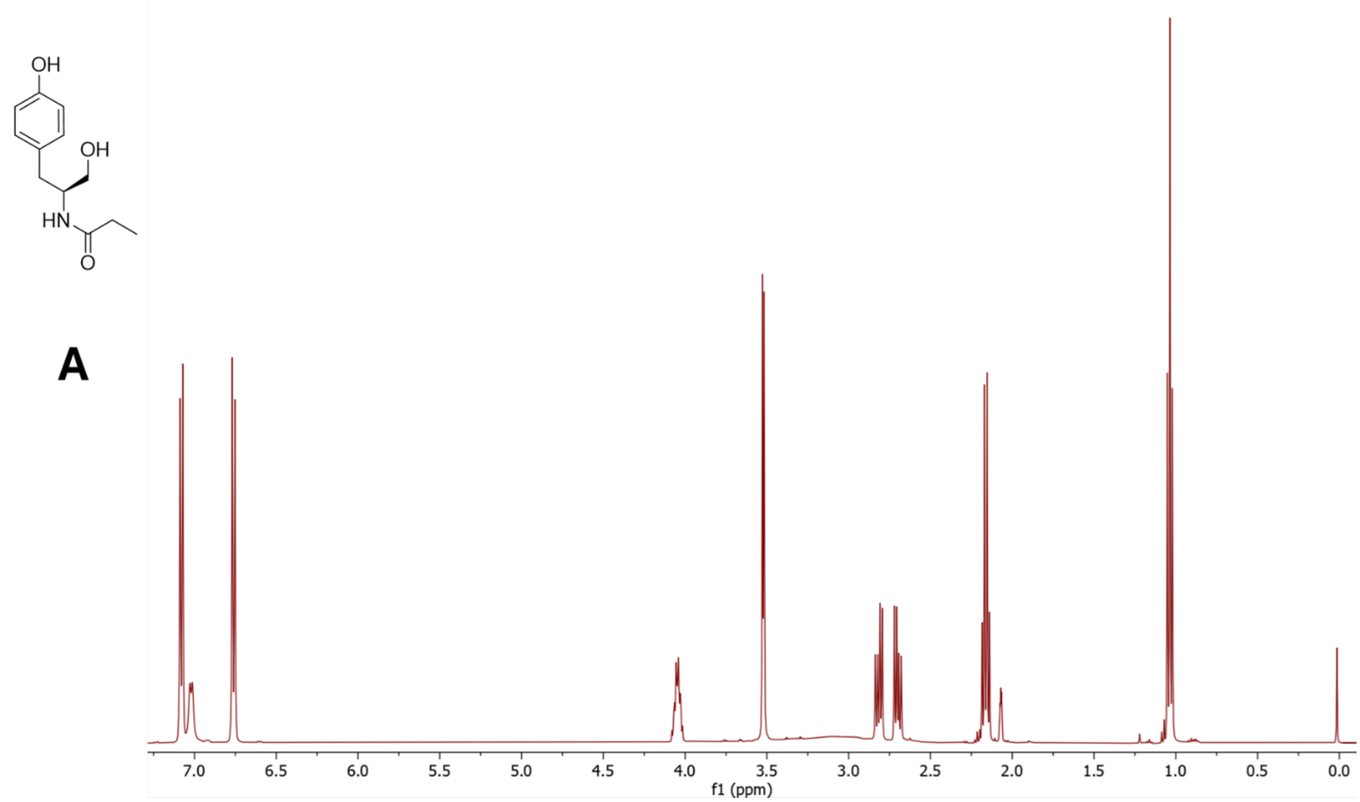


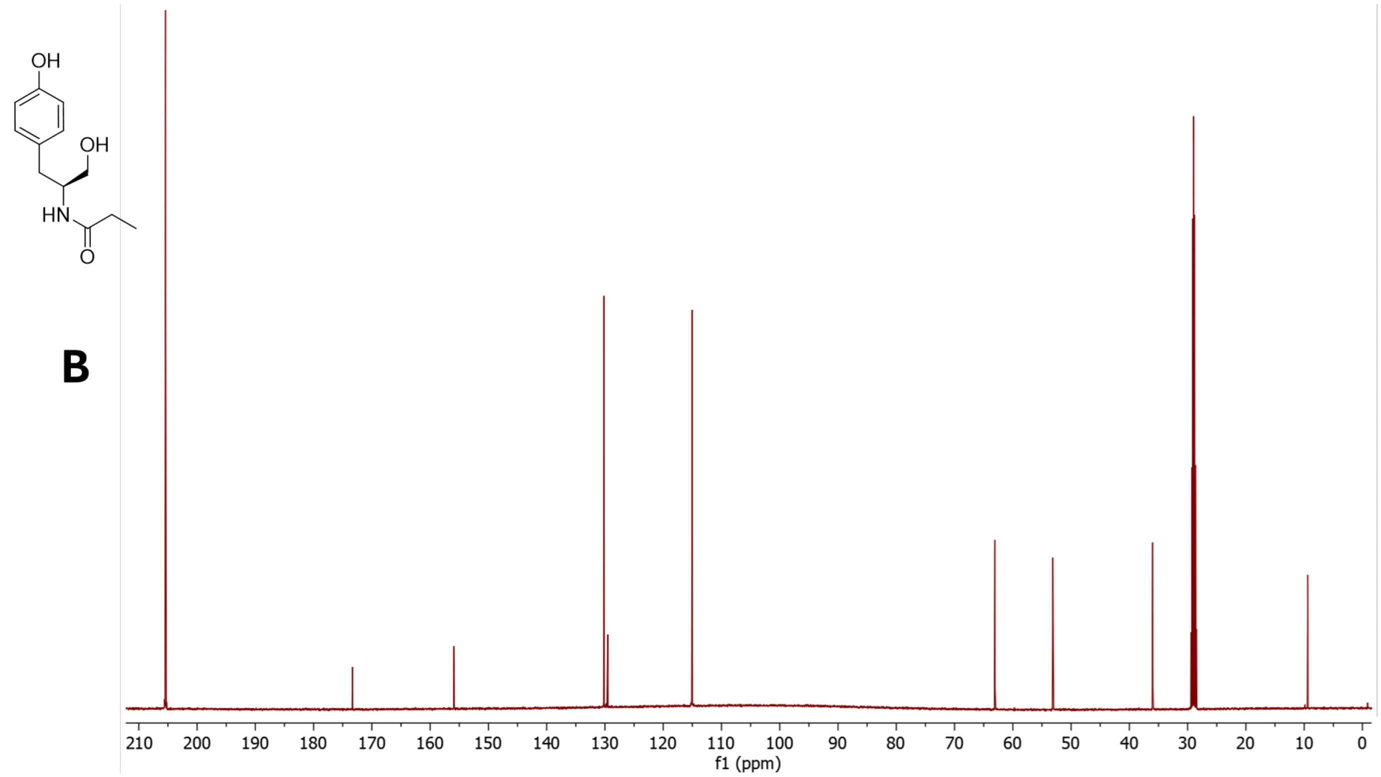

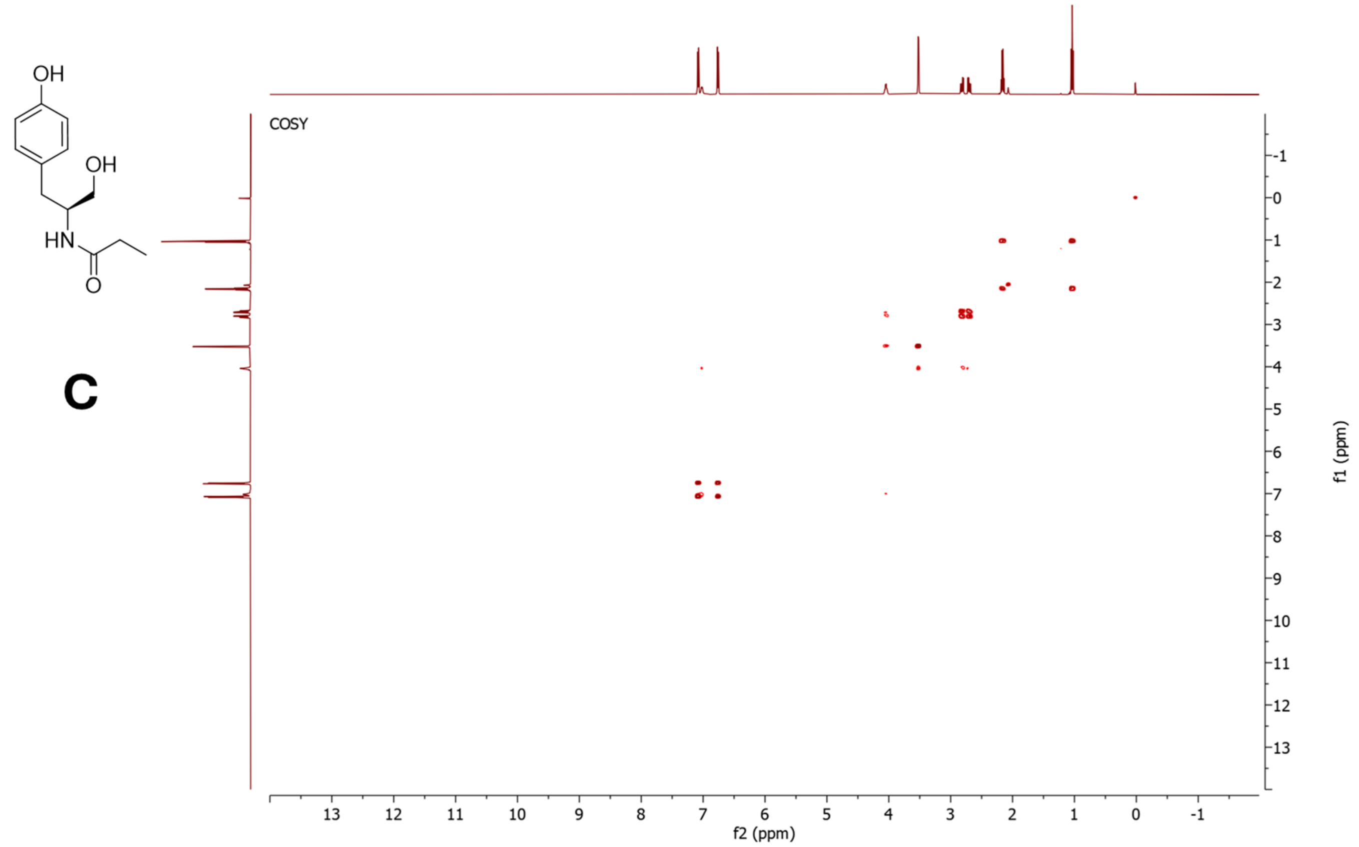


**C**

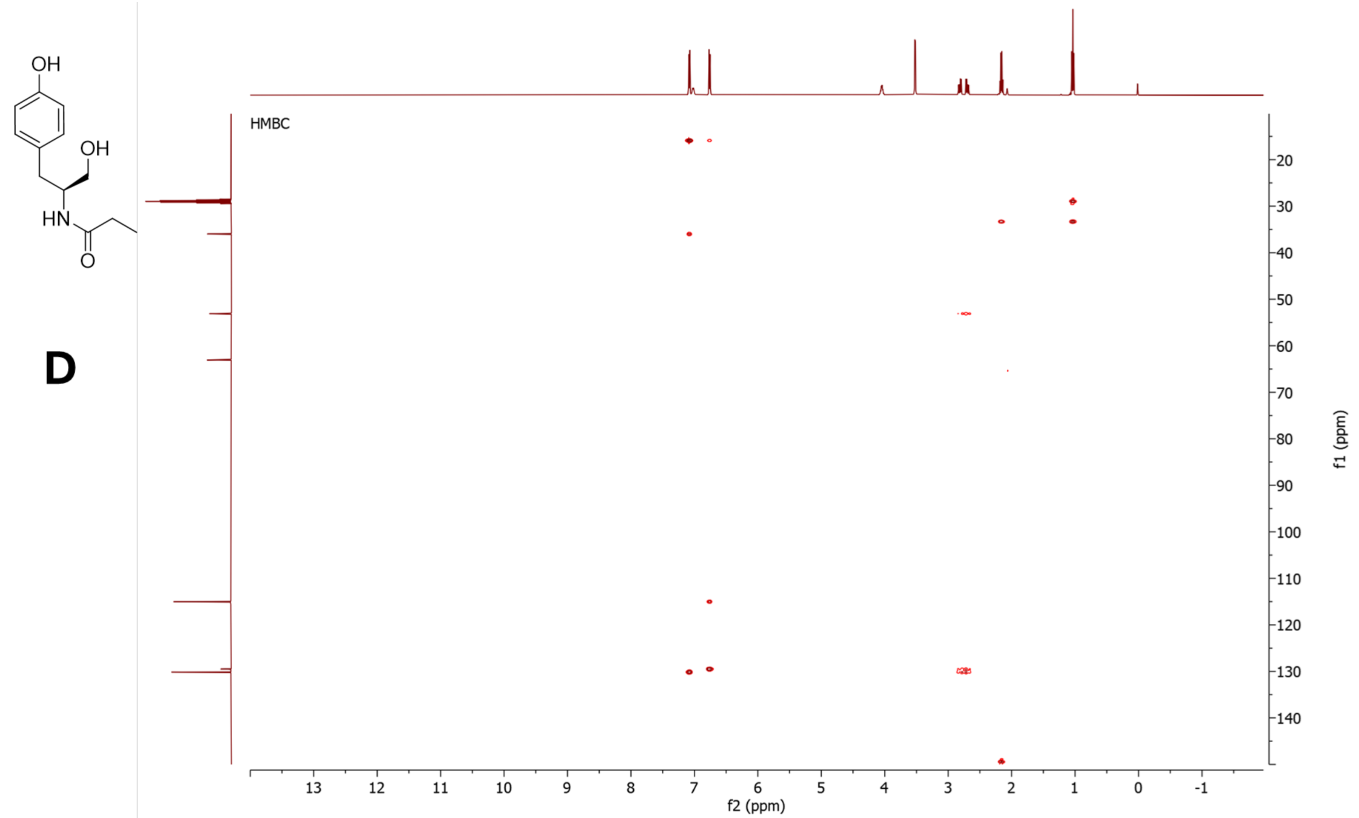


**D**

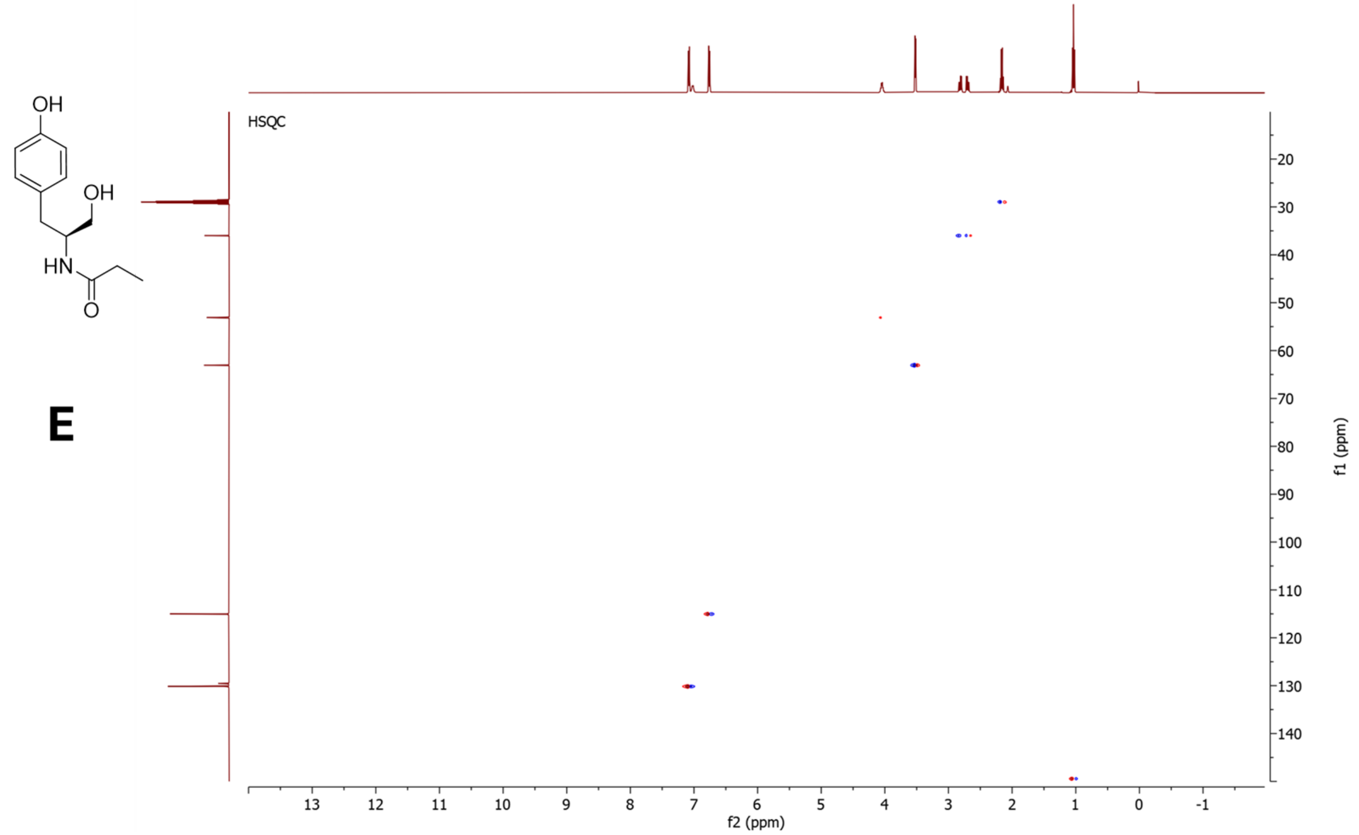


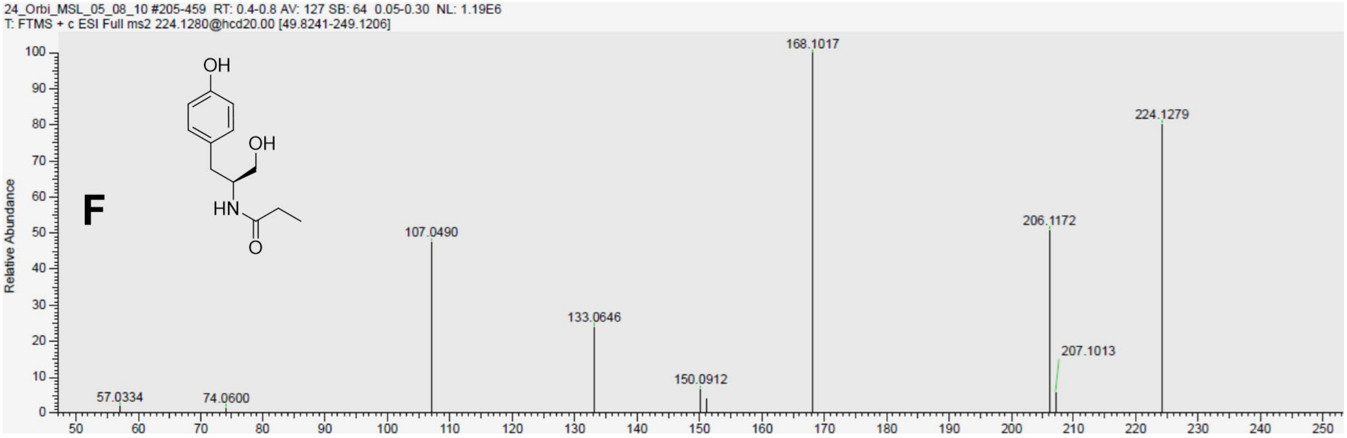


**Figure S50**. NMR data for **4d** in acetone-$d_6$, 500 MHz. **A)** $^{1}$H-NMR spectrum. **B)** $^{13}$C-NMR spectrum. **C)** COSY spectrum. **C)** HMBC spectrum. **E)** HSQC spectrum. **F)** HR-MS/MS spectrum.

5.2.4 NMR and HR-MS/MS data for **4e**

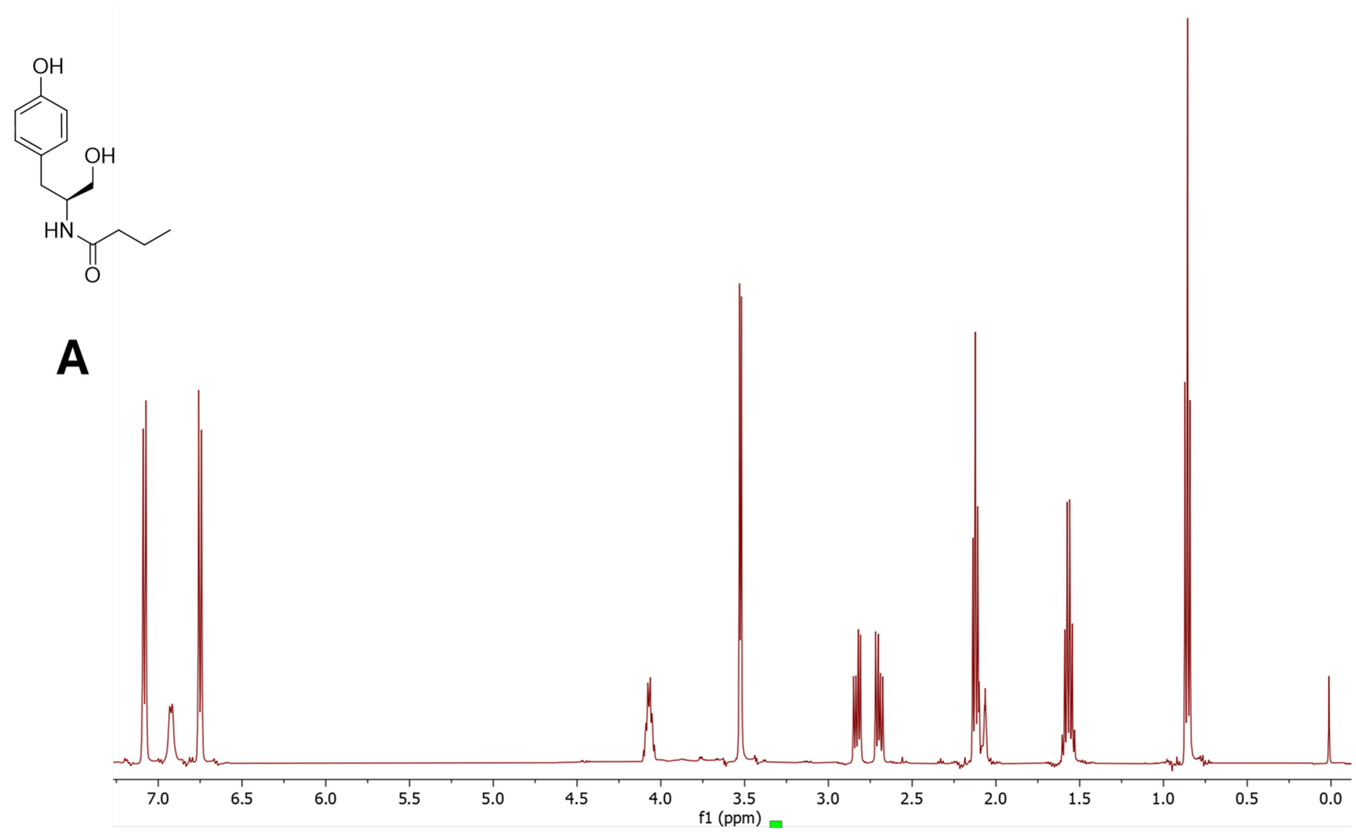


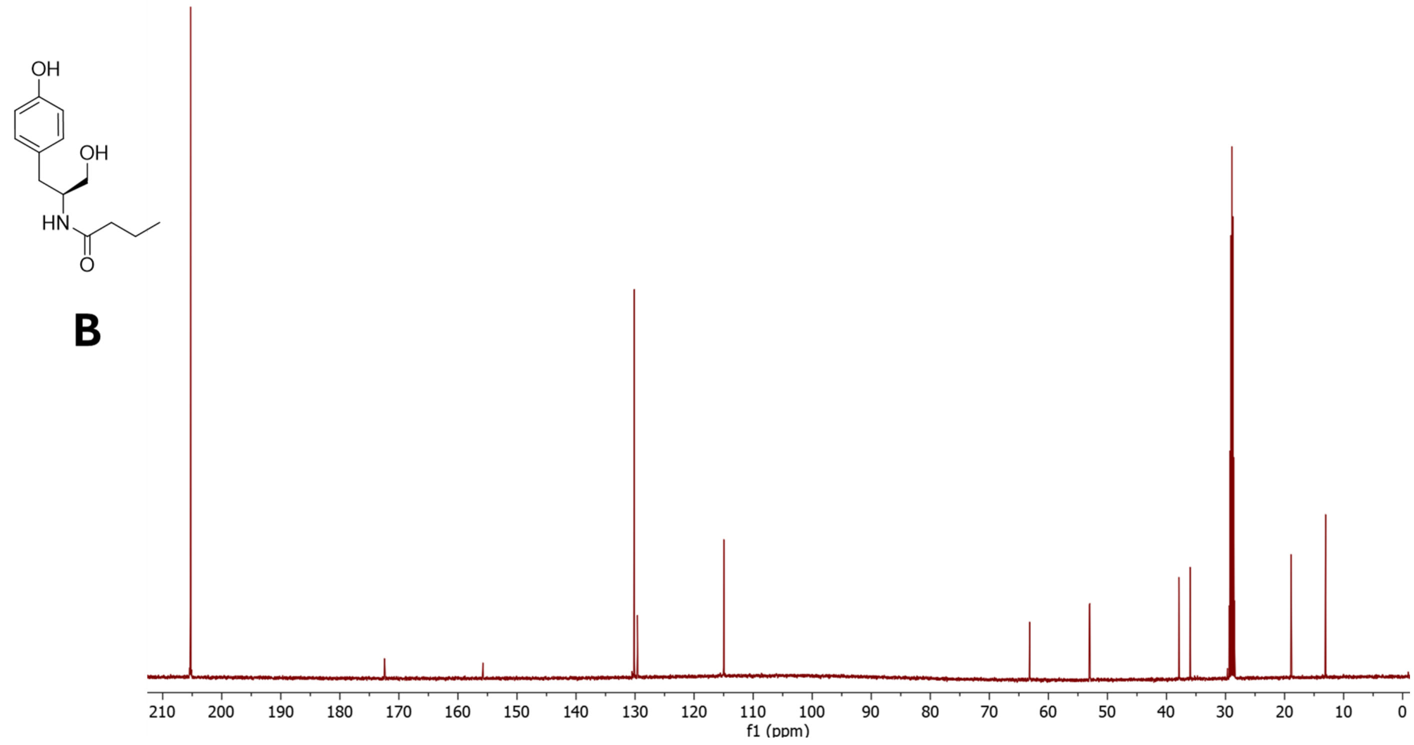

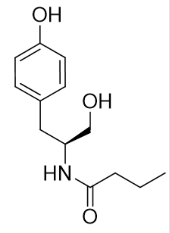


C

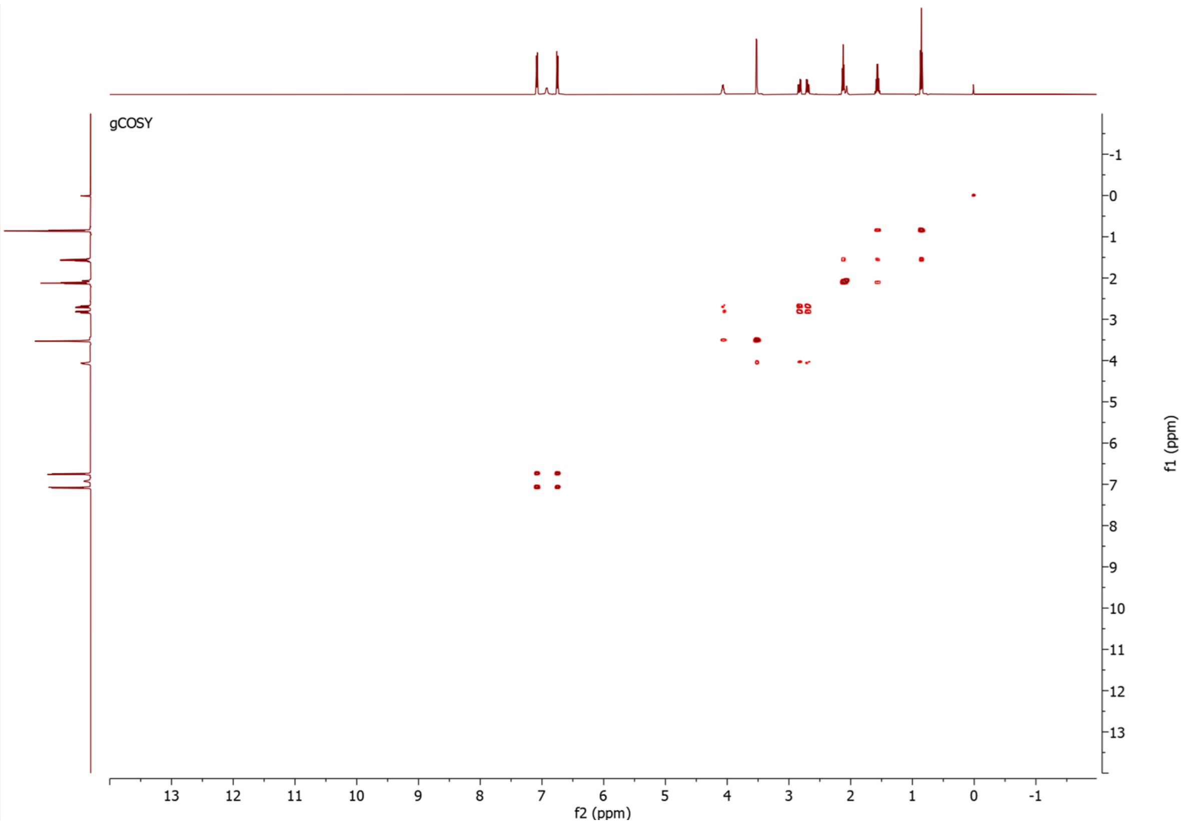


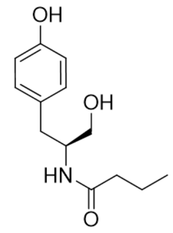


D

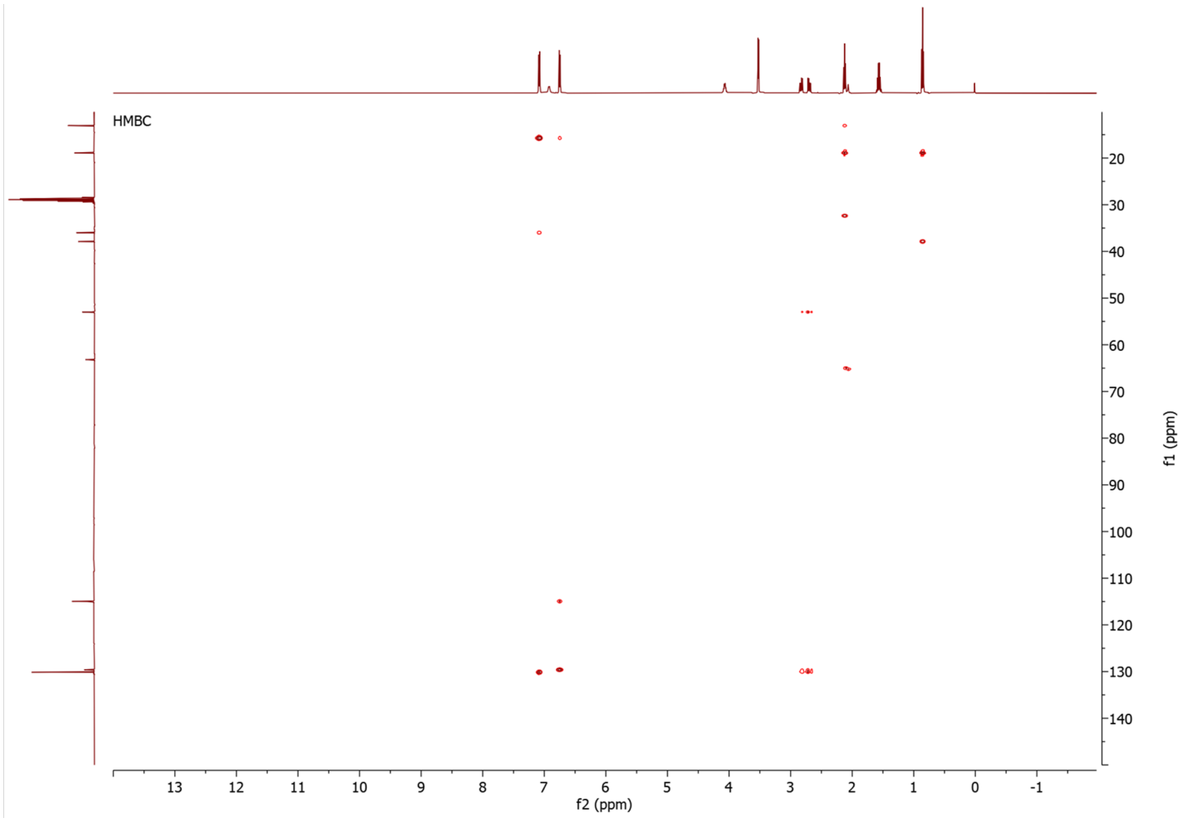

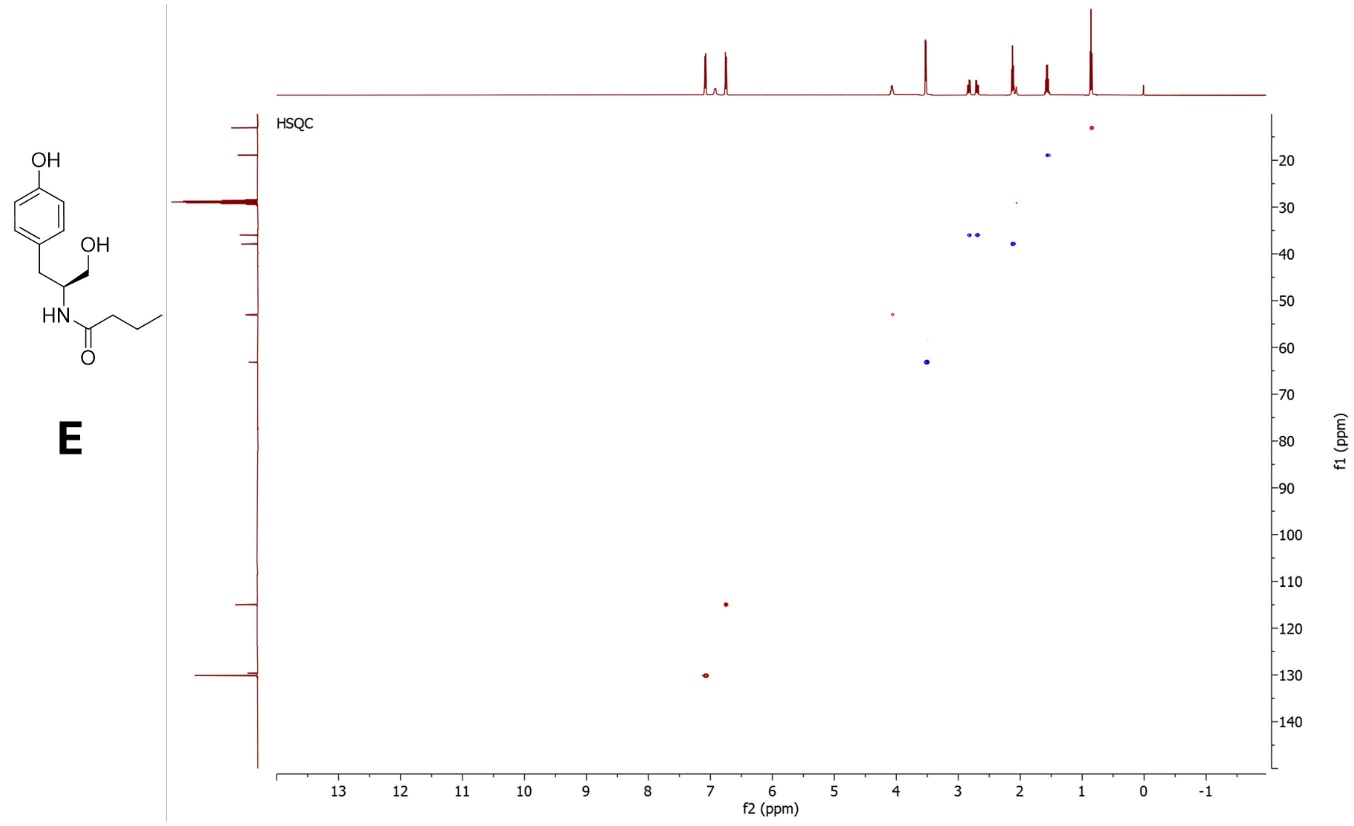


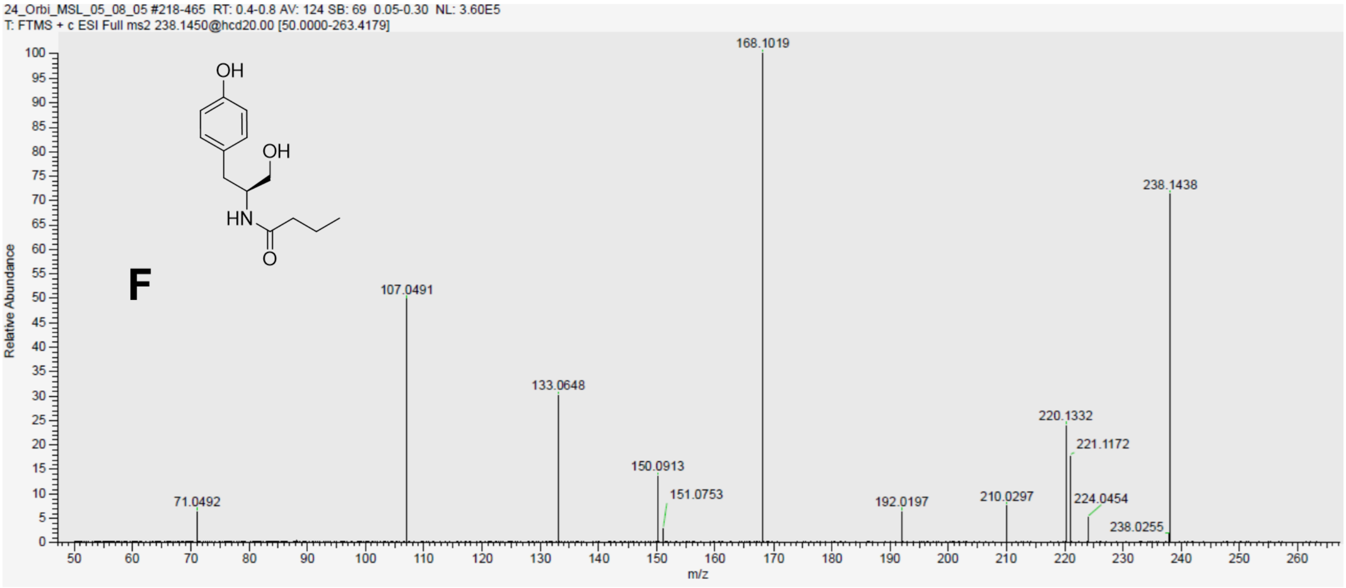


**Figure S51**. NMR data for **4e** in acetone-$d_6$, 500 MHz. **A)** $^{1}$H-NMR spectrum. **B)** $^{13}$C-NMR spectrum. **C)** COSY spectrum. **C)** HMBC spectrum. **E)** HSQC spectrum. **F)** HR-MS/MS spectrum.

5.2.5 HR-MS/MS data for **4b**, **4c***, **4f**

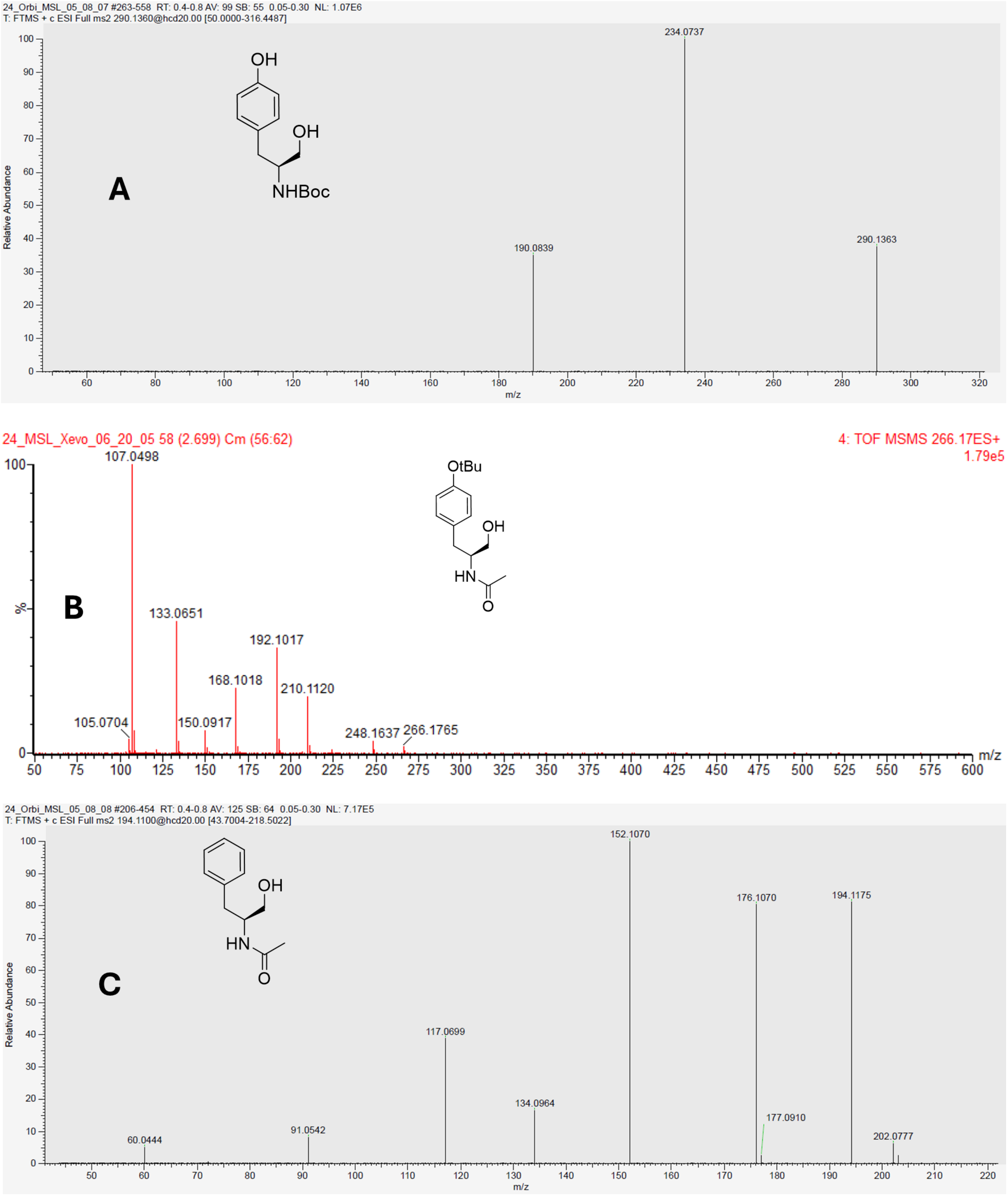


**Figure S52**. HR-MS/MS spectrum of **A) 4b**. **B) 4c***. **C) 4f**.

5.3 NMR data for *N*-acyl-L-tyrosyl coenzyme A thioesters

5.3.1 NMR data for **3Ca**

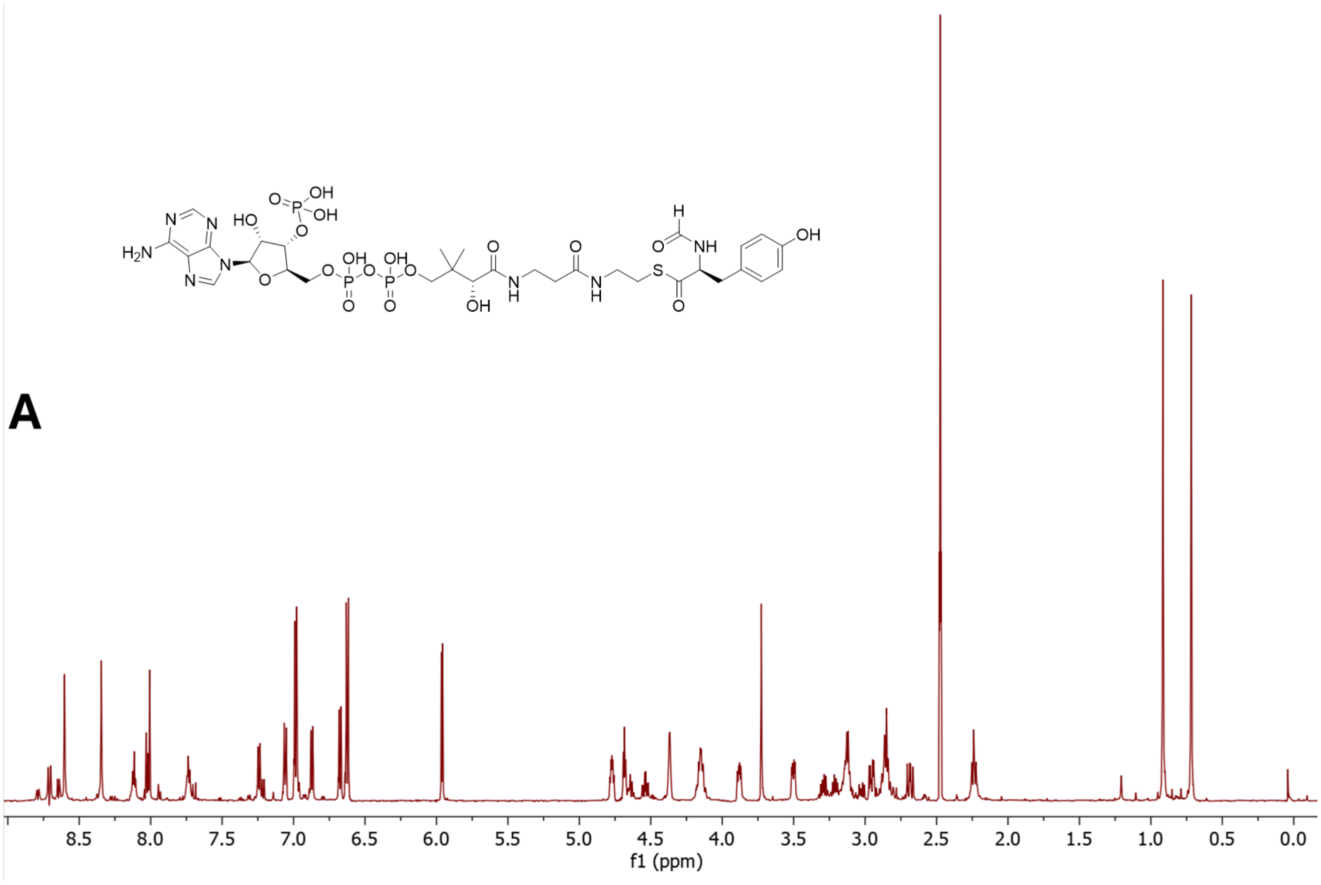

**B**

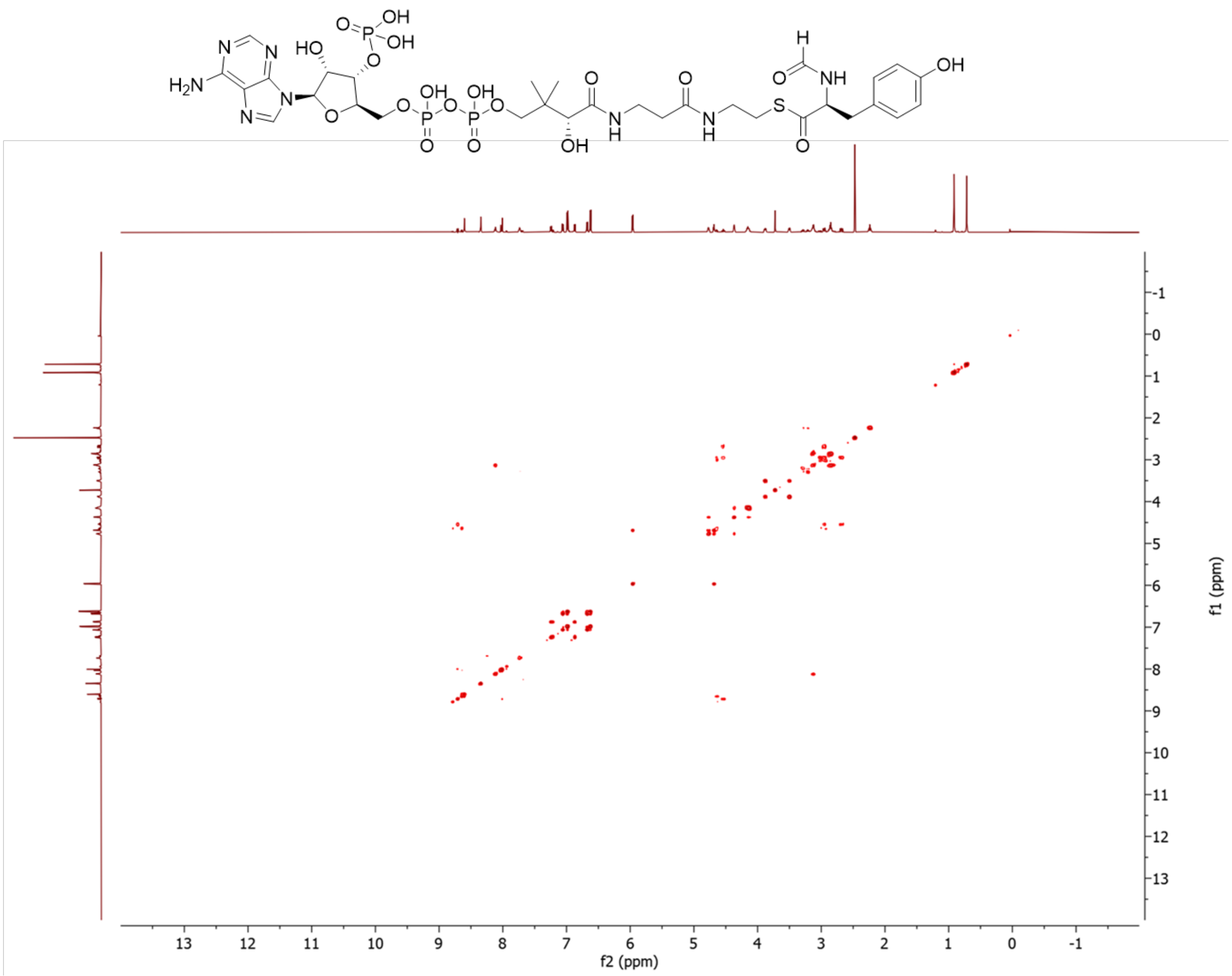

C

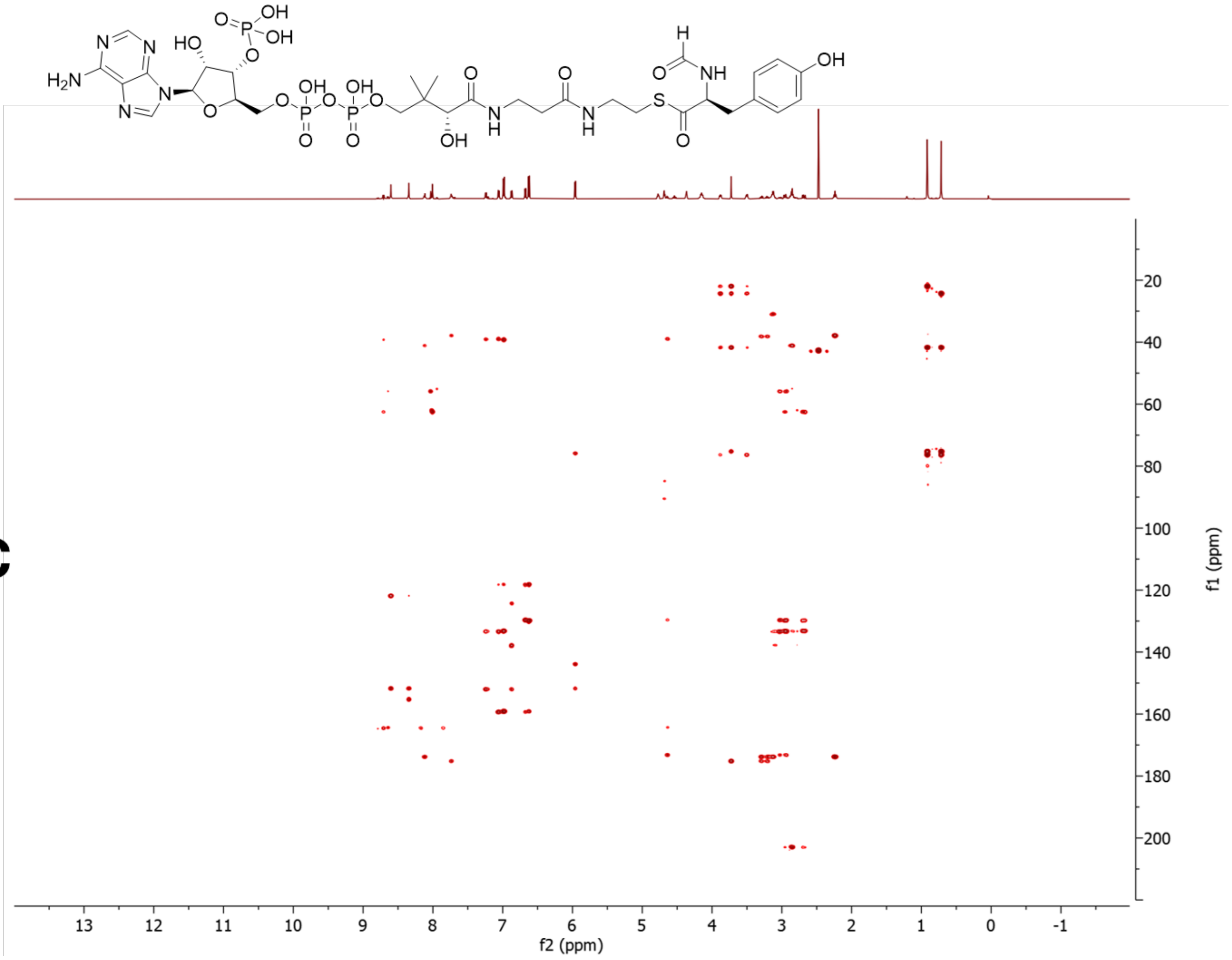

D

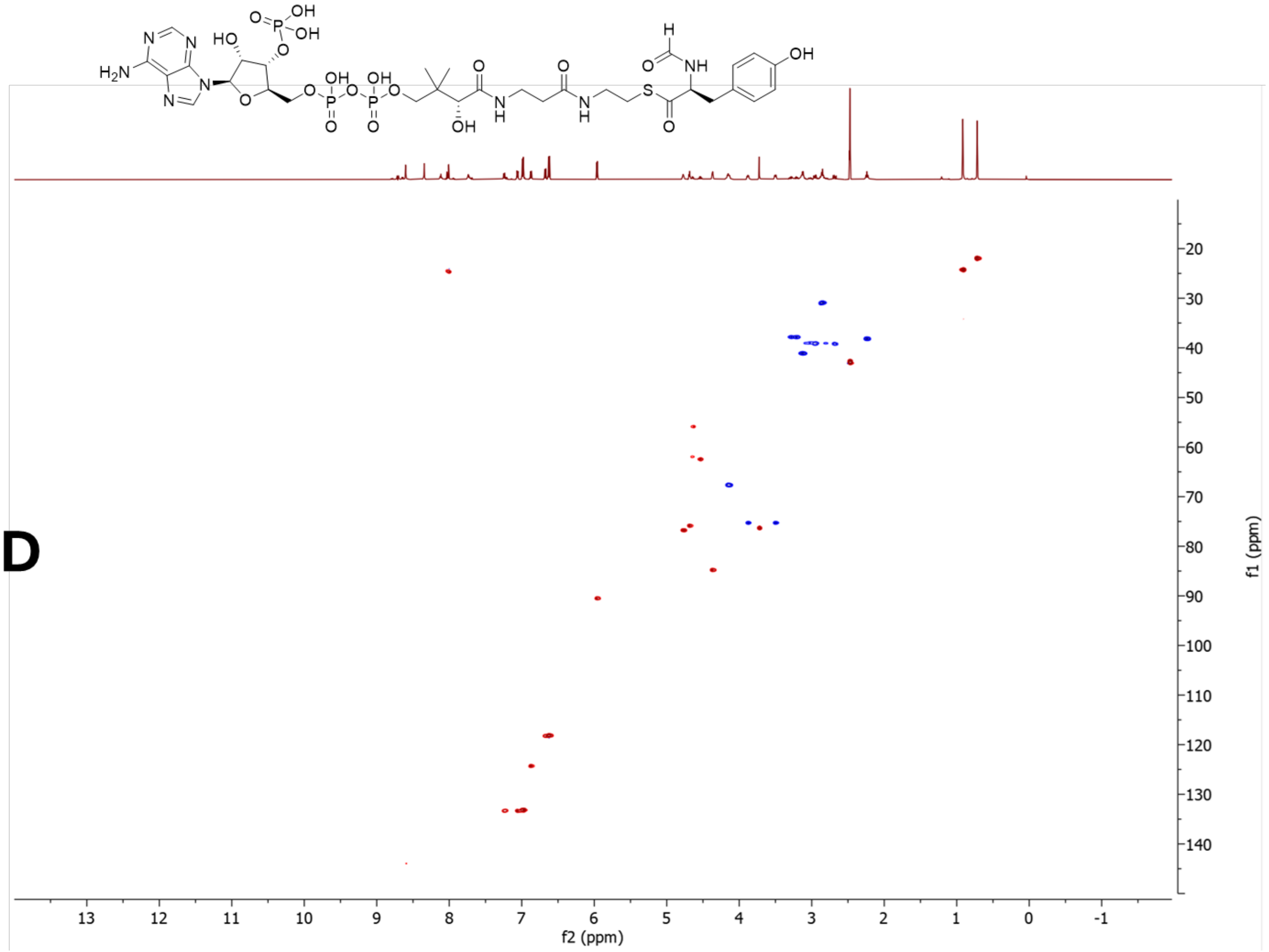


**Figure S53:** NMR data for **3Ca** in DMSO-$d_6$, 600 MHz. **A)** $^1$H-NMR spectrum. **B)** COSY spectrum. **C)** HMBC spectrum. **D)** HSQC spectrum. The CHO carbon in the HSQC is aliased.

5.3.2 NMR data for **3Cb**

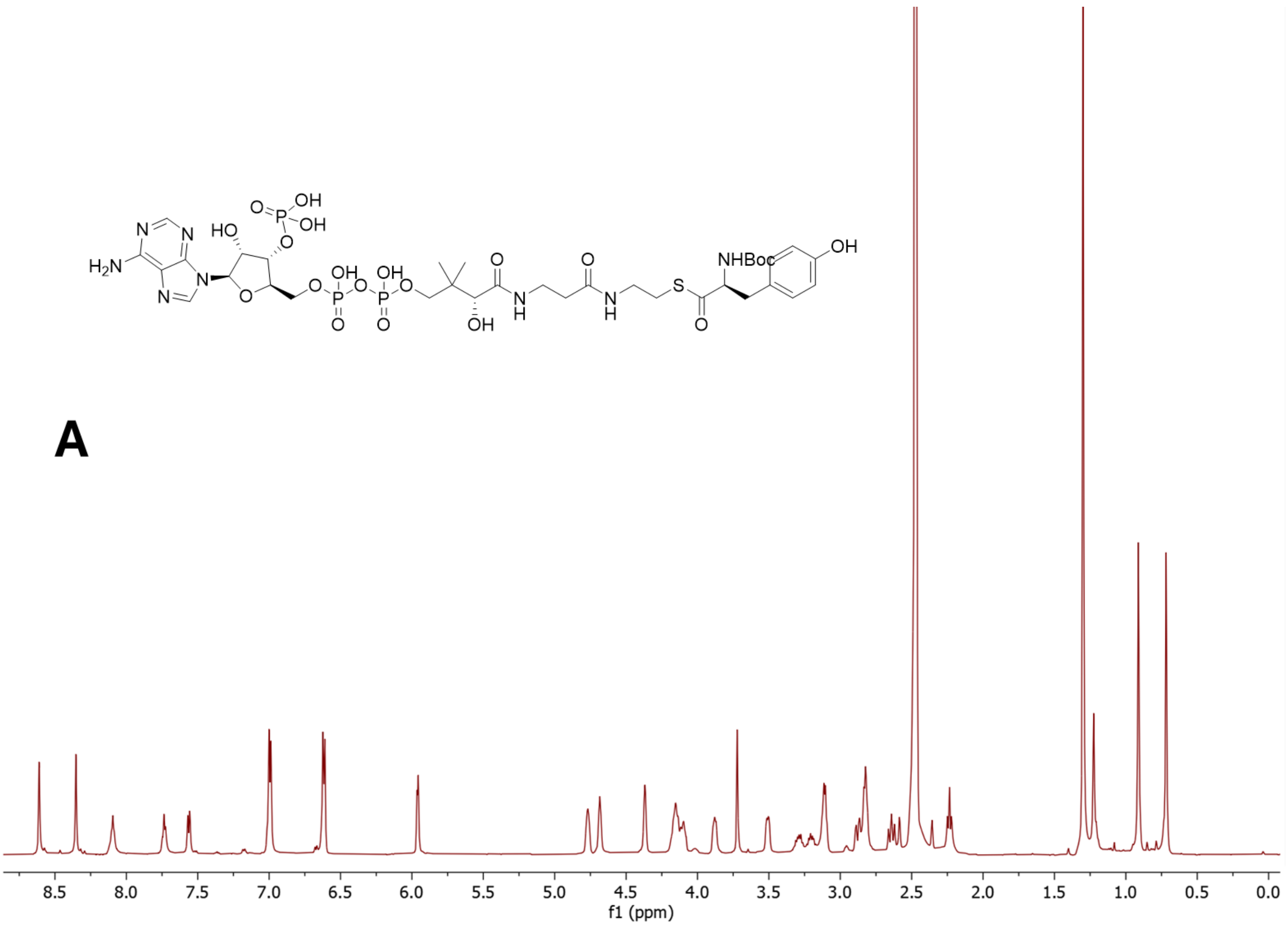

B

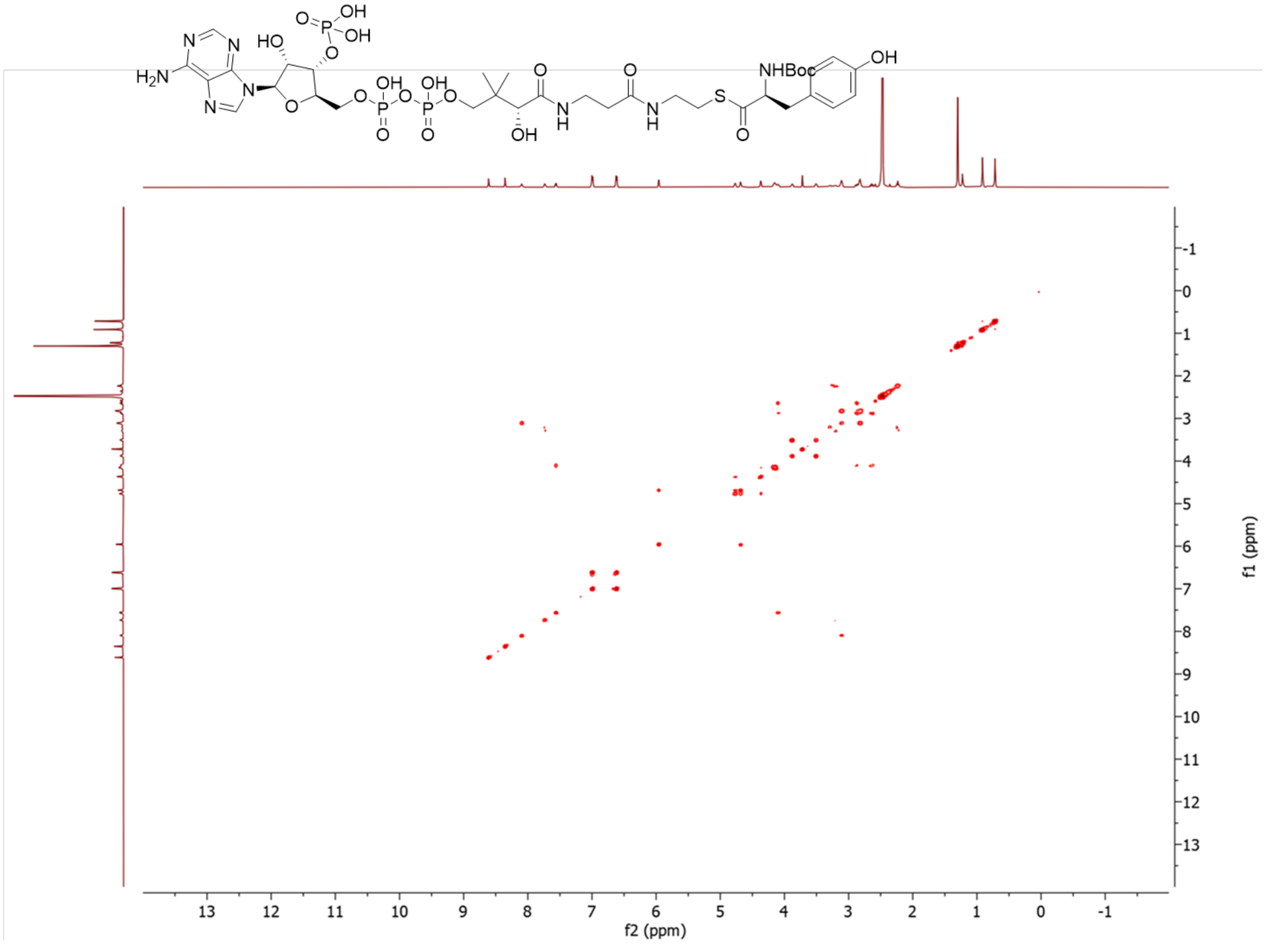

C

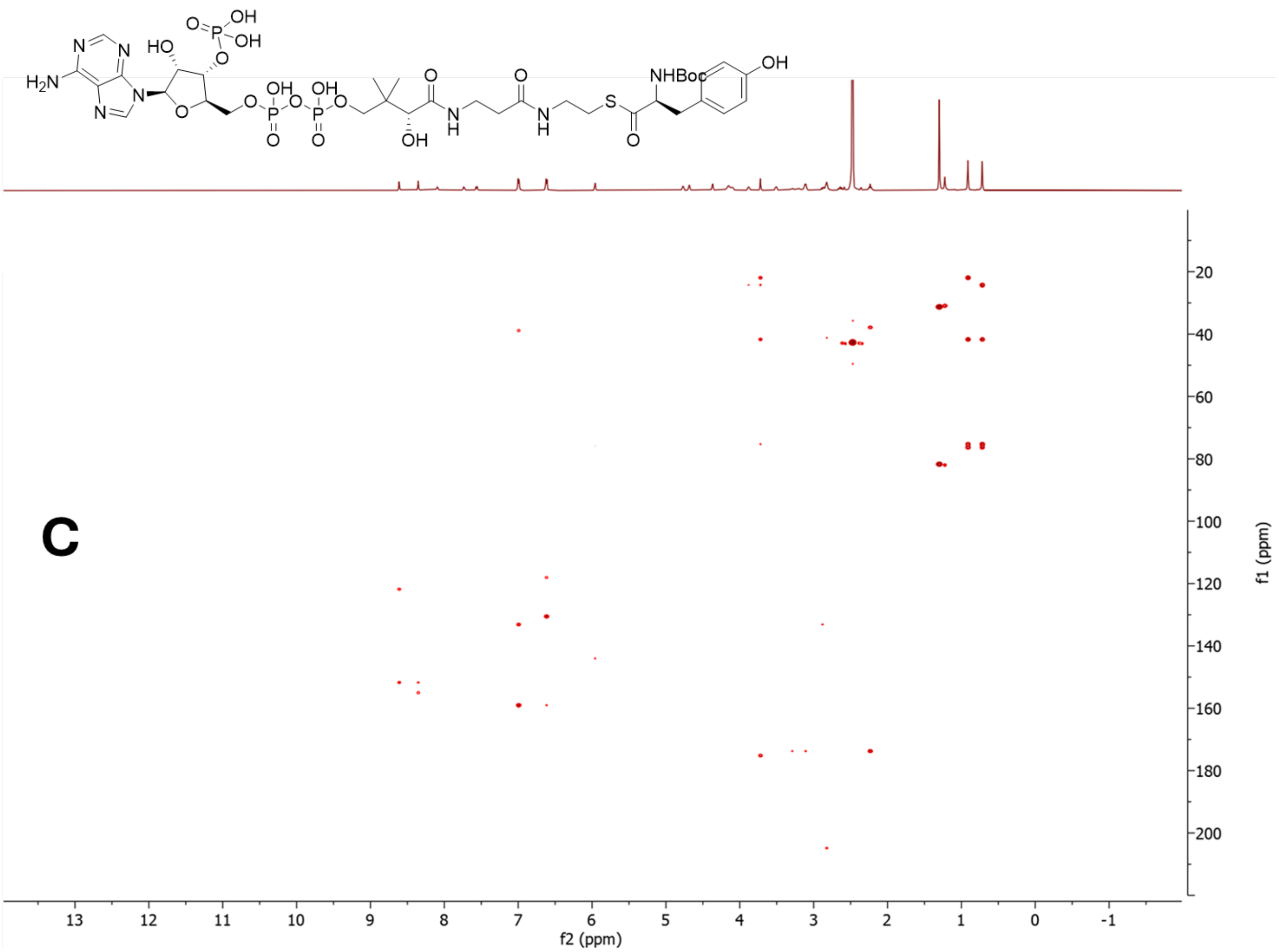

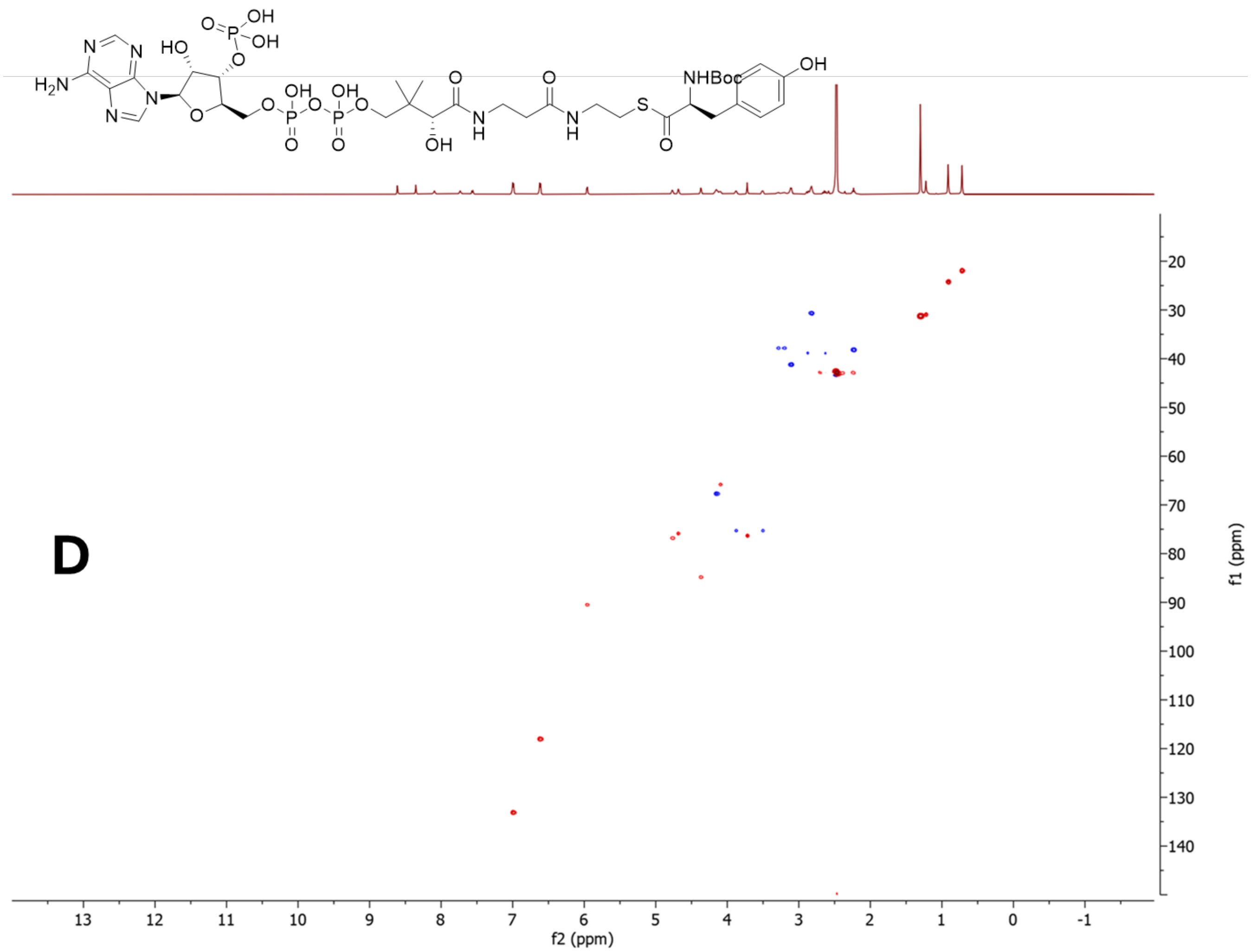


**Figure S54:** NMR data for **3Cb** in DMSO-d$_6$, 600 MHz. **A)** $^1$H-NMR spectrum. **B)** COSY spectrum. **C)** HMBC spectrum. **D)** HSQC spectrum.

5.3.3 NMR data for **3Cc**

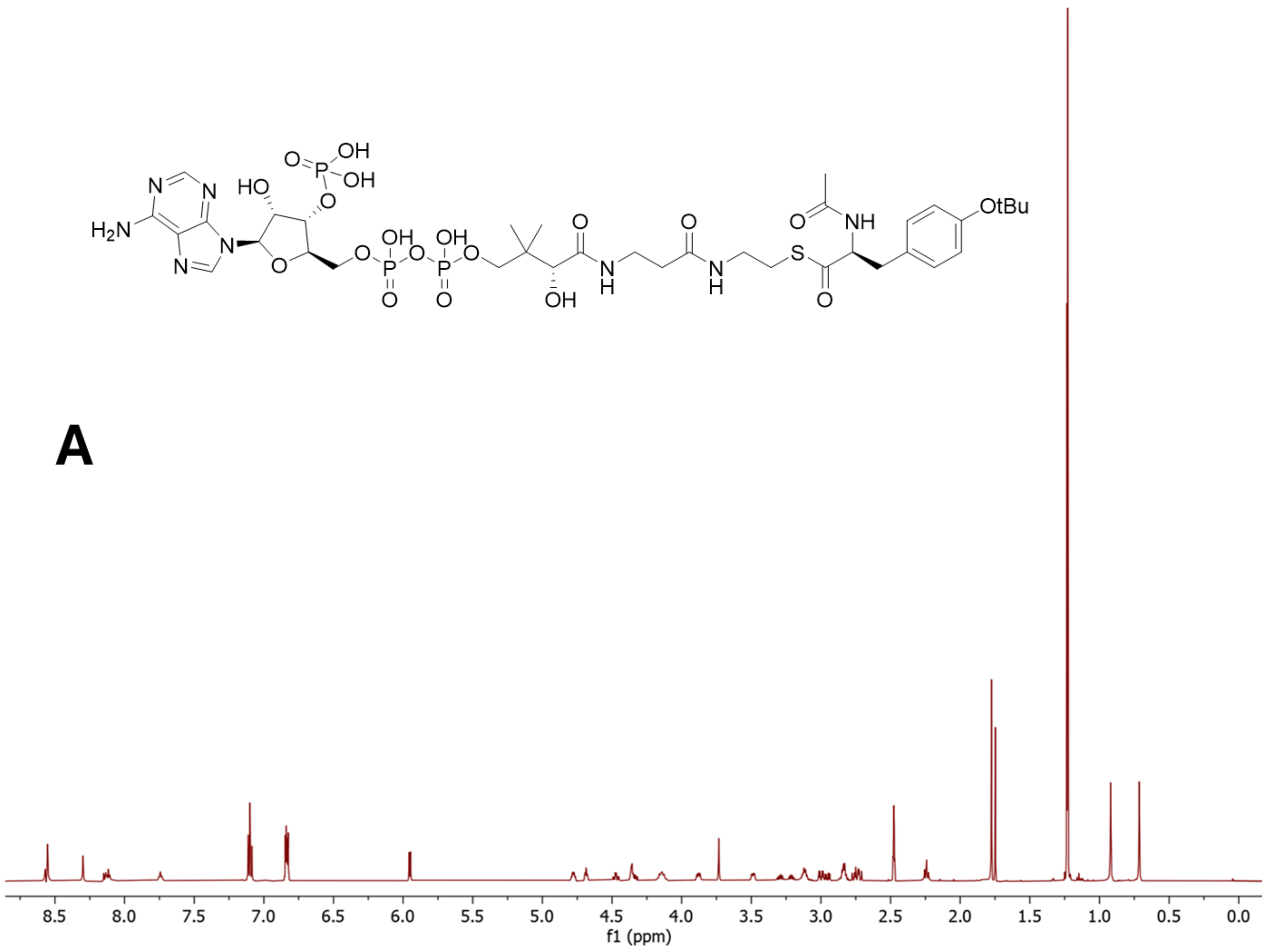

B

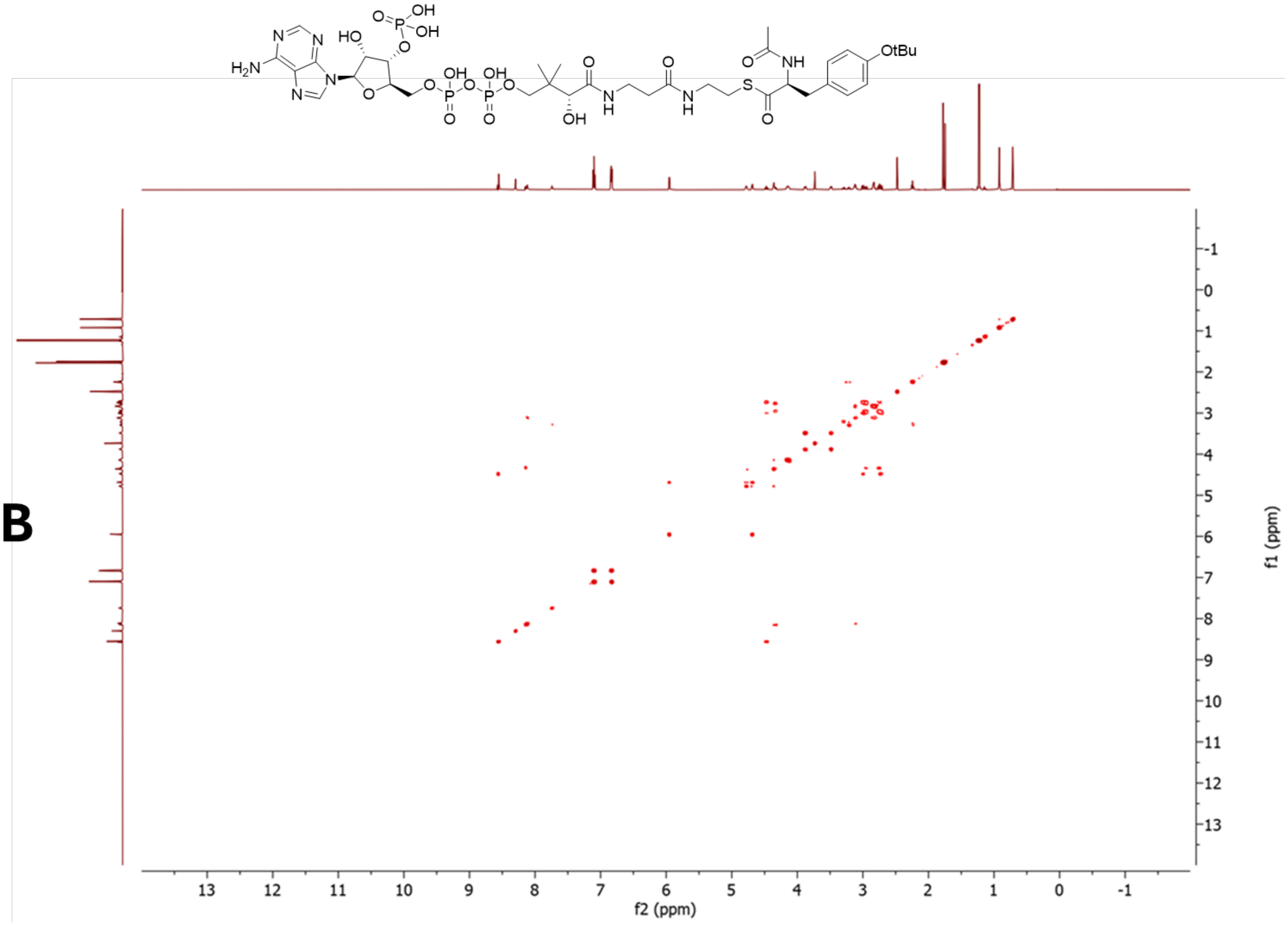

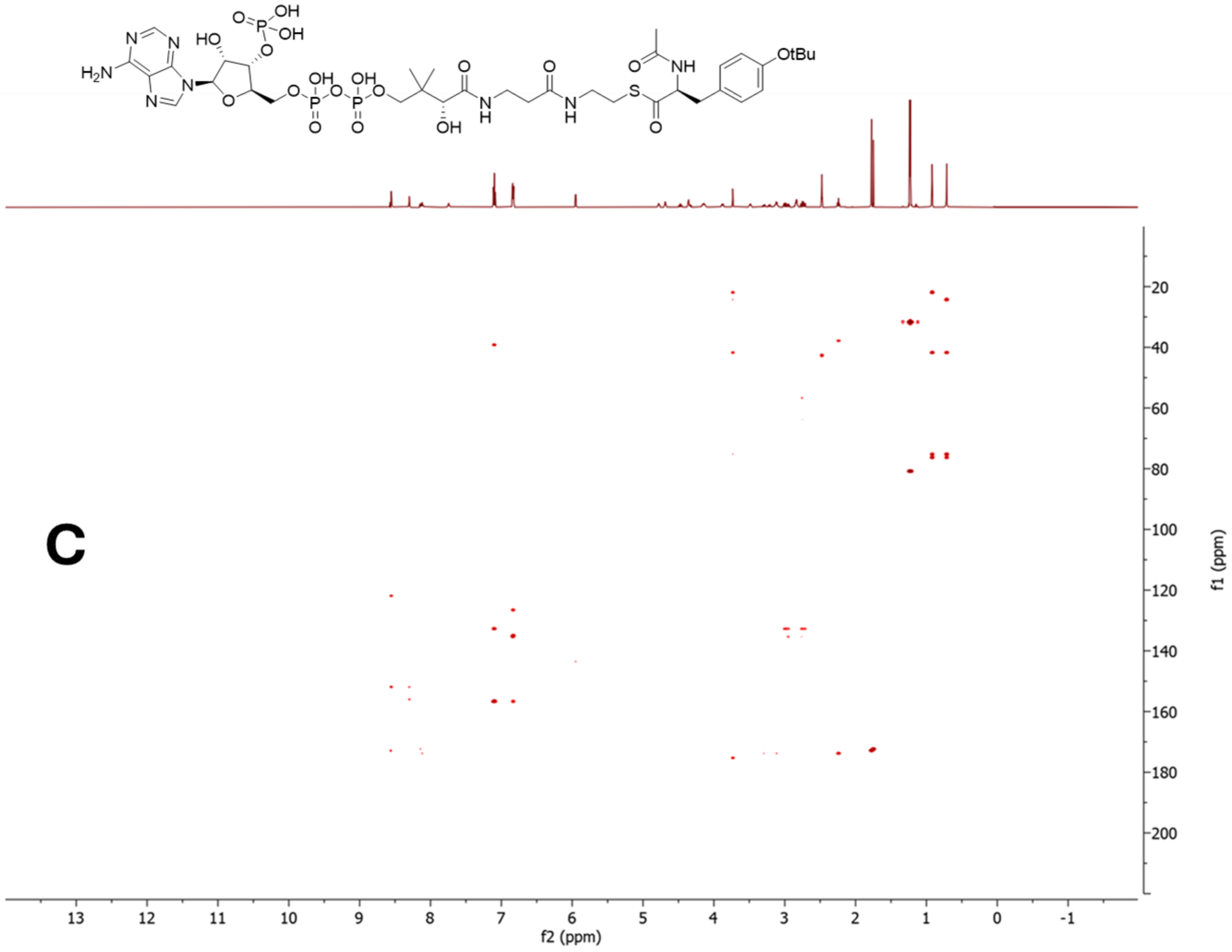
OH
O=P
OH
N
N
HO
O
H2N
N
N
O
OH
OH
O
O
O
NH
OtBu
O
P
O
P
O
N
H
N
H
S
O
O
O
OH
O
C
20
40
60
80
100
120
140
160
180
200
f1 (ppm)
13
12
11
10
9
8
7
6
5
4
3
2
1
0
-1
f2 (ppm)

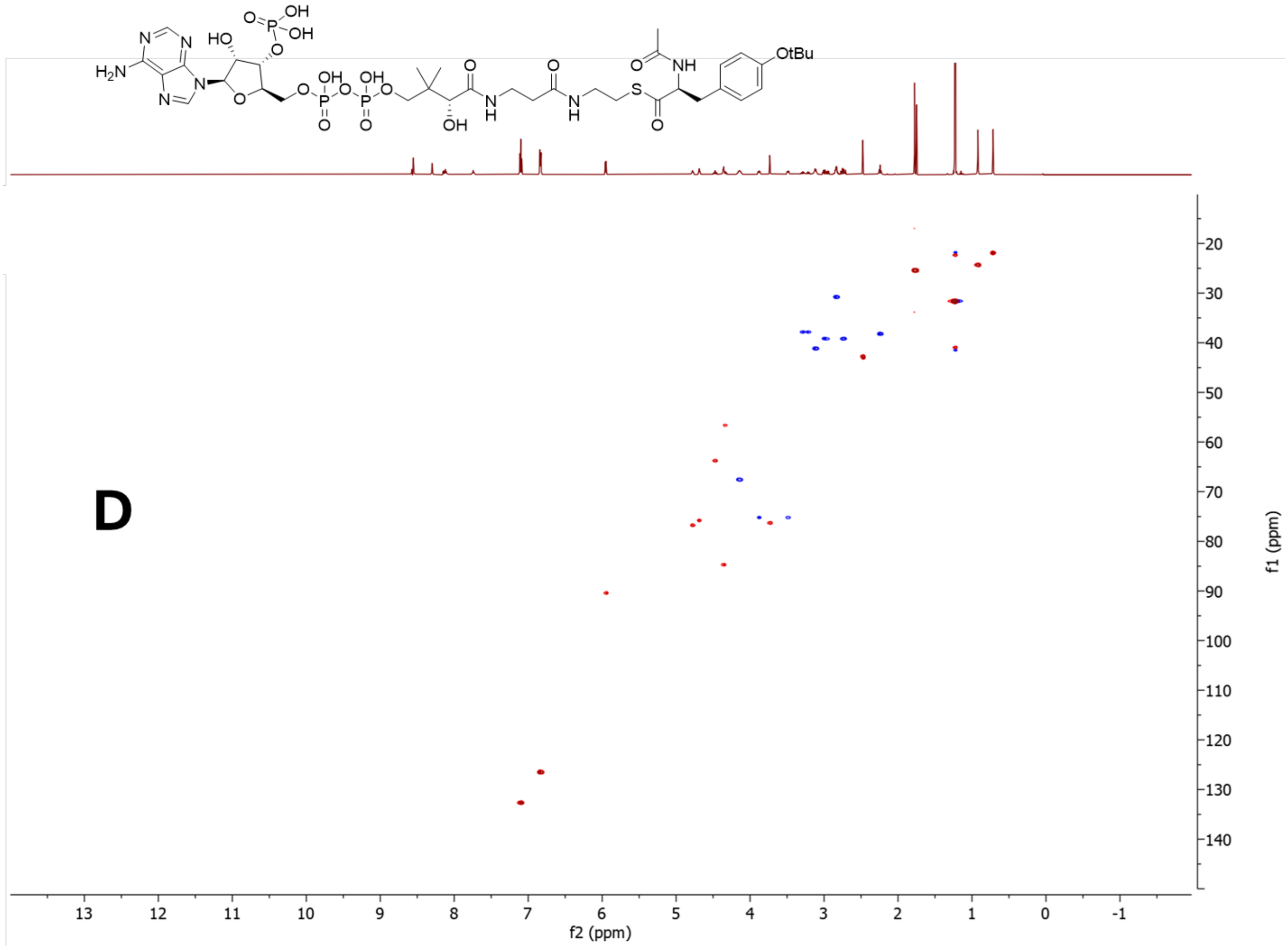


**Figure S55:** NMR data for **3Cc** in DMSO-d$_6$, 600 MHz. **A)** $^1$H-NMR spectrum. **B)** COSY spectrum. **C)** HMBC spectrum. **D)** HSQC spectrum.

5.4 NMR data for *N*-acyl-L-tyrosyl *N*-acetyl cysteamine thioesters

5.4.1 NMR data for **3Sa**

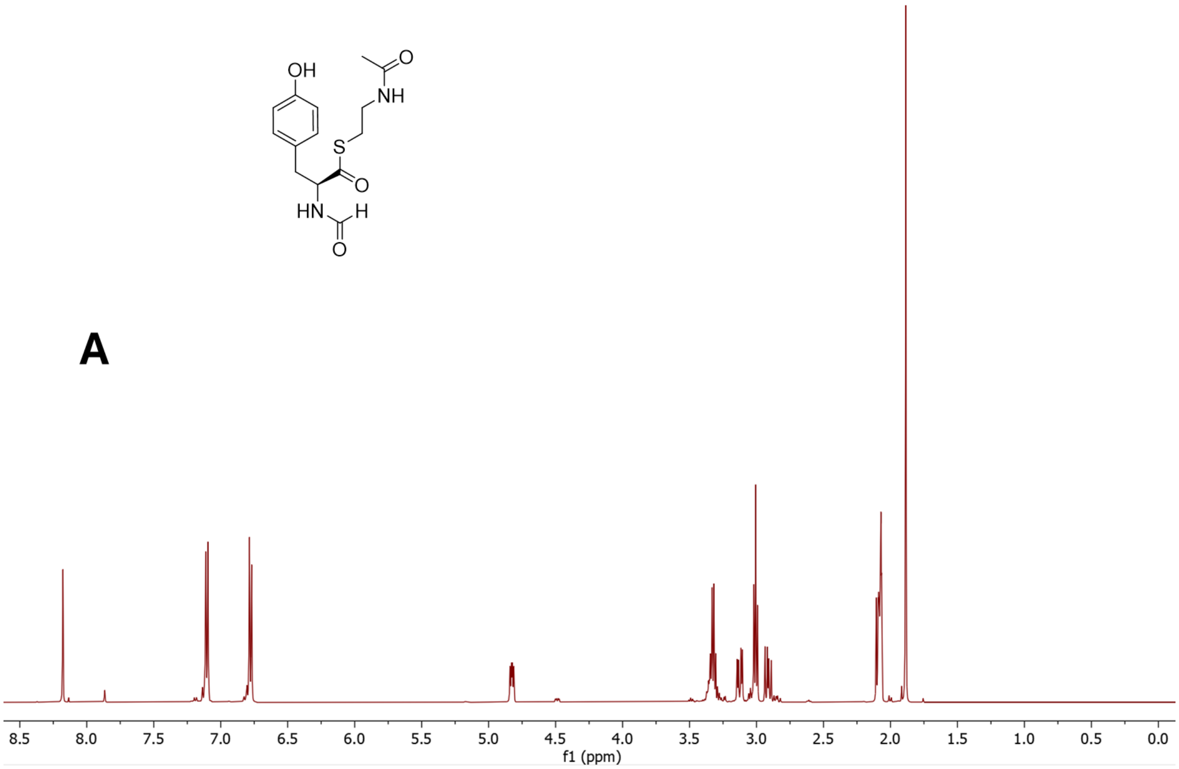


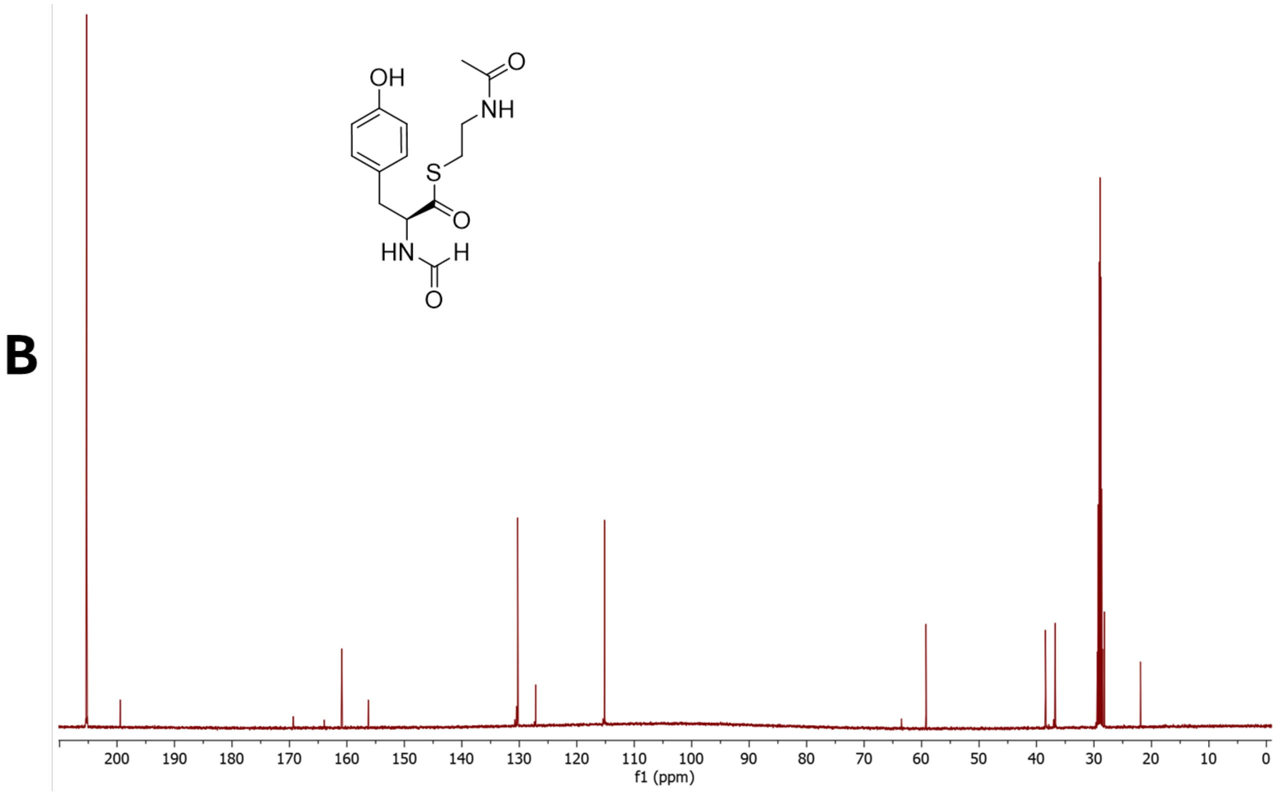

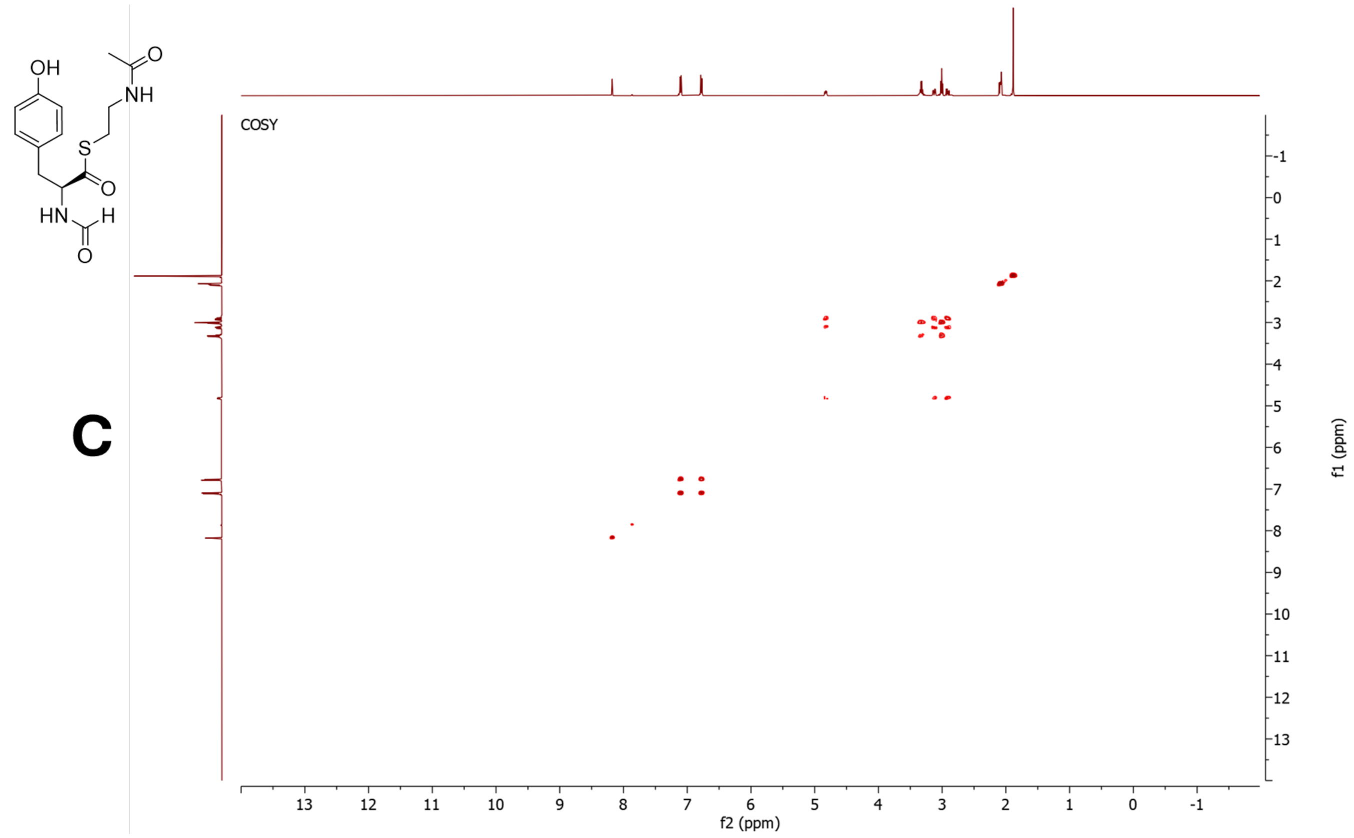
OH
NH
O
S
O
HN
H
O
COSY
f1 (ppm)
f2 (ppm)

C

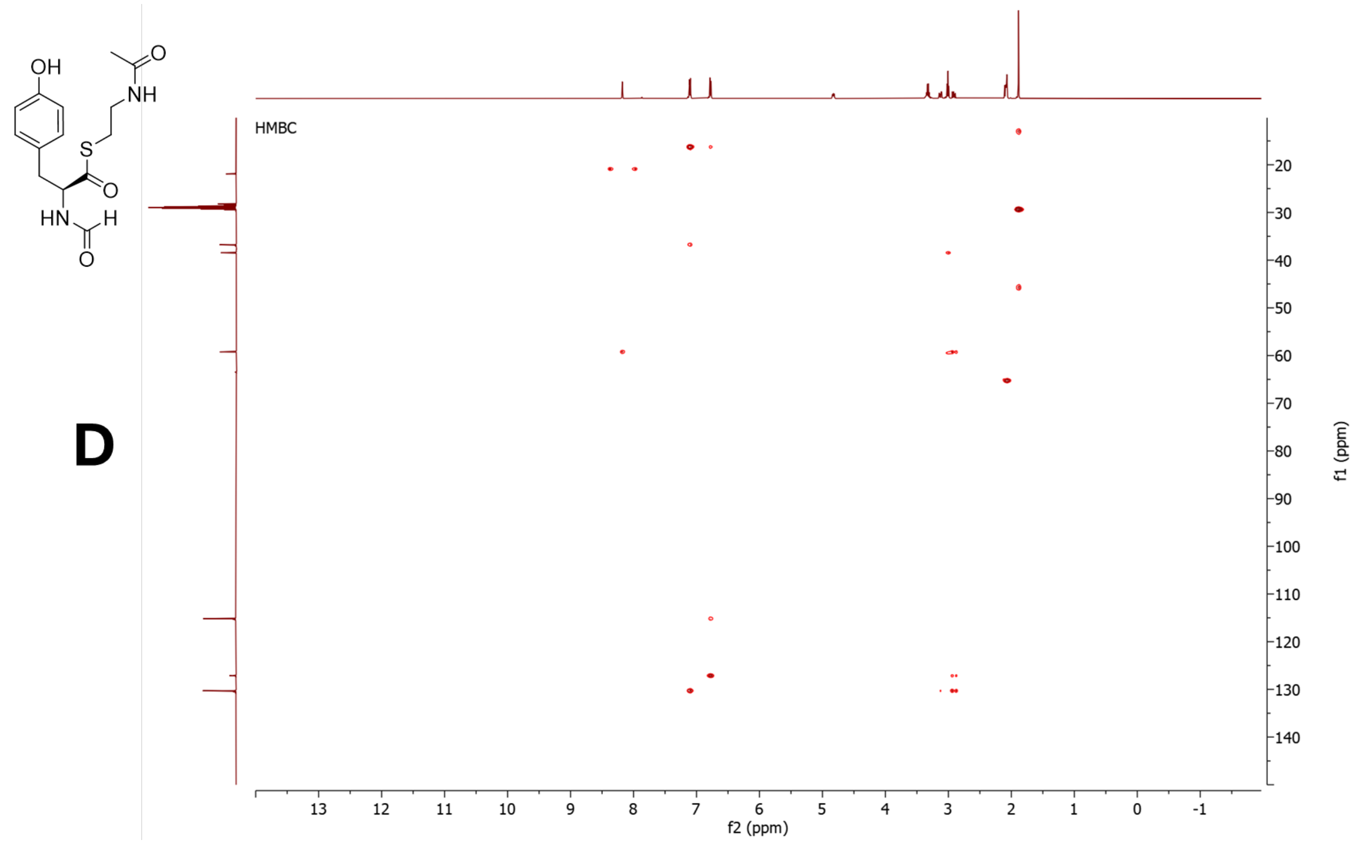
OH
NH
O
S
O
HN
H
O
HMBC
f1 (ppm)
f2 (ppm)

D

E

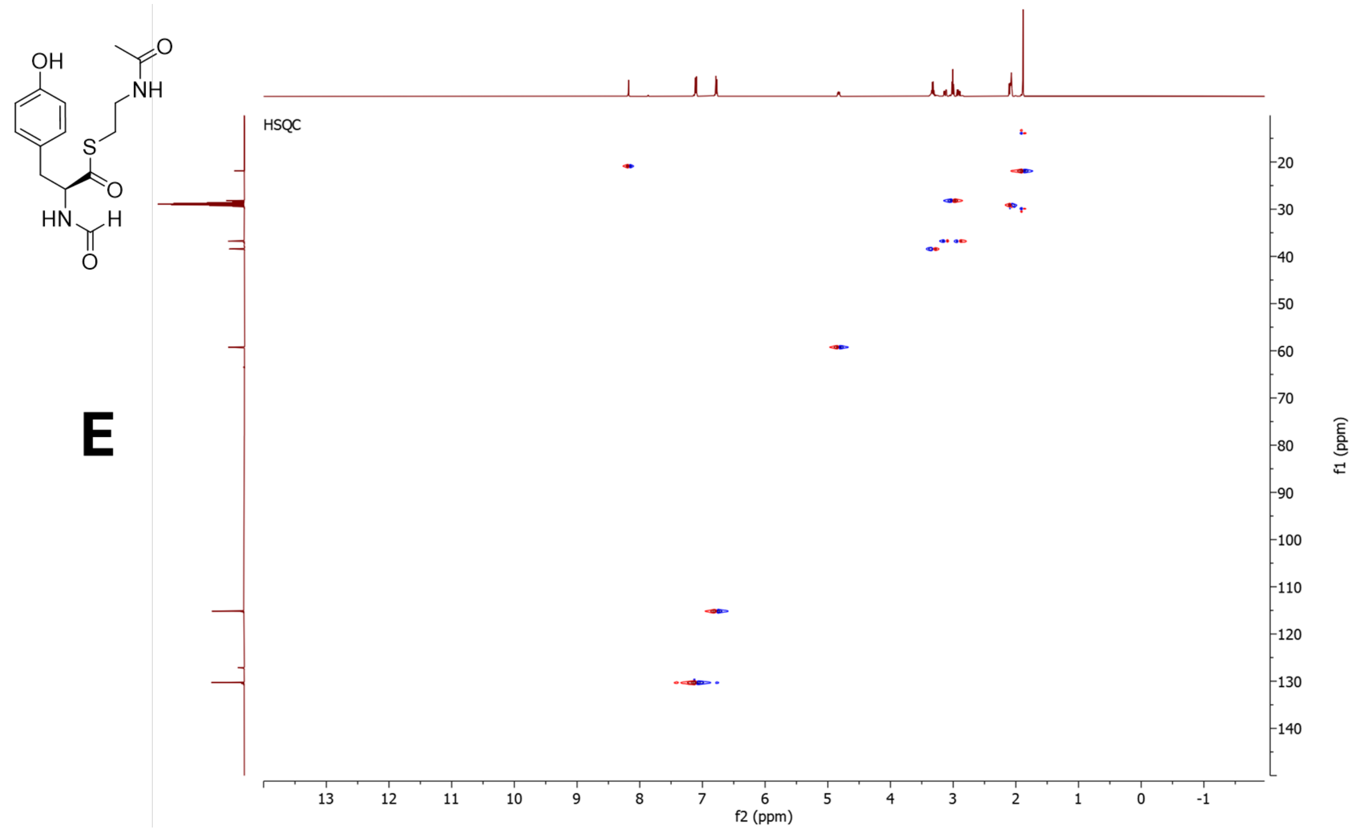


**Figure S56**. NMR data for **3Sa** in acetone-$d_6$, 500 MHz. **A)** $^{1}$H-NMR spectrum. **B)** $^{13}$C-NMR spectrum. **C)** COSY spectrum. **C)** HMBC spectrum. **E)** HSQC spectrum. The CHO carbon in the HSQC is aliased

5.4.2 NMR data for **3Sc**

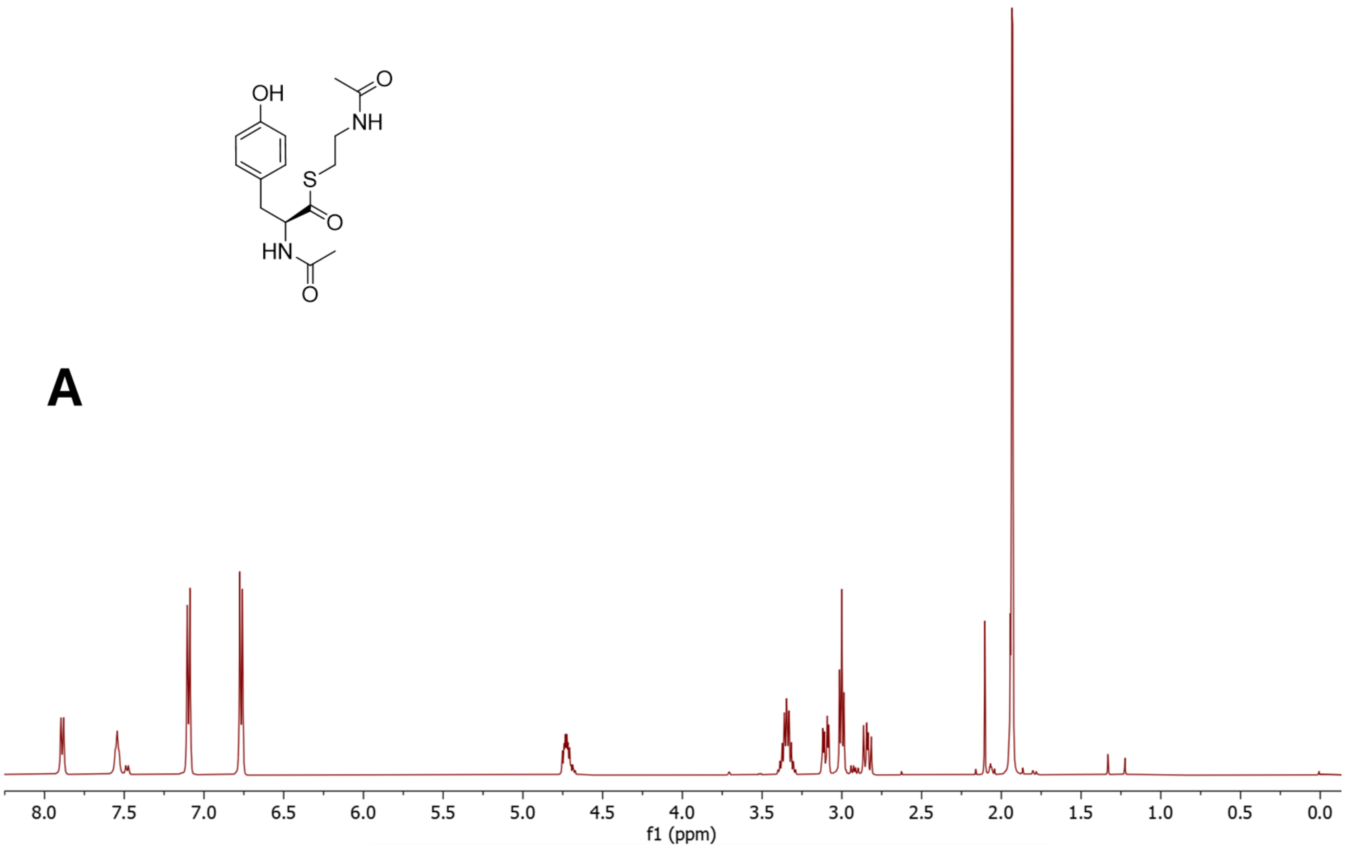


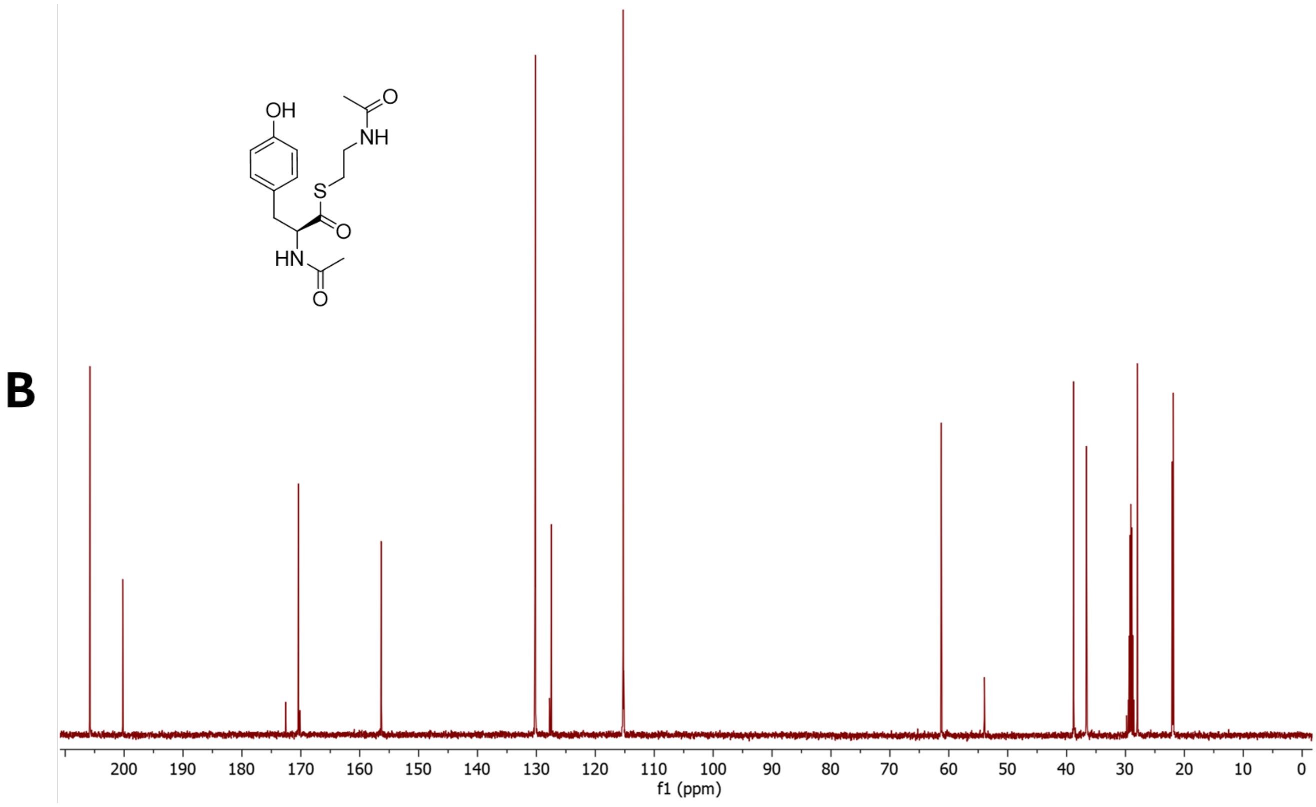

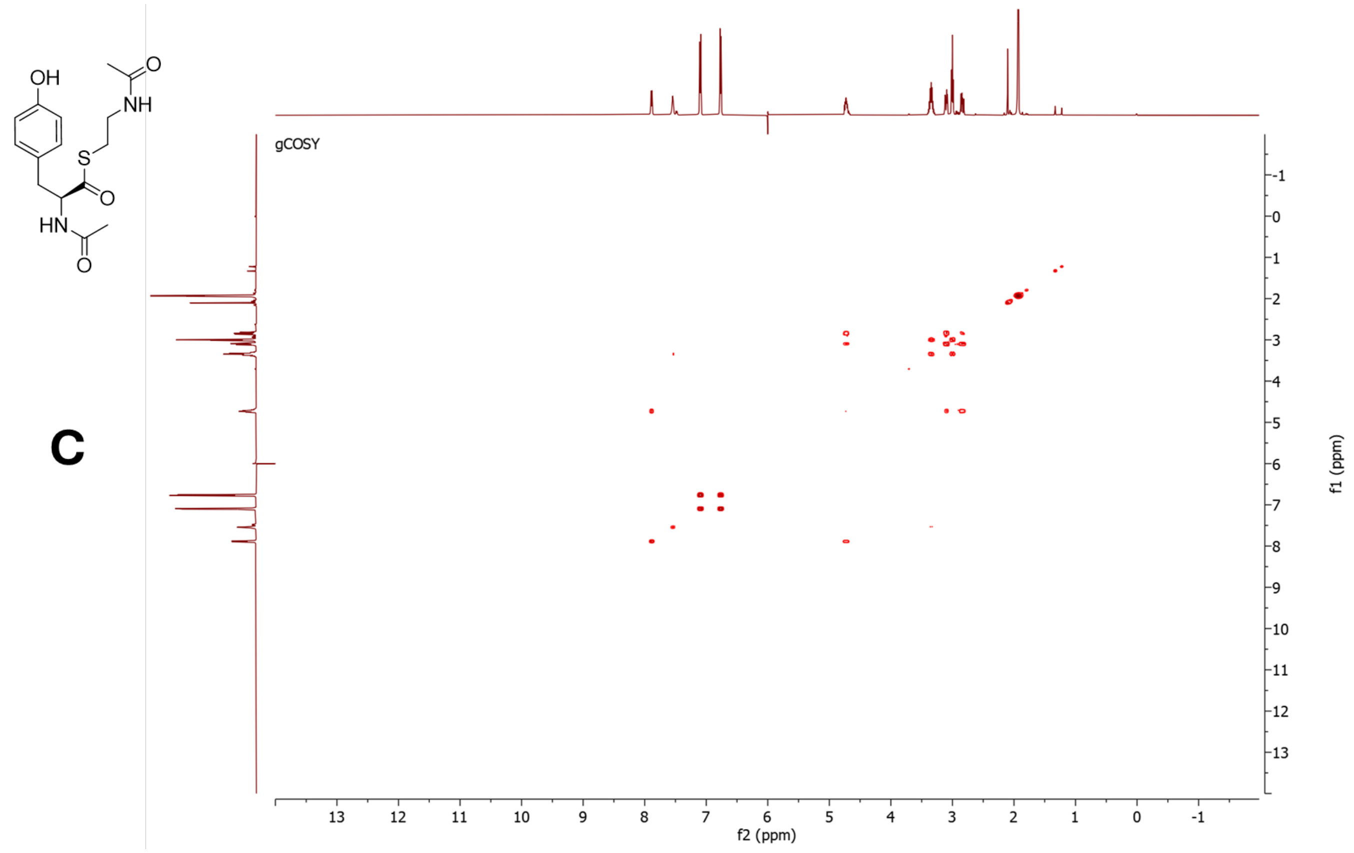


C

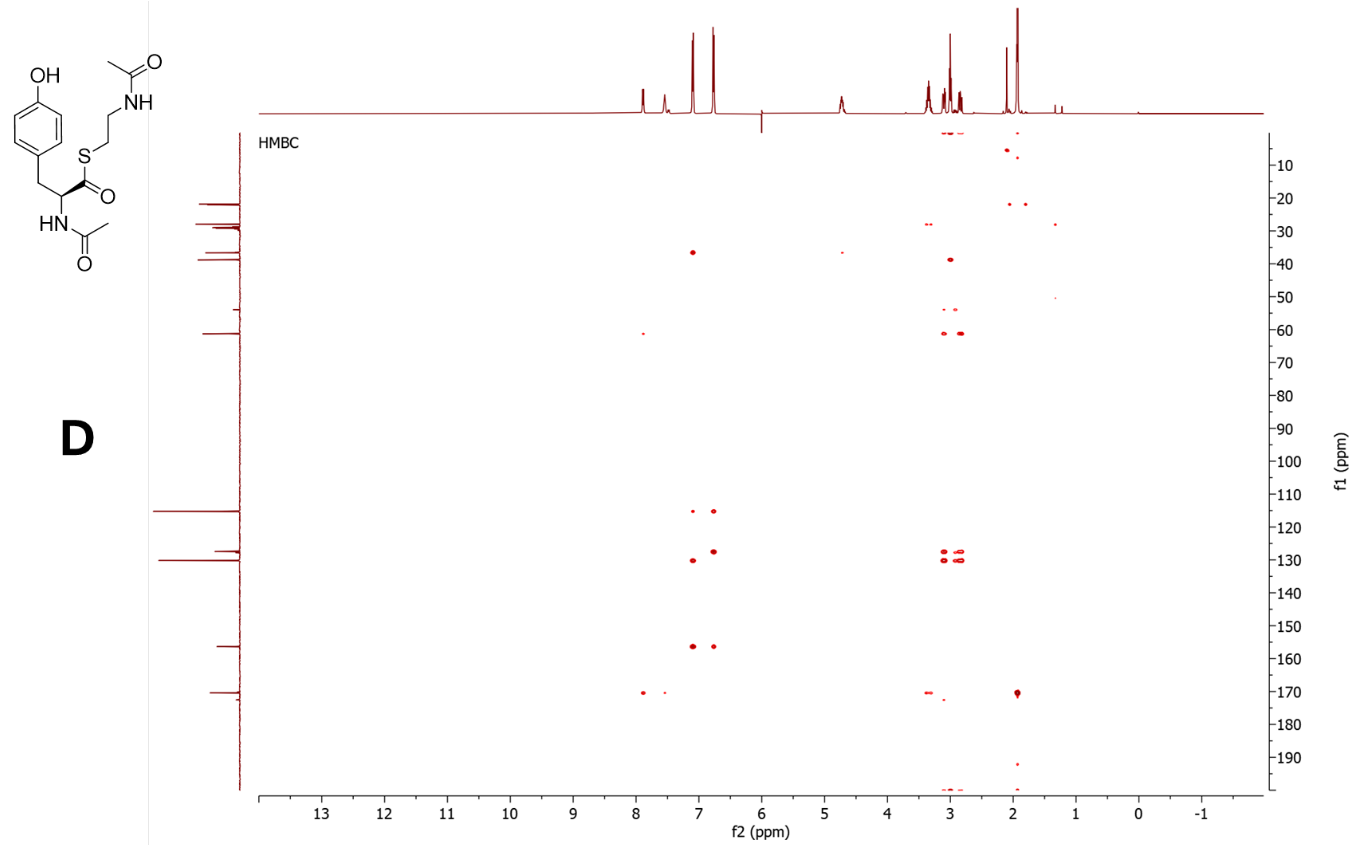


D

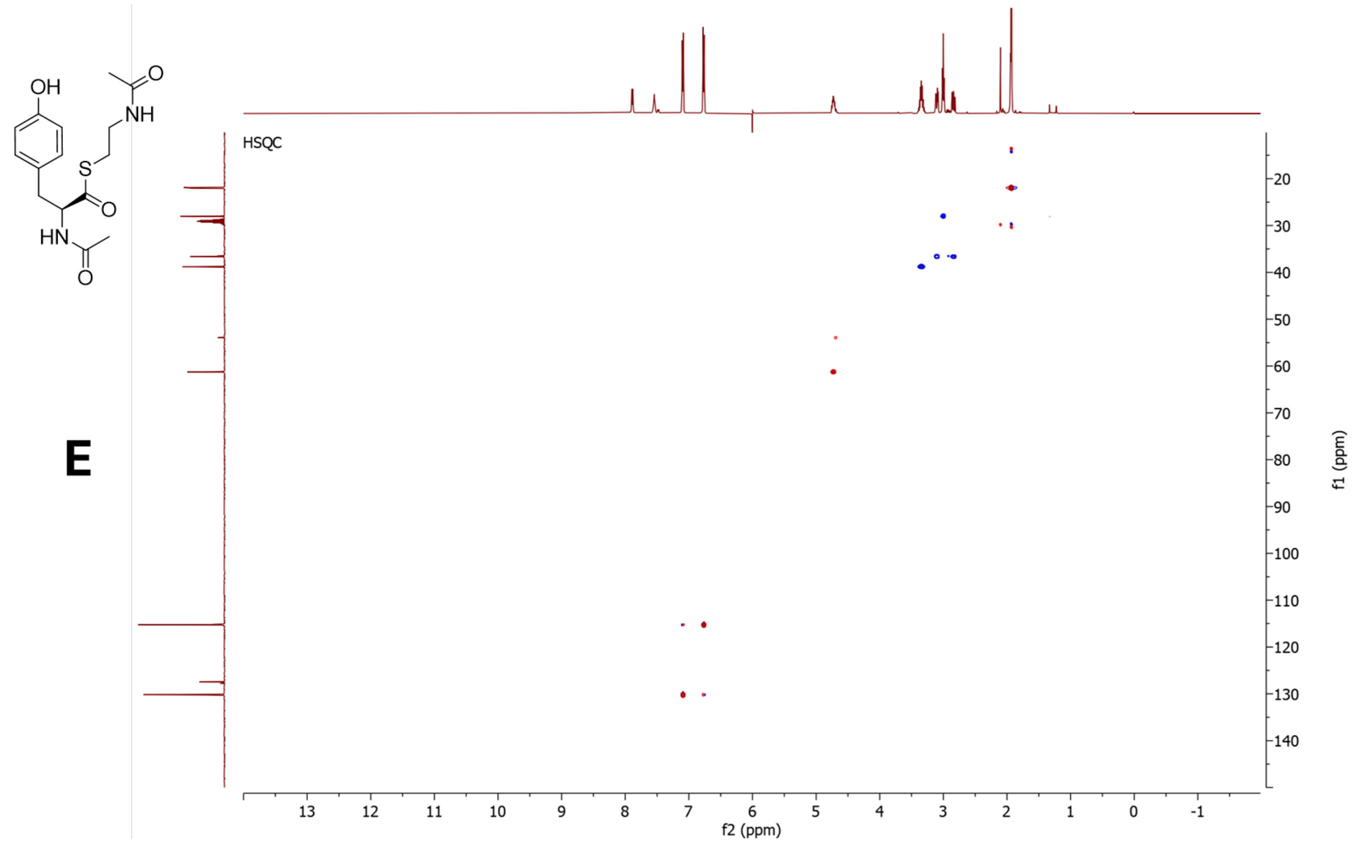


**Figure S57**. NMR data for **3Sc** in acetone-*d*6, 500 MHz. **A)** [1]H-NMR spectrum. **B)** [13]C-NMR spectrum. **C)** COSY spectrum. **C)** HMBC spectrum. **E)** HSQC spectrum.

5.4.3 NMR data for **3Sd**

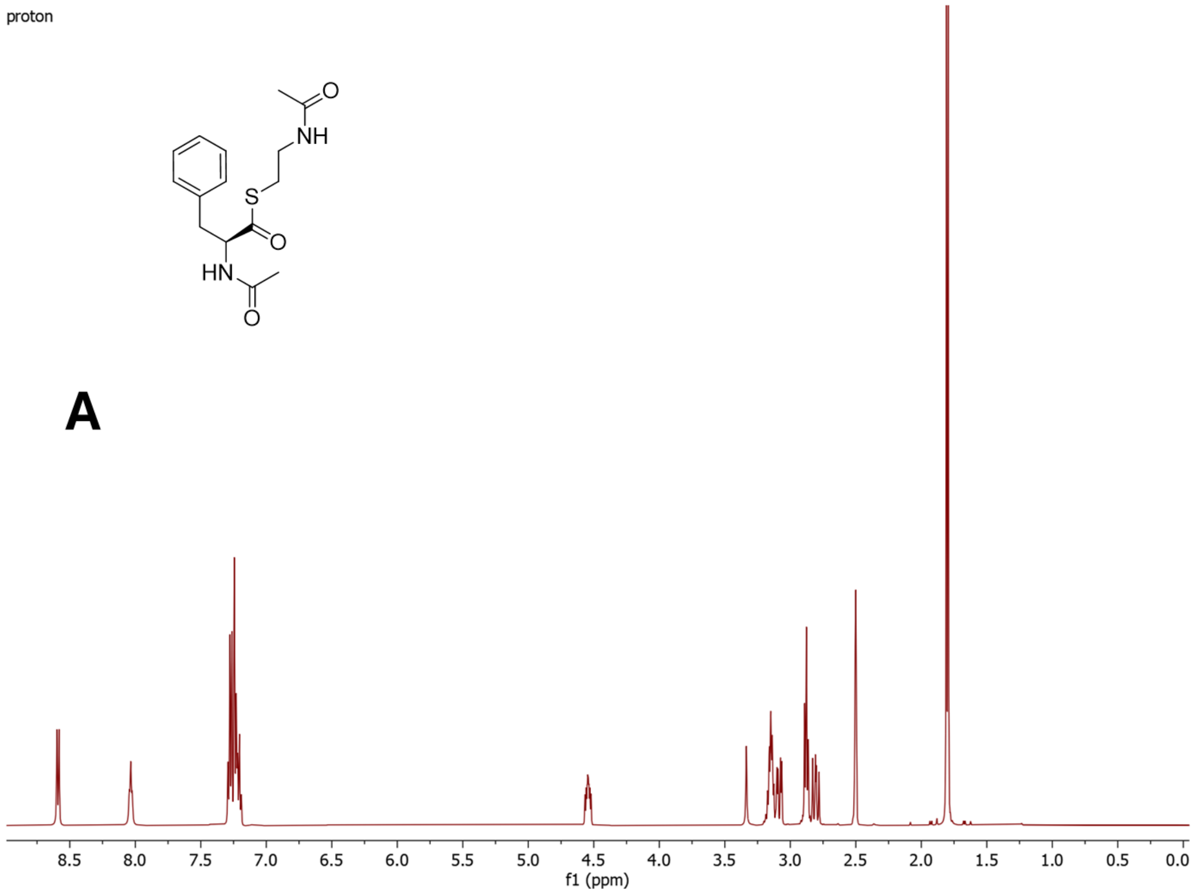


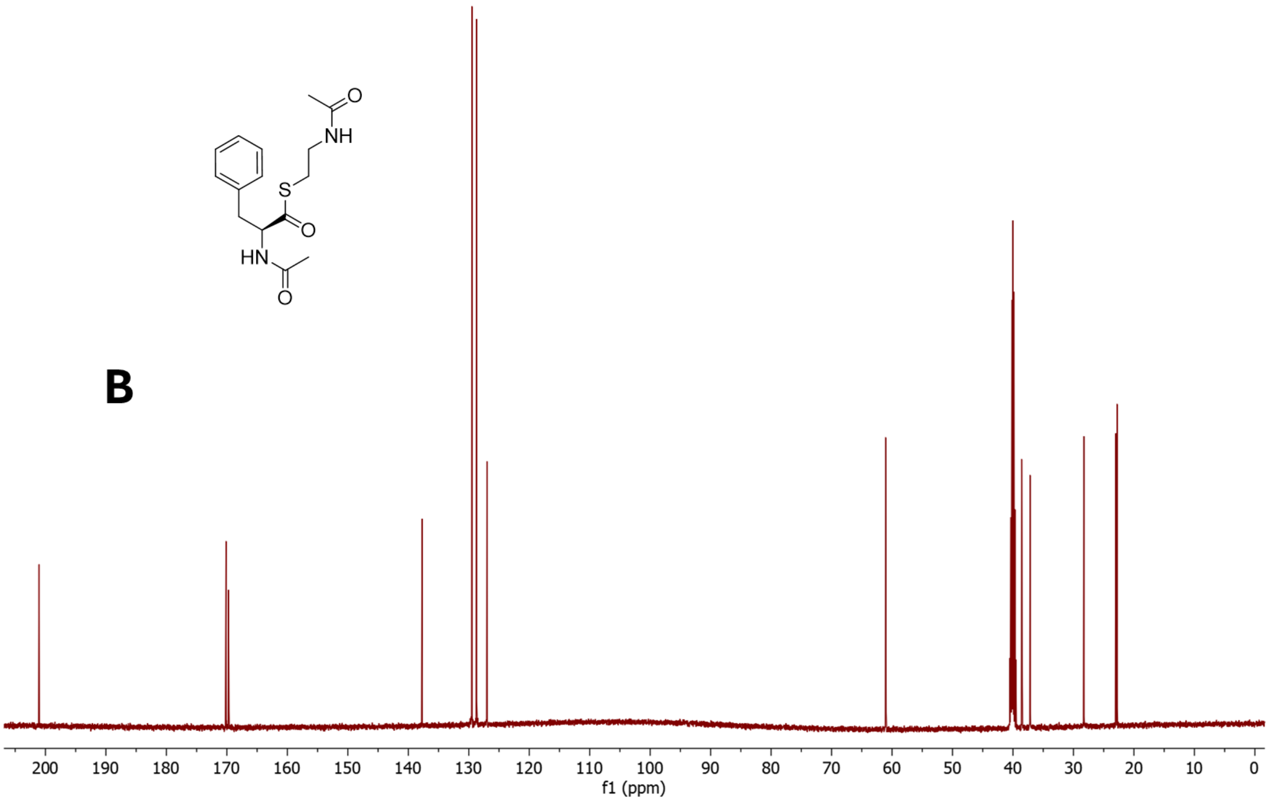

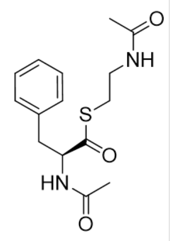


**C**

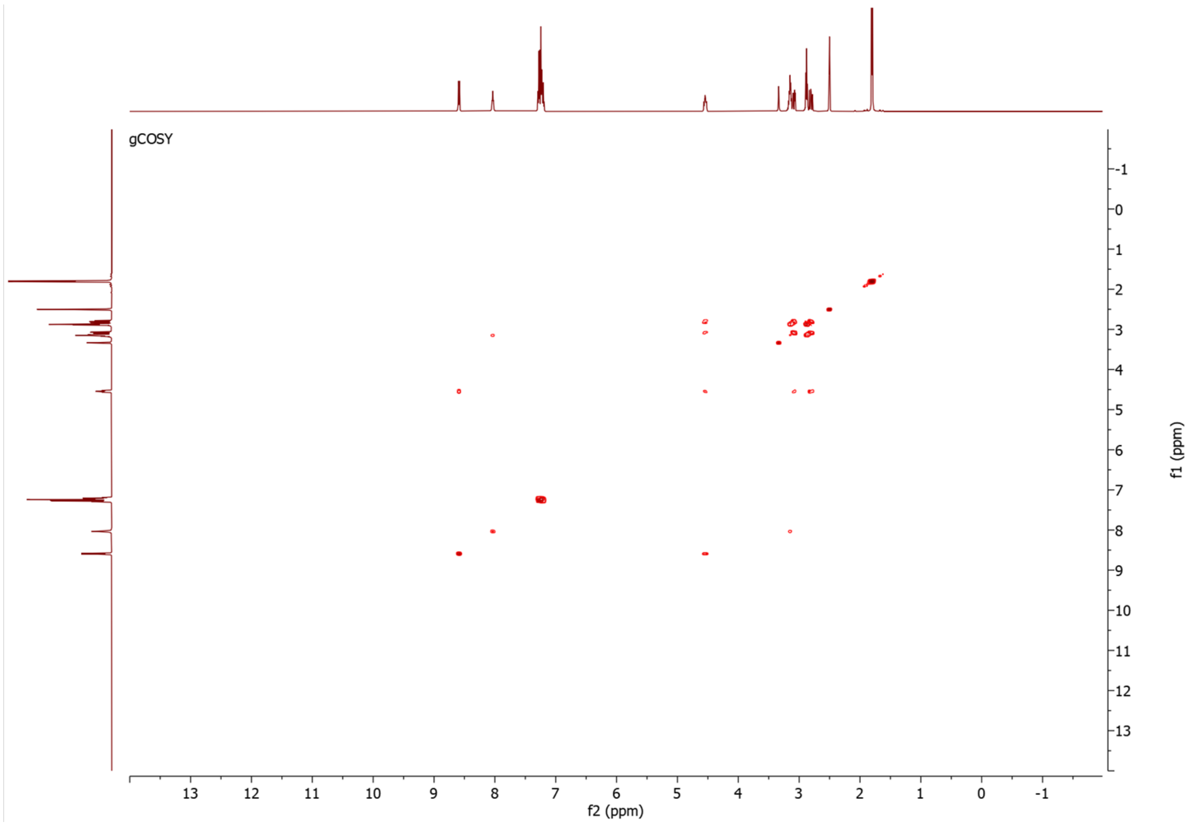


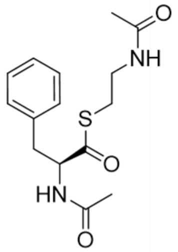


**D**

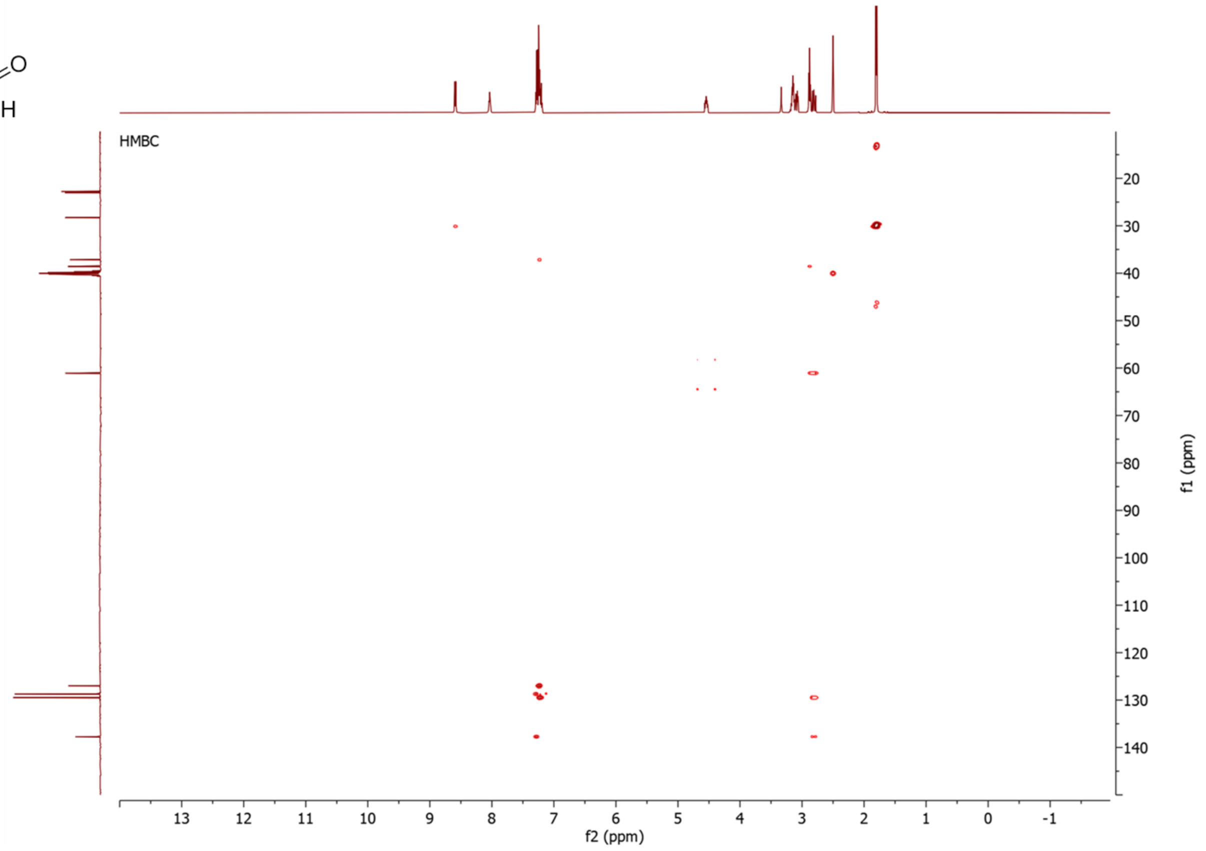

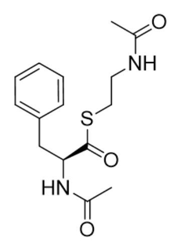

E

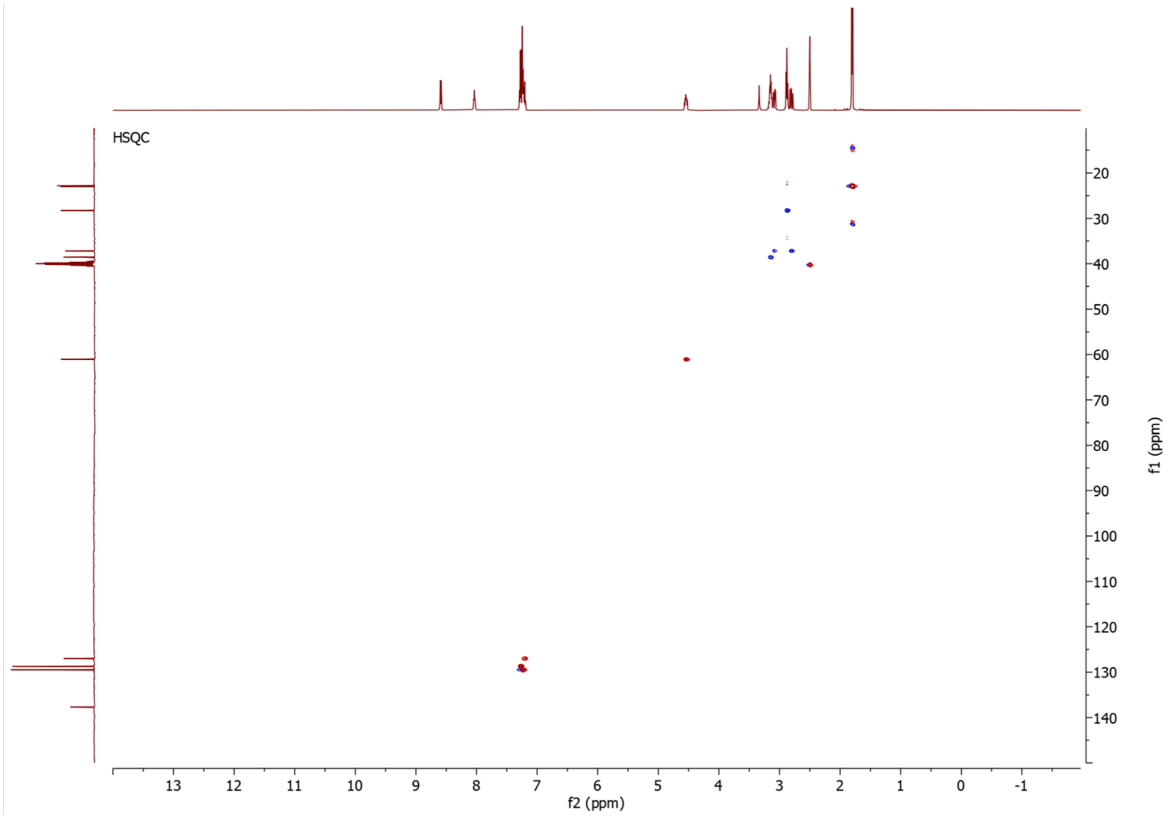


**Figure S58**. NMR data for **3Sd** in DMSO-$d_6$, 500 MHz. **A)** $^{1}$H-NMR spectrum. **B)** $^{13}$C-NMR spectrum. **C)** COSY spectrum. **C)** HMBC spectrum. **E)** HSQC spectrum.

5.4.4 NMR data for **3Sb**

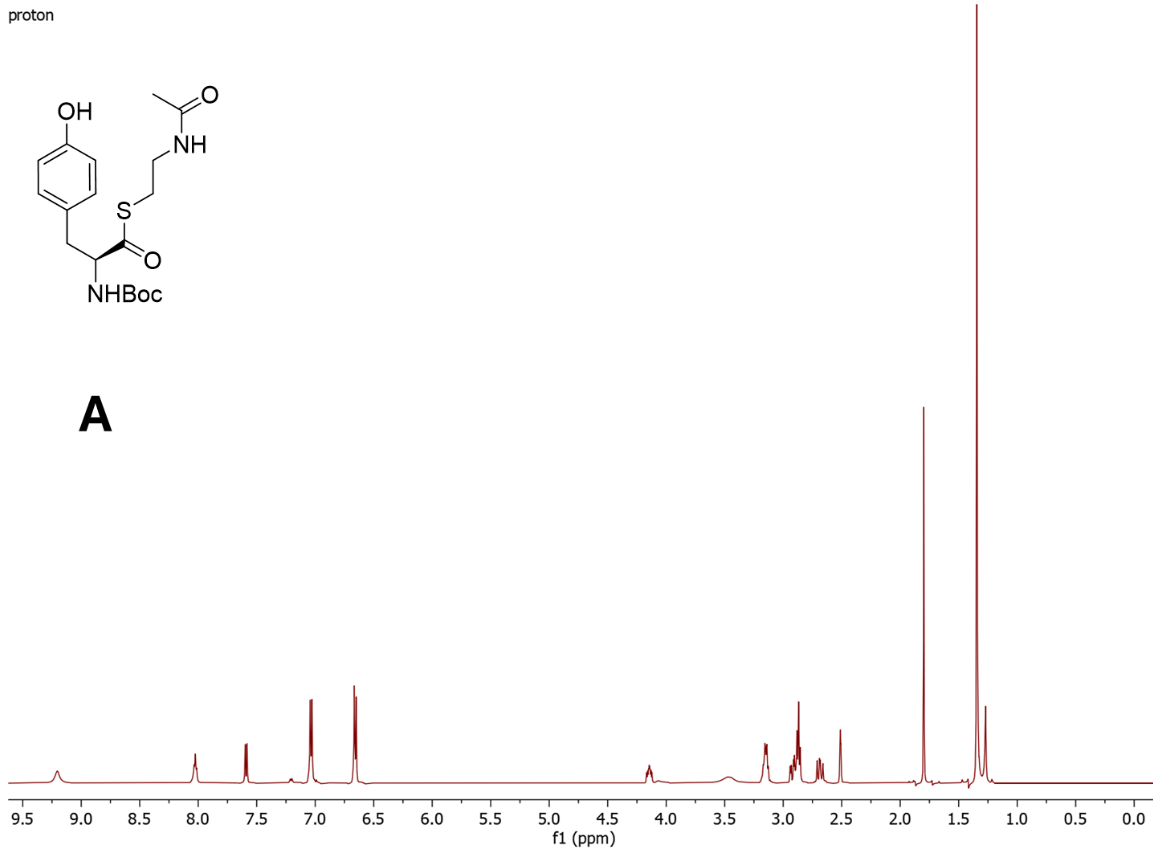


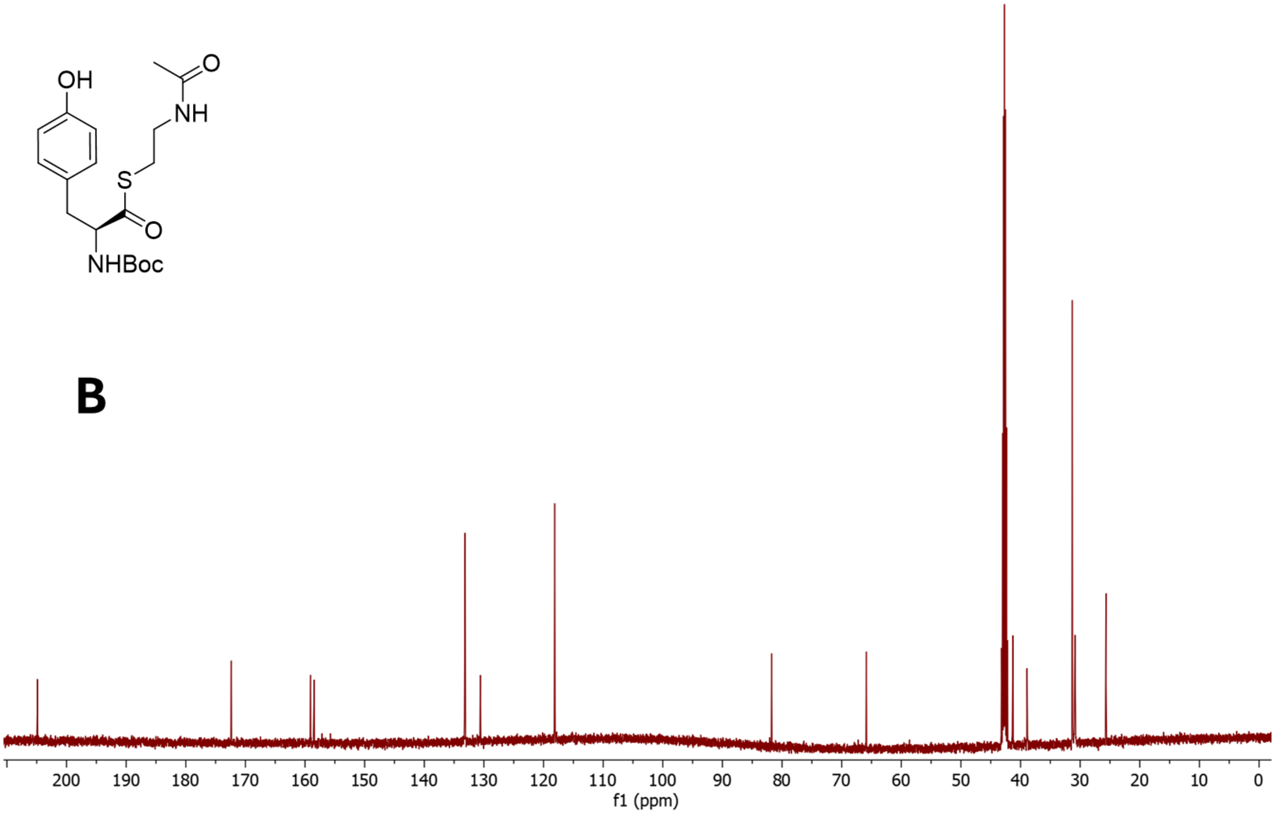

C

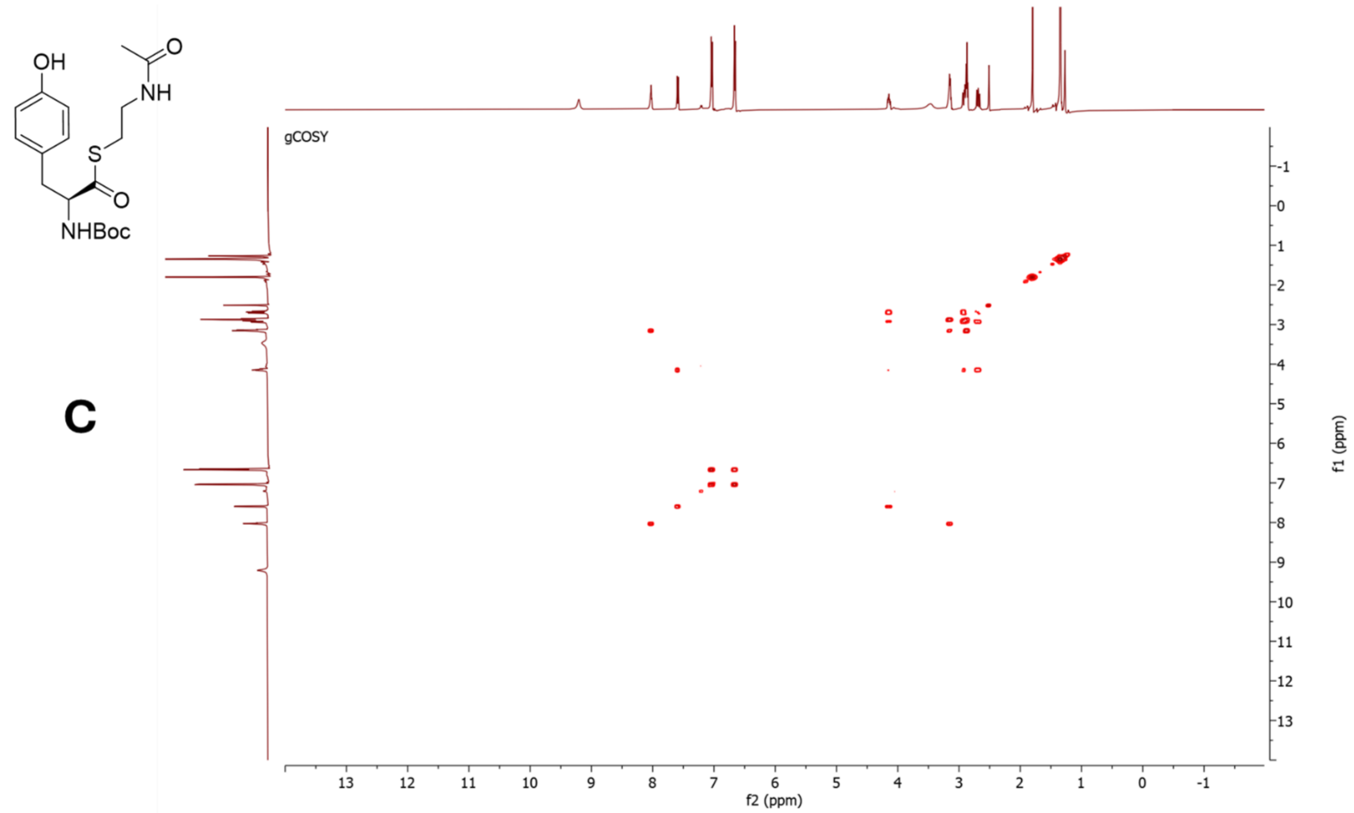
OH
O
NH
S
O
NHBoc
gCOSY
f2 (ppm)
f1 (ppm)

D

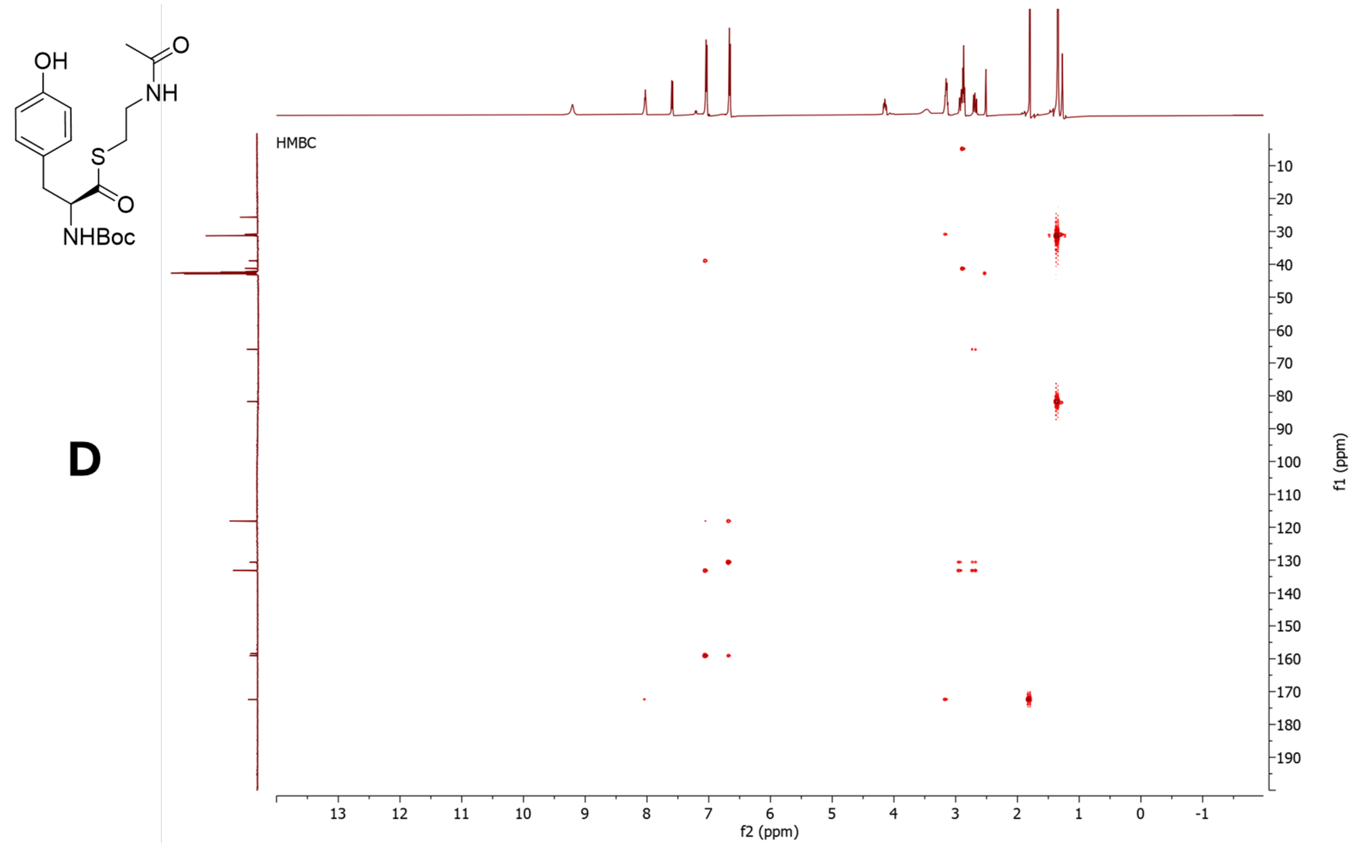
OH
O
NH
S
O
NHBoc
HMBC
f2 (ppm)
f1 (ppm)

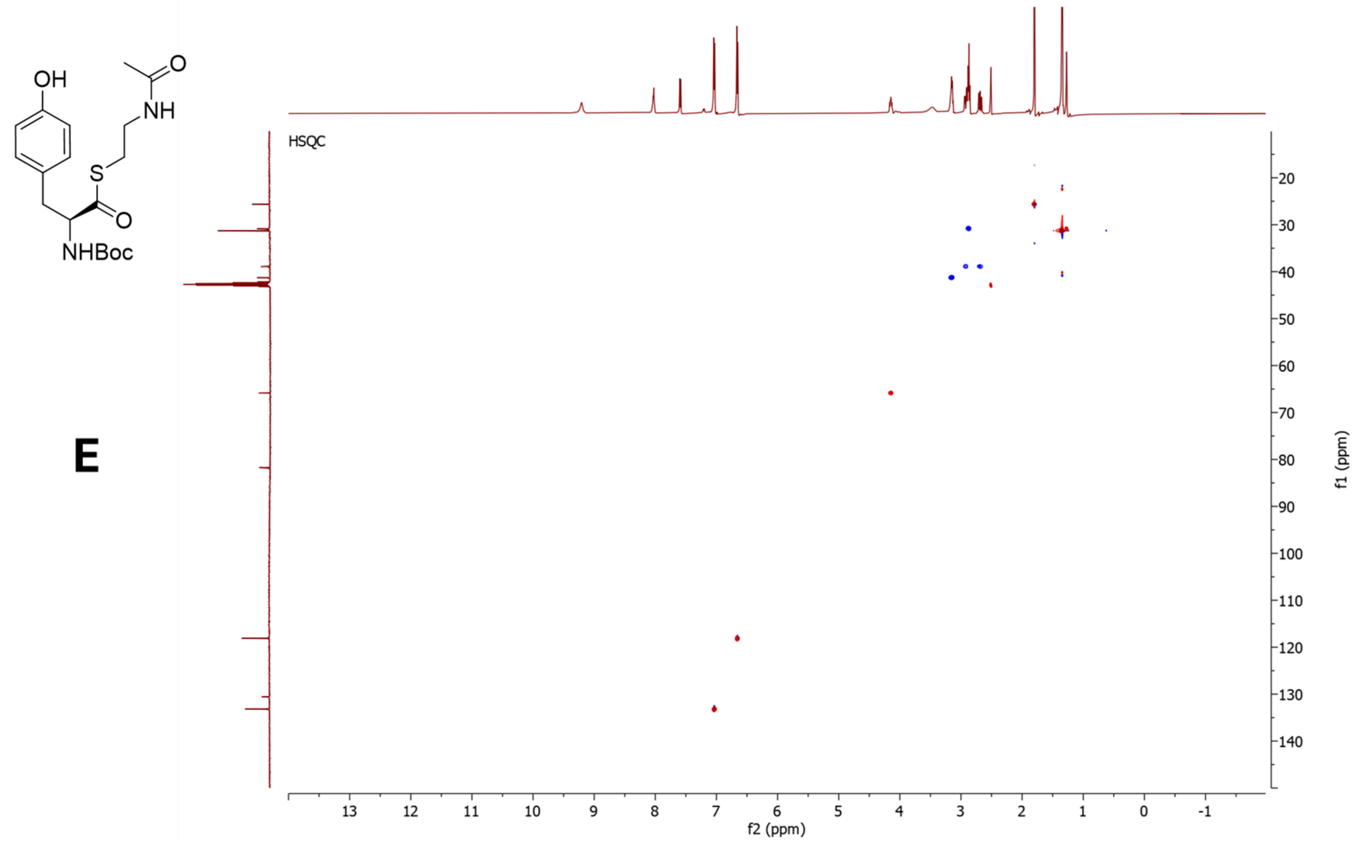


**Figure S59**. NMR data for **3Sb** in DMSO-$d_6$, 500 MHz. **A)** $^{1}$H-NMR spectrum. **B)** $^{13}$C-NMR spectrum. **C)** COSY spectrum. **C)** HMBC spectrum. **E)** HSQC spectrum.

5.5 NMR data for *N*-acyl-L-tyrosyl thiophenolates

5.5.1 NMR data for **7a**

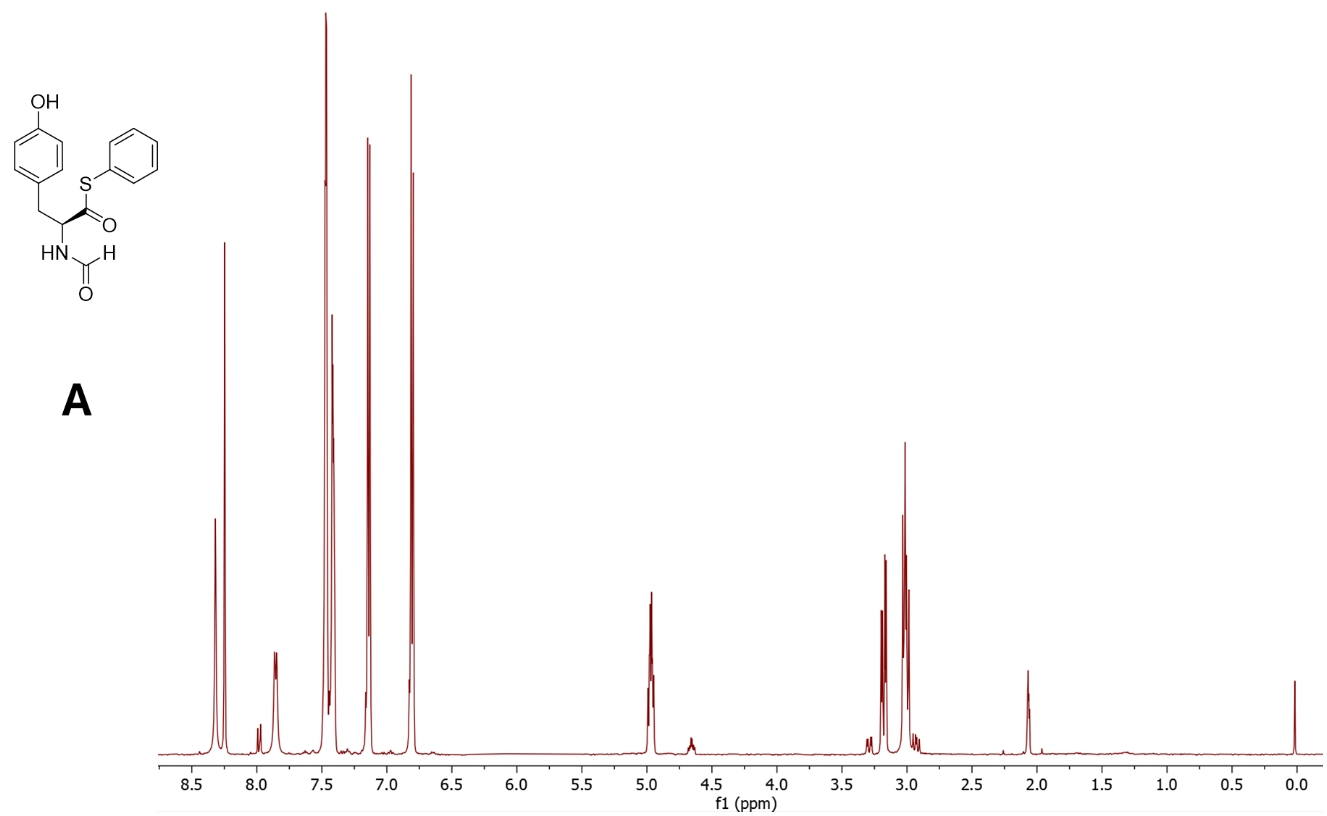


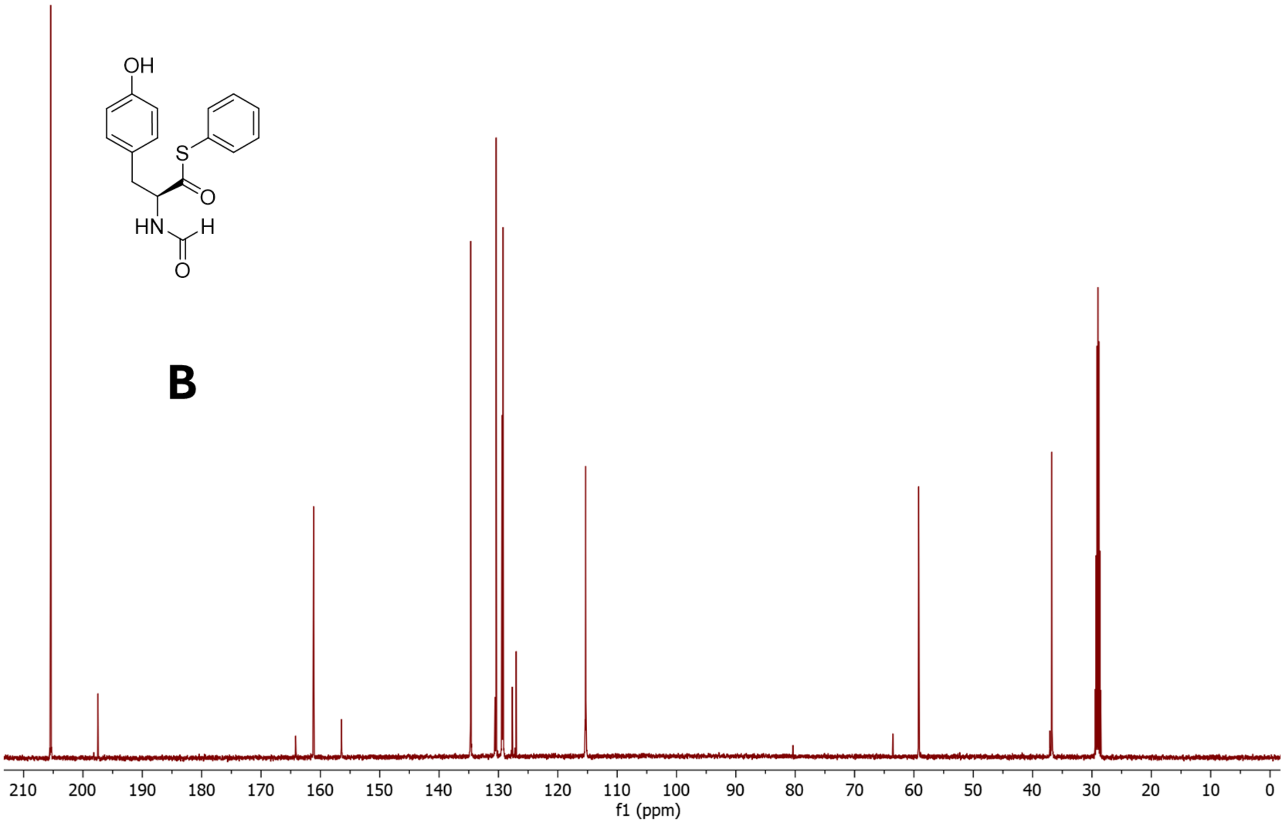

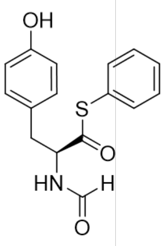

**C**

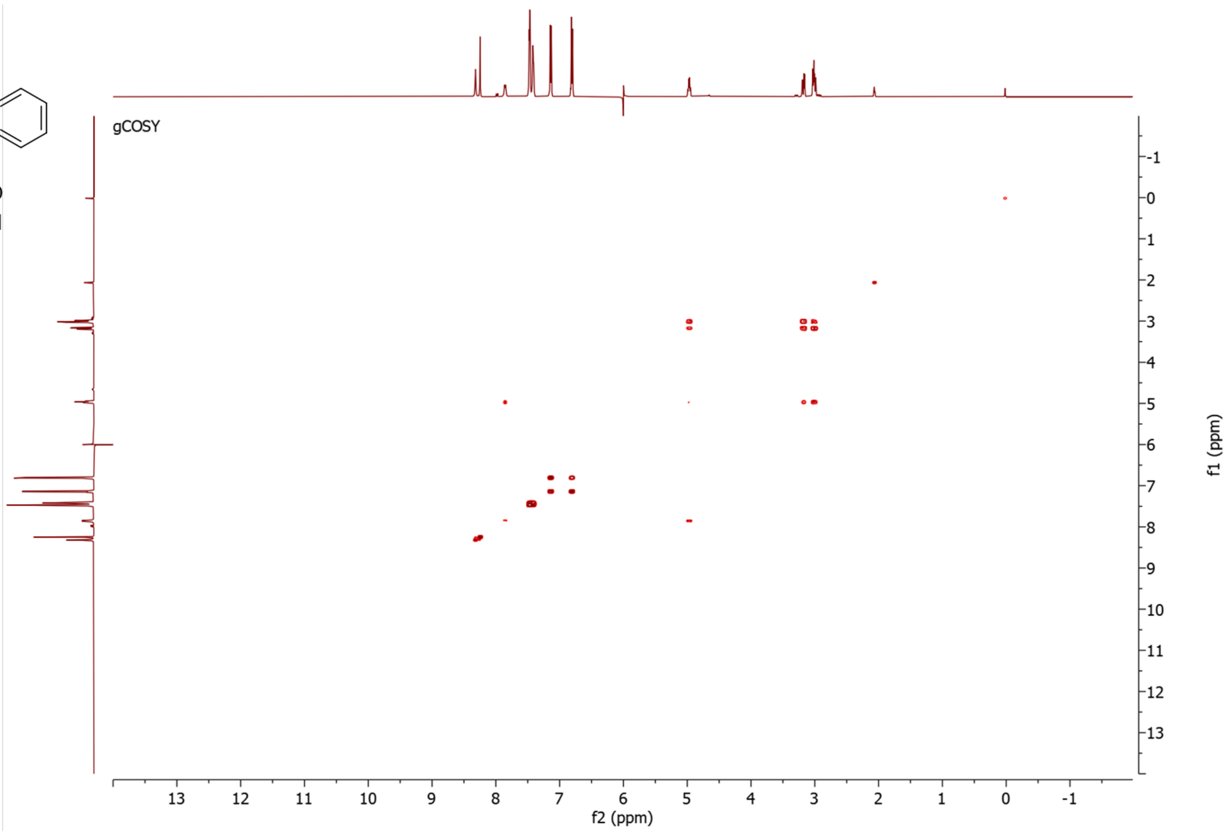


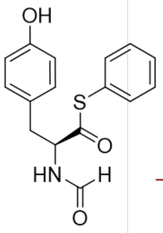

**D**

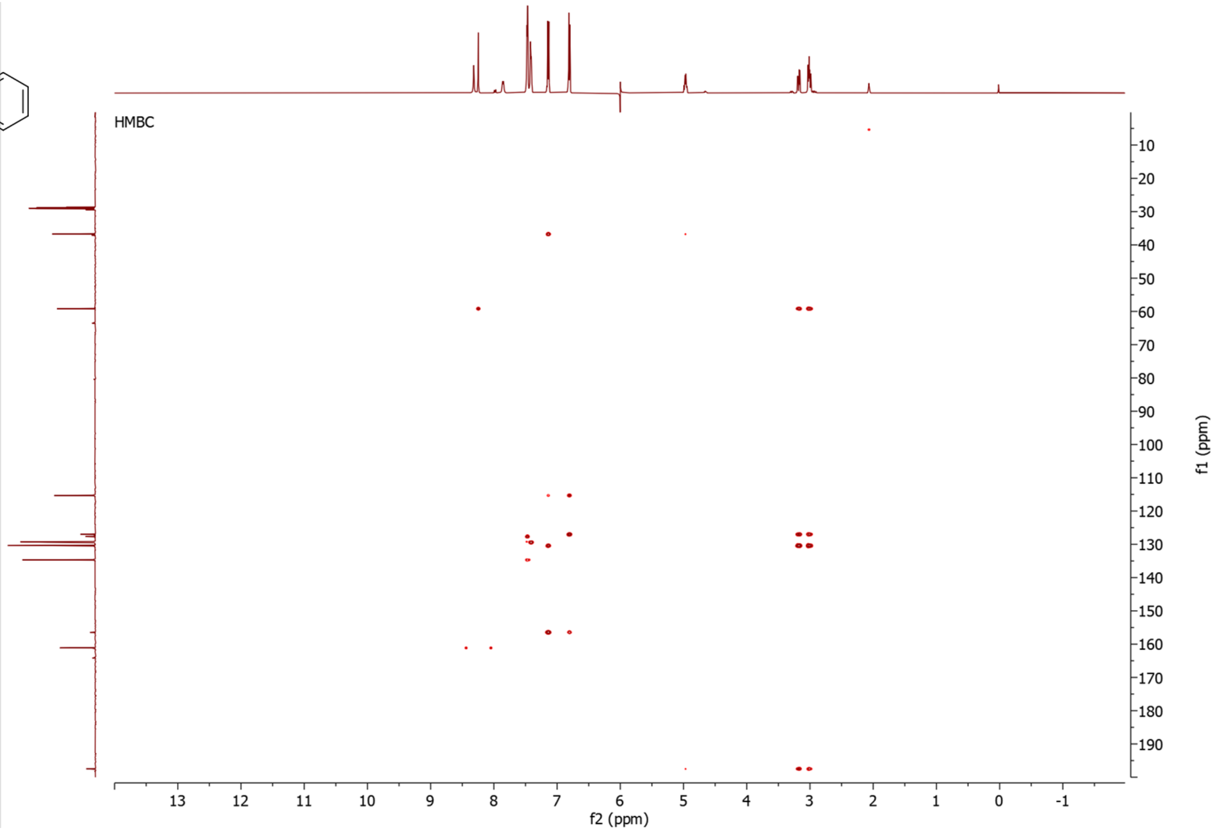

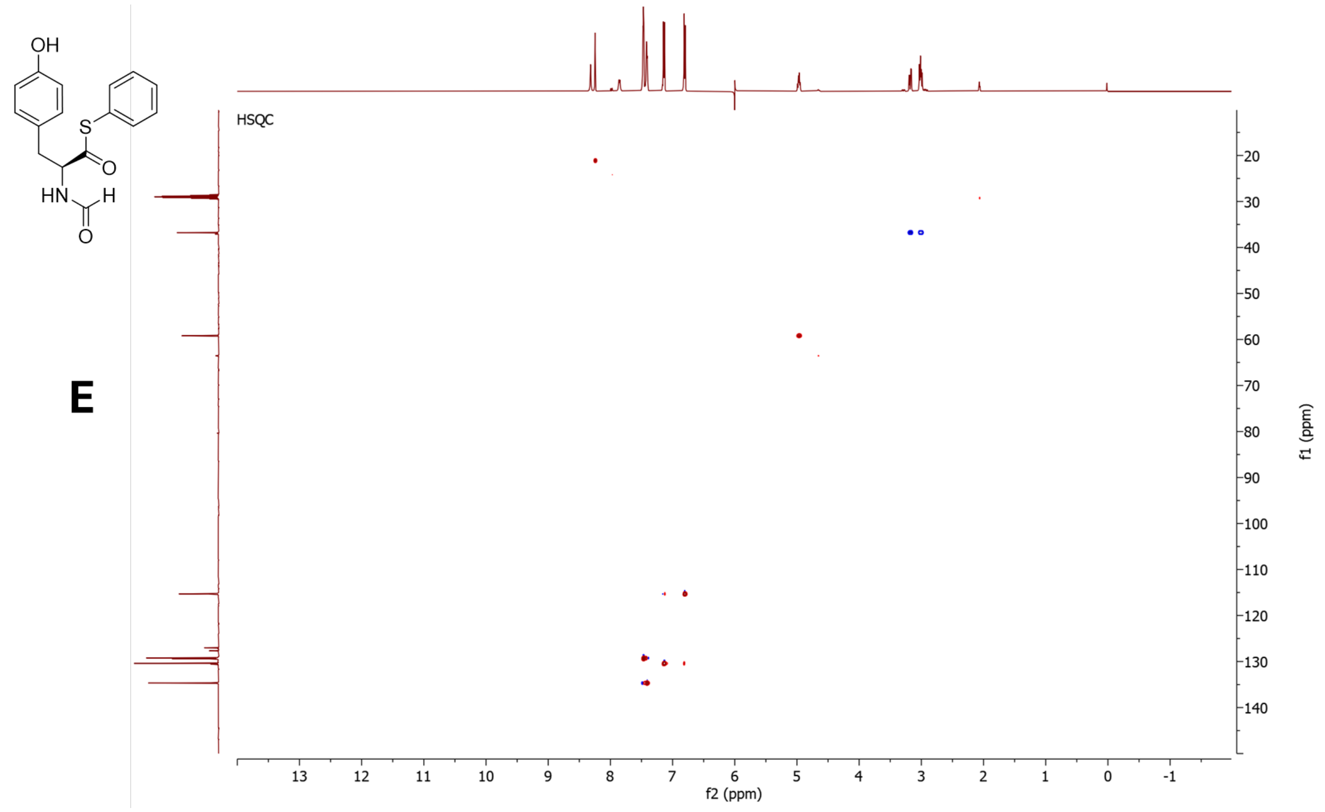


**Figure S60**. NMR data for **7a** in Acetone-$d_6$, 500 MHz. **A)** $^{1}$H-NMR spectrum. **B)** $^{13}$C-NMR spectrum. **C)** COSY spectrum. **C)** HMBC spectrum. **E)** HSQC spectrum. The CHO carbon in the HSQC is aliased

5.5.2 NMR data for **7b**

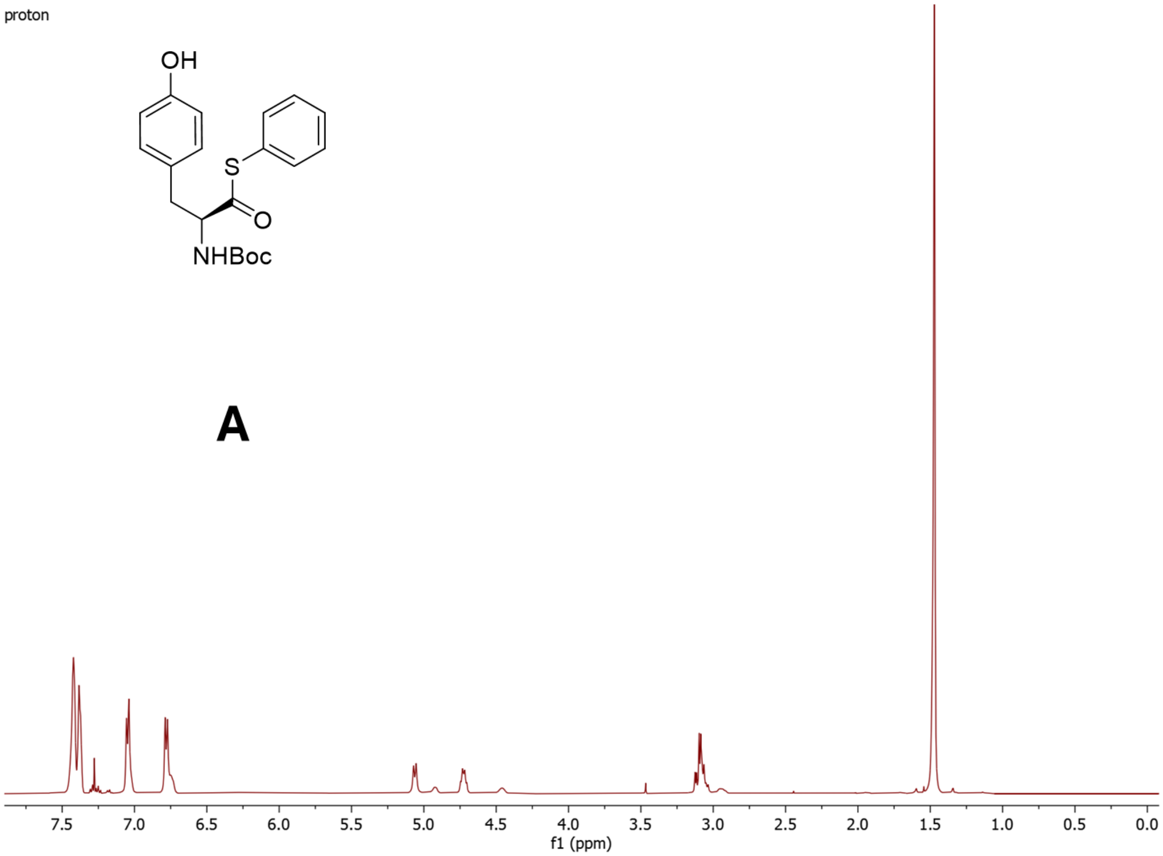


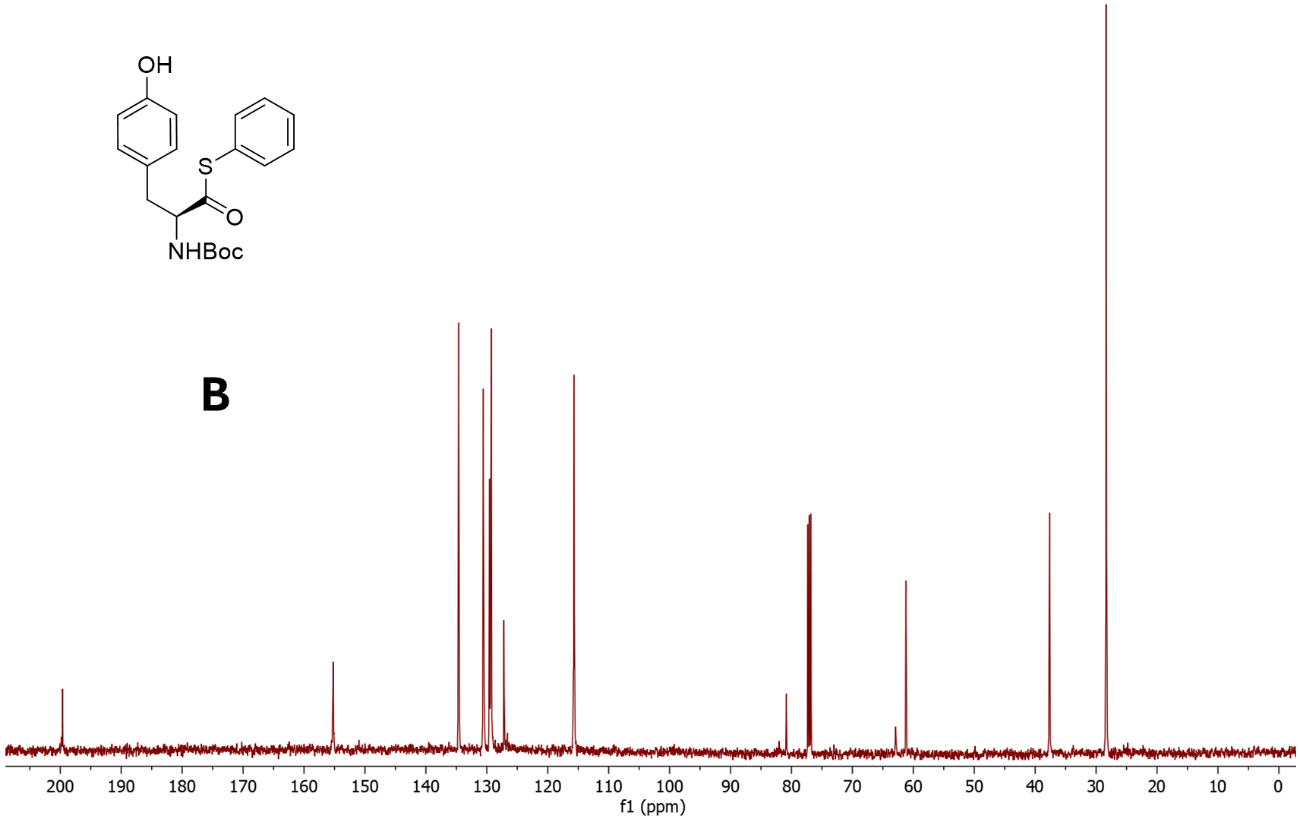

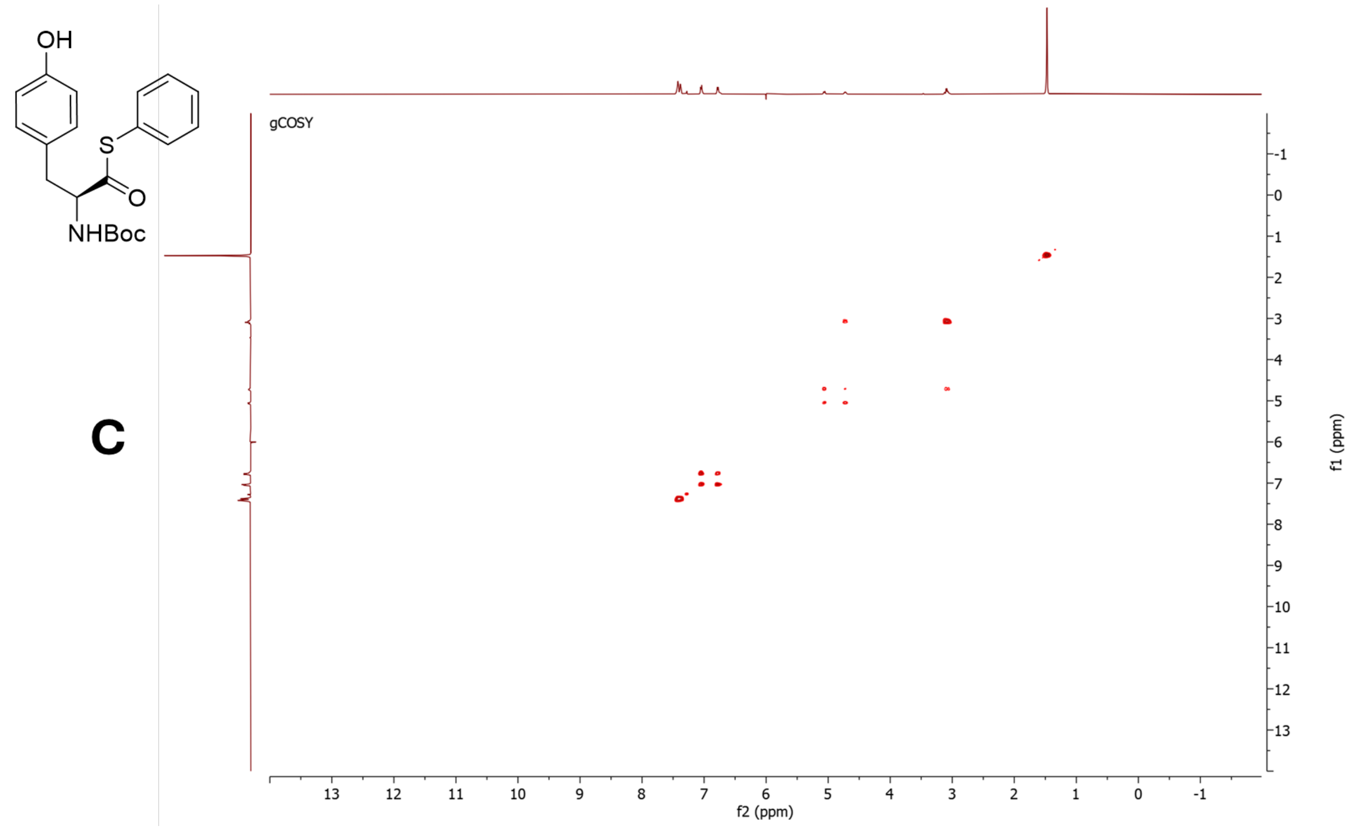


**C**

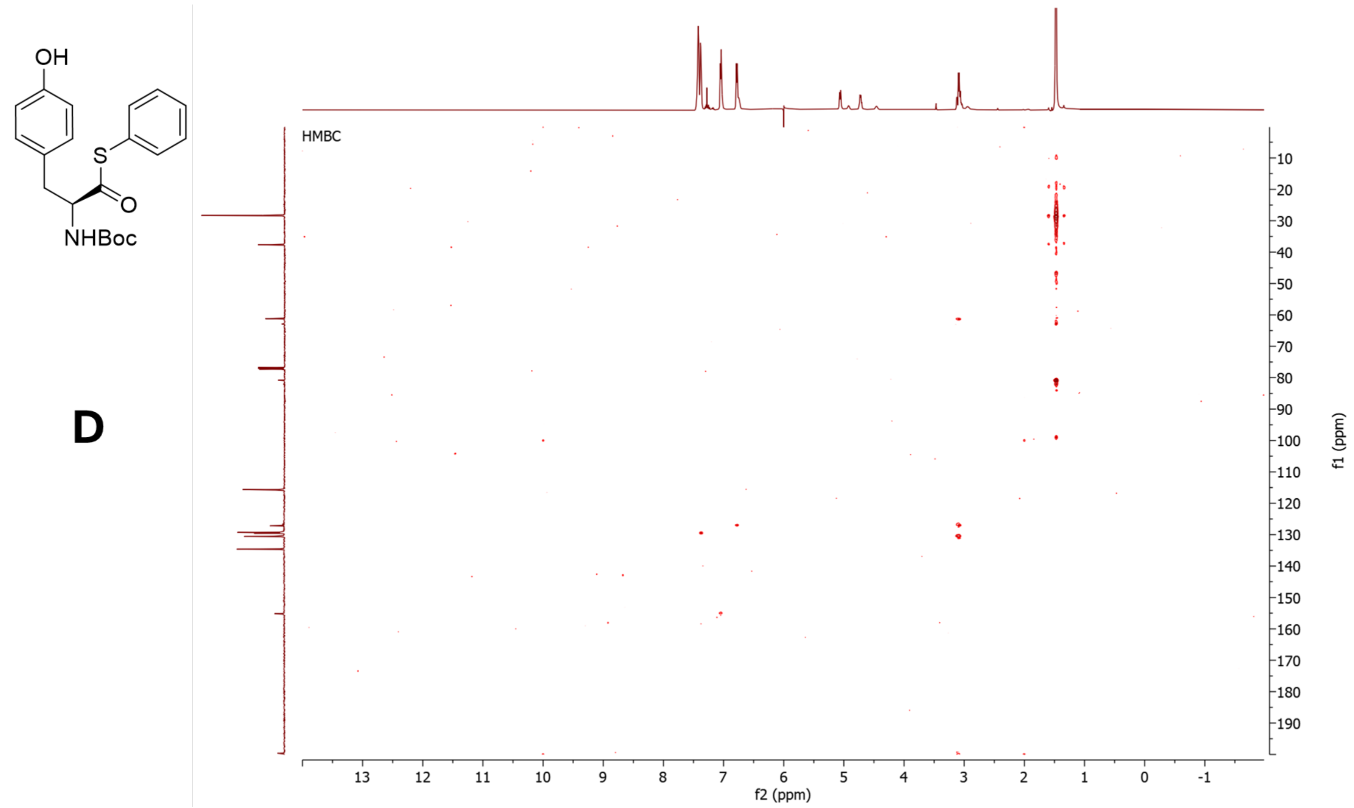


**D**

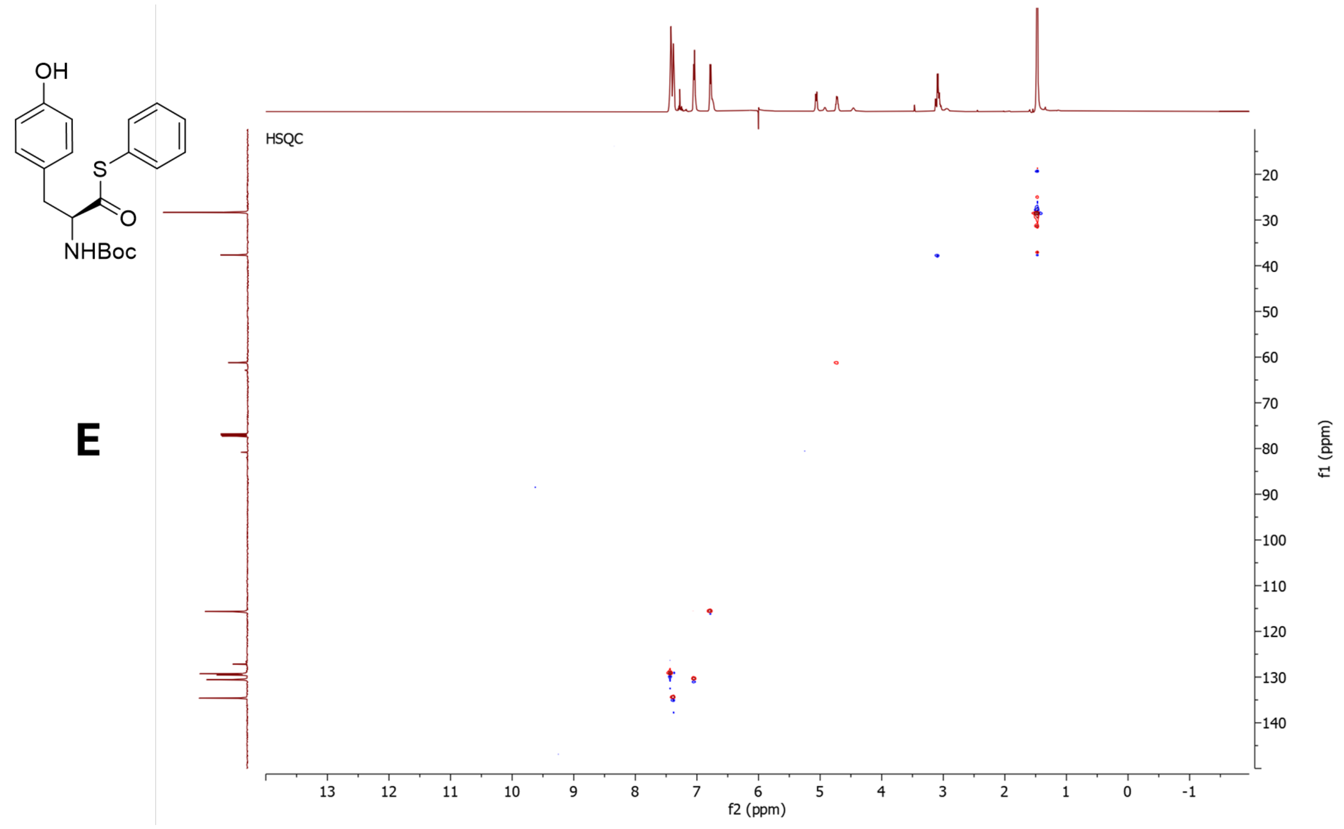


**Figure S61**. NMR data for **7b** in $CDCl_3$, 500 MHz. **A)** $^{1}$H-NMR spectrum. **B)** $^{13}$C-NMR spectrum. **C)** COSY spectrum. **C)** HMBC spectrum. **E)** HSQC spectrum.

5.5.3 NMR data for **7c**

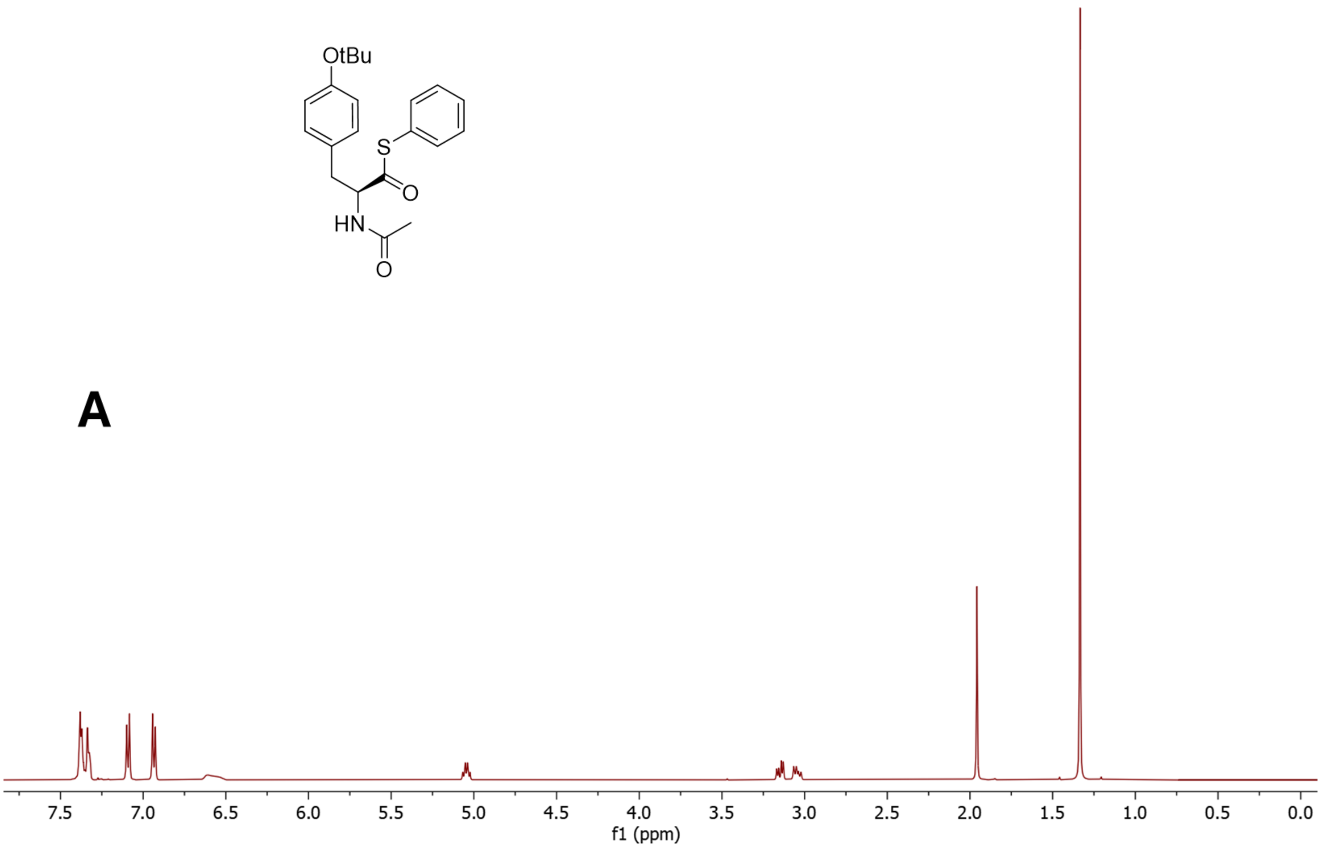


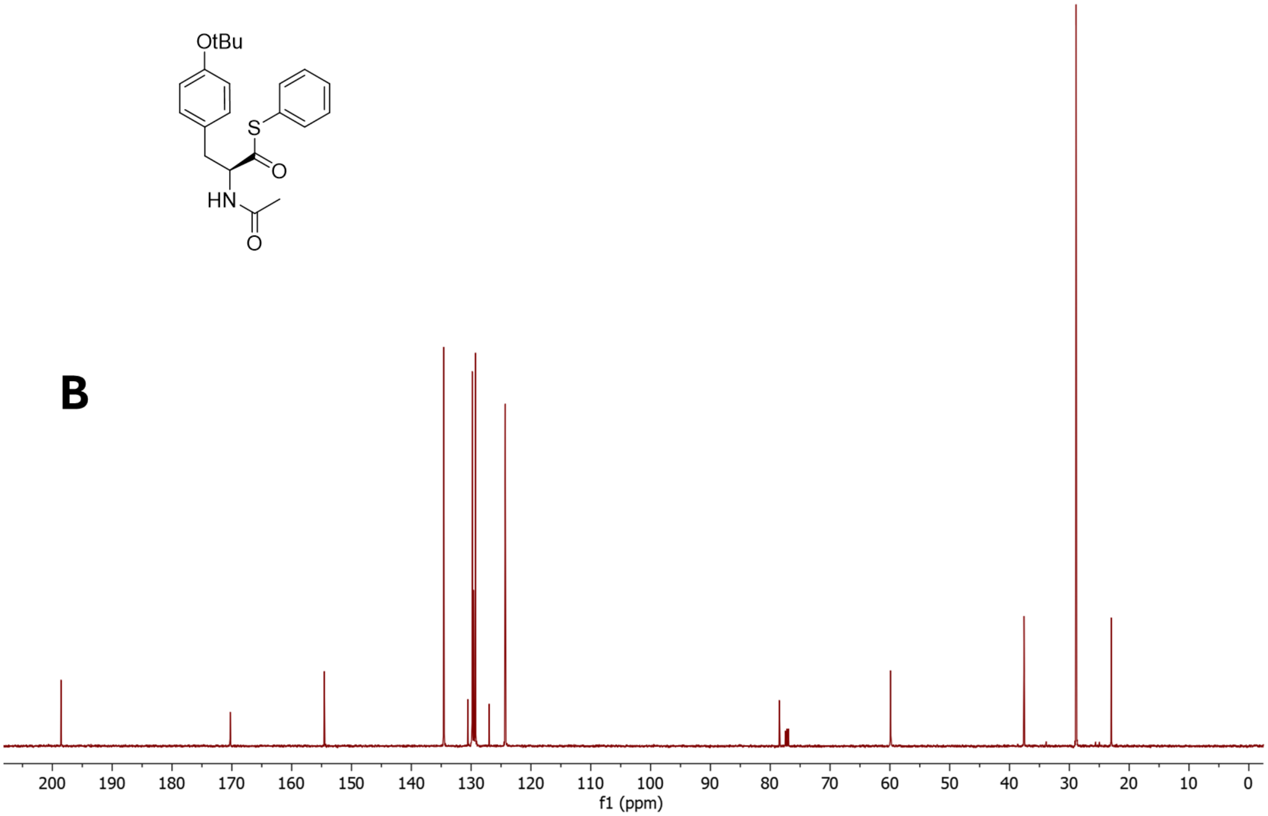

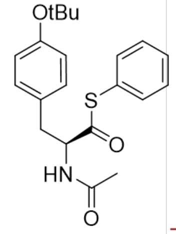


**C**

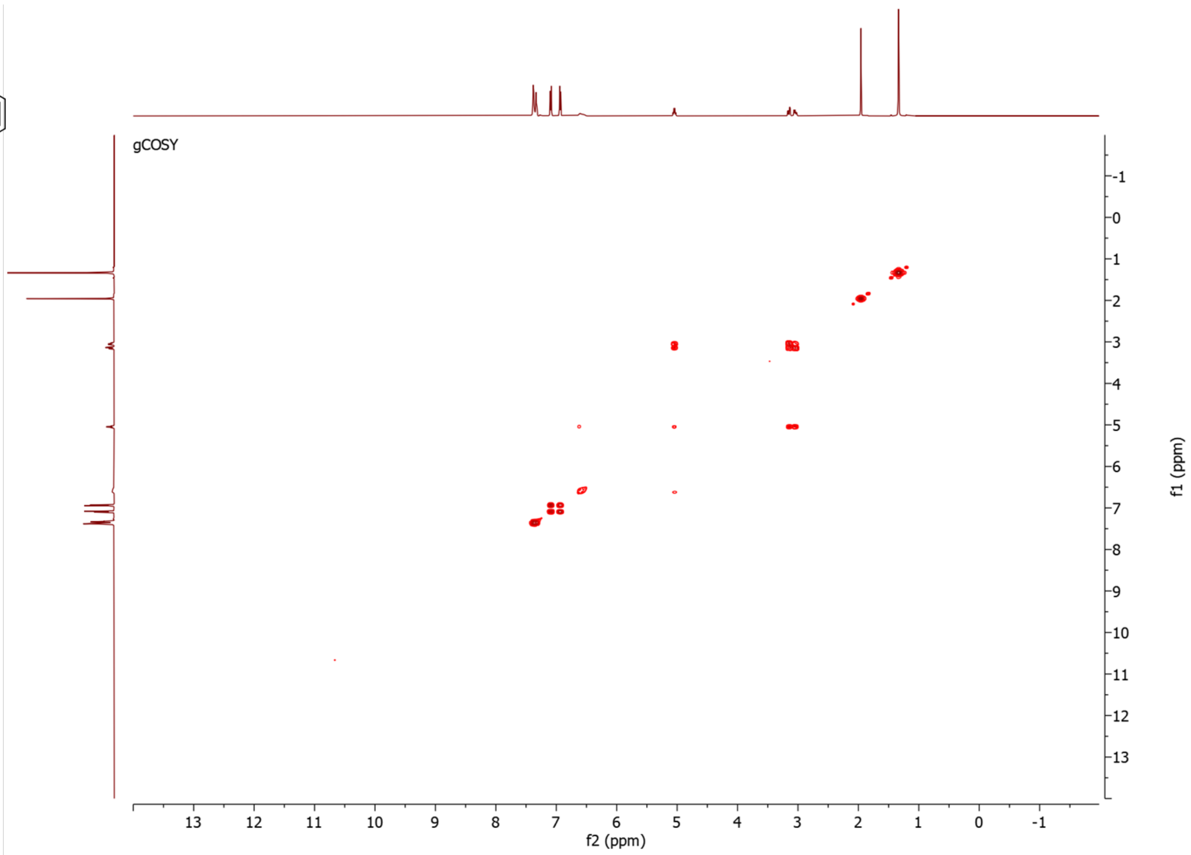


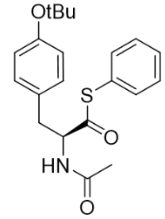


**D**

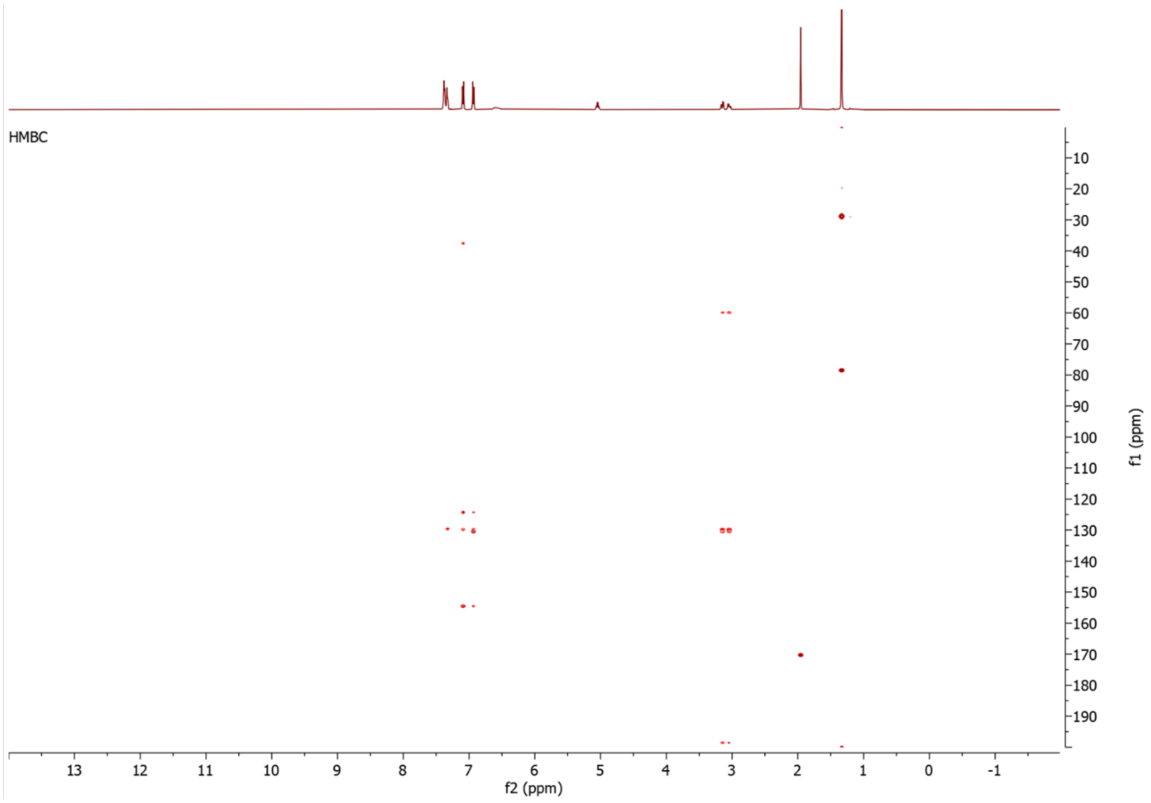

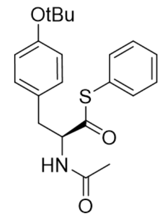


E

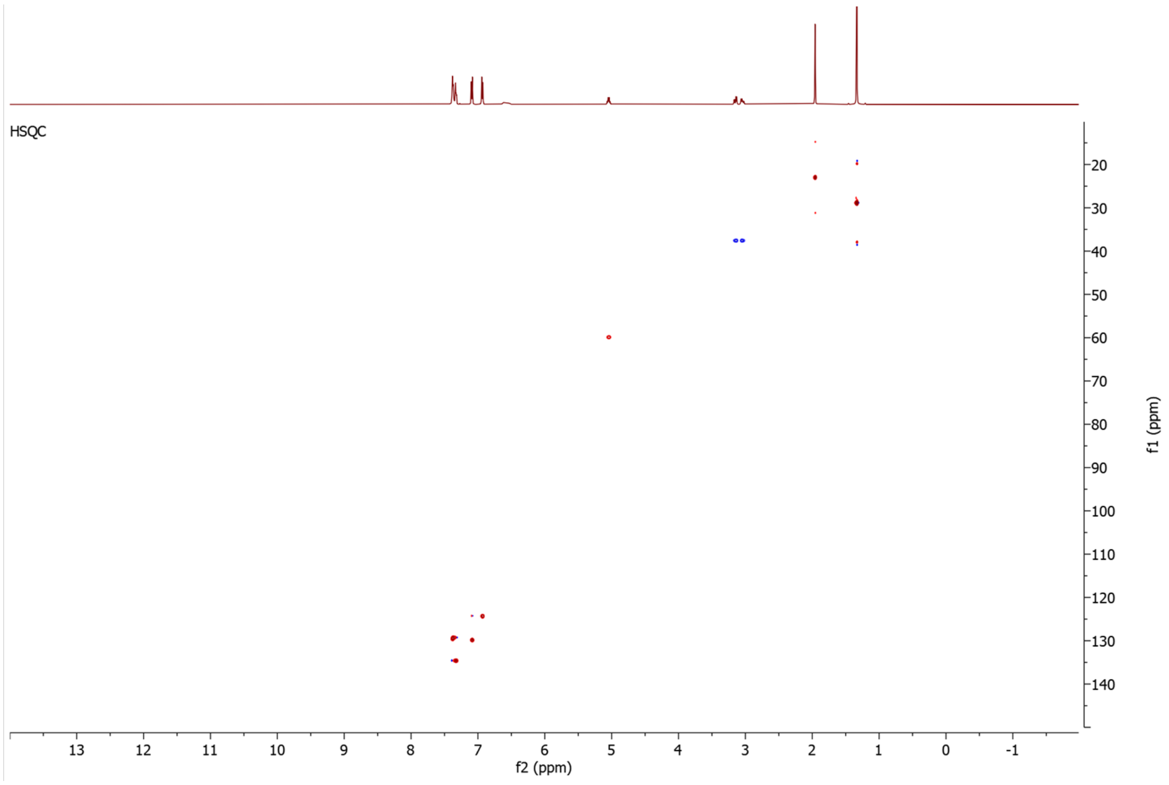


**Figure S62**. NMR data for **7c** in $CDCl_3$, 500 MHz. **A)** $^{1}H$-NMR spectrum. **B)** $^{13}C$-NMR spectrum. **C)** COSY spectrum. **C)** HMBC spectrum. **E)** HSQC spectrum.

#### 6. Supplementary tables

Table S1. Transcriptome SRA datasets containing *bur-ox* homologs (protein sequences used in this study were retrieved from assemblies generated from the SRAs highlighted in green).

| SRA accession | Scientific name of animal |
|---|---|
| SRR1424831 | *Mytilus edulis* |
| **SRR12010104** | ***Mytilus californianus*** |
| SRR13364369 | *Mytilus coruscus* |
| SRR12451931 | *Pinctada margaritifera* |
| SRR6934455 | *Mytilus trossulus* |
| **SRR5357623** | ***Crassostrea virginica*** |
| SRR18332320 | *Congeria kusceri* |
| SRR5188384 | *Limnoperna fortunei* |
| DRR077431 | *Gigantidas platifrons* |
| SRR6169098 | *Corbicula fluminea* |
| SRR24634822 | *Fusconaia mitchelli* |
| ERR3600911 | *Bathymodiolus azoricus* |
| SRR5515063, SRR13983649 | *Tegillarca granosa* |
| SRR17649340 | *Aplysia californica* |
| SRR8134414 | *Mercenaria mercenaria* |
| SRR12212519 | *Panopea generosa* |
| SRR12456155, SRR12456157 | *Bellamya purificata* |
| SRR7009105 | *Lyrodus pedicellatus* |

Table S2: Published bivalve genomes containing homologs of the putative bursatellin biosynthesis genes.

| Bivalve type | Scientific name | Accession number of genomic contigs |
|---|---|---|
| Oyster | *Crassostrea virginica* | NC_035783.1 + NC_035780.1 |
| Oyster | *Pinctada fucata* | AP027107.1 |
| Soft shell clam | *Mya arenaria* | NC_069132.1 |
| Cockle | *Cerastoderma edule* | OX401404.1 + OX401415.1 |
| Oyster | *Crassostrea gigas* | NC_047561.1 + NC_047565.1 + NW_022994918.1 |
| Mussel | *Mytilus galloprovincialis* | CM046166.1 + CM046170.1 |
| Clam | *Tegillarca granosa* | CM056595.1 |
| Oyster | *Crassostrea hongkongensis* | CM027472.1 + CM027471.1 |
| Mussel | *Mytilus edulis* | CM034350.1 + OY782958.1 |

| | | |
|---|---|---|
| Mussel | *Sinohyriopsis cumingii* | CM068825.1 |
| Mussel | *Mytilus coruscus* | CM029599.1 |
| Scallop | *Pecten maximus* | NC_047017.1 |
| Mussel | *Perna viridis* | CM074512.1 |
| Oyster | *Ostrea edulis* | NC_079166.1 + NC_079168.1 |
| Hard clam | *Mercenaria mercenaria* | NC_069361.1 + NC_069363.1 |
| Mussel | *Mytilisepta virgata* | CM051518.1 |
| Oyster | *Crassostrea angulata* | NC_069113.1 + NC_069115.1 |
| Mussel | *Dreissena polymorpha* | NC_068357.1 |
| Clam | *Tridacna crocea* | OX031058.1 + OX031064.1 |
| Oyster | *Crassostrea ariakensis* | CM035816.1 + CM035812.1 |
| Mussel | *Limnoperna fortunei* | OX104113.1 |
| Oyster | *Pteria penguin* | CM068587.1 |
| Mussel | *Bathymodiolus septemdierum* | OY726519.1 |
| Clam | *Panopea generosa* | CM055897.1 |
| Mussel | *Congeria kusceri* | CM051028.1 |
| Oyster | *Magallana gigas* | OY970750.1 + OY970753.1 |
| Oyster | *Crassostrea nippona* | CM065923.1 + CM065919.1 |
| Mussel | *Mytilus trossulus* | NC_086375.1 |
| Clam | *Archivesica marissinica* | CM026404.1 |
| Clam | *Solen grandis* | CM037786.1 + CM037777.1 |
| Clam | *Gari tellinella* | OV277857.1 |
| Clam | *Spisula solida* | OX365945.1 |
| Clam | *Tridacna gigas* | OX244029.2 + OX244040.2 |
| Clam | *Hippopus hippopus* | OX328097.1 + OX328106.1 |
| Clam | *Tridacna derasa* | OY723416.1 + OY723424.1 |
| Clam | *Spisula subtruncata* | OY787661.1 |
| Mussel | *Bathymodiolus brooksi* | OY804576.1 |
| Cockle | *Fragum whitleyi* | OX411544.1 + OX411555.1 |
| Cockle | *Fragum fragum* | OX336366.1 + OX336374.1 |
| Mussel | *Arcuatula senhousia* | OZ020304.1 |
| Clam | *Anadara kagoshimensis* | CM037858.1 |
| Clam | *Lucinisca nassula* | OY757012.1 |
| Cockle | *Fragum sueziense* | OY804593.1 + OY804601.1 |
| Clam | *Ruditapes philippinarum* | NW_026853759.1 + NW_026855673.1 |
| Clam | *Conchocele bisecta* | CM055269.1 |

| | | |
|---|---|---|
| Scallop | *Argopecten irradians* | SAYR01058937.1 |
| Scallop | *Mizuhopecten yessoensis* | NW_018407753.1 |
| Mussel | *Gigantidas platifrons* | MJUT01058264.1 |
| Oyster | *Pinctada imbricata* | VSWD01000009.1 |
| Oyster | *Pinctada margaritifera* | CAXHYN010001331.1 + CAXHYN010001676.1 |
| Mussel | *Mytilus californianus* | NW_026262690.1 |
| Oyster | *Saccostrea glomerata* | PRKT01000059.1 + PRKT01000492.1 |
| Mussel | *Margaritifera margaritifera* | JAQPZY010001606.1 |
| Oyster | *Ostrea lurida* | CAJDSM010054183.1 + CAJDSM010033138.1 + CAJDSM010033137.1 |
| Clam | *Saxidomus purpurata* | JAFMSR010000001.1 |
| Pinnid | *Pinna nobilis* | JACSZP010087345.1 + JACSZP010023644.1 + JACSZP010067024.1 |
| Oyster | *Saccostrea echinata* | NW_026889758.1 + NW_026889756.1 |
| Clam | *Lutraria rhynchaena* | VIBL01000002.1 + VIBL01000270.1 |
| Clam | *Lutraria lutraria* | CAXIEH010000924.1 + CAXIEH010000893.1 |
| Cockle | *Serripes groenlandicus* | JASEVX010000001.1 + JASEVX010000023.1 |
| Clam | *Cyrtodaria siliqua* | JAWPHD010002704.1 + JAWPHD010000445.1 |
| Mussel | *Geukensia demissa* | JARWCS010092333.1 + JARWCS010334910.1 |
| Scallop | *Crassadoma gigantea* | JAULCA010449267.1 + JAULCA010372609.1 + JAULCA010305786.1 |
| Mussel | *Dreissena rostriformis* | VMBQ01001933.1 + VMBQ01002005.1 |
| Mussel | *Unio pictorum* | JARLTB010000022.1 |
| Mussel | *Unio delphinus* | JAQISU010000010.1 |
| Clam | *Cyclina sinensis* | JAAONU010000011.1 |
| Mussel | *Potamilus streckersoni* | JAEAOA010001436.1 + JAEAOA010000659.1 |
| Scallop | *Ylistrum balloti* | NW_026782274.1 |
| Mussel | *Modiolus philippinarum* | MJUU01004508.1 + MJUU01039902.1 |
| Cockle | *Clinocardium nuttallii* | JAUELF010183073.1 + JAUELF010045067.1 |
| Oyster | *Ostrea denselamellosa* | JAMZEH010000010.1 + JAMZEH010000028.1 |
| Pinnid | *Atrina japonica* | BROG01000650.1 |
| Mussel | *Megalonaias nervosa* | JAECUM010014877.1 + JAECUM010013324.1 |
| Mussel | *Venustaconcha ellipsiformis* | QKMX01092772.1 + QKMX01284489.1 + QKMX01350545.1 |
| Scallop | *Chlamys rubida* | JAUMHH010117908.1 + JAUMHH010212981.1 + JAUMHH010148904.1 |
| Oyster | *Magallana hongkongensis* | WFKH01006014.1 + WFKH01011925.1 |
| Clam | *Pododesmus macrochisma* | JAUMGZ010238297.1 + JAUMGZ010166265.1 + JAUMGZ010160049.1 |
| Mussel | *Geukensia granosissima* | JAULBM010312432.1 + JAULBM010326986.1 + JAULBM010032995.1 |
| Clam | *Saxidomus gigantea* | JAUMHA010304471.1 + JAUMHA010121608.1 |
| Mussel | *Lithophaga antillarum* | JAOXYI010577611.1 + JAOXYI010377393.1 + JAOXYI010606623.1 |

| | | |
|---|---|---|
| Mussel | *Botula fusca* | JAOXYH010050630.1 + JAOXYH010694795.1 + JAOXYH010156300.1 |
| Clam | *Rugalucina vietnamica* | CAXHKZ010001552.1 + CAXHKZ010001478 |
| Mussel | *Ctena decussata* | OZ022499.1 |
| Clam | *Anodontia alba* | CAXHKK010009825.1 + CAXHKK010005800.1 + CAXHKK010002056.1 |
| Clam | *Loripinus fragilis* | OZ026514.1 |
| Clam | *Corbicula japonica* | JAZAYU010000006.1 + JAZAYU010000078.1 |
| Oyster | *Crassostrea gasar* | JBEEQF010000007.1 + JBEEQF010000032.1 |
| Oyster | *Crassostrea rhizophorae* | JBEOLP010000152.1 + JBEOLP010000129.1 |
| Clam | *Mysia undata* | OZ067060.1 |
| Clam | *Lucinella divaricata* | OZ077285.1 |
| Clam | *Stewartia floridana* | CAXMME010000765.1 + CAXMME010004148.1 |
| Clam | *Venus verrucosa* | OZ121644.1 |
| Clam | *Tivela stultorum* | JBCDMG010000002.1 |
| Clam | *Arca noae* | OZ178933.1 |
| Clam | *Heteranomia squamula* | CAXVUU010000068.1 |
| Clam | *Ctenoides ales* | CM090420.1 |
| Clam | *Anadara tuberculosa* | JBHJMB010000028.1 + JBHJMB010000023.1 |
| Tellin | *Macomangulus tenuis* | OZ060682.1 |
| Mussel | *Solenaia oleivora* | CM078438.1 |
| Clam | *Donax trunculus* | OZ187422.1 |
| Clam | *Americardia media* | CAYAAN010000001.1 |

Table S3: Published gastropod genomes containing homologs of the putative bursatellin biosynthesis genes.

| **Gastropoda** | **Scientific name** | **Accession number of genomic contigs** |
|---|---|---|
| Caenogastropoda | *Littorina saxatilis* | CM074575.1 |
| Vetigastropoda | *Haliotis asinina* | CM074533.1 + CM074528.1 |
| Heterobranchia | *Onchidella celtica* | OZ007569.1 |
| Caenogastropoda | *Semisulcospira habei* | BTPG01000004.1 |
| Vetigastropoda | *Lepetodrilus gordensis* | JAVCKX010185751.1 + JAVCKX010122874.1 + JAVCKX010122435.1 |
| Vetigastropoda | *Lepetodrilus elevatus* | JAVCKZ010068827.1 + JAVCKZ010094824.1 |
| Heterobranchia | *Berghia stephanieae* | CM067442.1 |
| Heterobranchia | *Ellobium chinense* | JAWQUT010000032.1 |
| Caenogastropoda | *Alviniconcha strummeri* | OY757543.1 |
| Vetigastropoda | *Clypeosectus delectus* | JAULBO010026924.1 + JAULBO010135183.1 |
| Vetigastropoda | *Lepetodrilus pustulosus* | JAULBP010109531.1 + JAULBP010117495.1 + JAULBP010096792.1 |
| Vetigastropoda | *Lepetodrilus ovalis* | JAUTBQ010165611.1 + JAUTBQ010143669.1 + JAUTBQ010175210.1 |

| | | |
|---|---|---|
| Vetigastropoda | *Lepetodrilus galriftensis* | JAUTBZ010087442.1 + JAUTBZ010035238.1 + JAUTBZ010011212.1 |
| Vetigastropoda | *Lepetodrilus schrolli* | JAUTBT010071510.1 + JAUTBT010062808.1 |
| Vetigastropoda | *Lepetodrilus fucensis* | JAUTBV010074313.1 + JAUTBV010111966.1 + JAUTBV010109127.1 |
| Vetigastropoda | *Lepetodrilus tevnianus* | JAUTBX010034294.1 + JAUTBX010117487.1 + JAUTBX010070023.1 |
| Neomphaliones | *Nodopelta heminoda* | JAUTCA010249081.1 + JAUTCA010268385.1 + JAUTCA010021302.1 |
| Vetigastropoda | *Diodora aspera* | JAUELE010218786.1 + JAUELE010134255.1 + JAUELE010100431.1 |
| Caenogastropoda | *Sinotaia purificata* | CM054177.1 |
| Vetigastropoda | *Haliotis rufescens* | JALGQB010000028.1 + JALGQB010000011.1 |
| Vetigastropoda | *Haliotis cracherodii* | JAJLRC010000005.1 + JAJLRC010000001.1 |
| Vetigastropoda | *Haliotis laevigata* | VKKT01011081.1 + VKKT01006944.1 |
| Nudibranchia | *Phyllidia flava* | OZ184099.1 + OZ184103.1 |
| Caenogastropoda | *Littorina brevicula* | CM093034.1 |

Table S4: PCR primers used in this study.

| **Primer name** | **Primer DNA sequence** |
|---|---|
| Cv-TR_fwd | 5’ – ataagactagtgatgaatctattcatgaacaaac – 3’ |
| Cv-TR_rev | 5’ – tgatgcacgtgttcagtcaaaccaaaaaattgt – 3’ |
| CvATR_fwd | 5’ – gagatacctaaacaaaccg – 3’ |
| Flag-TS_rev | 5’ – actagtcttatcatcatcatc – 3’ |

Table S5. HSQC data for CoA esters **3Ca**-**3Cc** (500 MHz, DMSO-$d_6$)

| CoA deriv. | *N*-Boc-L-Tyr (**3Cb**) | | *N*-formyl-L-Tyr (**3Ca**) | | *N*-Ac-*O*-tBu-Tyr (**3Cc**) | |
|---|---|---|---|---|---|---|
| #[a] | δH | δC | δH | δC | δH | δC |
| R | 1.34 (3CH3) | 27.6 | 8.03 (CH) | 160.4 (folded) | 1.81 (CH3) | 21.8 |
| *O*-tBu | - | - | - | - | 1.25 (3CH3) | 28.0 |
| A[b] | - | 201.2 | - | 199.3 | - | 200.1 |
| B | 4.12 (CH) | 62.0 | 4.56 (CH) | 58.8 | 4.50 (CH) | 60.0 |
| C | 2.91 (CH2)<br>2.66 | 35.1 | 3.06 (CH2)<br>2.84 | 35.3 | 3.03 (CH2)<br>2.78 | 35.4 |
| E | 7.02 (2CH) | 129.4 | 7.09 (2CH) | 129.6 | 7.13 (2CH) | 128.9 |
| F | 6.64 (2CH) | 114.2 | 6.66 (2CH) | 114.4 | 6.87 (2CH) | 122.8 |
| 8[c] | 8.62 (CH) | 140.3 | 8.62 (CH) | 140.1 | 8.60 (CH) | 140.0 |
| 1' | 6.00 (CH) | 86.8 | 6.00 (CH) | 86.8 | 5.97 (CH) | 86.7 |
| 2' | 4.72 (CH) | 72.2 | 4.72 (CH | 72.2 | 4.72 (CH) | 72.0 |
| 3' | 4.78 (CH) | 73.1 | 4.81 (CH) | 73.0 | 4.81 (CH) | 73.0 |
| 4' | 4.40 | 81.1 | 4.41 (CH) | 81.1 | 4.38 (CH) | 80.1 |
| 5' | 4.19 (CH2) | 63.9 | 4.19 (CH2) | 64.0 | 4.19 (CH2) | 63.4 |
| 1" | 3.91 (CH2)<br>3.53 | 71.6 | 3.53 (CH2) | 71.5 | 3.91 (CH2)<br>3.53 | 71.5 |
| 3" | 3.75 (CH) | 72.7 | 3.75 (CH) | 72.6 | 3.75 (CH) | 72.6 |
| 5" | 3.31 (CH2)<br>3.22 | 34.2 | 3.31 (CH2)<br>3.25 | 35.2 | 3.31 (CH2)<br>3.25 | 34.2 |
| 6" | 2.25 (CH2) | 34.5 | 2.28 (CH2) | 34.5 | 2.28 (CH2) | 34.5 |
| 8" | 3.12 (CH2) | 37.4 | 3.16 (CH2) | 37.5 | 3.16 (CH2) | 37.5 |
| 9" | 2.84 (CH2) | 26.9 | 2.88 (CH2) | 27.2 | 2.88 (CH2) | 27.1 |
| 10" | 0.94 (CH3) | 20.5 | 0.94 (CH3) | 20.5 | 0.94 (CH3) | 20.6 |
| 11" | 0.75 (CH3) | 18.2 | 0.75 (CH3) | 18.2 | 0.75 (CH3) | 18.2 |

[a]Numbering and chemical shift assignment comparison: Wu et al., *J. Am. Chem. Soc.* **1998**, *120*, 39, 9988–9994.

[b]Called using HMBC data between A and 9" and A and C.

[c]H-2 could be observed in 1D spectrum but did not give an HMBC signal.

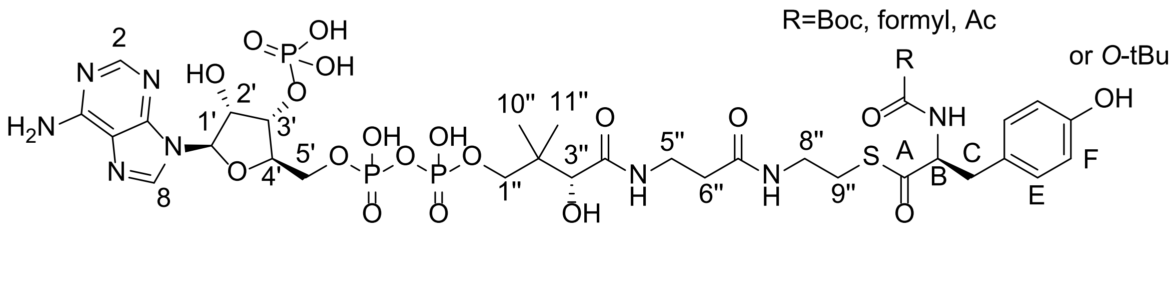

### 7. Miscellaneous data

#### 7.1. Codon-optimized DNA sequences

*Cvfmt-atr*

ATGATGAAAGATGCTTCTGCTTCTTCTATTGCTATGGAAGAAGATATTTGTTGTGCGGTAATCGGTGAAGGGAATATCCTTCTTAGCTGT
CTTAAAATATTGGAAAAATCTAACATAAAAATAGTGGCTGTTTTTACTGAAACTCCATCTATTAGATCTTATTGTGATGAATCTTGTATTC
AATGGTGGGAGCGGACTTCAAGACTAGAAGAAATCTTGAAGGATAAGAAGATACATTATTTGTTCTCTATTTCTAATCCACGGATATTGA
AGGAGAATGAACTTAATGTTCCAAAGATTTTGACTATTAATTATCATGATTCTCCATTGCCTGCTTATGCTGGTGTTCATGCTACTTCTTG
GGCTATTATTAATGGTGAAACTGAACATGGTATTTCTTGGCATGTTGTTGAACCTGGTATTGATACCGGTGACATTCTGAAGTTTAAGAA
AATTGAAATTGATAAAAAAGAATCTGCGCTGGCTTTGAATTTGAAATGTCAAGAAGCTGGTATCCAATGTTTTGAGGAGCTGGTGAATG
ACATCAAAACTAATTCTATTGTTAGATCTCCACAACCAAAAAACGGTAGATCTTATTTTGGTTTGCATGCTATTCCACCAAATTTGGGTG
TGTTGTCTTTTGATAGAAAATCTAAAGATACTTTTAATTTGGCTAGAGCTTTGGAATTTGGTCATCATGAAAATTCTTTGGGTTCTGCTA
AATTGTTGACTTCTTCTGGTGAATTTTTGATGGTTATTTCTACTGAAGTTGCTTGTAGATTTCCAACTTCTGATACTAAACCTGGTACTA
TTTTGGATATTGTTGATGATTCTATTATTGTTTCAACTTTGGGTGAACCATTGTTGATGTCTGTTGCTAAATTGGATGGTGCTTCTATTC
CACAATGTAATTTTAGAAAACATGGTTTGGTTACTGGTTTGCGTCTGGATTCATCTTTCACTATTGACACGGAATGCCTTTTGCAGATAC
GCAAGAGGGAATCTTTTTGGAAAAGAAAACTAGAAAGATACGAGCCAACTGTATTTTTTAAACAAAGAATGAACGTAGTTGCTAATATT
CTTGACTTGTCTGATGAAGTACATTTGGAAACTAAAACTGTCAAACTCCCATTCGTTTCTAATGATGATCATACTTTTGTTAAATCTGCT
TTGGTTGCTTTTATTGGTAGAACTTGTTGTACTCCTGATGTTAATATTGGTTTGATTGCTAATAAATCTCAAATTCCAAGAAATGCTGAA
ACTTTGTATTCTGATATTTGCCCTGCTGTTTTTAGATTGAACATGACTGACGAAATTAATGATGTGATTGGAACATGTCAAAAGGACCTA
ATGAAATATGAAGAATCTATGTCCTTTTTGGTTGATGTGTTTTACAGATATCCTGAATTACGTGAAAGAAAGCAAACTCCACATCATAAT
ATTGTTATCGGCACTTCTGTCGATGTTTTATCTGATATACAAATCGGTAAGAAAGCCTTGCAAGACTGCAATATTTTGCTTCTGTACTCCC
ACGATATGAATGAGATTTATATTCATTACTCTATGAGACCGGGTCAAGACCTATCCTTTATCTTGGATGTTTTCAAACACTTTCCCATTTT
CTTGGAGTCTGTGACTAAAAACCCTATTCAATTGTTGTGTGAAGTTTCTTTTGTTCCATTGGATGAATTGAAAGTTTTGTATCCATCTCC
GACTGAGATACCTAAACAAACCGGTTCTCAAATTATTAATATTTTTGAGAACTGCGTTGAATGTTATAGATCTAGAATTGCTTTGCAAAC
TTCTACTCTATCTGCTACTTATGAAGAAGTTTTGCAAATGGTTGAGAATTTGACTGAATTGTTGAGAGAAACTGCTTTGAATGGTAGAA
AAAGAGTTATTGGCTTGCACTTGCCTAACAGTATTGCTTACGTTATTTCTGTTTTGTCTGTTTTGAAATGTAAACATGTTTTTTTGCCAT
TGCCATTGGATTATCCACATGAAAGATTGGTTTTTACTATGAAAGATTCTAGAGTACAAACTGTTGTGACTACTAAGGAAAGAGTTGAAC
AGATTGACTTTAACGAGTTGTCTGCTAATCCAAAAATTGTTAATGCCGGAACGGTAGATGGGATAGAGTTGTATATTGTGCGGTTTTATG
AGTTGGATGACTTTGAGAGATGGAACTCTAAATTGCCGCAGCCAAAGGACAATTTTGAGAATTTAGCATATATTATGTATACTTCTGGTT
CTACTGGTAAACCAAAAGGTGTTAAAGTTAAAGAATCTTCTGTTGTAAATTTGGCTAAAGCGCAAATTCAATTGTGGGATTTGAATCCA
AAAGACGTCATTGGTCAATTCGCTTCTGTTGGTTTTGATGCTACTATTTCTGAAATTTTTACTTCTTTGTTTTCTGGTGCTTCTTTGGTT
GTTTTTGAAGAAAAAGAAAGATTGGGACAAGAGTTCTTAATGGCTATGAACAAGCATAAAATCACTACTATTACTTTGCCACCGTCTTT
GCTTAACATCTATTCTCCAAAAGATTTGCCATATTTGAAAAAAATTGTTACTGCTGGTGAAGCTTGTACTTTGTCTACTGCTGTTAAATG
GGCTGTTAAAAATGAAAGAAGATTTTTTAATGCTTATGGTCCAACTGAAGCTACTGTTTGTGCTACTTGTTTTGAATTTATGCCTGAAAA
TAAACATGAAGATGTTAATTGTGAATTGCCAATTGGTATTGGTATTCCTGGTGTTGATGTTTATTTGTTTGATGATTATTTGAAACCATT
GCCACCTGGTGTTATCGGTGAAATTTACATCGGTGGCTTGGGTTTGTCTGAAGGTTATCATGGTCATGCTTCTCATTTGACTAAAGAAAA
ATTTGTTCAACATCCATTGGTTAATTCTCCGTCTTTGTTGTATAGAACTGGTGATCATGGTTTGCAAGATTCTTCTGGTAATATTACTTTT
GTTGGTAGATTGGATGATATGGTTAAAATTAGAGGTCAAAGAGTTGATTTGTCTGAAATTGAACAAGTTTTGATTCAACATCCAAAAGT
AGATGTTGCAGTTGTTGTTCCACATAGATGTATTAATAATAATGAAATTTCAATTGCCGCGTTTGTTGCTCCAACTTTTATTTTTTCTTC
AGAACTCCGTGAGTATTTAGTTAAAGTATTGCCAAAATTTATGATTCCCACTTTTATTAAAAAACTTGATCTCAGCGACTTTCCCAAAAC
CATAAACGGTAAATTGGATCGCAAGAAATTGGGCAAAGATGAATCTATTCATGAACAAACTGAATCTGTTGGTCATTCTCATTTGAATG
AATCTCAGTTGCAGATCGCACAAATCTGGTGTACTATCCTCAAATTGAATTCTTCTTACGCTTACTCTTTGCATAGAAAATCTTCTTTTT
CTGAATTGGGTGGTAATTCTTTGCAATTGGTTTTGTTGCAAAGACATTTGGAAGAAACTTTTGGTTTGGTGTTGAGTTTTACTGACATT
GGTTCTGCTGACACAATCGAGGAGTTCTCTGATGTCGTCAAAAGAAAAAAGGACATTCTACAAAAGAACGAACAAACTCATCCTAAGGA
TGAAGAGGACTTGAGAACTCTCATTATTAATGACTCTGAATTTACTTCTGATTTGTTTTTGCCACAAGCTAGAAGAGGTTCTGTTGCTAT

TCCTCATACTTCTGGGACGAATCCAAAGTCTTACCTTAGATATCCGAAAAACATTTTGATTTCTGGTGTTACTGGTTTTTTGGGTGCTTT
TTTGTTGTCTGAATTGTTGGAAAAATCAAATGCGCATATTTGTTGTATGGTTAGAGAAACTTCTGAAACTAGAGGTGTTGGTCGGATTG
TAGAAAACTTGCGACGGTATAATTTGTGGAAATTTGAATACACTAACAGAATTGCTGTTGTTATTTCTGATTTGTCTCAACCAAGATTGG
GTATTGCTCCTGATATTTATAATTCTTTGTGTAATTCTATCGATGCTGTATTTATGAACGCGGCTATGATGAACTTTAATACTGACTACCA
AGACCATAGAACTGCTAATGTGTTGTCGACTAAGGAATTCATCAAATTTGCTTTGACTGGTGTTCAAAAAATGTTGTTTTCTACTAGCAC
TCTGGGTGTTTTTTTGTTTCCACCAAAACCTGGTCCTGGTGAACCAATGCATCCAATTATGTTTGAATGTGATGAAGTTGAAGATCCATC
TGATATTGCTGGTGGTTATGGTCAATCTAAATGGGCTTCTGAAAGATTGATTATGCAAGCTTTGGATTTGTTGCCTGGTGGTGCTATTTT
TAGACCTGCTAGAATTTCTGGTTGTACTACTTCTGGTATTGGTCCAAGAAATGATTTGTTTGCTTCTACTATTATTGGTATGAGAAAATT
GGGTTGTTTTCCTGATATGGATTTTCCATATGATTTGACTCCTGTTGATTTTGTTGCTAAAGCCATCGTTGAGATTTCTTTGAAAATCTG
TAACGACCGCGAAAATAGTTATGAGAGAATCTTCCATTTGTTTAATAAGAACACAATGCCTTTTAATAGATTGTTTGATGGTGAAAATAT
TGAACCATTGTCTTTGGAAGAATGGAGAAAAGTTTTGAAATCTGCTCCTGAAGATAATAAAGAATTGATTCCATTGACTCCATTTTTTT
TTTCTTCTTTTTGGGATAGATCTCCATACTGGCCAATTTTCGATACTACAAATACAGACTCATTGATTTCTAATGAAACTAAAGAACTCT
TGAAACCGTCTGAAGAATTGTTGGTTGTTTATAAACAATTTTTTGGTTTGACTGAA

_Cv-tr_

GATGAATCTATTCATGAACAAACTGAATCTGTTGGTCATTCTCATTTGAATGAATCTCAGTTGCAGATCGCACAAATCTGGTGTACTATC
CTCAAATTGAATTCTTCTTACGCTTACTCTTTGCATAGAAAATCTTCTTTTTCTGAATTGGGTGGTAATTCTTTGCAATTGGTTTTGTTG
CAAAGACATTTGGAAGAAACTTTTGGTTTGGTGTTGAGTTTTACTGACATTGGTTCTGCTGACACAATCGAGGAGTTCTCTGATGTCGT
CAAAAGAAAAAAGGACATTCTACAAAAGAACGAACAAACTCATCCTAAGGATGAAGAGGACTTGAGAACTCTCATTATTAATGACTCTG
AATTTACTTCTGATTTGTTTTTGCCACAAGCTAGAAGAGGTTCTGTTGCTATTCCTCATACTTCTGGGACGAATCCAAAGTCTTACCTTA
GATATCCGAAAAACATTTTGATTTCTGGTGTTACTGGTTTTTTGGGTGCTTTTTTGTTGTCTGAATTGTTGGAAAAATCAAATGCGCAT
ATTTGTTGTATGGTTAGAGAAACTTCTGAAACTAGAGGTGTTGGTCGGATTGTAGAAAACTTGCGACGGTATAATTTGTGGAAATTTGA
ATACACTAACAGAATTGCTGTTGTTATTTCTGATTTGTCTCAACCAAGATTGGGTATTGCTCCTGATATTTATAATTCTTTGTGTAATTCT
ATCGATGCTGTATTTATGAACGCGGCTATGATGAACTTTAATACTGACTACCAAGACCATAGAACTGCTAATGTGTTGTCGACTAAGGAAT
TCATCAAATTTGCTTTGACTGGTGTTCAAAAAATGTTGTTTTCTACTAGCACTCTGGGTGTTTTTTTGTTTCCACCAAAACCTGGTCCTG
GTGAACCAATGCATCCAATTATGTTTGAATGTGATGAAGTTGAAGATCCATCTGATATTGCTGGTGGTTATGGTCAATCTAAATGGGCTT
CTGAAAGATTGATTATGCAAGCTTTGGATTTGTTGCCTGGTGGTGCTATTTTTAGACCTGCTAGAATTTCTGGTTGTACTACTTCTGGTA
TTGGTCCAAGAAATGATTTGTTTGCTTCTACTATTATTGGTATGAGAAAATTGGGTTGTTTTCCTGATATGGATTTTCCATATGATTTGA
CTCCTGTTGATTTTGTTGCTAAAGCCATCGTTGAGATTTCTTTGAAAATCTGTAACGACCGCGAAAATAGTTATGAGAGAATCTTCCATT
TGTTTAATAAGAACACAATGCCTTTTAATAGATTGTTTGATGGTGAAAATATTGAACCATTGTCTTTGGAAGAATGGAGAAAAGTTTTG
AAATCTGCTCCTGAAGATAATAAAGAATTGATTCCATTGACTCCATTTTTTTTTTCTTCTTTTTGGGATAGATCTCCATACTGGCCAATT
TTCGATACTACAAATACAGACTCATTGATTTCTAATGAAACTAAAGAACTCTTGAAACCGTCTGAAGAATTGTTGGTTGTTTATAAACAA
TTTTTTGGTTTGACTGAA

_Mcal-act_

ATGGCGGATAAAAACGCGCCGGTGCTGACCGATGCGATTAAAAGCTATGATGGCATTTTTAAAGGCATGTATGATGATGATCTGCTGGAA
AAATTTCTGCGCATTCGCCGCCATCTGGCGGAAGGTCAGCTGAAAAAAGCGCAAGCGCTGCTGGAAGAAAGCCTGAGCGAATGCGGCGA
ACATCCGAGCGATCGCAGCACCCTGGAACTGCTGAAATATGAACTGGATCGCTATATTACCTTTCATATTACCAACAGCTATAAAATTGTG
GATGATGTGAACTGGACCGAAATTAACACCCGCAAACTGGAACGCCTGACCGAAAAACATTGGGCGCTGCCGCAGAGCGTGTTTTTTCT
GACCTTTAACATTGGCCCGAGCAAATATCGCCTGGTGGGCCGCAAAATTCTGAGCGATCTGTGGATTCAGAACCGCATTCCGGAAATTCA
TTGCAGCTTTGTGAGCGCGGGCATGATTCTGGGCGTGTTTAGCGATCAGCGCATTACCGAAACCTTTGTGAACGTGTTTTATCATCCGCT
GACCAACGGCGAAAAACTGAAAGTGATGAAAGAAATTACCGATCATAAAATTGATAACGTGGAAATTGTGTTTCCGTTTAGCCTGCTGC
GCGAAAGCACCCGCAAAGAACTGTGCAGCCCGAGCGAAGGCTGGGTGGTGGATGATGCGTTTGATGTGATGCTGGGCTGCGGCGAAGAA
CGCATTCGCGCGTATACCATTAAATTTCTGAAAACCCTGAGCAGTCAGAAACTGGTGCTGTTTGATCCGGCGTGCAGCACCGGCGTGTTT
CTGAGCACCCTGAAAAAAAGCTTTCCGGAAAGCTTTACCATTGGCCAAGATCTGAGCAAACAGATGGTGGGCTTTAGCAAAACCCGCGT
GGATGAAGTGCACTGCGGCAACGCGATGGAACCGAAAATTAGCCCGGCGACCGCGGATGTGGTGTTTATTCGCTTTCTGAACAGCGAAG
TGGTGACGAGCGCGGAAGCGGAACACCTGATTGGCGCGCTGCTGCCGACCGTGAAAAAAGGCGGCTATATGGTGATTTTTGGCCACACC
CCGGTGCTGCTGAGCAGCGCGAACTTTAGCCTGATGGGCAACTTTGTGGTGAAACACTGCATTGGCATTAGCGATGATACGAGCGGCATT
TTTCAGTATTATACCCTGCAGCGTCAG

7.2. Plasmid maps

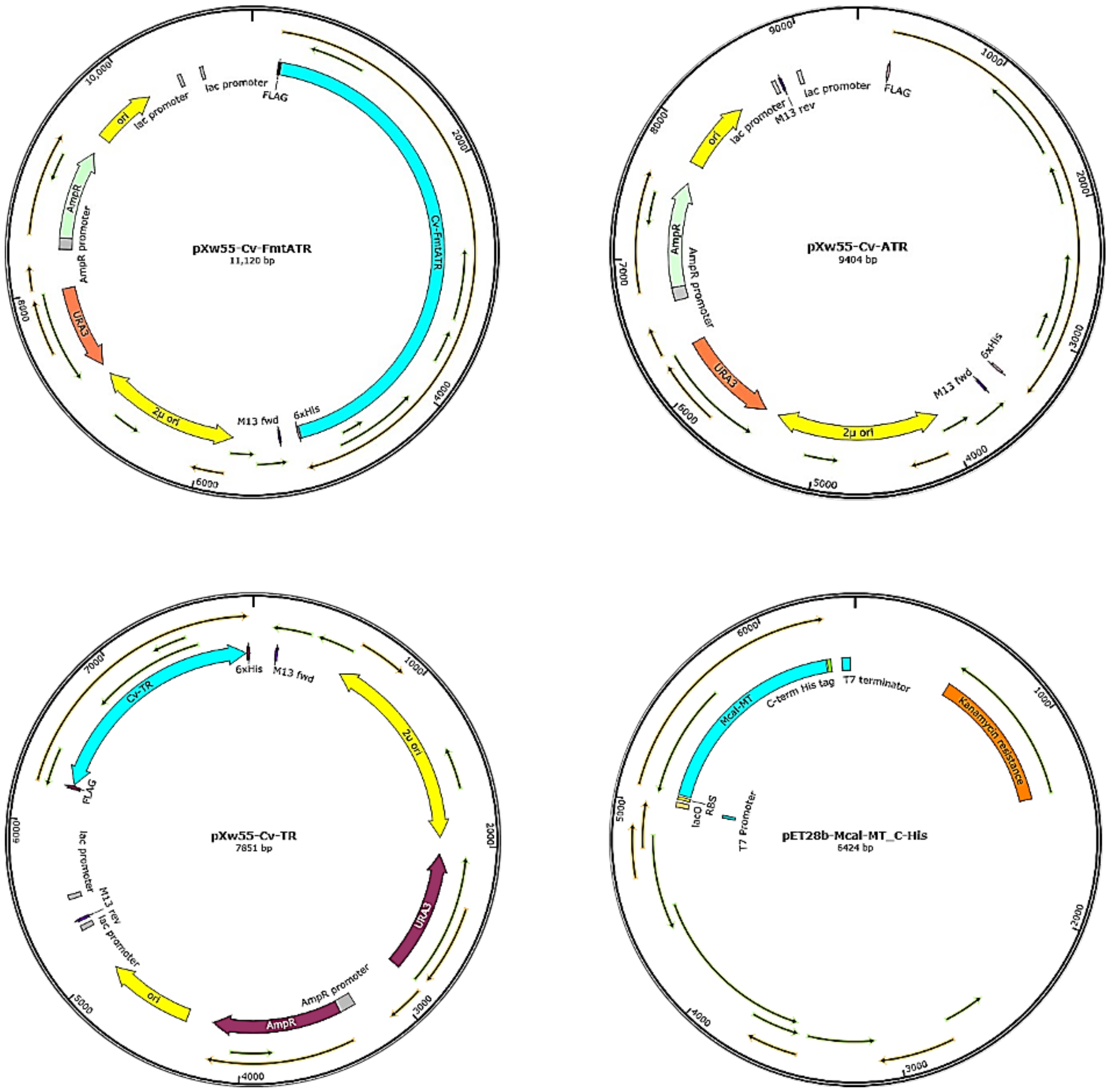


**Figure S63**. Maps of constructs used in this study. *Cvfmt-atr* and *Cv-tr* fragment PCR amplified from *Cvfmt-atr* were cloned into pXw55 backbone using SpeI and PmlI sites. pXw55-CvATR was generated from pXw55-CvFmtATR by site-directed mutagenesis. *Mcal-act* was cloned into pET28b backbone using XhoI and NcoI sites.

7.3. Standard curves

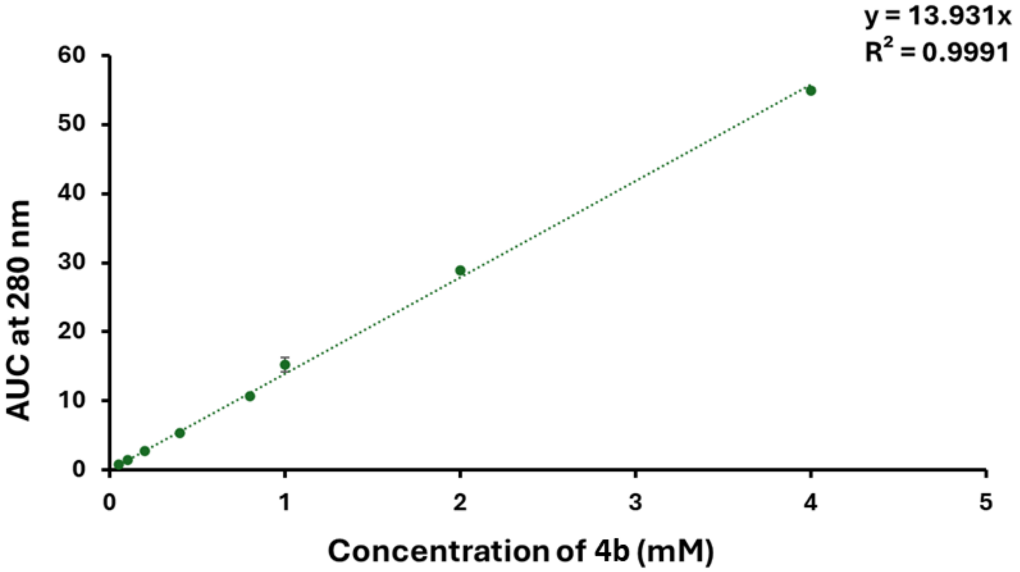


**Figure S64**. *N*-Boc-L-tyrosinol (**4b**) standard curve.
